# Spatial omics representation and functional tissue module inference using graph Fourier transform

**DOI:** 10.1101/2022.12.10.519929

**Authors:** Yuzhou Chang, Jixin Liu, Anjun Ma, Sizun Jiang, Jordan Krull, Yao Yu Yeo, Yang Liu, Scott J. Rodig, Dan H. Barouch, Rong Fan, Dong Xu, Garry Nolan, Zihai Li, Bingqiang Liu, Qin Ma

## Abstract

Tissue module (TM) is a spatially organized tissue region and executes specialized biological functions, recurring and varying at different tissue sites. However, the computational identification of TMs poses challenges due to their convoluted biological functions, poorly-defined molecular features, and varying spatially organized patterns. Here, we present a hypothesis-free graph Fourier transform model, SpaGFT, to represent spatially organized features using the Fourier coefficients, leading to an accurate representation of spatially variable genes and proteins and the characterization of TM at a fast computational speed. We implemented sequencing-based and imaging-based spatial transcriptomics, spatial-CITE-seq, and spatial proteomics to identify spatially variable genes and proteins, define TM identities, and infer convoluted functions among TMs in mouse brains and human lymph nodes. We collected a human tonsil sample and performed CODEX to accurately demonstrate molecular and cellular variability within the secondary follicle structure. The superior accuracy, scalability, and interpretability of SpaGFT indicate that it is an effective representation of spatially-resolved omics data and an essential tool for bringing new insights into molecular tissue biology.

## Introduction

Spatially-resolved omics data, including transcriptome^1–4^, proteomics^5,6^, spatial-CITE-seq^7^, and spatial ATAC-seq^8^, enable a profound understanding of cellular organizations and interactions within a tissue of interest^9^. Among different spatial omics technologies, spatially-resolved transcriptomics (SRT) was selected as the Method of Year 2020^10^, and can simultaneously measure gene expression and spatial locations of cells/spots. It has been widely applied to empower the investigation of tissue functions or cell phenotype at a cellular/subcellular level^5,11^. Various studies were conducted to investigate spatially variable genes (SVGs)^12^, spatial domains^13^, and cell-cell communications^14^ from SRT data. Specifically, a spatial domain is a segment of tissue architecture, referring to a coherent region in gene expression and histology in well-structured tissues^15^. Spatial domains identified by SpaGCN^13^ and BayesSpace^16^ enable a better understanding of cell organization and how cells interact with other domains to perform their functions. However, spatially-organized regions could be incoherent and discontinuous, such as the germinal center (GC) in the lymph node^3^, where the spatial domain concept is stretched.

Hence, tissue module (TM) has been proposed to investigate molecular tissue biology based on molecule compositions and functions in various tissues^3,5^. It was defined as a spatially-organized region formed from cell types co-regulated by recruitment factors, such as cytokines^5^ (**Terminology Boxes 1 and 2**). Then, TM was interpreted as a conserved functional niche across multiple samples with recurrent cellular communities to execute the corresponding functions^3^. Identifying TM empowers the investigation of spatially-organized functional niches, the underlying cell-cell communications mechanism, and the understanding of convoluted molecular functions from multiple TMs. To our knowledge, there is no rigorous computational formulation of TM identification, mainly because: (*i*) the molecular signatures (e.g., SVGs) of a TM are not fully elucidated; (*ii*) TMs exhibit a wide range of length scales and irregular boundaries; (*iii*) there is no repository for TM spatial patterns; (*iv*) the relevant feature crosstalk of convoluted TMs has not yet been well-defined^3^. For example, the GC in the lymph node is a typical TM for B cell maturation. However, the accurate inference of GC is still under-investigated since GC involves multiple cell types and initiates a series of immune reactions for producing long-lived memory B cells and plasma cells^17^.

TM identification can be formulated as an approximate *k*-bandlimited graph signal inference problem with the theoretical foundation in **Supplementary Note 1**^18–20^. Here we introduce a hypothesis-free and efficient graph Fourier transform framework for TM identification, SpaGFT, based on a novel data representation of the molecular signatures embedded in the TM. Specifically, one molecular signature of a TM is a smooth signal and can be represented as the linear combination of *k* low-frequency Fourier mode (FMs), where a low-frequency FM contributes to a slow and smooth graph signal variation leading to a more organized (rather than random) spatial pattern^20^. For example, if a gene is an SVG, its low-frequency FMs usually have more significant contributions than high-frequency FMs in our investigations. Herein, SVG identification can be converted to recognizing *k* low-frequency FMs and their corresponding Fourier coefficient (FCs). This transform does not rely on any predefined spatial patterns^20^, hence ensuring generalizability in identifying both well-defined and irregular SVG patterns. Intuitively, the recurrent SVG patterns are approximate representations of the corresponding TM, like multi-views of the same TM, and provide a natural avenue to TM identification. Using this novel graph representation of spatial omics data, the SpaGFT framework can identify TMs not only from gene expression but also from signals isolated from protein expression, chromatin accessibility, host-residue microbiome abundance, and histology information (**Fig. 1a**).

**Fig. 1.**
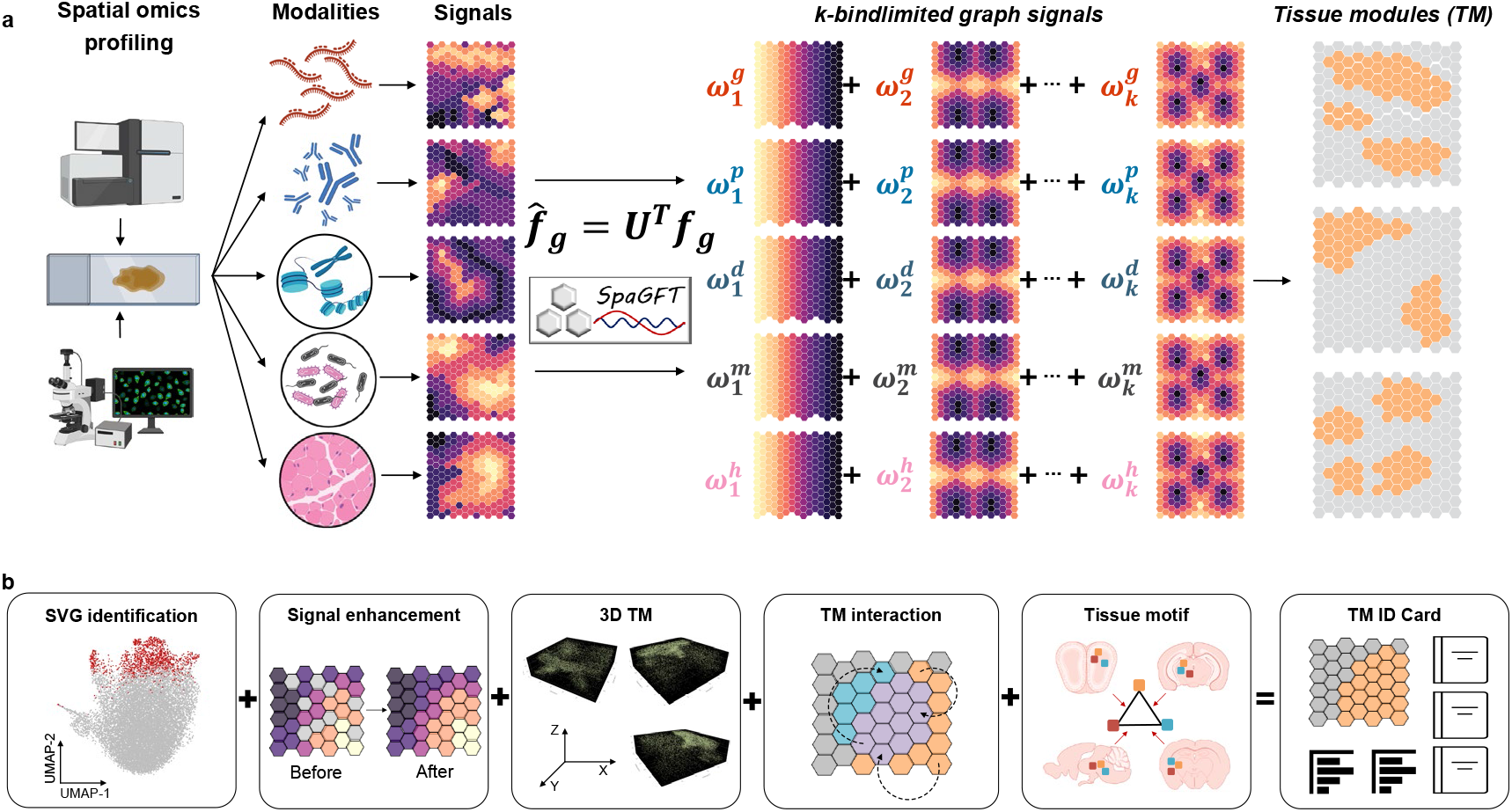
Conceptual workflow and enabled functions of SpaGFT. **a.**Schematic diagram of tissue identification. A tissue or organ sample can be profiled by sequencing-based (Visium), imaging-based spatial transcriptomics (e.g., MERFISH), spatial epigenomics (e.g., spatial-CITE-seq), histology (e.g., H&E staining), and spatial proteomics (e.g., CODEX). Each modality will produce signals to support functional regions that potentially form tissue modules (TM). Specifically, activated TM will execute biological functions and tend to form spatially organized patterns. Such spatially organized patterns can be regarded as an approximate *k*-bandlimited graph signal. By implementing the graph Fourier transform, the k-bandlimited graph signal can be represented by a linear combination of a series of first *k* low-frequency Fourier Modes (FM). The weights of *k* low-frequency Fourier Modes are defined as Fourier coefficients (FC) and used for characterizing TMs. **b.**The figure demonstrates the functionalities of SpaGFT, which include SVG identification, signal enhancement, 3D TM construction, TM interaction investigation, and tissue motif inference. Eventually, TM’s key information is displayed on the TM ID Card.

To provide a comprehensive characterization of TMs, SpaGFT generates an ID Card for each TM, showcasing the spatial organization of a TM, corresponding SVGs (other molecular signatures if available), gene signal enhancement, interactions among different TMs, and the high-level tissue motif structures (**Fig. 1b**). To test the performance and efficiency of the SpaGFT in elucidating SVGs of a TM, 31 public datasets were used to compare between SpaGFT and other state-of-the-art tools. We further apply three cases to showcase SpaGFT applications on mouse brain and human lymph node and tonsil regarding TM interpretation. In the mouse brain case, we defined and interpreted TMs across multiple samples based on Visium and MERFISH. For the lymph node case, we investigated relations and their convoluted functions of overlapped TMs among B follicle, T cell zone, and GC based on Visium and spatial-CITE-seq^21^. For the tonsil case, we generated new CODEX data to showcase a variety of secondary follicles regarding morphological patterns, molecular signatures, and cellular components at the pixel level.

## SpaGFT identifies SVGs accurately from SRT data in support of TM prediction

Accurate SVG identification is a prerequisite to predicting TMs from SRT data, with hypothesis-driven statistical frameworks and graph neural networks developed recently^3,22^. Although these methods demonstrate performance, are equipped with rigorous statistical evaluation, and provide valuable biological insights, they have two main limitations: (*i*) they can identify well-defined patterns (e.g., radial hotspot, curve belt, or gradient streak), but exhibit a lesser detection performance for irregular patterns, such as GC in the lymph node^3^; and (*ii*) the prediction accuracy decreases if a tool significantly improves the efficiency^23^ for datasets with a large number of spots/cells^24,25^. SpaGFT represents the gene expression (defined as graph signals *f_g_*)in a Fourier space (**Fig. 2a**) through a linear combination of FMs *U* (**Fig. 2b**). The FMs are orthogonal and determined by spatial graph topology in increasing order of their frequency, with FM1 having the lowest frequency (**Supplementary Fig. 1a**). Intuitively, the expression pattern of an SVG should be recovered by a set of low-frequency FMs, other than high-frequency FMs (**Supplementary Figs. 1b-c**). SpaGFT calculates a Fourier coefficient 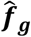 (FC) for each FM, and the sum of low-frequency FCs is defined as the GFTscore for evaluating contribution of low-frequency FMs. Overall, a gene is identified as an SVG in SpaGFT (**Supplementary Note 1**) if (*i*) its GFTscore is greater than the inflection point determined using the Kneedle algorithm from the GFTscore distribution and (*ii*) it has significantly higher FCs of low-frequency FMs than those of high-frequency FMs (adjusted *p*-value < 0.05 in a Wilcoxon test).

**Fig. 2.**
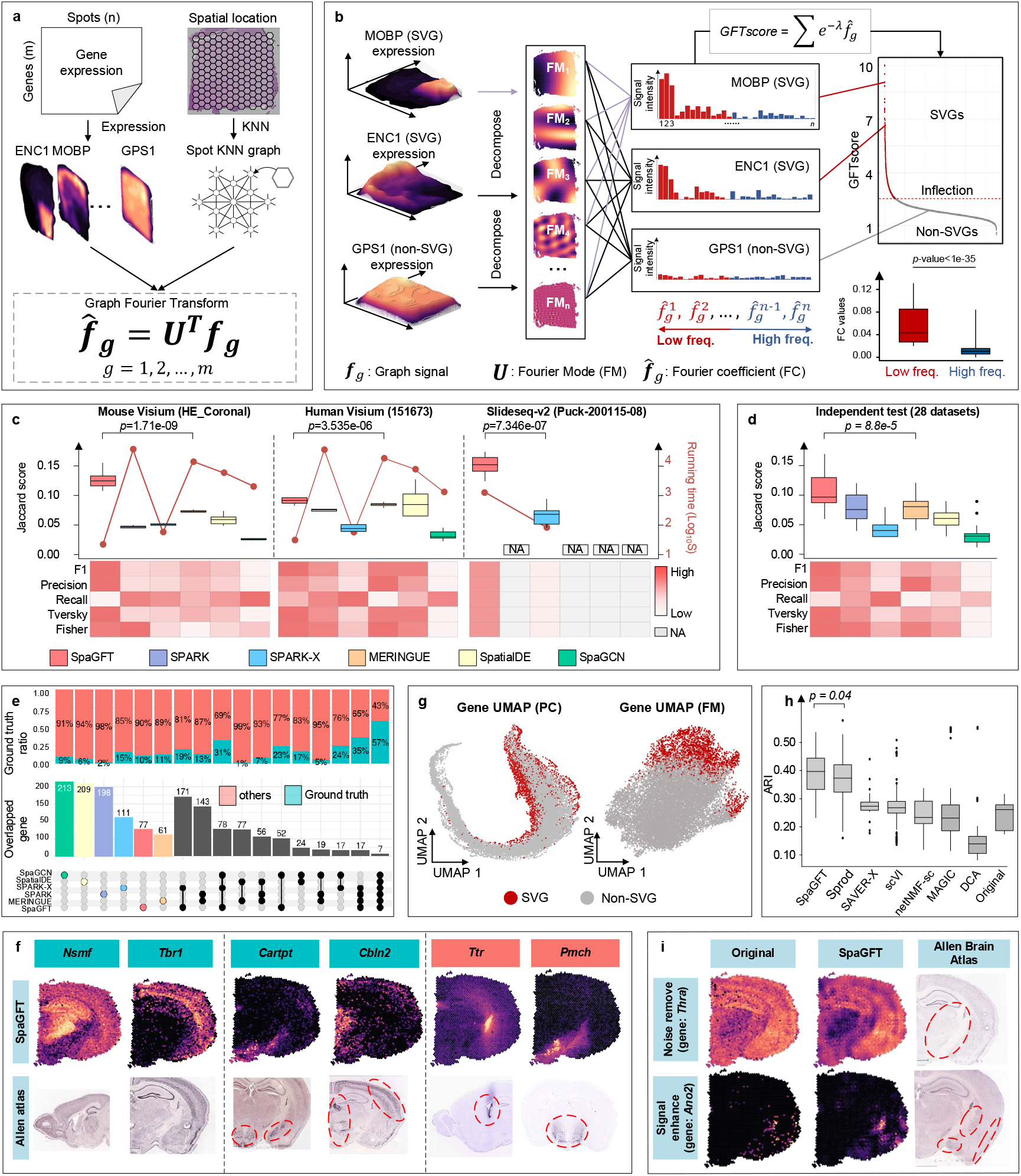
Overview of SpaGFT framework design and SVG identification performance comparison. **a.**SpaGFT considers a gene-spot expression count matrix (*m × n*) and spatial locations as input data, with *ENC1, MOBP*, and *GPS1* listed as examples. A KNN spot graph is generated by calculating the Euclidean distance among spots based on spatial locations between any two spots. Combining gene expressions and spot KNN graph, the graph signal *f_g_* of gene *g* can be projected to a series of FM *U* and transformed into a frequency signal 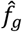 using a graph Fourier transform. **b.**Two known SVGs (*MOBP* and *ENC1*) and one non-SVG (*GPS1*) are shown as examples. A gene can be decomposed into multiple FMs (a series of periodic signals with gradually faded patterns) and corresponding frequency signals. The FMs can be separated into low-frequency (red) and high-frequency (blue) domains. For each gene, a *GFTscore* was designed to measure the FCs in the low-frequency domain quantitatively. Boxplot in the right-bottom corner shows that the low-frequency FCs of 1,902 SVGs from sample 151673 are significantly higher than high-frequency FCs (with a *p*-value<1e^-35^ in the Wilcoxon rank-sum test). **c.**The SVG prediction evaluation was compared to five benchmarking tools in terms of reference-based evaluation methods, the Jaccard Index, Tversky Index, Fisher Statistic, Precision, Recall, and F1 score, and reference-free evaluation methods, such as Moran’s I and Geary’s C. The running time (seconds with log transformation) of each tool is represented as red lines. In addition, the F1 score, precision, recall, Tversky index, and odds ratio of Fisher’s exact test on all parameter combinations for each tool is shown as heatmaps. The statistical test method was the same as panel b to calculate the *p*-value for the highest two tools (i.e., parameter combination number N = 16 and N = 53 for SpaGFT and MERINGUE in HE_coronal data; N = 16 and N = 54 for SpaGFT and MERINGUE in 151673 data; N = 16 and N = 18 for SpaGFT and SPARK-X in Puck-200115-08 data). Particularly, regarding larger datasets such as Slide-seqV2, the statistical methods might not be able to identify SVG in a reasonable time, showing NA in Fig. 2c. **d.**The parameter combination showing the highest median Jaccard scores among all three benchmark datasets was selected as the default parameter for each tool. Using such a parameter selection, the SVG prediction performance of SpaGFT on additional 28 independent datasets was compared to those of the five benchmark tools. The black line in each box indicates the median Jaccard score of all 28 datasets. The statistical test method was the same as panel b to calculate the *p*-values for the highest two tools (N =28). **e.**Shared SVGs among six computational tools using the HE-coronal dataset. The bottom upset plot indicates uniquely identified SVGs and overlapped SVGs, and the bar plot in the middle shows the corresponding SVG number. The ground truth ratio panel (top) demonstrates the proportion of ground-truth SVG among shared SVGs across six tools. **f**. SVG examples that all tools can identify (validated ground truth, left panel), uniquely identified by SpaGFT (validated ground truth, middle panel), and uniquely identified by SpaGFT (not validated, right panel). If the coronal plane of ISH validation data was unavailable, the sagittal plane was used. **g.**Comparison of the UMAPs obtained using the top 207 PCs (left) and the top 207 FCs (right) of the Mouse Visium data (HE-coronal, 2,702 spots). PCA dimensions were generated directly from the gene-spot expression matrix using Scanpy. Red dots indicate the 2,157 SVGs identified by SpaGFT using default settings, whereas the grey color suggests non-SVGs. Red circles in the ISH data indicate the expression region of the mouse brain. **h**. Boxplot showcases the performance of SVG signal enhancement. The y-axis is the ARI value, and the y-axis is the imputation tool name. The statistical test method was the same as panel b to calculate the *p*-values for the highest two tools (N = 144). **i.**The spatial map shows the gene expression before and after enhancement and the IHC image from Allen Brain Atlas. Abbreviation: spatial variable gene (SVG); principle component (PC).

We assessed the performance of SVG identification using 31 public SRT datasets from human and mouse brains (**Supplementary Table 1**). As no golden-standard SVG database is available, we collected 849 SVG candidates from five existing studies^15,26–29^, and 458 of them were used as curated benchmarking SVGs based on cross-validation with the In Situ Hybridization (ISH) database of Allen Brain Atlas (**Supplementary Tables 2 and 3**, **Methods**). The SVG prediction performance of SpaGFT was compared with SPARK^30^, SPARK-X^23^, MERINGUE^31^, SpatialDE^32^, and SpaGCN^13^, in terms of six reference-based and two reference-free metrics. The grid-search of parameter combinations was conducted on three high-quality brain datasets to evaluate each tool’s performance, in which SpaGFT shows the highest median and peak scores (**Fig. 2c**, **Supplementary Fig. 2**, and **Supplementary Table 4**). In addition, the computational speed of SpaGFT was two-fold faster than that of SPARK-X and hundreds-fold faster than those of the other four tools on the two Visium datasets (**Supplementary Table 5**). Although SpaGFT exhibited a slower performance than SPARK-X on the Slide-seqV2 dataset, it showed a remarkably enhanced SVG prediction performance compared to SPARK-X (**Supplementary Table 5**). We then performed an independent test on additional 28 datasets using the parameter combination with the highest median Jaccard Index among three datasets from the grid-search test. The results revealed that SpaGFT promised supreme performance among the investigated tools based on the evaluation metrics (**Fig. 2d** and **Supplementary Table 6**).

Within the top 500 SVGs from each of the above six tools, SpaGFT identifies SVGs shared with other tools and also unique SVGs that are validated as ground true (**Fig. 2e** and **Supplementary Table 7**). For example, *Nsmf* and *Tbr1* were identified by all six tools and showed clear structures of the hippocampus, cortical region, and cerebral cortex (**Fig. 2f**). On the other hand*, Cartpt, Cbln2, Ttr*, and *Pmch* were uniquely identified by SpaGFT and showed key functions in the brain, such as *Cartpt* participating dopamine metabolism (**Supplementary Fig. 3** and **Annotation 1 of Supplementary Note 2**). These benchmarking results suggest that SpaGFT is robust and accurate in identifying SVGs from SRT data. SpaGFT takes advantage of FM representation of gene expression patterns in SVG identification, and the SVGs identified by SpaGFT were distinguishably separated from non-SVGs on the FM-based UMAP with a clear boundary, whereas SVGs were irregularly distributed on the principal component-based gene UMAP (**Fig. 2g**).

Notably, due to the dropout and other technical issues^33^, some SVGs retain a low expression or display noisy signals that may be hard to recognize. SpaGFT offers the SVG enhancement function to overcome the limitation, where the FCs of low-frequency FMs will be enhanced and those of high-frequency FMs will be diminished. Then it recovers the SVG spatial expression with an enhanced magnitude via an inverse graph Fourier transform (iGFT) (**Supplementary Fig. 4**). To test the performance of gene expression imputation and enhancement, we used 16 human brain SRT datasets with well-annotated spatial domains^15,34^. As a result, SpaGFT outperformed other gene enhancement tools, including Sprod^35^, SAVER-X, scVI, netNMF-sc, MAGIC, and DCA^36–40^ (**Fig. 2h and Supplementary Table 8**). For example, SpaGFT removed the noisy background for the neural cell broadly expressed gene, *Thra*, and enhanced the low-expression gene from the amygdala coronal view, such as *Ano2* (**Fig. 2i** and **Annotation 2 of Supplementary Note 2**).

## SpaGFT characterizes TM and tissue motifs in the mouse brain

As FC is a low-dimensional representation of SVG to describe spatially organized patterns quantitatively, they are ideal features for detecting SVG clusters and inferring TMs (**Supplementary Note 1**). Therefore, we implemented an optimization framework in SpaGFT to identify TMs by determining SVG clusters that share similar spatially organized patterns (**Supplementary Fig. 5a**). Taking the HE-coronal data as an example, 12 SVG clusters (**Fig. 3a and Supplementary Table 9**) were identified from a total of 2,118 SVGs by minimizing overlapped spots among identified TMs (**Supplementary Fig. 5b**), so that each TM has its representative biological functions. To characterize a TM, an ID Card is produced to showcase a TM spatial map, transformed FCs, enhanced SVGs, underlying biological pathways, overlapped regions with other TMs, and cell-type composition.

**Fig. 3.**
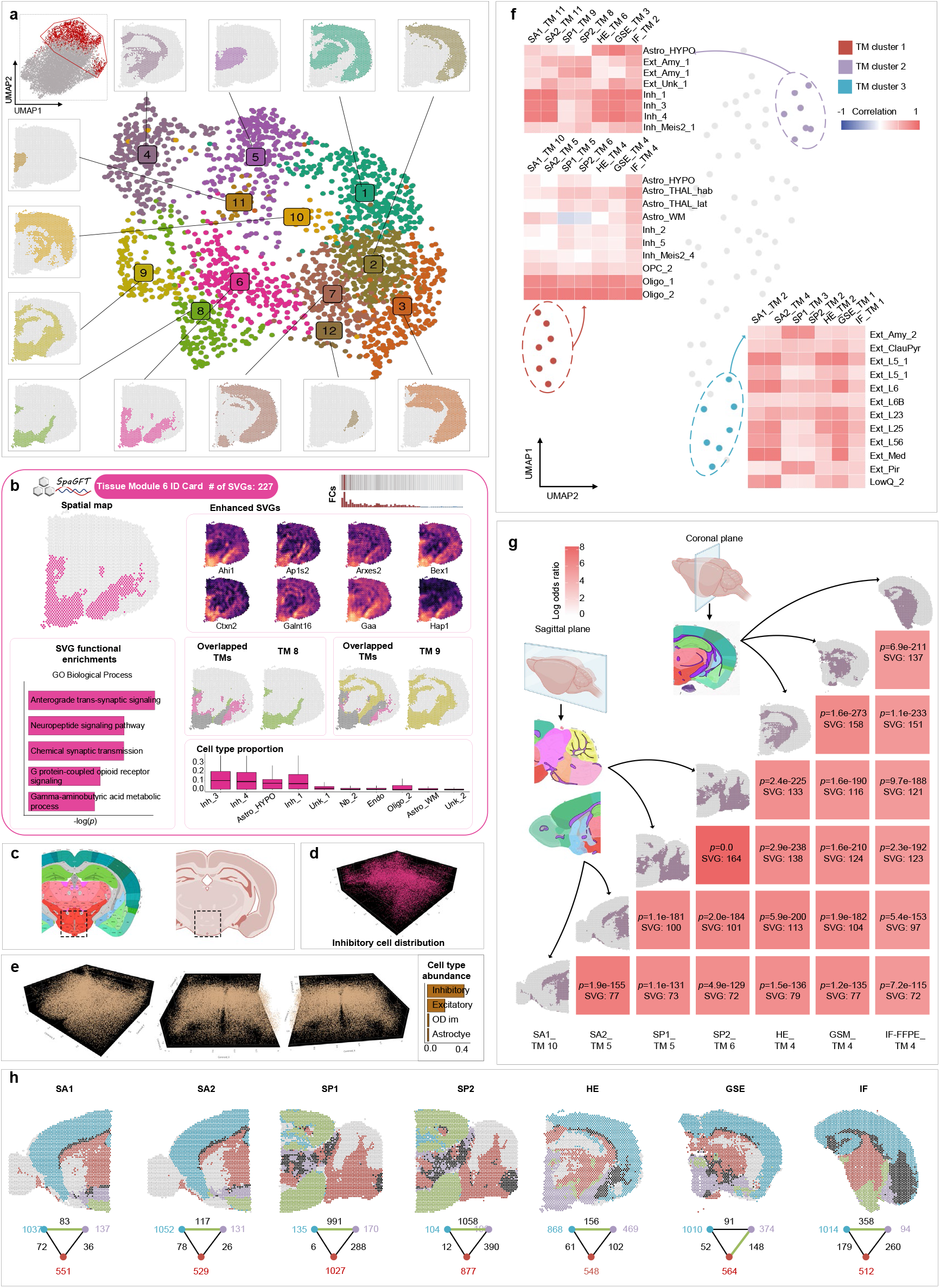
TM and tissue motif identification in mouse brains. **a.**All 2,118 SVGs identified in the Mouse Visium data (HE-coronal) were grouped into 12 clusters, representing 12 TMs. SVGs from Fig. 2g are shown in the left-up corner (red: SVGs, grey: non-SVGs). **b**. An ID Card is created to display the fundamental information about each TM. The spatial map shows the shape of TM 6 with 227 SVGs. The corresponding FC is displayed on the top right. The spatial maps of the enhanced SVGs are shown on the right, including *Ahi1, Ap1s2, Arxes2, Bex1, Ctxn2, Halnt16, Gaa*, and *Hap1*. Functional enrichment tests of the 227 SVGs are performed on Biological Process 2021 by Enrichr to provide insights on the functional and regulation information enriched in this TM. Overlapped TMs section shows that TMs 8 and 9 intersect with TM 6. The cell type proportion section shows the top enriched cell types. **c.**Twelve consecutive samples were generated from the mouse hypothalamic preoptic region using MERFISH data. **d.**3D spatial map shows one of the cell types distributions (inhibitory cell in this case). **e.**A 3D tissue module is displayed from three views. **f**. Results of clustering 76 TMs identified from seven samples. Three heatmaps show the binarized TM-cell type matrix, indicating consistent cell types shared within three TM clusters, where red means cell type highly correlated with the corresponding TM, and blue means a negative correlation. **g**. Interpretation of TMs from multiple samples with similar SVG components. Seven samples are used to demonstrate the commonality of SVG-similar TMs in multiple samples. The heatmap color indicates the log-odds ratio of the Fisher exact test. The FDR adjusted *p*-value and the number of shared SVGs between the two samples are shown on the heatmap. White matter anatomical structure is derived from Allen Brain Atlas and indicated by purple. **h**. The figure demonstrates two conserved tissue motifs shared by multiple samples. The spatial map indicates spot localization where a spot is colored according to TM cluster assignment (brown for TM cluster 1; purple for TM cluster 2; and blue TM cluster 3). The tissue motif below each spatial map demonstrates the colocalization of TM clusters. A node represents one specific TM cluster, and the value of the node means the number of spots in the corresponding sample of a TM cluster. An edge will be added if overlapped spots exist between the two nodes. The weight of an edge is the number of overlapped spots between the two nodes. The green edge denotes the edge with the largest weight in one tissue motif.

As a demonstration, we showcased the ID Card for TM 6 (see other in **Supplementary Data 1**), which displayed a discontinuous pattern distributed in the striatum, thalamus, and hypothalamus regions (**Fig. 3b**). The eight enhanced SVGs were selected to support the spatial expression pattern of TM 6. The enrichment of ontologies (**Supplementary Fig. 5c**) was also performed for TM-associated SVGs to elucidate the underlying functions of the TM 6, and the results indicated the function of regulatory neurons for the hypothalamus and amygdala (more details in **Annotation 3 of Supplementary Note 2**). In addition, we performed cell type deconvolution analysis based on 59 mouse brain cell types using cell2location (**Supplementary Table 10**)^33,41^ to illustrate cellular components. As a result, TM 6 contained a high abundance of inhibitory neuron and astrocyte subtypes, whose distribution pattern (**Supplementary Fig. 5d** and **Annotation 3 of Supplementary Note 2**) and functions also supported the anatomical structures of TM 6. In addition to 2D TM characterization, SpaGFT can infer 3D TMs. Based on the 12 consecutive slides of mouse preoptic hypothalamus region^1^ (**Fig. 3c**) profiled by MERFISH, we identified nine 3D TMs using 3D coordinates to construct the spatial graph for the SpaGFT framework. For example, inhibitory neuron cells were visualized in **Fig. 3d**, and TM 2 in **Fig. 3e** had a high abundance of such a cell type (see more in **Supplementary Data 2**).

As the 3D TM can be characterized using consecutive sample slides, we hypothesized that if one TM’s 3D structure was irregular and complex, multiple 2D views of this TM would be represented by a group of SVG components with synergetic functions, regardless of sampling strategies (e.g., different anatomical plane) or sources. To verify the hypothesis, seven mouse brain samples were collected from the Visium website and one independent study^42,43^, including sagittal and coronal planes (**Supplementary Fig. 6a**). SpaGFT identified 76 TMs among the seven samples (**Supplementary Table 11**). The 76 TMs were grouped into eight clusters based on their associated SVGs (**Supplementary Fig. 6b** and **Methods**). As a result, we focused on the three colored TM clusters (TM clusters 1, 2, and 3 in **Fig. 3f**), each of which contained conserved TMs across seven samples (**Supplementary Table 12**). First, the cell2location results show that TM clusters 1, 2, and 3 were highly correlated with non-neuronal cells, inhibitory neurons, and excitatory neurons, respectively (**Fig. 3f**). TM cluster 1 was enriched with oligodendrocytes and also in agreement with the white matter anatomical structure from the Allen Brain Atlas (**Fig. 3g**). This was also supported by white matter signature genes (e.g., *Mbp;* **Supplementary Table 13**), which shows a high agreement across all seven samples^15^ (**Fig. 3g**). TM clusters 2 and 3 were highly correlated with the hypothalamus and partial cerebrum regions, respectively (**Supplementary Fig. 6c**). We concluded that conserved TMs forming one TM cluster typically contained conserved cell types and SVGs regardless of sample resources.

Based on the discovery of conserved TMs across multi-samples, we further reason that if one organ or tissue has multiple conserved TMs, those TMs tend to form a high-order functional structure to organize and support the main functions. This high-order functional structure is defined as a tissue motif introduced earlier by the previous study^11^. Therefore, we defined TM clusters 1, 2, and 3 as one tissue motif (**Fig. 3h**). Based on the spot numbers assigned by the TM clusters (triangle graph in **Fig. 3h**), we found that the tissue motif co-occurred and was conserved in all seven samples and showed a collaborative neuronal circuit activity (details in **Annotation 4 of Supplementary Note 2**). From multiple anatomical views of mouse brain samples, our results demonstrate that SpaGFT provided a novel gene-centric perspective for investigating and interpreting conserved TMs among multiple samples and their convolution^33^.

## SpaGFT identifies short-length scale TMs and their crosstalks in human lymph node

Lymph node is a secondary lymphoid organ containing several recurrent short-length scaled regions, such as T cell zones, B follicles, and GCs^3^. We applied SpaGFT to Visium data of human tonsil^43^ to investigate the organization of functional regions and their crosstalks. SpaGFT identified 1,346 SVGs and characterized nine TMs (**Fig. 4a** and **Supplementary Table 14**). To recognize the T cell zone, B follicle, and GC, we first used cell2location^33,41^ to determine the cell proportion (**Supplementary Fig. 7a** and **Supplementary Table 15**) for the nine TMs and investigate function enrichment (**Supplementary Figs. 7b-d**) for three selected TMs. Based on the molecular, cellular, and functional signatures of three regions^41^, we found that TMs 3, 5, and 7 (**Fig. 4b**) were associated with the T cell zone, GC, and B follicle, respectively (**Annotation 5 of Supplementary Note 2**).

**Fig. 4.**
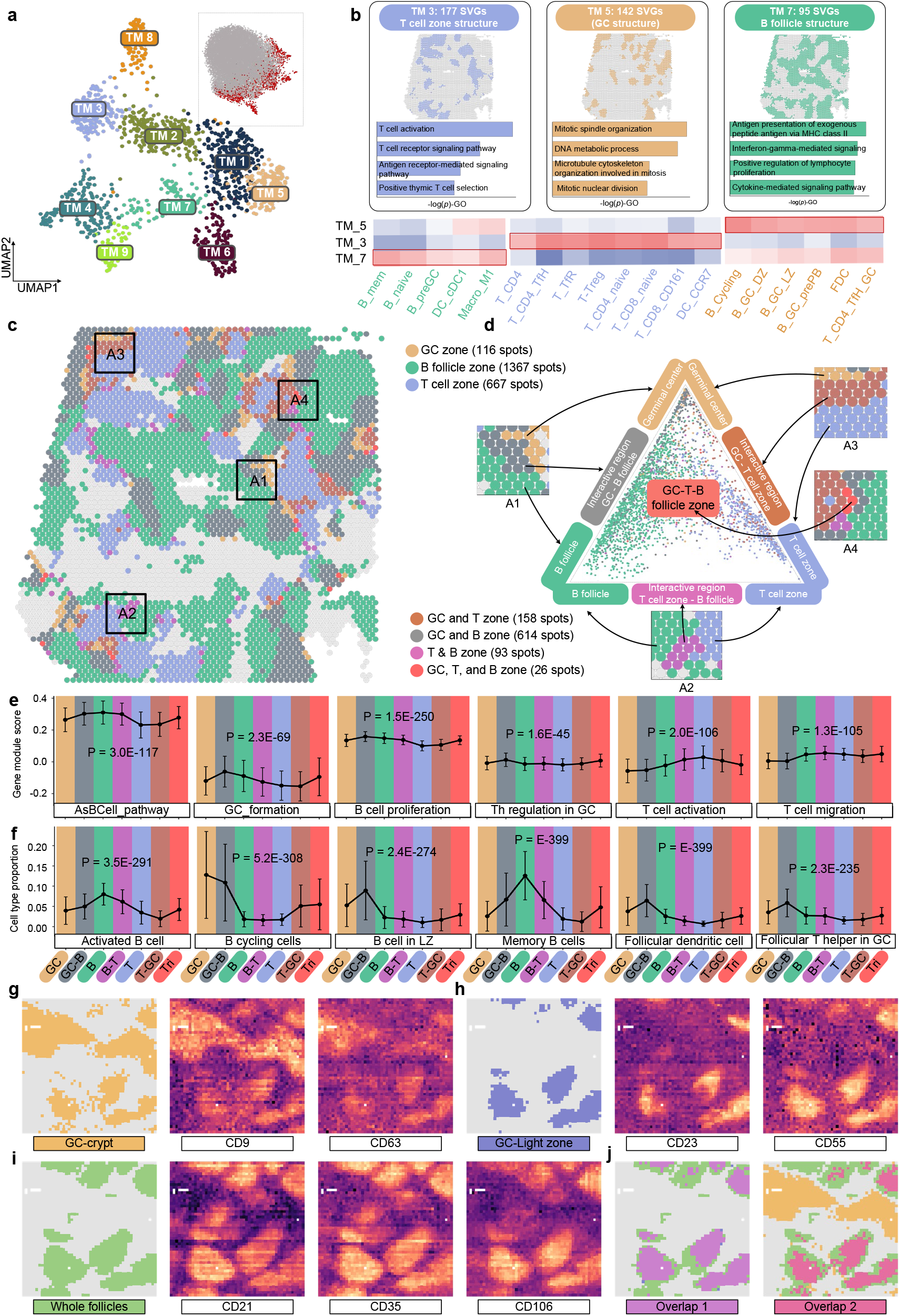
SpaGFT revealed functional and cellular variation of the T cell zone, GC, and B follicle and their interactive regions. **a.**UMAP visualization of nine TMs from the human lymph node. Each dot represented SVGs. Upright UMAP showed SVGs in red color and non-SVG in gray color. **b.**TMs 3, 5, and 7 were highly associated with the T cell zone, GC, and B follicle cell components based on molecular and functional signatures. The heatmap visualized the TM-cell type correlation matrix. **c.**The spatial map overlaid three TMs and displayed the overlapped spot and unique spots. As different colors corresponded to spots, we selected four areas to showcase the region-to-region interaction. A1 showcased GC, GC-B interaction region, and B follicle. A2 showcased the B follicle, B-T interaction region, and T cell zone. A3 showcased the GC, GC-T interaction zone, and T cell zone. A4 displayed a B-GC-T interaction zone. **d**. The barycentric coordinate plot shows cell-type components and the abundance of spots in interactive and functional regions. If the spot is closer to the vertical of the equilateral triangle, the cell type composition of the spot tends to be signature cell types of the functional region. The spots were colored by functional region and interactive region categories. **e** and **f**. The three plots displayed changes in enriched functions and cell type components across seven regions (GC, GC-B, B, B-T, T, T-GC, T-GC-B), respectively. The P-value was calculated using one-way ANOVA to test the differences among the means of seven regions. **g-i**, Three GC-relevant TMs and corresponding spatially variable proteins were identified. **j**. Two overlapped regions included (1) whole follicle and GC-light zone and (2) whole follicle and GC-crypt. Abbreviately, AsBcell_pathway stands for antigen-dependent B cell activation. Th is helper T cell in **e**.

In contrast to spatial domain detection tools, SpaGFT does not distinguish a rigorous boundary to segment tissue. Instead, SpaGFT allows the occurrence of the overlapped regions of TMs and contributes to understanding the functional coherence and collaboration among different TMs. Therefore, we projected TMs 3, 5, and 7 on the spatial map and observed that they were spatially close to or overlapped with each other, indicating potential regions of polyfunctionality among these three TMs (four regions were selected to visualize unique and overlapped TMs regions from **Fig. 4c**). To reveal the crosstalk among these three regions, we projected spots (assigned to all three regions) to the Barycentric coordinates (the equilateral triangle in **Fig. 4d**) which displays relations and abundance of the unique and overlapped regions regarding cell type components^11^. As a result (**Supplementary Table 16**), we found 614 spots overlapped with B follicle and GC, 158 spots overlapped with GC and T zone, 93 spots overlapped with T zone and B follicle, and 26 spots overlapped across three TMs. Furthermore, we hypothesized that the spots from the overlapped region would vary in functions and cell components to support the polyfunctionality of these regions. Hence, we investigated the changes in enriched functions (**Supplementary Table 17**) and cell types (**Supplementary Fig. 8**) across seven regions (i.e., GC, GC-B, B, B-T, T, T-GC, and T-B-GC). As a result, lymph node-relevant pathways and cell types are significantly varied across those regions, such as B and T cell activity and functions (**Figs. 4e-f**, **Annotation 6 of Supplementary Note 2**).

As the resolution (~55 *μ*m per spot) of Visium might limit the insight discovery and interpretation, we next implemented SpaGFT on human tonsils profiled by spatial-CITE-seq^7^ (with genes and proteins) to explore the interactive region of finer structures in follicles at a higher resolution (~25 *μ*m per pixel). First, we discovered three TMs for GC-crypt (**Fig. 4g**), GC-light zone (**Fig. 4h**), and clear whole follicle region (**Fig. 4i**) among several other TMs (**Supplementary Table 18**). Second, we overlapped GC-crypt and GC-light zone with the whole follicle region, giving rise to two overlapped regions (**Fig. 4j**). Interestingly, we found a potential B cell migration between follicle and GC-crypt regions which might be promoted by tetraspanins such as CD9 and CD63 on B cell membrane^44^. More biological interpretations of these regions were in **Annotation 7 of Supplementary Note 2**.

## SpaGFT revealed secondary follicle variability based on CODEX data

The results of spatial-CITE-seq in **Fig. 4** showcased that SpaGFT could identify TMs at the protein level. However, although spatial-CITE-seq had 25 *μ*m per pixel size and more than 200 proteins, it is currently limited in its ability to interpret the variability of finer follicle structures and corresponding functions at the cellular level. Hence, we additionally imaged a human tonsil using a 49-plex CODEX panel at a ~0.37 *μ*m per pixel resolution (**Fig. 5a**) to better characterize and interpret follicle variability. Based on the anatomical patterns highlighted by B (e.g., CD20) and T cell (e.g., CD4) protein markers, we selected fields of view (FOV) that would allow for a good representation of the complex tissue structures present in the tonsil (i.e., GC and interfollicular space^45^) while still allowing for variability in follicle structure^46^. We first identified cell types by DeepCell^47^, FlowSOM^48^, and Marker Enrichment modeling^49^ to identify the cell phenotypes present in the data (**Fig. 5b**). Interestingly, we observed that the clear arrangement of T and B cell patterns (e.g., A3, A5, and A6) informed identifiable GC regions within the follicular structure, compared to others (e.g., A4) without clear T and B cell spatial organization (**Fig. 5b**). We hypothesized that A4 might be comprised of multiple follicles unlike A5 and A6; hence, it could represent a more spatially complex FOV.

**Fig.5.**
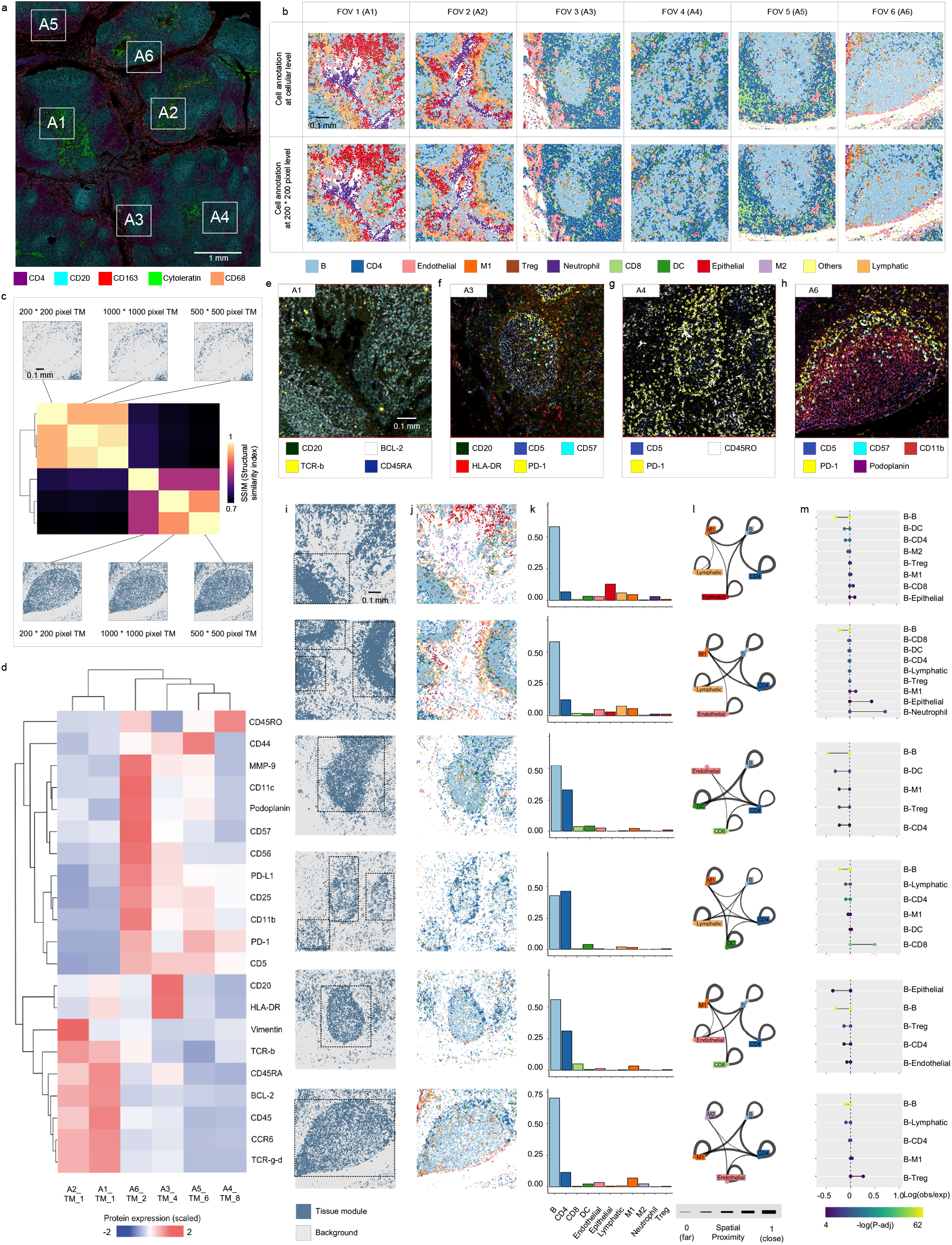
Cellular and molecular variability of secondary follicles using high-resolution CODEX. **a**. A 49-plex CODEX data was generated from human tonsil tissue at a 0.37 μm/pixel resolution. Six FOVs were selected based on their varying tissue microenvironment and cellular organization. **b**. Cell phenotype maps for each of the six FOVs, depicting the cellular composition and organization. **c**. A representative view of the application of TM to characterize the gradient pixel-level images for A6. The heatmap depicts the SSIM score, where a higher score corresponds to a lighter color and greater structural similarity. **d**. A heatmap showcasing the protein expression of each of the six TMs, which were identified as TMs resembling secondary follicles. The values in the heatmap are scaled by z-scores of protein expression. **e-h**. Overlays of CODEX images for TM-associated SVPs for FOVs 1, 3, 4, and 6, respectively. **i**. Spatial maps depicting the patterns of secondary follicle TMs from six FOVs. Dash rectangles indicate the identified follicle regions. Note that panels d to h are ordered by FOV 1, 2, 3, 4, 5, and 6. **j**. Cell phenotype maps of the TMs identified in **i**. **k.**Barplots depicting the cell components of the identified TM in i. The cell type colors were depicted in **b**. **l.**The graph network depicting the spatial proximity of the top 5 abundant cell types in the TM identified in **i**, as calculated by 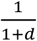, where *d* represents the average distance between any two cell types. **m**. Dumbbell plots indicated significant cell-cell interaction among B cells and others. If the observed distance is significantly smaller than the expected distance, the two cell types tend to be contacted and interact. Line length represents relative distances subtracting the expected distance from the observed distance. The point size was scaled by adjusted *p*-value.

We demonstrated that SpaGFT could effectively characterize TMs regardless of cell arrangement patterns beyond qualitative patterns observed visually. We directly took the raw images as inputs without requiring single-cell segmentation for a scalable approach toward identifying TMs formed from protein signatures within the tissue environment^50^. To verify whether size-reduced CODEX image (**Supplementary Fig. 9a**) would lose the power of characterizing TM, we first used FOV 6 to generate three images with different pixel sizes, including 1,000-by-1,000 pixel image (~0.8 *μ*m per pixel size), 500-by-500 pixel image (~1.6 *μ* m per pixel size), and 200-by-200 pixel image. As a result, although three pixel-level images would produce different spatial variation patterns regarding low- and high-frequency TM (**Supplementary Fig. 9b**), SpaGFT still characterized TMs with consistent patterns (**Supplementary Fig. 9c**).

Subsequently, we calculated the structural similarity score (SSIM) to evaluate pattern similarity among identified TMs quantitatively. Each gradient pixel size image identified six TMs, and those TMs showed pairwise consistency (**Fig. 5c and Supplementary Fig. 9d**), suggesting that 200-by-200 pixel images were sufficient in characterizing TMs.

We next implemented SpaGFT to characterize TMs for the six FOVs with 200-by-200 pixel images and annotated follicles for each FOV based on cell components (**Supplementary Fig. 10**) and molecular signatures (**Supplementary Table 19; Supplementary Figs. 11a-b**). Specifically, TM1 of A1 and TM1 of A2 displayed the morphological features of the partial mantle zone (MZ). Molecularly, we discovered the B cell-specific marker^51^ (CD20) and B cell antiapoptotic factor (BCL-2)^52^ were spatially variable proteins (SVP) for these two TMs of A1 and A2 (**Figs. 5d-e and Supplementary Fig. 11c**). As a result, CD20 confirmed the MZ structure, and the expression of BCL-2 suggested a novel marker on MZ structures^53^. In another case, TM 4 of A3, TM 9 of A4, and TM 4 of A5 displayed GC-specific T cell signatures (**Figs. 5f–5g** and **Supplementary Fig. 11d**) and corresponding molecular features, including PD-1^54^ and CD57^55^, indicating the presence of well-characterized GC-specific T follicular helper cells^56^. For TM 2 of A6, we observed a complex molecular environment, where Podoplanin, CD11c, and CD11b were SVPs and showcased the existence of follicular dendritic cell (FDC)^57^ and GC-centric macrophages^58^ (**Fig. 5h**). In addition to molecular heterogeneity, we further captured their variability in terms of length-scale and morphology (**Fig. 5i**), cell type (**Figs. 5j and k**), cell-cell distance (**Fig. 5l**), and cell-cell interactions (**Fig. 5m**). For example, from the tissue morphology perspective, A3, A4, A5, and A6 captured clear oval shape patterns with different length-scales, but A1 and A2 captured multiple partial MZ patterns (**Fig. 5i**). Although we hardly distinguished morphological pattern of GC in A4 (**Fig. 3b**), SpaGFT still enabled to determine the three small length-scale GC patterns at the molecular level (**Fig. 5i**).

Regarding cellular characteristics, six TMs (i.e., two MZ from A1 and A2; four GCs from A3 to A6) were dominated by B and CD4 T cells with varying proportions (**Figs. 5j-k**; **Supplementary Table 20**). Specifically, MZ TMs from A1 and A2 showed an average of 58% B and 10% CD4 T cells. GC TMs from A3 and A5 with similar length-scale showed an average of 54% B and 32% CD4 T cells. A4 captured three length-scale GC and showcased 43% B and 46% CD4 T cells, while the large-scale GC from A6 contained 70% B and 12% T cells, indicating B and T cell proportions varying in different length-scale GC. We could also infer cell-cell interaction based on distance (**Figs. 5l-m**). In general, all TMs from A1 and A2 show that the observed B-B distance was smaller than the expected distance, which suggests the homogeneous biological process of the significant B-B interaction in the GC region. In addition, cell-cell interaction also shows heterogeneity for these TMs. The interactions between CD4 T cell and B cell were observed in TMs from A1 and A2, showcasing that CD4 T cell might contact and pass through B cell clusters in mantle zone^59^. DC-B and CD4 T-B cell interaction in A3 and A4 suggested the light zone functions for B cell selection^60,61^. Macrophage-B cell interaction in GC in A6 potentially indicated macrophage regulation on B cells (e.g., B cells failed to pass the cell selection process and were taken up by macrophage^62^). Overall, we can apply SpaGFT to low-resolution high-plex spatial proteomics to identify and characterize secondary follicle regions to uncover cellular and molecular variability of secondary follicles that were confirmed at a higher resolution. We also affirmed that TM identified by SpaGFT was not a simple cell aggregation region but reflected both the cellular and regional activity and cell-cell interactions based on spatially orchestrated molecular signatures.

## Discussion

Tissue module is an essential concept for studying tissue function and morphology from a complex tissue or organ. We present SpaGFT as a fast and accurate tool for SVG identification and TM characterization using spatial omics data. SpaGFT introduced a graph Fourier transform framework to analyze spatial omics data, which can transform complex structures on irregular topologies into simple but informative topological features. In this paper, we provide a systematic study to demonstrate SpaGFT’s mathematical rigor, computational design, biological applications, and insights discovery from spatial omics data. The benchmarking results of 31 spatial transcriptome datasets revealed that SpaGFT achieved superior SVG detection performances compared to existing tools, indicating that the FC can effectively capture spatially organized gene expression signals and distinguish SVGs and non-SVGs. In addition, TMs were confirmed to maintain coherence and convoluted biological functions with recurrent cell communities. Moreover, three case studies provided biological insights from the TM interpretation. We also demonstrated the capability of identifying TMs from diverse technologies, tissue motifs across multi-sample, the crosstalk among TMs, and the molecular and cellular variability of secondary follicles.

We also expect that SpaGFT can be potentially used for other spatial multi-omics data harmonization and integration by discovering conserved spatial FCs patterns of metabolic, proteomic, morphogenetic, and epigenetics regardless of healthy and pathological states. Meanwhile, there is an increasing need for building connections between spatial spots using multi-omics at the single-cell level. Based on the SpaGFT framework, FC is a reliable representation of spatially organized spatial-omics features and can be used as a regularizer to constrain alignment between spot and single cells within\across TMs. Such an alignment can provide further insight into understanding the underlying gene regulatory networks in TMs and facilitate the identification of cell-cell communications using the spatial information within a TM or between TMs. In addition, with the broad usability and accumulation of CODEX data on the clinical side^63^, we expect that SpaGFT can discover new features (e.g., phenotype, morphology, and spatial distribution) associated with clinical outcomes regarding clinical variables, such as demographic and therapeutical information.

However, there is still room for improving prediction performance and understanding the TM mechanism. First, although the SpaGFT computation speed is very competitive, it can be further enhanced by reducing the computational complexity from *O*(*n*^2^) to *O*(*n*×*log*(*n*)) using fast Fourier transform algorithms^64^. Second, the alteration of the spot graph and TM topology represents a potential challenge in identifying TMs across spatial samples from different tissues or experiments, which results in diverse FM spaces and renders the FCs incomparable. This is similar to the “batch effect” issue in multiple single-cell RNA sequencing (scRNA-seq) integration analyses. One possible solution to this challenge is to embed and align spatial data points to a fixed topological space using machine learning frameworks, such as optimal transport. Third, SpaGFT implementation on the CODEX image relies on experts’ knowledge to pre-select functional regions. The future direction of analyzing multiplexed images is to develop a topological learning framework to automatically detect and segment functional objects based on SpaGFT feature representation.

## Online Methods

We introduce Spatial Graph Fourier Transform (SpaGFT) to identify SVGs and characterize TMs based on spatial omics data. The core concept of SpaGFT is to transform spatial gene expressions into Fourier coefficients (FC) for downstream analysis, such as expression signal enhancement and TM characterization. The main framework of SpaGFT includes three major steps: graph signal transform, SVG identification, and TM characterization. The detailed theoretical foundation can be found in **Supplementary Note 1**.

### Graph signal transform

#### K-nearest neighbor (KNN) Graph construction

Given a gene expression matrix containing *n* spots, including their spatial coordinates and *m* genes, SpaGFT calculates the Euclidean distances between each pair of spots based on spatial coordinates first. In the following, an undirected graph *G* = (*V*,*E*) will be constructed, where *V* = {*v*_1_, *v*_2_,…, *v*_n_} is the node set corresponding to *n* spots; *E* is the edge set while there exists an edge *e_ij_* between *v_i_* and *v_j_* in *E* if and only if *v_i_* is the KNN of *v_i_* or *v_j_* is the KNN of *v_i_* based on Euclidean distance, where *i,j* = 1,2,…,*n*; and *i* ≠ *j*. Based on the `benchmarking`, the default `K is defined` as 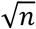. Note that all the notations of matrices and vectors are bolded, and all the vectors are treated as column vectors in the following description. An adjacent binary matrix *A* = (*a_ij_*) with rows and columns as *n* spots is defined as:

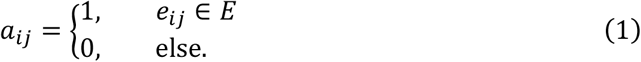

A diagonal matrix ***D*** = *diag*(*d*_1_, *d*_2_,…, *d*_n_), where 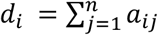 represents the degree of *v_i_*.

#### Fourier mode calculation

Using matrices ***A*** and ***D***, a Laplacian matrix ***L*** can be obtained by

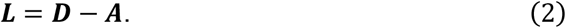

The Laplacian matrix ***L*** can be decomposed using spectral decomposition

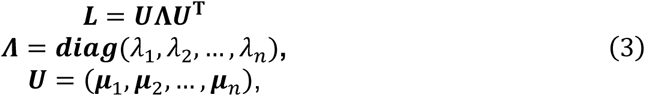

where the diagonal elements of ***Λ*** are the eigenvalues of ***L*** with *λ*_1_ ≤ *λ*_2_ ≤… ≤ *λ_n_*, where *λ_1_* is always equal to 0 regardless of graph topology. Thus, *λ*_1_ is excluded for the following analysis. The columns of *U* are the unit eigenvector of ***L. μ**_k_* is the *k^th^* Fourier mode (FM), 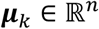, *k* = 1, 2,…, *n*, and the set {***μ***_1_, ***μ***_2_,…, ***μ**_k_*} is an orthogonal basis for the linear space (**Supplementary Figs. 1a and 1c**). For 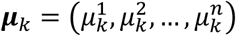, where 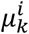 indicates the value of the *k^th^* FM on node *v_i_*, the smoothness of ***μ**_k_* reflects the total variation of the *k^th^* FM in all mutual adjacent spots, which can be formulated as

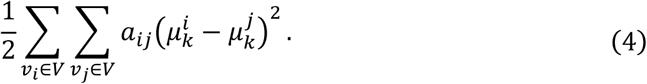

The form can be derived by matrix multiplication as

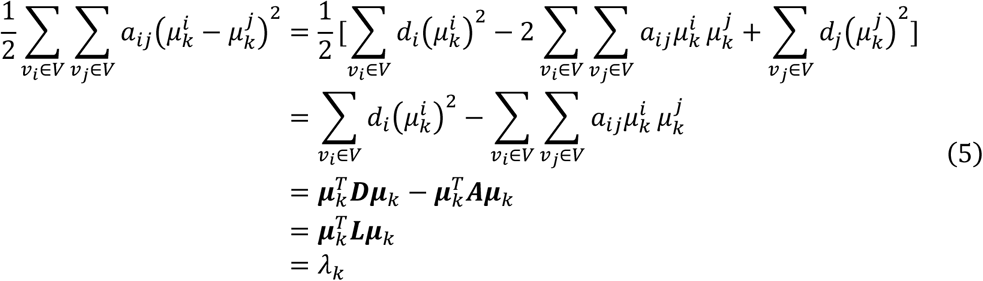

where 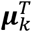 is the transpose of ***μ**_k_*. According to the definition of smoothness, if an eigenvector corresponds to a small eigenvalue, it indicates the variation of FM values on adjacent nodes is low. The increasing trend of eigenvalues corresponds to an increasing trend of oscillations of eigenvectors; hence, the eigenvalues and eigenvectors of ***L*** are used as frequencies and FMs in our SpaGFT, respectively. Intuitively, a small eigenvalue corresponds to a low-frequency FM, while a large eigenvalue corresponds to a high-frequency FM.

#### Graph Fourier transform

The graph signal of a gene *g* is defined as 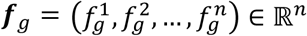, which is a *n*-dimensional vector and represents the gene expression values across *n* spots. The graph signal ***f**_g_* is transformed into a Fourier coefficient 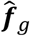 by

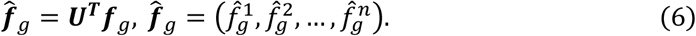

In such a way, 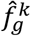 is the projection of ***f**_g_* on FM ***μ**_k_*, representing the contribution of FM ***μ**_k_* to graph signal ***f**_g_*, *k* is the index of ***f**_g_* (e.g., *k* = 1,2,…,*n*). This Fourier transform harmonizes gene expression and its spatial distribution to represent gene *g* in the frequency domain. The details of SVG identification using 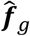 can be found below.

### SVG identification

#### GFTscore definition

We designed a *GFTscore* to quantitatively measure the randomness of gene expressions distributed in the spatial domain, defined as

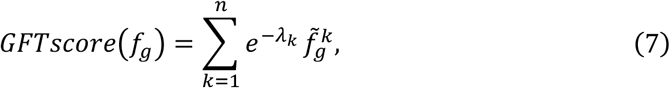

where *λ_k_* is the pre-calculated eigenvalue of ***L***, and the normalized Fourier coefficient 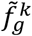 is defined as:

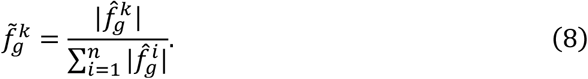

The gene with a high *GFTscore* tends to be SVG and vice versa. Therefore, all *m*genes are decreasingly ranked based on their *GFTscore* from high to low and denote these *GFTscore* values as *y*_1_ ≥ *y*_2_ ≥…≥ *y_m_*. In order to determine the cutoff *y_z_* to distinguish SVG and non-SVGs based on *GFTscore*, we applied the Kneedle algorithm^65^ to search for the inflection point of a *GFTscore* curve described below. The *GFTscore y_t_* of gene *g_t_* is converted by *y_c_t__* = *max*{*y*_1_, *y*_2_,…, *y_m_*} – *y_t_*, *t* = 1,2,…, *m*, where *y_c_t__* is the converted value of *y_t_*. Each point (*x_C_t__* = *t, y_c_t__*), where *x_c_t__* is the rank number of *y_c_t__*, is processed by a smoothing spline to preserve the curve shape and obtain (*x_S_t__, y_S_t__*), *t* = 1,2,…, *m*. Denote coordinate set 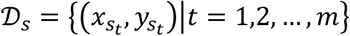 and can be normalized to corrdinate set 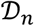 as follows:

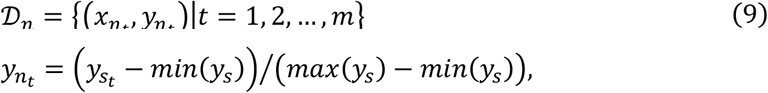

where *min*(*x_S_*) and *max*(*x_S_*) are the minimum and maximum in {*x*_*S*_1__, *x*_*S*_2__,…, *x_S_m__*}, respectively. Analogously, *min*(*y_s_*) and *max*(*y_s_*) are the minimum and maximum in {*y*_*S*_1__, *y*_*S*_2__,…, *y_S_m__*}, respectively. In addition, let 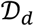 represents the set of differences between the *x*- and *y*-values, and one has:

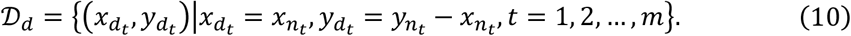

In the following, the question of determining the cutoff *y_z_* can be converted to the determination of the inflection point *y_z_* if it satisfies *y_d_z–1__ < y_d_z__*, *y*_*d*_*z*+1__ < *y_d_z__* and *y_d_h__* < *y_z_*, *h* = *z,z* + 1,…, *m*, where

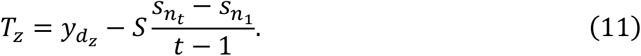

In Equation (11), *S* is a coefficient that can be used to determine the level of aggression for the inflection point.

#### Wilcoxon rank-sum test implementation for determining SVGs

Although the above GFTscore is an indicator to rank and evaluate the potential SVGs, a rigorous statistical test is needed to calculate the *p*-value for SVGs and control type I error. First, SpaGFT determines low-frequency FM and high-frequency FMs and corresponding FCs by applying the Kneedle algorithm to the eigenvalues of *L* (**Supplementary Fig. 1b**). The first and second inflection points are used for determining low-frequency and high-frequency FMs, respectively. Second, the Wilcoxon rank-sum test is utilized to test the differences between low-frequency FCs and high-frequency FCs to obtain statistical significance. If a gene has a high GFTscore and significantly adjusted *p*-value, the gene can be regarded as an SVG. We use *f*_1_, *f*_2_,…, *f_n_* to represent the expression of a random signal on *n* spots. *f_i_* is followed by Gaussian distributions and can be regarded as independent and identically distributed (*i.i.d*.) random variables^32^. By implementing GFT on (*f*_1_, *f*_2_,…,*f_n_*), we obtain Fourier coefficients *FC*_1_, *FC*_2_,…,*FC_p_*, where *p* is the number of low-frequency FCs and reflects the contributions from low-frequency FMs. We also obtain the *FC*_*p*+1_,*FC*_*p*+2_,…, *FC*_*p*+*q*_, where *q* is the number of high-frequency FCs and reflects the contributions from noise. Hence, we form the null hypothesis that no difference exists between low-frequency FCs and high-frequency FCs (**Proof can be found in C.2.3.3 of Supplementary Note 1**). Accordingly, a non-parametrical test (i.e., Wilcoxon rank-sum test) is used for testing the difference between median values of low-frequency FCs and high-frequency FCs. Especially, the null hypothesis is that the median of low-frequency FCs of an SVG is equal to or lower than the median of high-frequency FCs. The alternative hypothesis is that the median of low-frequency FCs of a SVG is higher than the median of high-frequency FCs. The *p*-value of each gene is calculated based on *Wilcoxon* one-sided rank-sum test and then adjusted using the false discovery rate (FDR) method. Eventually, a gene with *GFTscore* higher than *y_z_* and adjusted *p*-value less than 0.05 is considered an SVG.

### Benchmarking data setup

#### Dataset description

Thirty-two spatial transcriptome datasets were collected from the public domain, including 30 10X Visium datasets (18 human brain data, 11 mouse brain data, and one human lymph node data) and two Slide-seqV2 datasets (mouse brain). More details can be found in **Supplementary Table 1**. Those samples were sequenced by two different spatial technologies: 10X Visium measures ~55 *μ* m diameter per spot, and Slide-seqV2 measures ~10 *μ* m diameter per spot. Three datasets were selected as the training sets for grid-search parameter optimization in SpaGFT, including two highest read-depth datasets in Visium (HE-coronal) and Slide-seqV2 (Puck-200115-08), one signature dataset in Maynard’s study^15^. The remaining 28 datasets (excluding lymph node data) were used as independent test datasets.

#### Data preprocessing

For all the 32 datasets, we adopt the same preprocessing steps based on *squidpy^66^* (version 1.2.1), including filtering genes that have expression values in less than ten spots, normalizing the raw count matrix by counts per million reads method and implementing log-transformation to the normalized count matrix. No specific preprocessing step was performed on the spatial location data.

#### Benchmarking SVG collection

We collected SVG candidates from five publications^15,26–29^, with data from either human or mouse brain subregions. (*i*) A total of 130 layer signature genes were collected from Maynard’s study^15^. These genes are potential multiple-layer markers validated in the human dorsolateral prefrontal cortex region. (*ii*) A total of 397 cell-type-specific (CTS) genes in the adult mouse cortex were collected from Tasic’s study (2016 version)^29^. The authors performed scRNA-seq on the dissected target region, identified 49 cell types, and constructed a cellular taxonomy of the primary visual cortex in the adult mouse. (*iii*) A total of 182 CTS genes in mouse neocortex were collected from Tasic’s study^28^. Altogether, 133 cell types were identified from multiple cortical areas at single-cell resolution. (*iv*) A total of 260 signature genes across different major regions of the adult mouse brain were collected from Ortiz’s study^26^. The authors’ utilized spatial transcriptomics data to systematically profile subregions and delivered the subregional genes using consecutive coronal tissue sections. (*v*) A total of 86 signature genes in the cortical region shared by humans and mice were collected from Hodge’s study^27^. Collectively, a total of 849 genes were obtained, among which 153 genes were documented by multiple papers. More details, such as gene names, targeted regions, and sources, can be found in **Supplementary Table 2**.

Next, the above 849 genes were manually validated on the *in-situ* hybridization (ISH) database deployed on the Allen Brain Atlas (https://mouse.brain-map.org/). The ISH database provided ISH mouse brain data across 12 anatomical structures (i.e., Isocortex, Olfactory area, Hippocampal formation, Cortical subplate, Striatum, Pallidum, Thalamus, Hypothalamus, Midbrain, Pons, Medulla, and Cerebellum). We filtered the 849 genes as follows: (*i*) If a gene is showcased in multiple anatomical plane experiments (i.e., coronal plane and sagittal plane), it will be counted multiple times with different expressions in the corresponding experiments, such that 1,327 genes were archived (**Supplementary Table 3**). (*ii*) All 1,327 genes were first filtered by low gene expressions (cutoff is 1.0), and the *FindVariableFeatures* function (*“vst”* method) in the Seurat (v4.0.5) was used for identifying highly variable genes across twelve anatomical structures. Eventually, 458 genes were kept and considered as curated benchmarking SVGs.

### SpaGFT implementation and grid search of parameter optimization

A grid-search was set to test for six parameters, including *ratio_neighbors* (0.5, 1, 1.5, 2) for KNN selection and *S* (4, 5, 6, 8) for the inflection point coefficient, resulting in 16 parameter combinations. We set 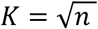 as the default parameter for constructing the KNN graphs in SpaGFT. SVGs were determined by genes with high *GFTscore* via the *KneeLocator* function (curve-convex’, direction-deceasing’, and S = 6) in the *kneed* package (version 0.7.0) and FDR (cutoff is less than 0.05). Detailed implementation and tutorial can be found on SpaGFT GitHub: https://github.com/OSU-BMBL/SpaGFT.

### Parameter setting of other tools

i. SpatialDE (version 1.1.3) is a method for identifying and describing SVGs based on Gaussian process regression used in geostatistics. *SpatialDE* consists of four steps, establishing the SpatialDE model, predicting statistical significance, selecting the model, and expressing histology automatically. We selected two key parameters, *design_formula* (‘0’ and ‘1’) in the *NaiveDE.regress_out* function and *kernel_space*(“{‘SE’:[5.,25.,50.],’const’:0}”, “{‘SE’:[6.,16.,36.],’const’:0}”, “{‘SE’:[7.,47.,57.],’const’:0}”, “{‘SE’:[4.,34.,64.],’const’:0}”, “{‘PER’:[5.,25.,50.],’const’:0}”, “{‘PER’:[6.,16.,36.],’const’:0}”, “{‘PER’:[7.,47.,57.],’const’:0}”, “{‘PER’:[4.,34.,64.],’const’:0}”, and “{‘linear’:0,’const’:0}”) in the *SpatialDE.run* function for parameter tunning, resulting in 18 parameter combinations.
ii. SPARK (version 1.1.1) is a statistical method for spatial count data analysis through generalized linear spatial models. Relying on statistical hypothesis testing, SPARX identifies SVGs via predefined kernels. First, raw count and spatial coordinates of spots were used to create the SPARK object via filtering low-quality spots (controlled by *min_total_counts*) or genes (controlled by *percentage*). Then the object was followed by fitting the count-based spatial model to estimate the parameters via *spark.vc* function, which is affected by the number of iterations (*fit.maxiter*) and models (*fit.model*). Lastly, ran *spark.test* function to test multiple kernel matrices and obtain the results. We selected four key parameters, *percentage* (0.05, 0.1, 0.15), *min_total_counts* (10, 100, 500) in *CreateSPARKObject* function, *fit.maxiter* (300, 500, 700), and *fit.model* (“poisson”, “gaussian”) in *spark.vc* function for parameter tuning, resulting in 54 parameter combinations.
iii. SPARK-X (version 1.1.1) is a non-parametric method that tests whether the expression level of the gene displays any spatial expression pattern via a general class of covariance tests. We selected three key parameters, *percentage* (0.05, 0.1, 0.15), *min_total_counts* (10, 100, 500) in the *CreateSPARKObject* function, and *option* (“single”, “mixture”) in the *sparkx* function for parameter tuning, resulting in 18 parameter combinations.
iv. SpaGCN (version 1.2.0) is a graph convolutional network approach that integrates gene expression, spatial location, and histology in spatial transcriptomics data analysis. *SpaGCN* consisted of four steps, integrating data into a chart, setting the graph convolutional layer, detecting spatial domains by clustering, and identifying SVGs in spatial domains. We selected two parameters, the value of ratio (1/3, 1/2, 2/3, and 5/6) in the *find_neighbor_cluster* function and *res* (0.8, 0.9, 1.0, 1.1, and 1.2) in the *SpaGCN.train* function for parameter tuning, resulting in 20 parameter combinations.
v. MERINGUE (version 1.0) is a computational framework based on spatial autocorrelation and cross-correlation analysis. It composes of three major steps to identify SVGs. Firstly, Voronoi tessellation was utilized to partition the graph to reflect the length scale of cellular density. Secondly, the adjacency matrix is defined using geodesic distance and the partitioned graph. Finally, gene-wise autocorrelation (e.g., Moran’s I) is conducted, and a permutation test is performed for significance calculation. We selected *min.read* (100, 500, 1000), *min.lib.size* (100, 500, 1000) in the *cleanCounts* function and *filterDist* (1.5, 2.5, 3.5, 7.5, 12.5, 15.5) in the *getSpatialNeighbors* function for parameter tuning, resulting in 54 parameter combinations.

### Metrics used in benchmarking experiments

Denote *P* = {*p*_1_, *p*_2_,…, *p_p_*}, where *p* is the total number of SVGs predicted by a tool in the performance comparison. The set of 458 curated benchmarking SVGs denoted as *R* = {*r*_1_, *r*_2_,…, *r_t_*}, where *t* = 458. In addition, *C* is the complete collection of all genes in a dataset. In addition, some notions are necessary to understand the following metrics, including *TP* = |*P* ⋂ *R*|, *FP* = |*P* – *P* ⋂ *R*|, *TN* = |(*C* – *P*) ⋂ (*C* – *R*)| and *FN* = |*R* – *P* ⋂ *R*|, where *TP*, *FP*, *TN*, and *FN* represent true positive, false positive, true negative, and false negative, respectively. The following metrics were used to test the performances of various methods. All scores were calculated using customized scripts unless specifically mentioned.

i. *Jaccard index*, also named the Jaccard similarity coefficient, is used to compare the similarity and differences between limited sample sets. Define the Jaccard index between sets *P* and *R* as:

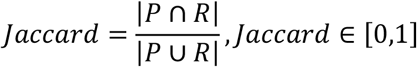 A larger *Jaccard* index indicates a higher similarity between the two sets.
ii. *Odds ratio* of the Fisher exact test is a statistical significance test used in the analysis of contingency tables, whose definition is:

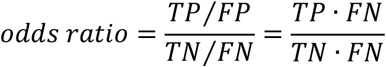 A higher *odds ratio* indicates a better prediction performance. Function *newGeneOverlap* and *testGeneOverlap* from R package *GeneOverlap* (Version 1.26.0) were used for the score calculation.
iii. *Precision* (also called positive predictive value) is the fraction of relevant instances among the retrieved instances, which is defined as:

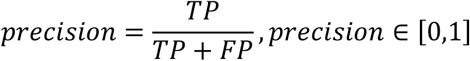 A higher precision indicates that an algorithm returns more relevant results than irrelevant ones.
iv. *Recall* (also known as sensitivity) is the fraction of relevant instances that were retrieved, which is defined as:

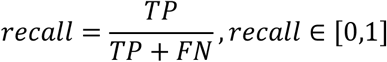 A higher recall indicates that an algorithm returns most of the relevant results (whether irrelevant ones are also returned).
v. F1 score is a measure of a test’s accuracy in statistical analysis of binary classification. It is calculated from the precision and recall of the test, defined as:

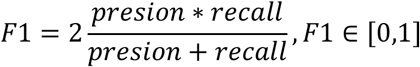 A higher F1 score indicates a better prediction performance of the algorithm.
vi. *Tversky index* is an asymmetric similarity measure on sets that compares a variant to a prototype, defined as

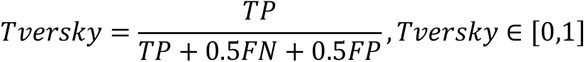 A higher *Tversky* signifies the better prediction performance of the algorithm. The *tversky.index* function from R package *tcR* (Version 2.3.2) was used for calculating the Tversky index.
vii. *Moran’s Index* used statistics to quantify the degree of spatial autocorrelation, defined as

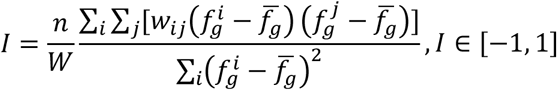

where 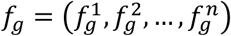 represents the gene expression values on *n* spots for gene *g*. *w_ij_* is the spatial weight between spots *i* and *j* calculated using the 2D spatial coordinates of the spots and *W* = ∑_*i*_∑*_j_W_ij_*. For each spot, we find the top *K* nearest neighbors according to Euclidean distances where *K* = 6 and *w_ij_* = 1 if spot *j* is one of the nearest neighbors of spot *i* while *w_ij_* = 0 otherwise. A *Moran’s Index* close to 1 indicates a clear spatial pattern, a value close to 0 indicates random spatial expressions, and a value close to −1 indicates a negative correlation between two adjacent spots. We applied the *moran.test* function from the *spdep* R package to generate the score.
viii. *Geary’s C* is a metric measuring spatial autocorrelation, defined as

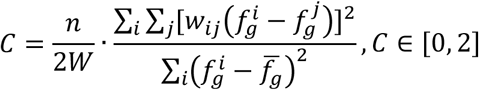

where a small *C* indicates strong spatial autocorrelation and all notations used here are the same as the notations when defining Moran’s Index. Generally, to convert it to range −1 to 1, the following formula is adopted

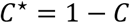 Here, the meaning of the value *C*\* is similar to Moran’s Index mentioned above. We used *geary.test* function from R package *spdep* to generate the score *C*, and then obtained *C*\* using a customized script.

### Visualization of frequency signal of SVGs in low-dimensional latent spaces

Mouse brain (i.e., HE coronal sample) with 2,702 spots was used for demonstrating FCs on distinguishing SVG and non-SVG in the 2D UMAP space. SpaGFT determined 207 low-frequency FMs using Kneedle Algorithm and computed corresponding FCs. PCA was also used for producing low-dimension representation. The transposed and normalized expression matrix was decomposed by using the *sc.tl.pca* function from the *scanpy* package (version 1.9.1). The first 207 principal components (PC) were selected for UMAP dimension reduction and visualization. The function *sc.tl.umap* was applied to conduct UMAP dimension reduction for FCs and PCs.

### SVG signal enhancement

An SVG may suffer from low expression or dropout issues due to technical bias^35^. To solve this problem, SpaGFT implemented the low-pass filter to enhance the SVG expressions. For a SVG with a measured expression value 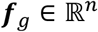, we define 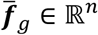 as the expected gene expression value of this SVG, and 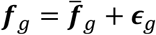, where 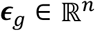 represents noises. SpaGFT estimates an approximated FCs 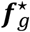 to expected gene expression 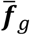 in the following way, resisting the noise *ϵ_g_*. The approximation has two requirements (*i*) the expected gene expression after enhancement should be similar to the originally measured gene expression, and (*ii*) keep low variation within estimated gene expression to prevent inducing new noises. Therefore, the following optimization problem is proposed to find an optimal solution 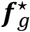 for 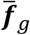

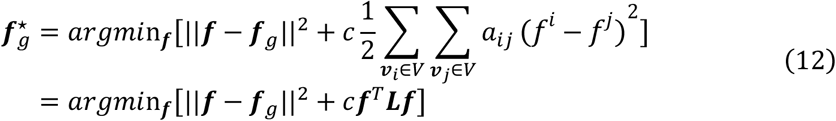

where ||·|| is the *L*2-norm, 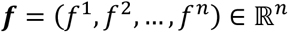 is the variable in solution space, and *i, j* = 1,2,…, *n. c* is a coefficient to determine the importance of variation of the estimated signals, and *c* >0. According to convex optimization, the optimal solution 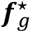 can be formulated as:

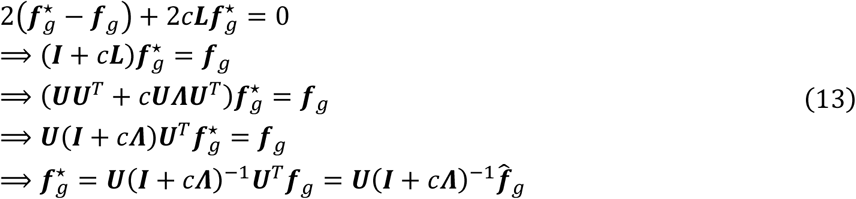

where *Λ* = *diag*(*λ*_1_, *λ*_2_,…, *λ*_n_), and ***I*** is an identity matrix. (*I* + *cΛ*)^-1^ is the low-pass filter and 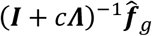 is the enhanced FCs. 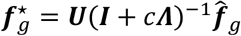 represents the enhanced SVG expression using inverse graph Fourier transform. Specifically, **in HE-coronal mouse brain data analysis, we selected** 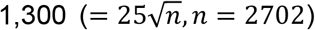 low-frequency FCs for enhancing signal and recovering spatial pattern by using inverse graph Fourier transform with *c* = 0.0001.

### SVG signal enhancement benchmarking

Sixteen human brain datasets with the well-annotated label were used for enhancement benchmarking^15,34^. SpaGFT can transform graph signals to FCs, and apply correspondence preprocessing in the frequency domain to realize signal enhancement of genes. Briefly, it composes of three major steps. Firstly, SpaGFT is implemented to obtain FCs. Secondly, a low-pass filter is applied to weigh and recalculate FCs. Lastly, SpaGFT implements iGFT to recover the enhanced FCs to graph signals. We select *c* (0.003, 0.005, 0.007) and *ratio_fms* (13, 15, 17) in the low_pass_enhancement function, resulting in 9 parameter combinations. For the parameters used for other computational tools, the details can be found as follows.

i. *SAVER-X* (version 1.0.2) is designed to improve data quality, which extracts genegene relationships by adopting a deep auto-encoder and a Bayesian model simultaneously. SAVER-X composes of three major steps roughly. Firstly, train the target data with an autoencoder without a chosen pretraining model. Secondly, filter unpredictable genes using cross-validation. Lastly, estimate the final denoised values with empirical Bayesian shrinkage. Two parameters are considered to explore the performance as well as the robustness of SAVER-X, including *batch_size* (32, 64, 128) in the *saverx* function and *fold* (4, 6, 8) in the *autoFilterCV* function, resulting in 9 parameter combinations.
ii. *Sprod* (version 1.0) is a computational framework based on latent graph learning of matched location and imaging data by leveraging information from the physical locations of sequencing to impute accurate SRT gene expression. The framework of Sprod can be divided into two major steps roughly, which are building a graph and optimizing objective function for such a graph to obtain the de-noised gene expression matrix. To validate its robustness, two parameters are adjusted, including *sprod_R* (0.1, 0.5) and *sprod_latent_dim* (8, 10, 12), to generate nine parameter combinations.
iii. *DCA* (version 0.3.1) is a deep count autoencoder network with specialized loss functions targeted to denoise scRNA-seq datasets. It uses the autoencoder framework to estimate three parameters (*μ, θ, π*) of zero-inflated negative binomial distribution conditioned on the input data for each gene. In particular, the autoencoder gives three output layers, representing for each gene the three parameters that make up the gene-specific loss function to compare to the original input of this gene. Finally, the mean (*μ*) of the negative binomial distribution represents denoised data as the main output. We set neurons of all hidden layers except for the bottleneck to (48, 64, 80) and neurons of bottleneck to (24, 32, 40) for parameter tuning, resulting in 9 parameter combinations.
iv. *MAGIC* (version 3.0.0) is a method that shares information across similar cells via data diffusion to denoise the cell count matrix and fill in missing transcripts. It composes of two major steps. Firstly, it builds its affinity matrix in four steps which include a data preprocessing step, converting distances to affinities using an adaptive Gaussian Kernel, converting the affinity matrix A into a Markov transition matrix M, and data diffusion through exponentiation of M. Once the affinity matrix is constructed, the imputation step of MAGIC involves sharing information between cells in the resulting neighborhoods through matrix multiplication. We applied the *knn* settings (3, 5, 7) and the level of *diffusion* (2, 3, 4) in the MAGIC initialization function for parameter tuning, resulting in 9 parameter combinations.
v. *scVI* (version 0.17.3) is a hierarchical Bayesian model based on a deep neural network, which is used for probabilistic representation and analysis of single-cell gene expression. It consists of two major steps. Firstly, the gene expression is compressed into a low-dimensional hidden space by the encoder, and then the hidden space is mapped to the posterior estimation of the gene expression distribution parameters by the neural network of the decoder. It uses random optimization and deep neural network to gather information on similar cells and genes, approximates the distribution of observed expression values, and consider the batch effect and limited sensitivity for batch correction, visualization, clustering, and differential expression. We selected *n_hidden* (64, 128, 256) and *gene_likelihood* (‘zinb’, ‘nb’, ‘poisson’) in the model.SCVI function for parameter tuning, resulting in 9 parameter combinations.
vi. *netNMF-sc* (version 0.0.1) is a non-negative matrix decomposition method for network regularization, which is designed for the imputation and dimensionality reduction of scRNA-seq analysis. It uses a priori gene network to obtain a more meaningful low-dimensional representation of genes, and network regularization uses a priori knowledge of gene-gene interaction to encourage gene pairs with known interactions to approach each other in low-dimensional representation. We selected *d* (8, 10, 12) and *alpha* (80, 100, 120) in the netNMFGD function for parameter tuning, resulting in 9 parameter combinations.

### TM identification

The pipeline is visualized in **Supplementary Fig. 5a**. As TMs can demonstrate functional coherence and indicate convolved collaboration among adjacent TMs, one TM is not necessarily displaying a clear boundary to its neighbor TMs. On the contrary, TMs allows the existence of overlapped region showing polyfunctional region. Computationally, the process of TM identification is to optimize the resolution parameter of the Louvain algorithm for obtaining a certain number of biology-informed TMs, which minimizes the overlapped area. Denote *G’* as the set of SVGs identified by SpaGFT. For each resolution parameter *res* > 0, *G’* can be partitioned to 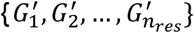 (i.e., 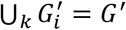 and 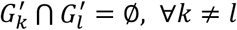) by applying the Louvain algorithm on FCs, and the resolution will be optimized by the loss function below. Denote 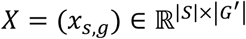 as the gene expression matrix, where *S* is the set of all spots. In the following, for each SVG group 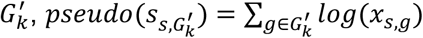 represents the pseudo expression value^13^ for spot *i*. Apply k-means algorithms with k=2 on 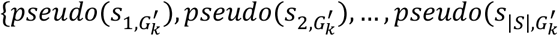 to pick out one spot cluster whose spots highly express genes in SVG group 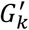 and such spot cluster is identified as a TM, denoted as *S_i_* ∈ *S*. Our objective function aims to find the best partition of *G’* such that the average overlap between any two *S_i_*, *S_j_* is minimized:

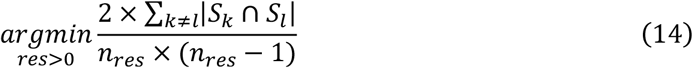

### SpaGFT implementation on mouse brain data and interpretation

We applied SpaGFT on the HE-coronal mouse data to identify TM and its associated SVGs using default parameters. To demonstrate the biological functions of identified TMs, pathway enrichment analysis (e.g., GO Biological Process 2021, KEGG 2021, and REACTOME 2022) was conducted using the Enrichr package^67^ based on the hypergeometric test for SVGs within individual TMs. Regarding MERFISH data, we use cell centroid 3D coordinates for constructing the KNN graph, and the following analyses, such as SVG identification and TM characterization, used default parameters.

### Cell2location deconvolution for generating TM-cell type correlation matrix

To generate the TM-cell type correlation matrix, we first followed the online tutorial of cell2location and calculated the cell proportion of each of the 59 cell types for the HE-coronal data across all spots (**Supplementary Table 10**). Then, pseudo-expression values across all spots for one TM were computed using the method from the **TM identification** section. Then, an element of the TM-cell type correlation matrix was calculated by computing the Pearson correlation coefficient between the proportion of a cell type and the pseudo-expression of a TM across all the spots. Lastly, the TM-cell type correlation matrix was obtained by calculating all elements as described above, with rows representing TMs and columns representing cell types.

### Identification of TM clusters among seven samples

Seven samples were collected to perform the analysis (**Supplementary Fig. 6a**). The four samples in the sagittal plane were obtained from frozen fresh samples. The other three samples were collected from the coronal plane. In addition, among three coronal plane samples, HE-coronal and GSM5519054 were frozen fresh samples but derived from different sources and sampling locations. IF-FFPE was preserved in formalin and paraffin and profiled by IF assay. The SVGs and TMs of HE-coronal had been identified from the previous Methods section (**HE-coronal mouse case analysis and interpretation**), and the SVGs and TMs of other six samples (SA1, SA2, SP1, SP2, GSM5519054, and IF-FFPE) were identified using SpaGFT with the default parameters. Then, SVGs identified from the seven samples were concatenated into an SVG-TM matrix (76 TMs in **Supplementary Table 12 and Supplementary Fig. 6b**), where values in the matrix were marked as 1 (existence) and 0 (not existence). The SVG-TM matrix was fit into PCA for dimension reduction and the Louvain algorithm for TM clustering, resulting in nine TM clusters. Among those nine TM clusters, three TM clusters contain conserved TMs from seven samples. TM-cell type correlation matrix was calculated for seven samples. To investigate the cell type composition of three TM clusters. In addition, Fisher’s exact test was performed to obtain the overlapping SVGs across identified TMs within one TM cluster, and the *p*-values were adjusted using the FDR method. The overlapped SVGs between any two TMs were calculated, and the odds ratio and adjusted *p*-values were shown on the heatmap (**Fig. 3g**).

### SpaGFT implementation on the lymph node Visium data and interpretation

#### Lymph node TM identification and interpretation

SVGs were identified on the human lymph node data (Visium) with the default setting of SpaGFT, and TMs and TM-associated SVGs were determined as described above. To demonstrate the relations between cell composition and TMs, cell2location^41^ was implemented to deconvolute spot and resolve fine-grained cell types in spatial transcriptomic data. Cell2location was used to generate the spot-cell type proportion matrix as described above, resulting in a cell proportion of 34 cell types. A TM-cell type correlation matrix was calculated using 34 lymph node cell types by the same method as previously described (the Method section of *Cell2location deconvolution for generating TM-cell type correlation matrix*). Then, the TM-cell type matrix was generated and visualized on a heatmap, and three major TMs in the lymph node were annotated, i.e., the T cell zone, GC, and B follicle.

#### Visualization of GC, T cell zone, and B follicles in the Barycentric coordinate system

Spot-cell proportion matrix was used to select and merge signature cell types of GC, T cell zone, and B follicles for generating a merged spot-cell type proportion matrix (an N-by-3 matrix, N is equal to the number of spots). For GC, B_Cycling, B_GC_DZ, B_GC_LZ, B_GC_prePB, FDC, and T_CD4_TfH_GC were selected as signature cell types. For T cell zone, T_CD4, T_CD4_TfH, T_TfR, T_Treg, T_CD4_naive, and T_CD8_naive were selected as signature cell types. For B follicle, B_mem, B_naive, and B_preGC were regarded as signature cell types. The merged spot-cell type proportion matrix was calculated by summing up the proportion of signature cell types for GC, T cell zone, and B follicle, respectively. Finally, annotated spots (spot assignment in **Supplementary Table 16**) were selected from the merged spot-cell type proportion matrix for visualization. The subset spots from the merged matrix were projected on an equilateral triangle via Barycentric coordinate project methods^11^. The projected spots were colored by TM assignment results. Unique and overlapped spots across seven regions (i.e., GC, GC-B, B, B-T, T, T-GC, and T-GC-B) from three TMs were assigned and visualized on the spatial map. Gene module scores were calculated using the *AddModuleScore* function from the Seurat (v4.0.5) package. Calculated gene module score and cell type proportion were then grouped by seven regions and visualized on the line plot (**Figs. 4e-f**). One-way ANOVA using function *aov* in R environment was conducted to test the difference among the means of seven regions regarding gene module scores and cell type proportions, respectively.

#### SpaGFT implementation on the Tonsil spatial-CITE-seq data and interpretation

SpaGFT was applied to the spatial-CITE-seq to identify TM and its associated SVGs. First, 147 SVPs were identified by the needle algorithm. Subsequently, these SVPs were selected for characterizing with resolution as 2.7, which is automatically determined based on the same objective function. TM 1 was the follicle’s main structure based on the morphology and protein signatures. Proteins such as CD9^7^ from TM 2 (**Supplementary Table 18**) showed a partially overlapped with GC, and TM 2 was selected for further investigation. To distinguish the finer structure of interested TMs (e.g., GC and crypt region), we further recluster SVPs in the interested TMs by setting resolution as 1.5. Unique and overlapped spots from TMs were then assigned and visualized on the spatial map.

#### CODEX tonsil tissue staining

An FFPE human tonsil tissue (provided by Dr. Scott Rodig, Brigham and Women’s Hospital Department of Pathology) was sectioned onto a No. 1 glass coverslip (22×22mm) pre-treated with Vectabond (SP-1800-7, Vector Labs). The tissue was deparaffinized by heating at 70°C for 1 hr and soaking in xylene 2x for 15 min each. The tissue was then rehydrated by incubating in the following sequence for 3 min each with gentle rocking: 100% EtOH twice, 95% EtOH twice, 80% EtOH once, 70% EtOH once, ddH_2_O thrice. To prepare for Heat-Induced Antigen Retrieval (HIER), a PT module (A80400012, ThermoFisher) was filled with 1X PBS, with a coverslip jar containing 1X Dako pH 9 Antigen Retrieval Buffer (S2375, Agilent) within. The PT module was then pre-warmed to 75°C. After rehydration, the tissue was placed in the pre-warmed coverslip jar, then the PT module was heated to 97°C for 20 min and cooled to 65°C. The coverslip jar was then removed from the PT module and cooled for ~15-20 min at room temperature. The tissue was then washed in rehydration buffer (232105, Akoya Biosciences) twice for 2 min each then incubated in CODEX staining buffer (232106, Akoya Biosciences) for 20 min while gently rocking. A hydrophobic barrier was then drawn on the perimeter of the coverslip with an ImmEdge Hydrophobic Barrier pen (310018, Vector Labs). The tissue was then transferred to a humidity chamber. The humidity chamber was made by filling an empty pipette tip box with paper towels and ddH_2_O, stacking the tip box on a cool box (432021, Corning) containing a −20°C ice block, then replacing the tip box lid with a 6-well plate lid. The tissue was then blocked with 200 *μ*L of blocking buffer.

The blocking buffer was made with 180 *μ*L BBDG block, 10 *μ*L oligo block, and 10 *μ*L sheared salmon sperm DNA; the BBDG block was prepared with 5% donkey serum, 0.1% Triton X-100, and 0.05% sodium azide prepared with 1X TBS IHC Wash buffer with Tween 20 (935B-09, Cell Marque); the oligo block was prepared by mixing 57 different custom oligos (IDT) in ddH_2_O with a final concentration of 0.5 *μ*m per oligo; the sheared salmon sperm DNA was added from its 10 mg/ml stock (AM9680, ThermoFisher). The tissue was blocked while photobleaching with a custom LED array for 2 hr. The LED array was set up by inclining two Happy Lights (6460231, Best Buy) against both sides of the cool box and positioning a LED Grow Light (B07C68N7PC, Amazon) above. The temperature was monitored to ensure that it remained under 35°C. The staining antibodies were then prepared during the 2 hr block.

DNA-conjugated antibodies at appropriate concentrations were added to 100 *μ*L of CODEX staining buffer, loaded into a 50-kDa centrifugal filter (UFC505096, Millipore) pre-wetted with CODEX staining buffer, and centrifuged at 12,500 *g* for 8 min. Concentrated antibodies were then transferred to a 0.1 *μ* m centrifugal filter (UFC30VV00, Millipore) pre-wetted with CODEX staining buffer, filled with extra CODEX staining buffer to a total volume of 181 *μ*l, added with 4.75 *μ*L of each Akoya blockers N (232108, Akoya), G (232109, Akoya), J (232110, Akoya), and S (232111, Akoya) to a total volume of 200 *μ*L, then centrifuged for 2 min at 12,500 *g* to remove antibody aggregates. The antibody flow through (99 *μl*) was used to stain the tissue overnight at 4°C in a humidity chamber covered with a foil-wrapped lid.

After the overnight antibody stain, the tissue was washed in CODEX staining buffer twice for 2 min each before fixing in 1.6% paraformaldehyde (PFA) for 10 min while gently rocking. The 1.6% PFA was prepared by diluting 16% PFA in CODEX storage buffer (232107, Akoya). After 1.6% PFA fixation, the tissue was rinsed in 1X PBS twice and washed in 1X PBS for 2 min while gently rocking. The tissue was then incubated in the cold (−20°C) 100% methanol on ice for 5 min without rocking for further fixation and then washed thrice in 1X PBS as before while gently rocking. The final fixation solution was then prepared by mixing 20 *μ*L of CODEX final fixative (232112, Akoya) in 1000 *μ*L of 1x PBS. The tissue was then fixed with 200 *μ*L of the final fixative solution at room temperature for 20 min in a humidity chamber. The tissue was then rinsed in 1X PBS and stored in 1X PBS at 4°C prior to CODEX imaging.

A black flat bottom 96-well plate (07-200-762, Corning) was used to store the reporter oligonucleotides, with each well corresponding to an imaging cycle. Each well contained two fluorescent oligonucleotides (Cy3 and Cy5, 5 *μ*L each) added to 240 *μ*L of plate master mix containing DAPI nuclear stain (1:600) (7000003, Akoya) and CODEX assay reagent (0.5 mg/ml) (7000002, Akoya). For the first and last blank cycles, an additional plate buffer was used to substitute for each fluorescent oligonucleotide. The 96-well plate was securely sealed with aluminum film (14-222-342, ThermoFisher) and kept at 4°C prior to CODEX imaging.

### CODEX antibody panel

The following antibodies, clones, and suppliers were used in this study:

BCL-2 (124, Novus Biologicals), CCR6 (polyclonal, Novus Biologicals), CD11b (EPR1344, Abcam), CD11c (EP1347Y, Abcam), CD15 (MMA, BD Biosciences), CD16 (D1N9L, Cell Signaling Technology), CD162 (HECA-452, Novus Biologicals), CD163 (EDHu-1, Novus Biologicals), CD2 (RPA-2.10, Biolegend), CD20 (rIGEL/773, Novus Biologicals), CD206 (polyclonal, R&D Systems), CD25 (4C9, Cell Marque), CD30 (BerH2, Cell Marque), CD31 (C31.3+C31.7+C31.10, Novus Biologicals), CD4 (EPR6855, Abcam), CD44 (IM-7, Biolegend), CD45 (B11+PD7/26, Novus Biologicals), CD45RA (HI100, Biolegend), CD45RO (UCH-L1, Biolegend), CD5 (UCHT2, Biolegend), CD56 (MRQ-42, Cell Marque), CD57 (HCD57, Biolegend), CD68 (KP-1, Biolegend), CD69 (polyclonal, R&D Systems), CD7 (MRQ-56, Cell Marque), CD8 (C8/144B, Novus Biologicals), collagen IV (polyclonal, Abcam), cytokeratin (C11, Biolegend), EGFR (D38B1, Cell Signaling Technology), FoxP3 (236A/E7, Abcam), granzyme B (EPR20129-217, Abcam), HLA-DR (EPR3692, Abcam), IDO-1 (D5J4E, Cell Signaling Technology), LAG-3 (D2G4O, Cell Signaling Technology), mast cell tryptase (AA1, Abcam), MMP-9 (L51/82, Biolegend), MUC-1 (955, Novus Biologicals), PD-1 (D4W2J, Cell Signaling Technology), PD-L1 (E1L3N, Cell Signaling Technology), podoplanin (D2-40, Biolegend), T-bet (D6N8B, Cell Signaling Technology), TCR β (G11, Santa Cruz Biotechnology), TCR-γ/δ (H-41, Santa Cruz Biotechnology), Tim-3 (polyclonal, Novus Biologicals), Vimentin (RV202, BD Biosciences), VISTA (D1L2G, Cell Signaling Technology), α-SMA (polyclonal, Abcam), and β-catenin (14, BD Biosciences).

### CODEX tonsil tissue imaging

The tonsil tissue coverslip and reporter plate were equilibrated to room temperature and placed on the CODEX microfluidics instrument. All buffer bottles were refilled (ddH_2_O, DMSO, 1X CODEX buffer (7000001, Akoya)), and the waste bottle was emptied before the run. To facilitate setting up of imaging areas and z planes, the tissue was stained with 750 *μ*L of nuclear stain solution (1 *μ*L of DAPI nuclear stain in 1500 *μ*L of 1X CODEX buffer) for 3 min, then washed with the CODEX fluidics device. For each imaging cycle, three images that corresponded to the DAPI, Cy3, and Cy5 channels were captured. The first and last blank imaging cycles did not contain any Cy3 or Cy5 oligos, and thus are used for background correction.

The CODEX imaging was operated using a 20x/0.75 objective (CFI Plan Apo Λ, Nikon) mounted to an inverted fluorescence microscope (BZ-X810, Keyence) which was connected to a CODEX microfluidics instrument and CODEX driver software (Akoya Biosciences). The acquired multiplexed images were stitched, and background corrected using the SINGER CODEX Processing Software (Akoya Biosciences). For this study, six independent 2,048×2,048 field-of-views (FOV) were cropped from the original 20,744×20,592 image. The FOVs were selected to include key cell types and tissue structures in tonsils, such as tonsillar crypts or lymphoid nodules.

### Cell segmentation

Custom ImageJ macros were used to normalize and cap nuclear and surface image signals at the 99.7th percentile to facilitate cell segmentation. Cell segmentation was performed using a local implementation of Mesmer from the DeepCell library (deepcell-tf 0.11.0)^47^, where the multiplex_segmentation.py script was modified to adjust the segmentation resolution (microns per pixel, mpp). model_mpp = 0.5 generated satisfactory segmentation results for this study. Single-cell features based on the cell segmentation mask were then scaled to cell size and extracted as FCS files.

### Cell clustering and annotation

Single-cell features were normalized to each FOV’s median DAPI signal to account for FOV signal variation, arcsinh transformed with cofactor = 150, capped between 1^st^ - 99^th^ percentile, and rescaled to 0-1. Sixteen markers (cytokeratin, podoplanin, CD31, αSMA, collagen IV, CD11b, CD11c, CD68, CD163, CD206, CD7, CD4, CD8, FoxP3, CD20, CD15) were used for unsupervised clustering using FlowSOM^48^ (66 output clusters). The cell type for each cluster was annotated based on its relative feature expression, as determined via Marker Enrichment Modeling^49^, and annotated clusters were visually compared to the original images to ensure accuracy and specificity. Cells belonging to indeterminable clusters were further clustered (20 output clusters) and annotated as above.

### SpaGFT implementation on tonsil CODEX data and interpretation

#### Resize CODEX images and SpaGFT implementation

As each FOV consisted of 2,048 by 2,048 pixels (~0.4 *μ*m per pixel size), the CODEX image needed to be scaled down to 200 by 200 pixels (~ 3.2 *μ*m per pixel size) to reduce the high computational burden (**Supplementary Fig. 9a**). Therefore, original CODEX images (2,048 by 2,048 pixels) were resized to 200 by 200 images by implementing function “resize” and selecting cubic interpolation from the imager package (v.42) in R environments. SpaGFT was then applied to the resized images by following default parameters.

#### Structural similarity (SSIM) calculation

The Structural Similarity (SSIM) score was a measurement for locally evaluating the similarity between two images regardless of image size^68^. The SSIM score ranged from 0 to 1; a higher score means more similarity between two images. It was defined as follows:

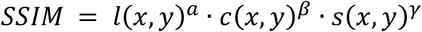

*x* and *y* were windows with 8 by 8 pixels; 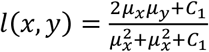 was the luminance comparison function for comparing the average brightness of the two images regarding pixels *x* and *y. C*_1_ is constant, and *α* is the weight factor of luminance comparison. 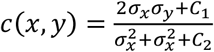 was the contrast comparison function for measuring the standard deviation of two images. *C*_2_ is constant, and *β* is the weight factor of contrast comparison. 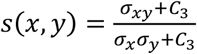 was the structure comparison by calculating the covariance between the two images. *C_3_* is constant, and *γ* is the weight factor of structure comparison.

#### Cell-cell distance and interaction analysis

To compute cell-cell distance within one TM, we first select cells assigned to each TM. An undirected cell graph was then constructed, where the cell was a node and edge connected by every two cells defined by the Delaunay triangulation using *the deldir* function from the deldir package (v.1.0-6). Subsequently, the edge represented the observed distance between the connected two cells, and Euclidean distance was used for calculating the distance based on the previous study^69^. Lastly, the average distance among different cell types was computed by taking the average of the observed cell-cell distance to generate the network plot. Regarding the determination of the cell-cell interaction, the spatial location of cells assigned in each TM was permutated and re-calculated cell-cell distance as expected distance. Based on the previous study, if the cell-cell distance was lower than 15 pm^70^ (~ 5 pixels in the 200 by 200-pixel image), the cells will contact and interact with each other. Wilcoxon rank-sum test was used for the computed p-value for expected distance and observed distance. If the expected distance was significantly smaller than the observed distance, it suggested that cells would interact with each other.

## Supporting information

Terminology Box 1

Terminology Box 2

## Data Availability

The 11 datasets from 10x Visium (ten mouse brain datasets and one human lymph node sample)^43^ can be accessed from https://www.10xgenomics.com/products/spatial-gene-expression. The GSM5519054_Visium_MouseBrain dataset is available from the GEO database with an accession number GSM5519054^42^. Regarding the human brain dataset^15^, twelve samples can be accessed via endpoint “jhpce#HumanPilot10x” on Globus data transfer platform at http://research.libd.org/globus/. The other six human brain datasets (2-3-AD_Visium_HumanBrain, 2-8-AD_Visium_HumanBrain, T4857-AD_Visium_HumanBrain, 2-5_Visium_HumanBrain, 18-64_Visium_HumanBrain, and 1-1_Visium_HumanBrain)^34^ can be downloaded from GSE220442 and https://bmbls.bmi.osumc.edu/scread/stofad-2. The two Slide-seqV2 datasets^4^ are available as accession number SCP815 in the Single Cell Portal via the link https://singlecell.broadinstitute.org/single_cell. MERFISH 3D mouse hypothalamic preoptic datasets were downloaded from https://doi.org/10.5061/dryad.8t8s248. Spatial-CITE-seq data is available from GSE213264. CODEX tonsil data is available upon request.

## Code Availability

SpaGFT is a python package for modeling and analyzing spatial transcriptomics data. The SpaGFT source code and the analysis scripts for generating results and figures in this paper are available at https://github.com/OSU-BMBL/SpaGFT.

## Acknowledgments

This work was supported by awards R01-GM131399, R21HG012482, and U54AG075931 from the National Institutes of Health. The work was also supported by the award NSF1945971 from the National Science Foundation. S.J. is funded by the Bill & Melinda Gates Foundation grant no. INV-002704, the National Institute Of Allergy And Infectious Diseases of the National Institutes of Health under award number DP2AI171139 and the Gilead Research Scholar in Hematologic Malignancies. This work was also supported by the Pelotonia Institute of Immuno-Oncology (PIIO). We thank the anatomical carton generated from the BioRender website.

## Author contributions

Conceptualization: Q.M.; methodology: J.L., Y.C., B.L., Q.M.; software coding: J.L. and Y. C.; data collection and investigation: Y.C.; experiment and interpretation: Y.Y.Y., S.J., S.R., G.N., Y.L., R.F.; data analysis and visualization: Y.C., A.M., S.J., Y.Y.Y., and J.L.; case study design and interpretation: Y.C., S.J., J.K., Y.L., A.M., and Z.L.; software testing and tutorial: J.L. and Y.C.; graphic demonstration: Y.C., J.K., S.J., A.M., and Z. L.; manuscript writing, review, and editing: all the authors.

## Supplementary information

**Supplementary Note 1 | Theoretical foundation and rationale of SpaGFT design.** This supplementary note describes (1) the biological rationale and mathematical formulation of TM characterization; (2) SVG identification regarding group Fourier transforms, GTFscore design, and statistical testing; (3) the foundation of TM characterization by a global optimized framework.

**Supplementary Note 2 | Gene function and tissue module (TM) annotation in benchmarking and case studies.** This supplementary note includes six annotations for gene and TM interpretation of different case studies.

**Supplementary Data 1 | TM ID card for all 12 TM from mouse brain.** This supplementary data includes all 12 TM ID cards for further demonstrating SVG, function enrichment, cell type proportion, and overlapped TMs.

**Supplementary Data 2 | 3D visualization of MERFISH data.** This supplementary data includes cell type annotation 3D maple and nine 3D TMs.

**Terminology Box 1 | Definition of terminologies used in SpaGFT.** This terminology box describes frequently used terminology in this paper in terms of biological terminologies and computational terminologies.

**Terminology Box 2 | Comparison of TM and spatial domain.** This terminology box compares TM and spatial domain regarding Literature description, examples, figure illustrations, applicable technologies, computational formulation, and high-order structure along with examples.

## Supplementary Tables

**Supplementary Table 1 | Data information.** The table includes information on 32 spatial transcriptome datasets from the public domain. The first column shows the data ID in the original paper or data source; the second column shows the use of the data (i.e., for grid-search optimization, independent test, or case study); the third column shows the sequencing platform; the fourth to the sixth columns show the sample information, including species, conditions, and tissue sources; the rest of the columns shows the statistical information of each data, including the number of spots, the number of genes, the number of total reads, the mean read per spot, the standard deviation of the number of reads per spot, the mean number of genes per spot, and the standard deviation of genes per spots.

**Supplementary Table 2 | 849 SVG candidates collected from the public domain.** The table collects 849 unique cell-type- or layer-specific markers from five different kinds of literature. The first column records the mouse gene symbol. The second column records the paper source. The third column records the experiment object in each gene, where “M,”“H,” and “M&H” represent mouse, human, and both. The fourth column records the human gene symbol. The fifth column records the original source in the paper for each gene, either figures or supplementary files.

**Supplementary Table 3 | 458 curated benchmarking SVGs validated by the Allen Brain Atlas.** The first six columns correspond to general information on gene identifiers, including gene symbol (mouse), gene symbol (human), UniqueID, probe name, plane, and the experiment ID in the ISH database. The ISH intensity on 12 brain regions was recorded from column G to Column R, respectively, including Isocortex, Olfactory area (OLF), Hippocampal formation (HPF), Cortical subplate (CTXsp), Striatum (STR), Pallidum (PAL), Thalamus (TH), Hypothalamus (HY), Midbrain (MB), Pons (P), Medulla (MY), and Cerebellum (CB). All the records were downloaded from the ISH database. Column S records the mean ISH intensity of 12 mouse brain regions. Column T records whether the gene is considered a curated benchmarking SVG in this paper.

**Supplementary Table 4 | Grid-search of parameter combination for SVG prediction.** The table records the details of the performance comparison in terms of the grid-search of parameter optimization. The first four columns correspond to sample ID, tested software, sequence technology, and parameter combinations. The rest of the columns records eight evaluation matrices, including the Jaccard index, Tversky index, the odds ratio of Fisher’s exact test, precision, recall, F1 score, Moran’s I, and Geary’s C. If an element in this table is “NA,” the software shows an error or ran out of time (running time was greater than 48 hours) during testing.

**Supplementary Table 5 | Running time of SpaGFT and other tools on the three grid-search test data.** The table records the running time and memory cost of SpaGFT, SPARK, SPARK-X, MERINGUE, SpatialDE, and SpaGCN on the HE-coronal, 151673, and Puck-200115-08 datasets. All tools and experiments were carried out in the same computing environment introduced in Methods. Columns A and B show tool names and sample names; Column C and D records the running time with the unit as second (S) and log10(S), respectively; Column E is memory cost with the unit as a megabyte. For any experiments that spent over 24 hours, we labeled them as “NA.”

**Supplementary Table 6 | SVG prediction performance on 28 independent test datasets using default parameters.** The table records the details of the performance comparison in terms of the independent test. The first column indicates the dataset ID, corresponding to the Dataset ID in Supplementary Table 1. The second column shows eight evaluation matrices, including the Jaccard index, the Tversky index, the odds ratio of Fisher’s exact test, precision, recall, F1 score, Moran’s I, and Geary’s C. The other columns are the software. If an element in this table is “NA,” the software shows an error or runs out of time (running time was greater than 48 hours) during testing.

**Supplementary Table 7 | Summary of top 500 genes identified by SpaGFT and the fix benchmarking tools.** The table records the unique and consistent SVGs of the top 458 SVGs identified by six tools for mouse brain data (HE-coronal). The first column is the gene name; Columns B, C, D, E, F, and G are software names; The values in Columns B to G indicate whether the gene is identified by this tool. If the value is equal to 1, it means the gene is the output of the top 458 SVGs in this software, and vice versa; Column H is the sum of values from Columns B to G, indicating the consistency of identified genes (the higher value, the higher consistency). When the value in Column H is “1,” it means that this gene is uniquely identified by one of the tools from Columns B to G; Column I indicates whether this SVG is from 458 ground truth.

**Supplementary Table 8 | Gene enhancement results.** The table demonstrates the performance of different tools, including Sprod, SAVER-X, scVI, netNMF-sc, MAGIC, and DCA, using 18 human datasets (2-3, 2-5, 2-8, 18-64, 1-1, T4857, 151507, 151508, 151509, 151510, 151669, 151670, 151671, 151672, 151673, 151674, 151675, and 151676). The first column is the method name. The second column is the sample ID. The third column is the combination parameter. The fourth column is the ARI score calculated by inputting the ground truth label and predicted label.

**Supplementary Table 9 | SVG results in the HE coronal data.** The table records all SVGs predicted from SpaGFT on the HE-coronal data. Column A is the SVG name; Columns B-D are gene interpretations; Column E is the number of spots having this SVG expressed; Column F is the corresponding *GFTscore;* Column G is the ranking of GFTscore; Columns H and I are the *p*-value and *Q*-value of SVG, respectively; Columns J is the TM labels. SVGs are arranged based on the GFTscore from high to low.

**Supplementary Table 10 | Deconvolution results for HE-coronal sample.** The table shows the proportions of 59 cell types calculated by cell2location. The first column is the spot ID of the mouse sample. The rest of the columns are the cell proportions in 59 cell types, respectively.

**Supplementary Table 11 | SVG results in the seven mouse brain data.** The table records all SVGs predicted from SpaGFT in the seven mouse brain data. Column A is the gene name; Columns B-D are gene interpretations; Column E is the number of spots having this SVG expressed; Column F is the corresponding *GFTscore*; Column G is the ranking of GFTscore; Columns H and I are the *p*-value and *q*-value of SVG, respectively; Column J is the TM labels; Column L indicates the sample names.

**Supplementary Table 12 | TM-associated SVG and TM assignment to each spot from seven samples in terms of TM cluster 1, TM cluster 2, and TM cluster 3.** The table shows 690 SVGs, TMs, and two labels for TMs. The first row is the clustering results of the Louvain algorithm. The second row is the TM clusters label assignment, including TM cluster 1, TM cluster 2, and TM cluster 3. The rest rows are SVGs (690 SVGs).

**Supplementary Table 13 | The overlapped SVGs of all TMs across seven mouse brain samples.** The table records overlapped SVGs for all TMs identified from seven samples. The element in this table is the overlapped SVG name between any two TMs. If an element in this table is “NA,”it indicates no overlapping SVGs between two TM.

**Supplementary Table 14 | SVG results in the human lymph node data.** The table records all SVGs predicted from SpaGFT on the human lymph node data. Column A is the gene name; Columns B-D are gene interpretations; Column E is the number of spots having this SVG expressed; Column F is the corresponding *GFTscore*; Column G is the ranking of GFTscore; Columns H and I are the *p*-value and *q*-value of SVG, respectively; Column J is the TM labels. SVGs are arranged based on the GFTscore from high to low.

**Supplementary Table 15 | Deconvolution results for human lymph node sample.** The table shows the proportions of 34 cell types calculated by cell2location. The first column is the spot ID of the mouse sample. The rest of the columns are the cell proportions in 34 cell types.

**Supplementary Table 16 | TM assignment to each spot from lymph node data in terms of GC, T cell zone, and B follicle.** The table demonstrates GC, T cell zone, B follicle, and their interactive region assignment label. The first column is the spot ID. The second and third columns are spatial coordinates. The last column is the assignment label, where “0” is no assignment; “T.zone” is the spot assigned as T cell zone; “B.follicle” is the spot assigned as B follicle; “GC” is the spot assigned as germinal center; “T.zone-B.follicle” is the spot assigned as the interactive region between T cell zone and B follicle; “GC-T.zone” is the spot assigned as the interactive region between GC and T cell zone; “GC-B.follicle” is the spot assigned as the interactive region between GC and B follicle; “GC-T.zone-B.follicle” is the spot assigned as interactive region among GC, T zone, and B follicle.

**Supplementary Table 17 | Pathway and corresponding gene signatures.** The table documents the gene used for pathway enrichment from the GSEA database. The first column is gene names, and the second column is pathway names.

**Supplementary Table 18 | TM and corresponding protein signature based on spatial-CITE-seq.** The table records TMs predicted from SpaGFT on the spatial-CITE-seq tonsil data. Column A is the protein name (250 proteins); Column B is the number of spots having this protein expressed; Column C is the corresponding *GFTscore;*Column D is the ranking of GFTscore; Columns E and F are the *p*-value and *q*-value of protein, respectively; Column G is the TM label. Column H is the finer structure label for TM 1 and TM 2. Proteins are arranged based on tissue modules and finer structures.

**Supplementary Table 19 | TM characterization result for six FOVs from CODEX.** The table shows the SpaGFT prediction results. Column A is the protein name; Column B is the number of pixels, suggesting the number of pixels expressing this protein; Column C is GFTscore and shows low-frequency FM contributions; Column D is the GFT score rank; Colum E and F show the p-value and adjusted the p-value based on the false discovery rate method; Column G is the TM indicator; Column H is the FOV indicator.

**Supplementary Table 20 | Cell type proportions for TMs of each FOV.** The table shows the cell type proportions for each TM. Column A is the cell type name, including B cells, CD4 cells, CD8 cells, Dendritic cells, endothelial cells, epithelial cells, lymphatic cells, macrophage 1, macrophage 2, Neutrophil, and Tregs; Column B is the number of pixels, suggesting the number of pixels presenting this cell type; Column C is the percentage of this cell type; Column D and Column E are the FOV and TM indicators, respectively; Column F is the manual annotation of TM.

## Supplementary Figures

**Supplementary Fig. 1 | FM identification and visualization. a.** Workflow of FM identification. Spot graph is constructed by KNN, where *K* is equal to the number of spot *n*. The degree and adjacency matrix are generated, then the Laplacian matrix can be calculated by subtracting the degree matrix and adjacency matrix. Through decomposing the Laplacian matrix, eigenvalue and eigenvector are obtained, where eigenvectors are the FMs. **b**. The screen plot shows the rank of the eigenvalue (the x-axis) and corresponding values (the y-axis). The low-frequency FM can be determined by the first inflection point, and the high-frequency FM can be determined by the second inflection point. **c**. Visualizations of FM patterns in the different frequency domains of the Visium 151673 dataset, where LFM means low-frequency FM and HFM means high-frequency FM.

**Supplementary Fig. 2 | Performance Comparison.** The two panels show Moran’s I and Geary’s C score on the grid-search testing for the HE-coronal sample. The boxplot indicates Moran’s I and Geary’s C score distribution for six tools’ grid-search results, respectively. The Black line in the box indicates the median value.

**Supplementary Fig. 3 | ISH evidence of eight SVGs. a-f**. The ISH database webpage shows four major information, including experiment information (top left), ISH high-resolution image (right), 3D expression (middle left), and ISH intensity of 12 mouse brain regions (bottom). In addition, we used a dashed line to circle out ISH high-intensity regions on ISH high-resolution images.

**Supplementary Fig. 4 | Workflow of SVG enhancement.** The low SVG expression signal can be enhanced by a low-pass filter and iGFT using low-frequency FCs.

**Supplementary Fig. 5 | Workflow TM characterization and TM interpretation. a**. The figure shows the workflow of TM characterization: (i) enhanced FCs are utilized for clustering and generating SVG clusters based on the Louvain clustering algorithm using an initial resolution parameter (e.g., 0.1). (ii) SVGs in each SVG cluster are recovered to enhanced graph signals (i.e., enhanced SVGs’ expression), and the pseudo-expression value of one SVG cluster is calculated by averaging these corresponding enhanced SVGs’ expressions. (iii) the pseudo-expression of SVG clusters is used for characterizing TM candidates by utilizing the k-means algorithm. (iv) All TM candidates are used for pairwise calculating average overlapped spots; subsequently, the number of overlapped spots is optimized until SpaGFT finds a resolution to produce the minimal overlapped spots and eventually produce TMs. **b**. The optimization curve shows the changes in the average number of overlapped spots across identified TMs. The y-axis is the overlapped rate (average number of spots divided by the total number of spots). The x-axis is the value of the resolution parameter from the Louvain algorithm. **c**. KEGG and REACOME pathways show the functions of TM 6. **d**. The spatial map displays cell type proportion and distribution for TM 6.

**Supplementary Fig. 6 | Samples and the pipeline for characterizing TM cluster and forming tissue motif. a.** The figure demonstrates seven sample sources, including sagittal (Sagittal Anterior 1, Sagittal Anterior 2, Sagittal posterior 1, Sagittal posterior 2) and coronal planes (HE-coronal, GSM5519054-coronal, and IF-FFPE-coronal). **b**. The pipeline of TM cluster identification. SVGs of seven samples are concatenated for the SVG-TM matrix, where values in the matrix are marked as 1 (existence) and 0 (not existence). Subsequently, the SVG-TM matrix is implemented for PCA and the Louvain algorithm to identify TM clusters. **c**. The heatmap shows gene overlapping of 21 TMs derived from three TM clusters. The color indicates the log-odds ratio of the Fisher exact test. Adjusted *p*-value (FDR method) between two samples is showcased on the heatmap. Three structures (cerebrum, hypothalamus, and white matter) with sagittal and coronal planes are derived from Allen Brain Atlas, and targeted regions are indicated by the purple color.

**Supplementary Fig. 7 | Three TMs interpretation. a**. The heatmap visualizes the transposed TM-cell type correlation matrix (i.e., TM in the column, the cell type in the row, and each element means correlation of pseudo-expression value and cell type proportion across spots assigned in this TM). According to the transposed TM-cell type correlation matrix, TM 3, TM 5, and TM 7 correspond to the T cell zone, GC, and B follicle, respectively. **b-d**. The three figures showed the TM ID card for the T cell zone, GC, and B follicle, respectively. Each ID Card displays fundamental information about each TM, including a TM spatial map, the number of SVGs, samples of SVG, functional enrichment tests (e.g., Biological Process 2021 and REACTOME 2022), and cell type compositions.

**Supplementary Fig. 8 | Cell proportion changes across different regions**. The figure shows the other lymph node-relevant cell type changes across seven different regions.

**Supplementary Fig. 9 | Workflow of resizing CODEX image and gradient pixel image comparison. a.** The original codex image has 2,048 by 2,048 pixels and is resized down to the 200-by-200 pixel image. The resized pixel image is used for TM characterization based on the SpaGFT model. **b.** Low-frequency FMs and high-frequency FMs are visualized in terms of 1,000 by 1,000 pixel image, 500-by-500 pixel image, and 200-by-200 pixel image. **c.**The figure visualizes the comparison of the gradient pixel image. The zoom-in section shows the details of three gradient images. **d.**The heatmap shows the structural similarity (SSIM) score regarding all TMs characterized based on three gradient images. The higher SSIM corresponds to the lighter color and indicates the high similarity of the two compared images.

**Supplementary Fig. 10 | Overview of characterized TMs.**The figure shows the TM characterization results for six FOVs. The first row is the pixel-level cell type annotation, corresponding to 200-by-200 pixels. The other rows showcase TM and their cell type annotations. We label B-follicle, GC, and T zone for corresponding TMs.

**Supplementary Fig. 11 | Heterogeneous molecules for TMs associated with mantle zone and GC. a.**The binary heatmap shows TM-associated spatially variable proteins (SVP). The pink color means the corresponding protein belongs to this TM. **b.**The heatmap showed the scaled protein expression. Each block visualizes TM morphology along with protein expression, and the red rectangle points out the TM-associated SVP. **c**and **d**. Overlaid CODEX image visualizes TM using TM-associated SVPs for FOV 2 and 5, respectively.

## References

1 Moffitt, J. R. et al. Molecular, spatial, and functional single-cell profiling of the hypothalamic preoptic region. Science 362, eaau5324, doi:doi:10.1126/science.aau5324 (2018).

2 Salmén, F. et al. Barcoded solid-phase RNA capture for Spatial Transcriptomics profiling in mammalian tissue sections. Nat Protoc 13, 2501–2534, doi:10.1038/s41596-018-0045-2 (2018).

3 Palla, G., Fischer, D. S., Regev, A. & Theis, F. J. Spatial components of molecular tissue biology. Nature Biotechnology, doi:10.1038/s41587-021-01182-1 (2022).

4 Stickels, R. R. et al. Highly sensitive spatial transcriptomics at near-cellular resolution with Slide-seqV2. Nature Biotechnology 39, 313–319, doi:10.1038/s41587-020-0739-1 (2021).

5 Schürch, C. M. et al. Coordinated Cellular Neighborhoods Orchestrate Antitumoral Immunity at the Colorectal Cancer Invasive Front. Cell 182, 1341–1359.e1319, doi:https://doi.org/10.1016/j.cell.2020.07.005 (2020).

6 Zhu, B. et al. Robust single-cell matching and multimodal analysis using shared and distinct features. Nature Methods 20, 304–315, doi:10.1038/s41592-022-01709-7 (2023).

7 Liu, Y. et al. High-plex protein and whole transcriptome co-mapping at cellular resolution with spatial CITE-seq. Nature Biotechnology, doi:10.1038/s41587-023-01676-0 (2023).

8 Deng, Y. et al. Spatial profiling of chromatin accessibility in mouse and human tissues. Nature, doi:10.1038/s41586-022-05094-1 (2022).

9 Rodriques, S. G. et al. Slide-seq: A scalable technology for measuring genome-wide expression at high spatial resolution. Science 363, 1463–1467, doi:10.1126/science.aaw1219 (2019).

10 Method of the Year 2020: spatially resolved transcriptomics. Nature Methods 18, 1–1, doi:10.1038/s41592-020-01042-x (2021).

11 Bhate, S. S., Barlow, G. L., Schürch, C. M. & Nolan, G. P. Tissue schematics map the specialization of immune tissue motifs and their appropriation by tumors. Cell Systems 13, 109–130.e106, doi:https://doi.org/10.1016/j.cels.2021.09.012 (2022).

12 Moses, L. & Pachter, L. Museum of Spatial Transcriptomics. bioRxiv, 2021.2005.2011.443152, doi:10.1101/2021.05.11.443152 (2021).

13 Hu, J. et al. SpaGCN: Integrating gene expression, spatial location and histology to identify spatial domains and spatially variable genes by graph convolutional network. Nature Methods, doi:10.1038/s41592-021-01255-8 (2021).

14 Cang, Z. et al. Screening cell-cell communication in spatial transcriptomics via collective optimal transport. bioRxiv, 2022.2008.2024.505185, doi:10.1101/2022.08.24.505185 (2022).

15 Maynard, K. R. et al. Transcriptome-scale spatial gene expression in the human dorsolateral prefrontal cortex. Nature Neuroscience 24, 425–436, doi:10.1038/s41593-020-00787-0 (2021).

16 Zhao, E. et al. Spatial transcriptomics at subspot resolution with BayesSpace. Nat Biotechnol, doi:10.1038/s41587-021-00935-2 (2021).

17 Liu, X., Zhao, Y. & Qi, H. T-independent antigen induces humoral memory through germinal centers. Journal of Experimental Medicine 219, e20210527, doi:10.1084/jem.20210527 (2022).

18 Segarra, S., Marques, A. G., Leus, G. & Ribeiro, A. Reconstruction of graph signals through percolation from seeding nodes. IEEE Transactions on Signal Processing 64, 4363–4378 (2016).

19 Huang, L., Needell, D. & Tang, S. Robust recovery of bandlimited graph signals via randomized dynamical sampling. arXiv preprint arXiv:2109.14079 (2021).

20 Ricaud, B., Borgnat, P., Tremblay, N., Gonçalves, P. & Vandergheynst, P. Fourier could be a data scientist: From graph Fourier transform to signal processing on graphs. Comptes Rendus Physique 20, 474–488, doi:https://doi.org/10.1016/j.crhy.2019.08.003 (2019).

21 Liu, Y. et al. Spatial-CITE-seq: spatially resolved high-plex protein and whole transcriptome co-mapping. bioRxiv, 2022.2004.2001.486788, doi:10.1101/2022.04.01.486788 (2022).

22 Ma, Q. & Xu, D. Deep learning shapes single-cell data analysis. Nature Reviews Molecular Cell Biology, doi:10.1038/s41580-022-00466-x (2022).

23 Zhu, J., Sun, S. & Zhou, X. SPARK-X: non-parametric modeling enables scalable and robust detection of spatial expression patterns for large spatial transcriptomic studies. Genome Biology 22, 184, doi:10.1186/s13059-021-02404-0 (2021).

24 Liao, J., Lu, X., Shao, X., Zhu, L. & Fan, X. Uncovering an Organ’s Molecular Architecture at Single-Cell Resolution by Spatially Resolved Transcriptomics. Trends in Biotechnology, doi:10.1016/j.tibtech.2020.05.006 (2020).

25 Lewis, S. M. et al. Spatial omics and multiplexed imaging to explore cancer biology. Nature Methods 18, 997–1012, doi:10.1038/s41592-021-01203-6 (2021).

26 Ortiz, C. et al. Molecular atlas of the adult mouse brain. Sci Adv 6, eabb3446, doi:10.1126/sciadv.abb3446 (2020).

27 Hodge, R. D. et al. Conserved cell types with divergent features in human versus mouse cortex. Nature 573, 61–68, doi:10.1038/s41586-019-1506-7 (2019).

28 Tasic, B. et al. Shared and distinct transcriptomic cell types across neocortical areas. Nature 563, 72–78, doi:10.1038/s41586-018-0654-5 (2018).

29 Tasic, B. et al. Adult mouse cortical cell taxonomy revealed by single cell transcriptomics. Nature Neuroscience 19, 335–346, doi:10.1038/nn.4216 (2016).

30 Sun, S., Zhu, J. & Zhou, X. Statistical analysis of spatial expression patterns for spatially resolved transcriptomic studies. Nat Methods 17, 193–200, doi:10.1038/s41592-019-0701-7 (2020).

31 Miller, B. F., Bambah-Mukku, D., Dulac, C., Zhuang, X. & Fan, J. Characterizing spatial gene expression heterogeneity in spatially resolved single-cell transcriptomics data with nonuniform cellular densities. Genome Research, gr. 271288.271120 (2021).

32 Svensson, V., Teichmann, S. A. & Stegle, O. SpatialDE: identification of spatially variable genes. Nature Methods 15, 343–346, doi:10.1038/nmeth.4636 (2018).

33 Li, B. et al. Benchmarking spatial and single-cell transcriptomics integration methods for transcript distribution prediction and cell type deconvolution. Nature Methods, doi:10.1038/s41592-022-01480-9 (2022).

34 Chen, S. et al. Spatially resolved transcriptomics reveals genes associated with the vulnerability of middle temporal gyrus in Alzheimer’s disease. Acta Neuropathol Commun 10, 188, doi:10.1186/s40478-022-01494-6 (2022).

35 Wang, Y. et al. Sprod for de-noising spatially resolved transcriptomics data based on position and image information. Nature Methods 19, 950–958, doi:10.1038/s41592-022-01560-w (2022).

36 Wang, J. et al. Data denoising with transfer learning in single-cell transcriptomics. Nature Methods 16, 875–878, doi:10.1038/s41592-019-0537-1 (2019).

37 Lopez, R., Regier, J., Cole, M. B., Jordan, M. I. & Yosef, N. Deep generative modeling for single-cell transcriptomics. Nature Methods 15, 1053–1058, doi:10.1038/s41592-018-0229-2 (2018).

38 Elyanow, R., Dumitrascu, B., Engelhardt, B. E. & Raphael, B. J. netNMF-sc: leveraging gene-gene interactions for imputation and dimensionality reduction in single-cell expression analysis. Genome Res 30, 195–204, doi:10.1101/gr.251603.119 (2020).

39 van Dijk, D. et al. Recovering Gene Interactions from Single-Cell Data Using Data Diffusion. Cell 174, 716–729.e727, doi:10.1016/j.cell.2018.05.061 (2018).

40 Eraslan, G., Simon, L. M., Mircea, M., Mueller, N. S. & Theis, F. J. Single-cell RNA-seq denoising using a deep count autoencoder. Nature Communications 10, 390, doi:10.1038/s41467-018-07931-2 (2019).

41 Kleshchevnikov, V. et al. Cell2location maps fine-grained cell types in spatial transcriptomics. Nature Biotechnology, doi:10.1038/s41587-021-01139-4 (2022).

42 Buzzi, R. M. et al. Spatial transcriptome analysis defines heme as a hemopexin-targetable inflammatoxin in the brain. Free Radic Biol Med 179, 277–287, doi:10.1016/i.freeradbiomed.2021.11.011 (2022).

43 Genomics, X. Spatial Gene Expression Datasets, <https://www.10xgenomics.com/resources/datasets/> (2020).

44 Zou, F. et al. Expression and Function of Tetraspanins and Their Interacting Partners in B Cells. Frontiers in Immunology 9, doi:10.3389/fimmu.2018.01606 (2018).

45 Kerfoot, S. M. et al. Germinal center B cell and T follicular helper cell development initiates in the interfollicular zone. Immunity 34, 947–960, doi:10.1016/j.immuni.2011.03.024 (2011).

46 Natkunam, Y. The Biology of the Germinal Center. Hematology 2007, 210–215, doi:10.1182/asheducation-2007.1.210 (2007).

47 Greenwald, N. F. et al. Whole-cell segmentation of tissue images with human-level performance using large-scale data annotation and deep learning. Nature Biotechnology 40, 555–565, doi:10.1038/s41587-021-01094-0 (2022).

48 Van Gassen, S. et al. FlowSOM: Using self-organizing maps for visualization and interpretation of cytometry data. Cytometry A 87, 636–645, doi:10.1002/cyto.a.22625 (2015).

49 Diggins, K. E., Greenplate, A. R., Leelatian, N., Wogsland, C. E. & Irish, J. M. Characterizing cell subsets using marker enrichment modeling. Nature Methods 14, 275–278, doi:10.1038/nmeth.4149 (2017).

50 Liu, C. C. et al. Robust phenotyping of highly multiplexed tissue imaging data using pixel-level clustering. bioRxiv, 2022.2008.2016.504171, doi:10.1101/2022.08.16.504171 (2022).

51 Pavlasova, G. & Mraz, M. The regulation and function of CD20: an “enigma” of B-cell biology and targeted therapy. Haematologica 105, 1494–1506, doi:10.3324/haematol.2019.243543 (2020).

52 Meda, B. A. et al. BCL-2 Is Consistently Expressed in Hyperplastic Marginal Zones of the Spleen, Abdominal Lymph Nodes, and Ileal Lymphoid Tissue. The American Journal of Surgical Pathology 27 (2003).

53 Hockenbery, D. M., Zutter, M., Hickey, W., Nahm, M. & Korsmeyer, S. J. BCL2 protein is topographically restricted in tissues characterized by apoptotic cell death. Proc Natl Acad Sci U S A 88, 6961–6965, doi:10.1073/pnas.88.16.6961 (1991).

54 Heit, A. et al. Vaccination establishes clonal relatives of germinal center T cells in the blood of humans. Journal of Experimental Medicine 214, 2139–2152, doi:10.1084/jem.20161794 (2017).

55 Chtanova, T. et al. T Follicular Helper Cells Express a Distinctive Transcriptional Profile, Reflecting Their Role as Non-Th1/Th2 Effector Cells That Provide Help for B Cells1. The Journal of Immunology 173, 68–78, doi:10.4049/jimmunol.173.1.68 (2004).

56 Dorfman, D. M., Brown, J. A., Shahsafaei, A. & Freeman, G. J. Programmed death-1 (PD-1) is a marker of germinal center-associated T cells and angioimmunoblastic T-cell lymphoma. Am J Surg Pathol 30, 802–810, doi:10.1097/01.pas.0000209855.28282.ce (2006).

57 Marsee, D. K., Pinkus, G. S. & Hornick, J. L. Podoplanin (D2-40) is a highly effective marker of follicular dendritic cells. Appl Immunohistochem Mol Morphol 17, 102–107, doi:10.1097/PAl.0b013e318183a8e2 (2009).

58 Gray, E. E. & Cyster, J. G. Lymph node macrophages. J Innate Immun 4, 424–436, doi:10.1159/000337007 (2012).

59 Johansson-Lindbom, B., Ingvarsson, S. & Borrebaeck, C. A. Germinal centers regulate human Th2 development. J Immunol 171, 1657–1666, doi:10.4049/jimmunol.171.4.1657 (2003).

60 Nakagawa, R. & Calado, D. P. Positive Selection in the Light Zone of Germinal Centers. Frontiers in Immunology 12, doi:10.3389/fimmu.2021.661678 (2021).

61 Allen, C. D. C. et al. Germinal center dark and light zone organization is mediated by CXCR4 and CXCR5. Nature Immunology 5, 943–952, doi:10.1038/ni1100 (2004).

62 Allen, C. D., Okada, T. & Cyster, J. G. Germinal-center organization and cellular dynamics. Immunity 27, 190–202, doi:10.1016/j.immuni.2007.07.009 (2007).

63 Wu, Z. et al. Graph deep learning for the characterization of tumour microenvironments from spatial protein profiles in tissue specimens. Nature Biomedical Engineering 6, 1435–1448, doi:10.1038/s41551-022-00951-w (2022).

64 Lu, K.-S. & Ortega, A. Fast graph Fourier transforms based on graph symmetry and bipartition. IEEE Transactions on Signal Processing 67, 4855–4869 (2019).

65 Satopaa, V., Albrecht, J., Irwin, D. & Raghavan, B. in 2011 31st international conference on distributed computing systems workshops. 166–171 (IEEE).

66 Palla, G. et al. Squidpy: a scalable framework for spatial omics analysis. Nature Methods 19, 171–178, doi:10.1038/s41592-021-01358-2 (2022).

67 Kuleshov, M. V.et al. Enrichr: a comprehensive gene set enrichment analysis web server 2016 update. Nucleic Acids Res 44, W90–97, doi:10.1093/nar/gkw377 (2016).

68 Zhou, W., Bovik, A. C., Sheikh, H. R. & Simoncelli, E. P. Image quality assessment: from error visibility to structural similarity. IEEE Transactions on Image Processing 13, 600–612, doi:10.1109/TIP.2003.819861 (2004).

69 Jiang, S. et al. Combined protein and nucleic acid imaging reveals virus-dependent B cell and macrophage immunosuppression of tissue microenvironments. Immunity 55, 1118–1134.e1118, doi:10.1016/j.immuni.2022.03.020 (2022).

70 Fang, R. et al. Conservation and divergence of cortical cell organization in human and mouse revealed by MERFISH. Science 377, 56–62, doi:10.1126/science.abm1741 (2022).

