## Supplementary figures and images for "Spatial omics representation and functional tissue module inference using graph Fourier transform"

### Supp Fig 1

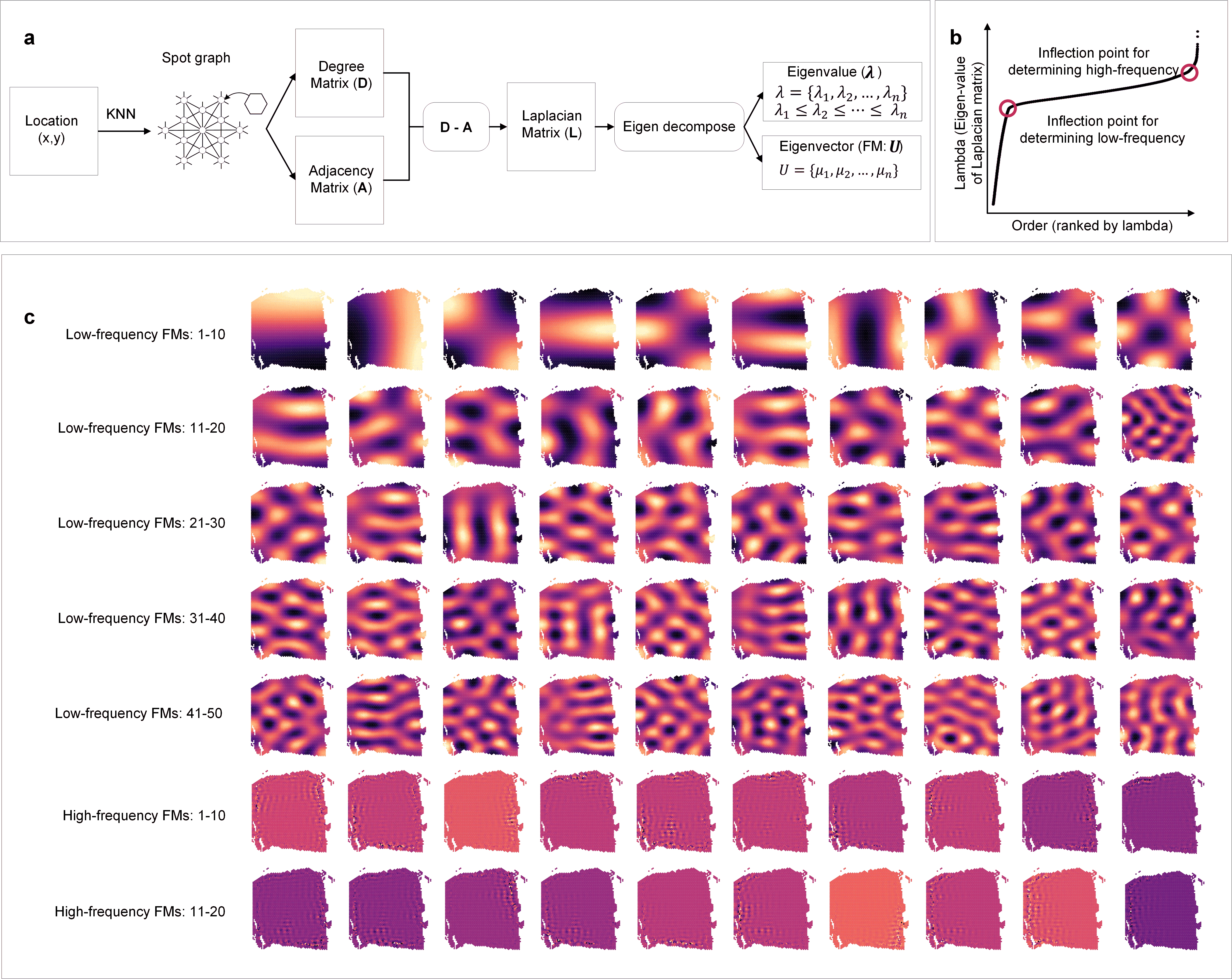

### Supp Fig 2

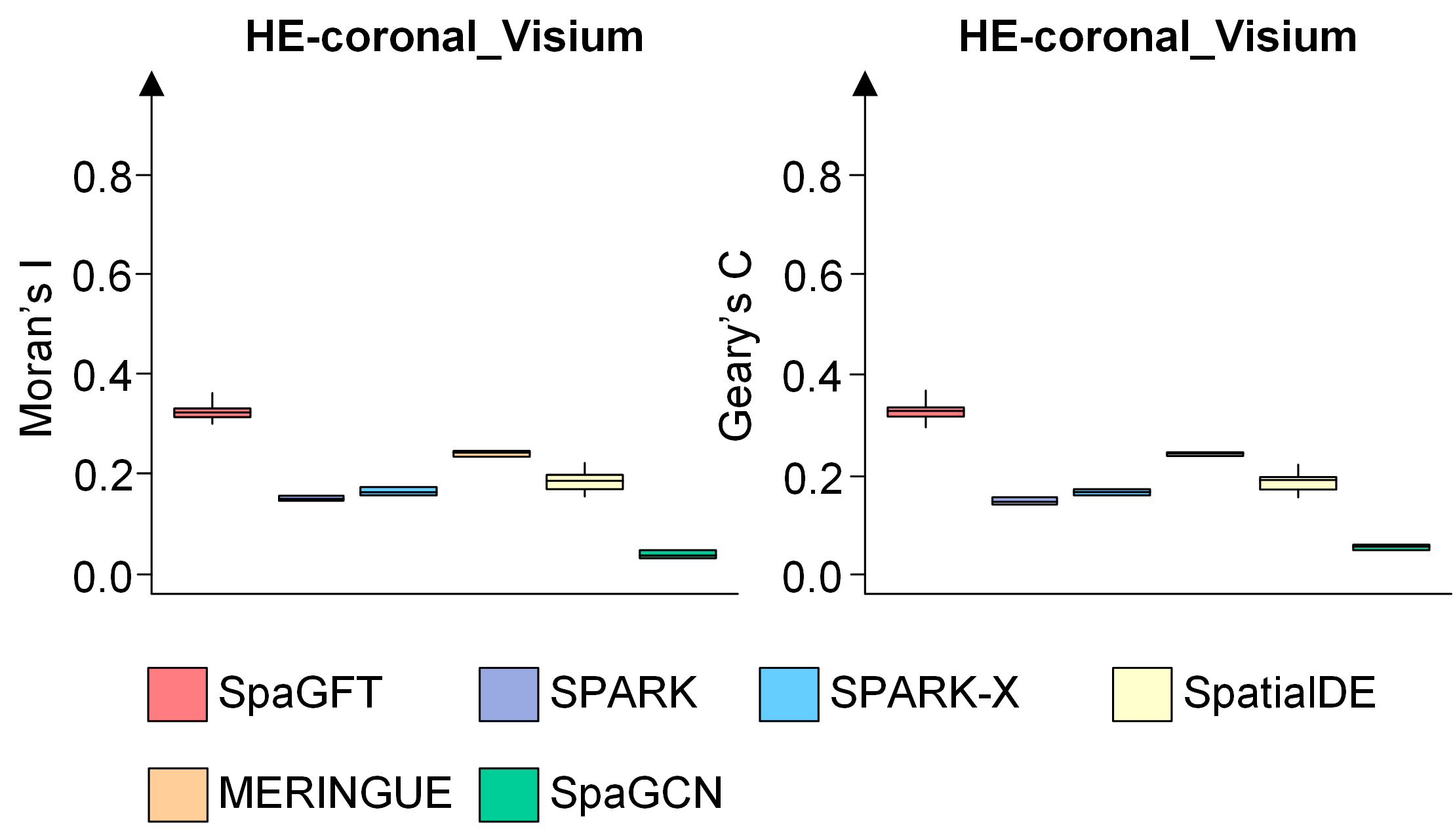

### Supp Fig 3

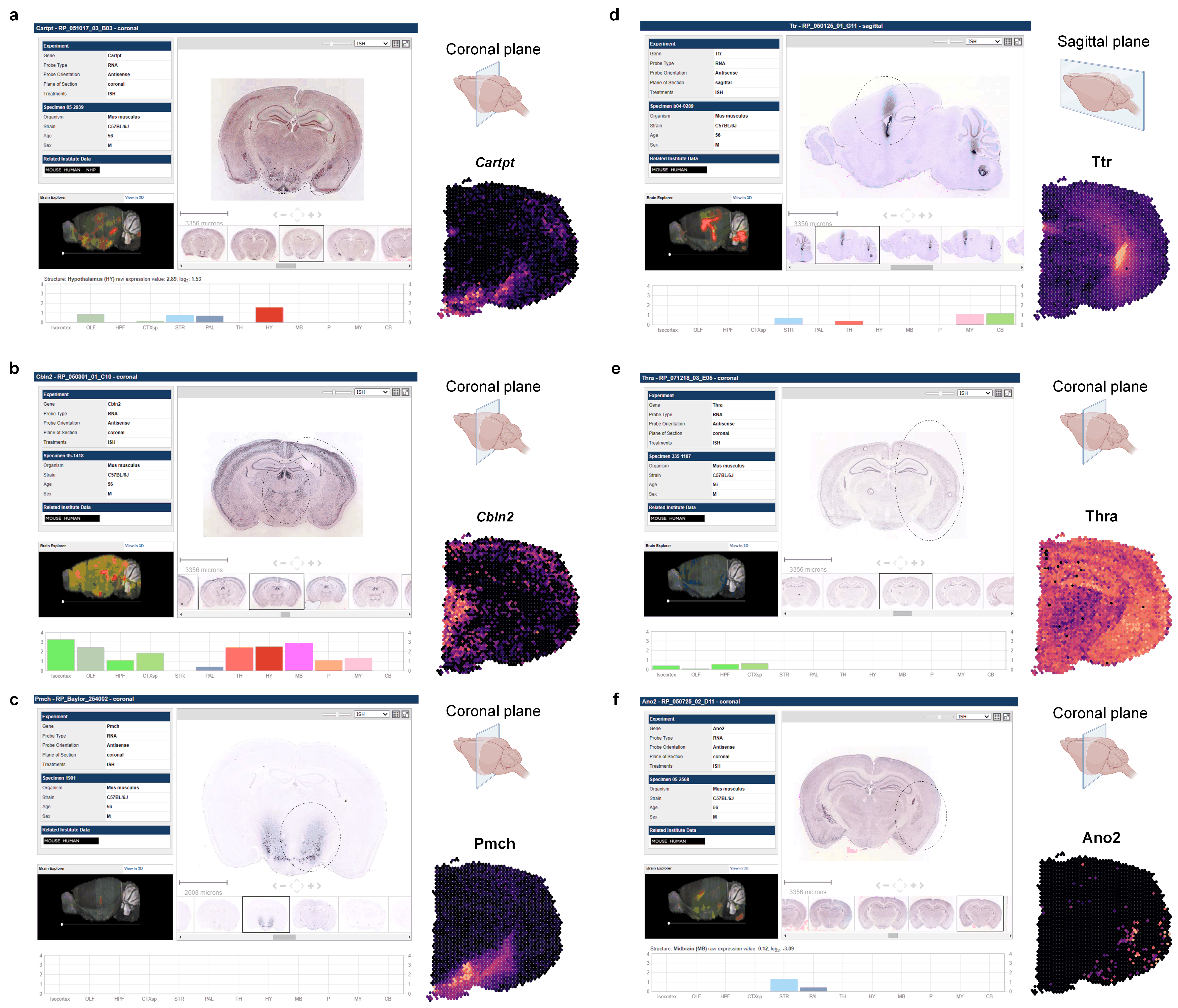

### Supp Fig 4

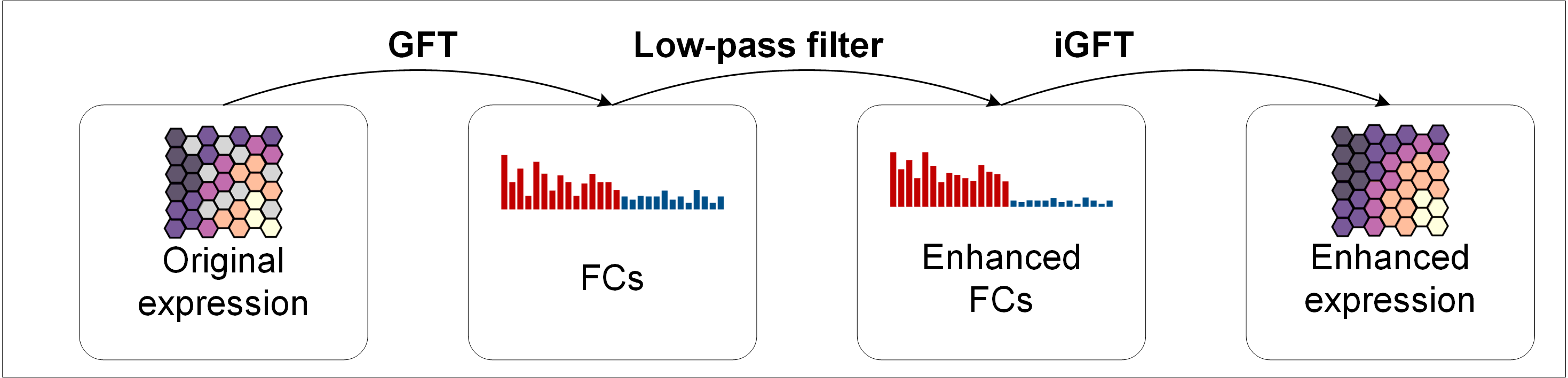

### Supp Fig 5

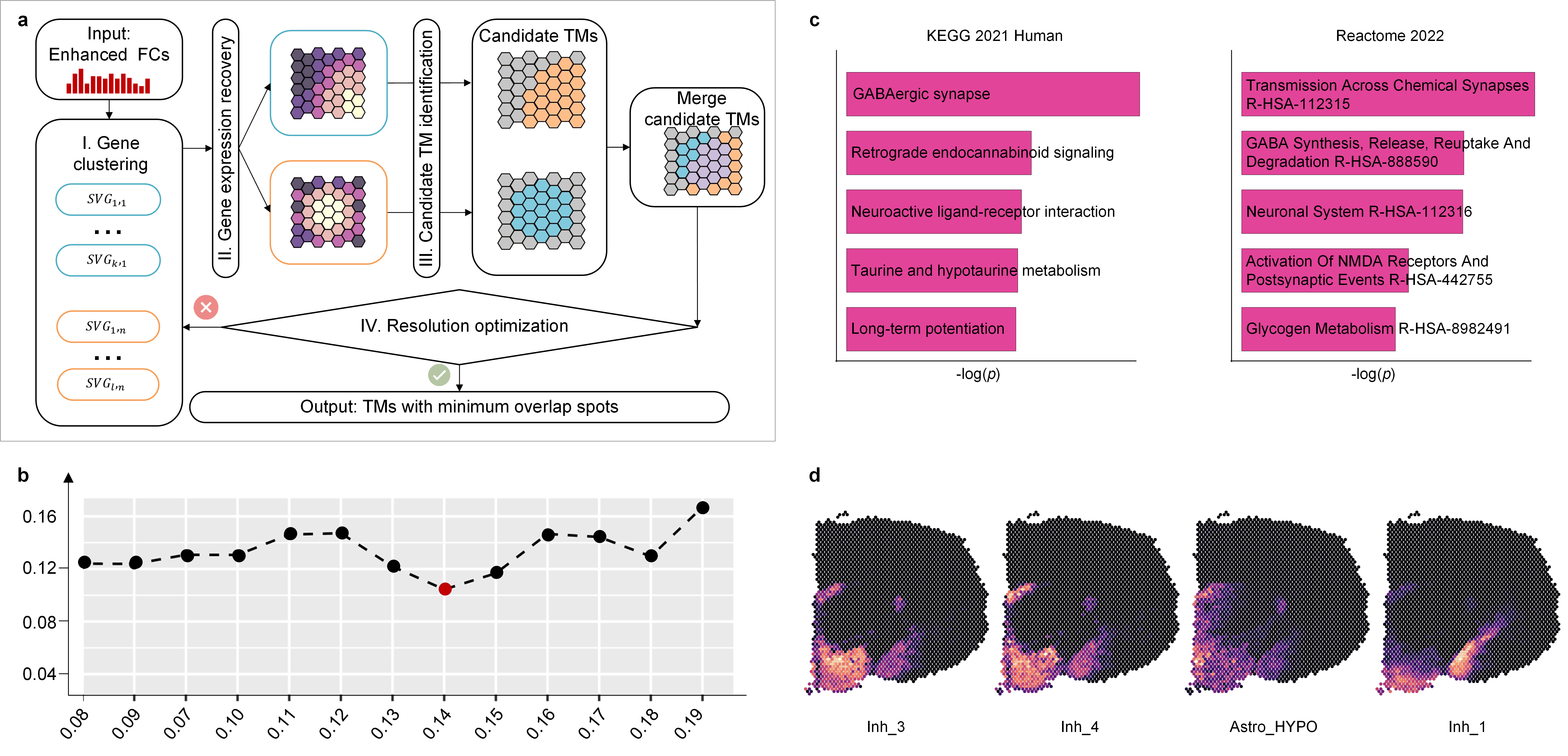

### Supp Fig 6

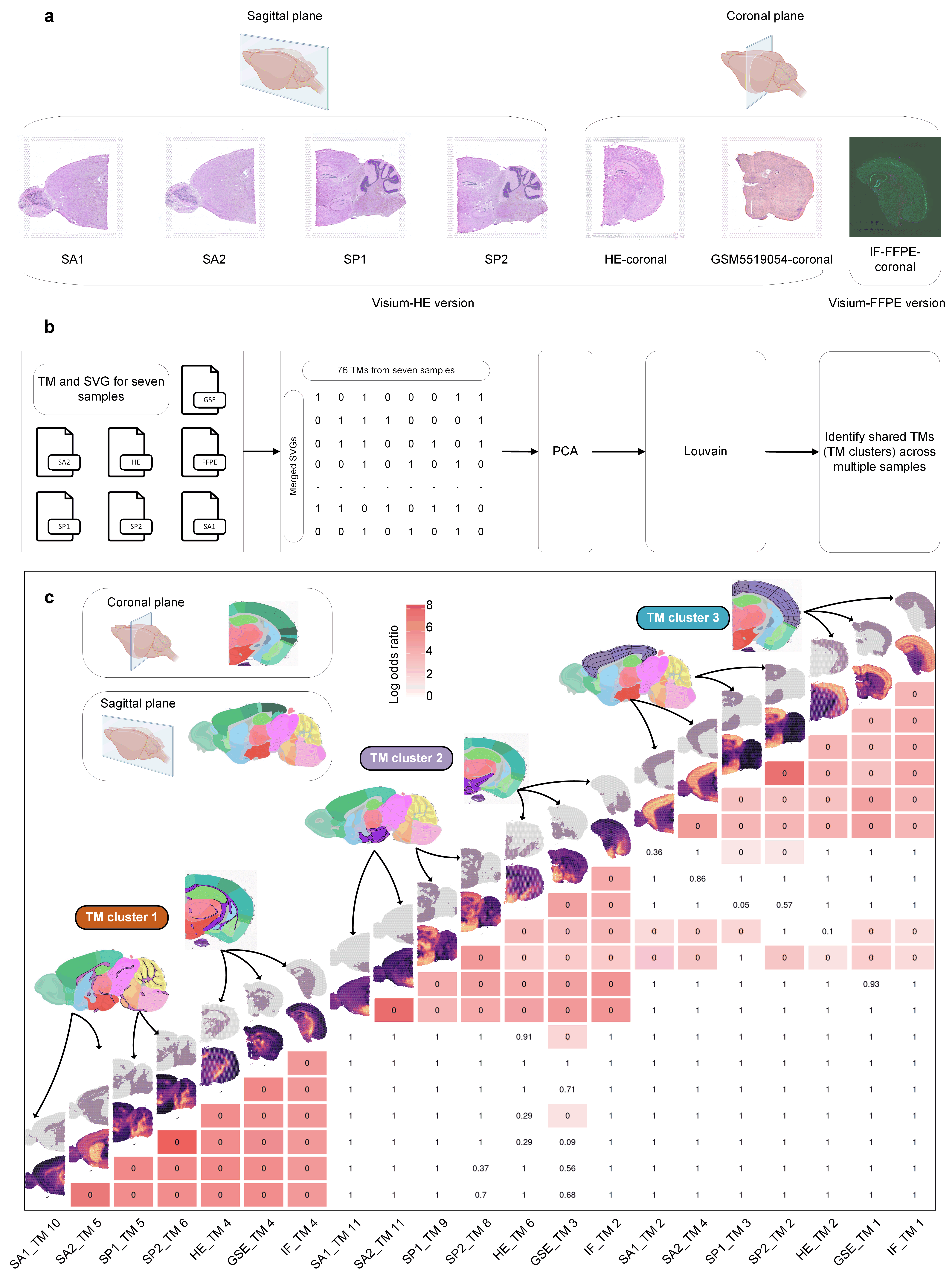

### Supp Fig 7

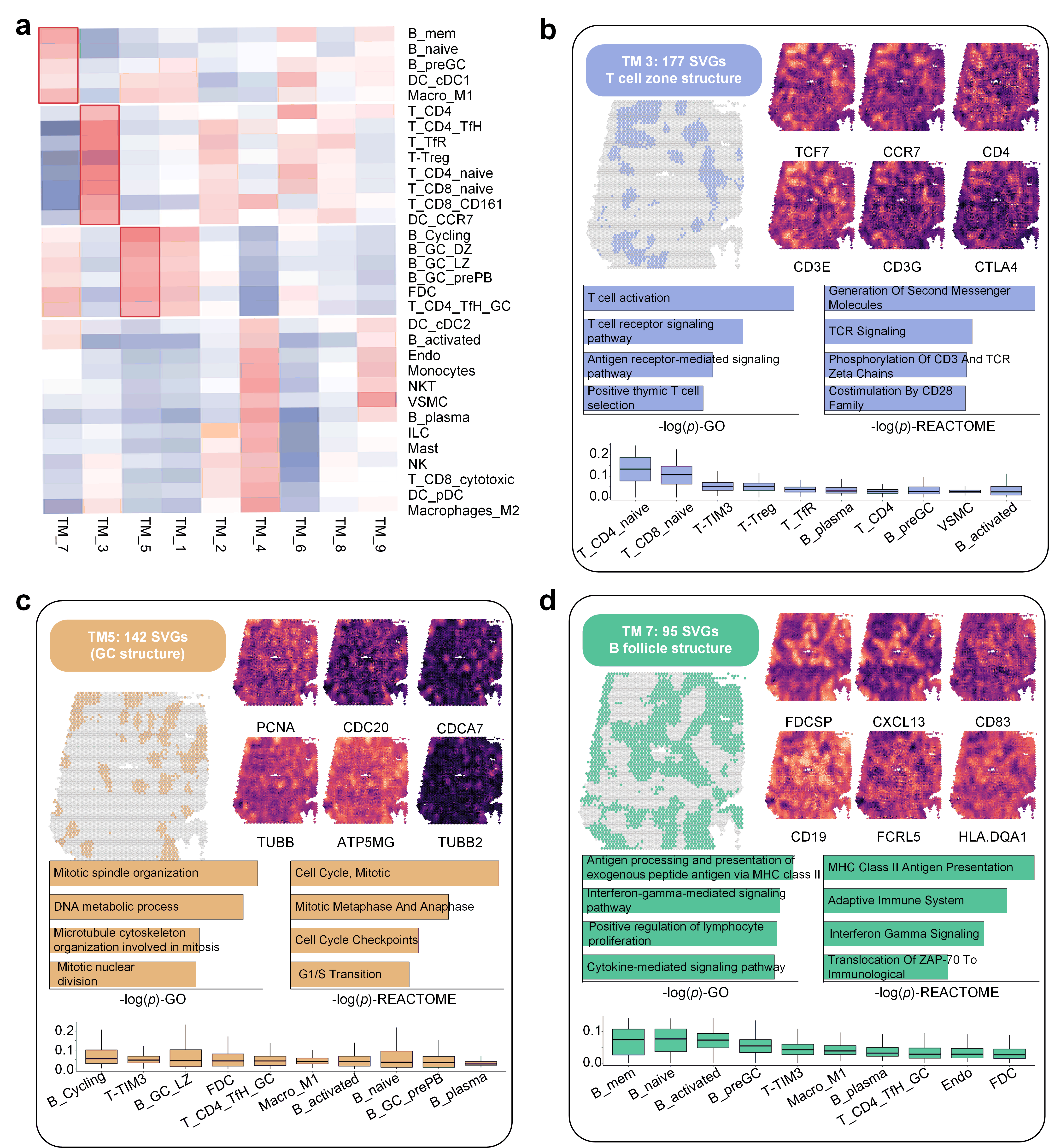

### Supp Fig 8

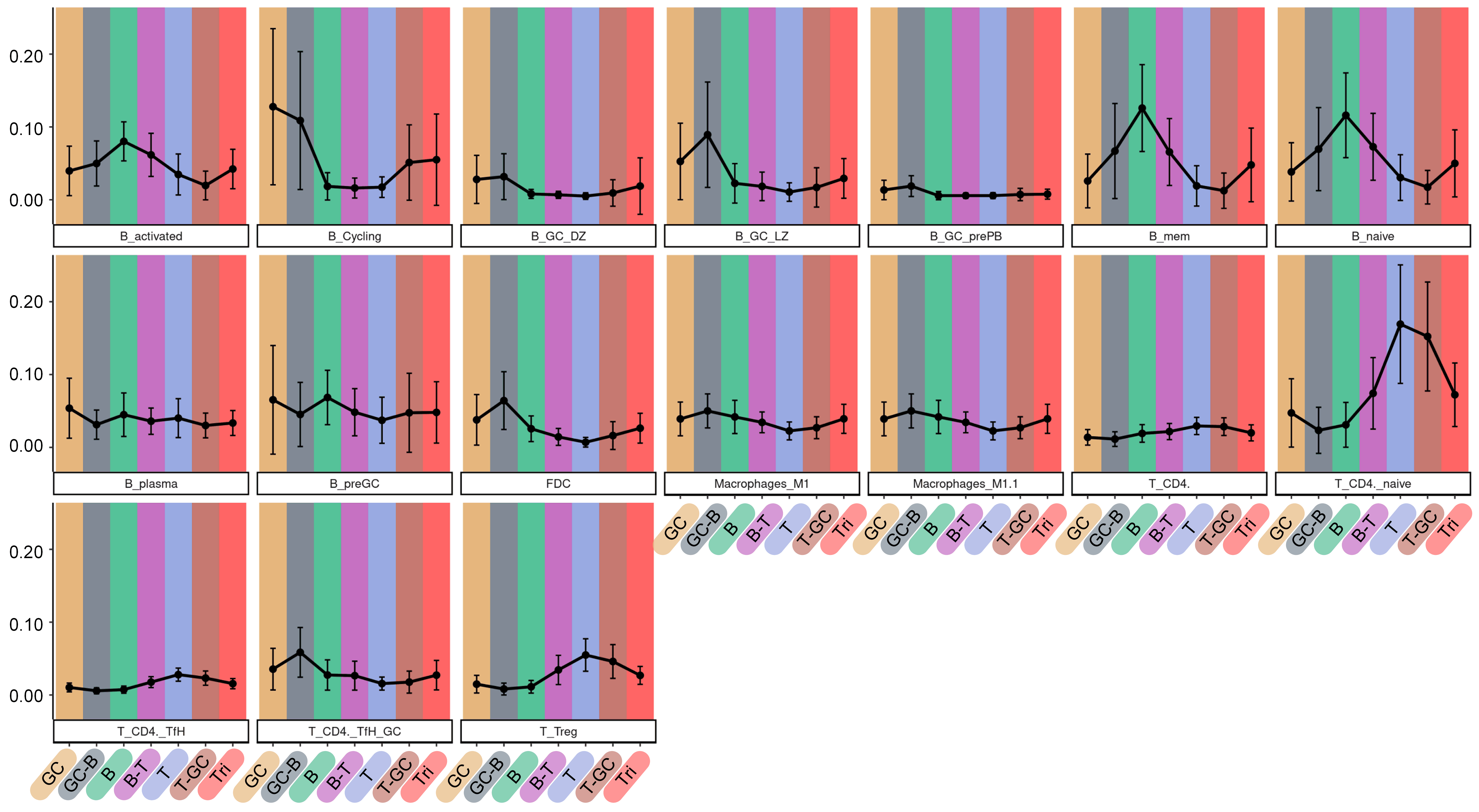

### Supp Fig 9

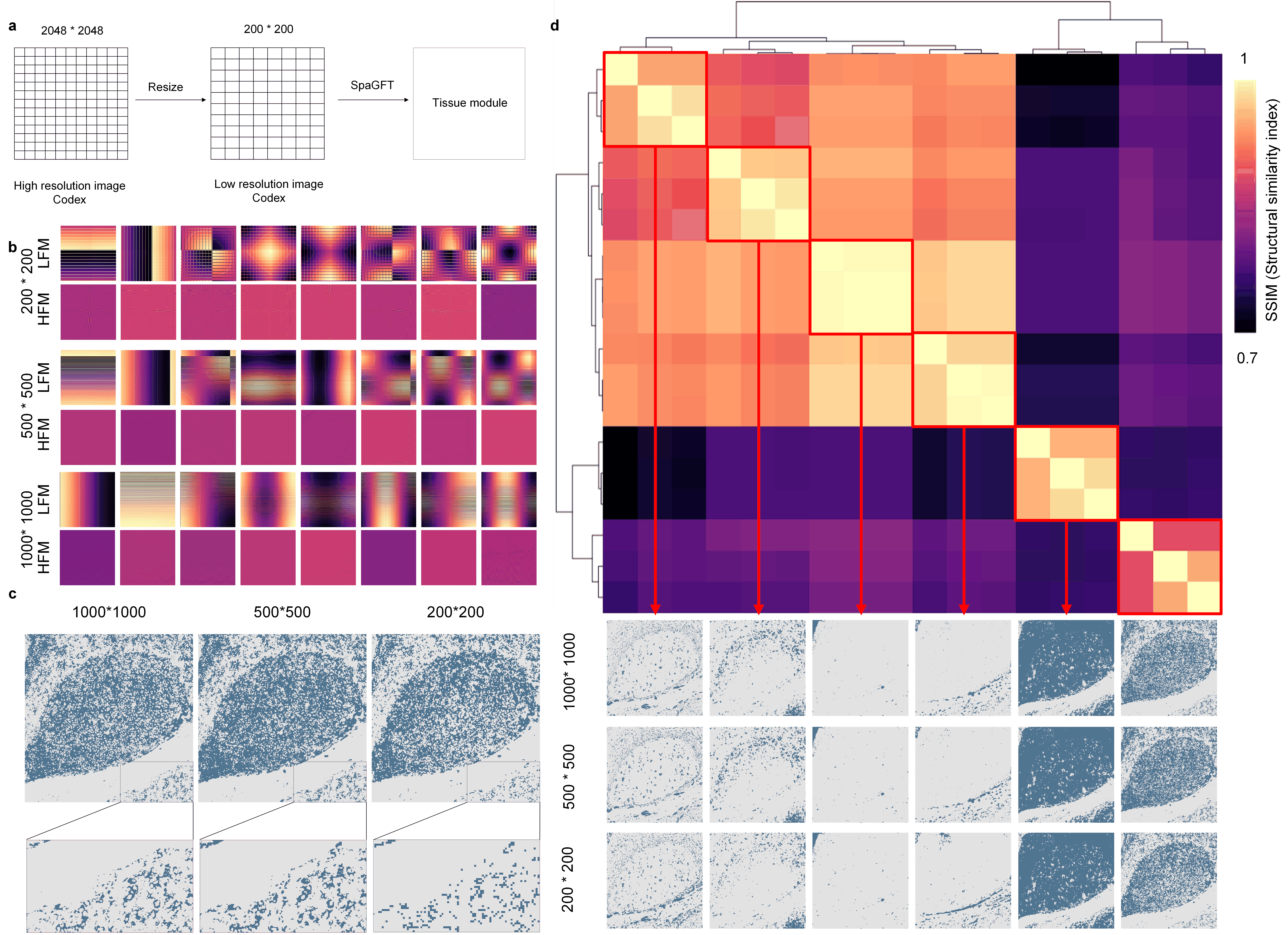

### Supp Fig 10

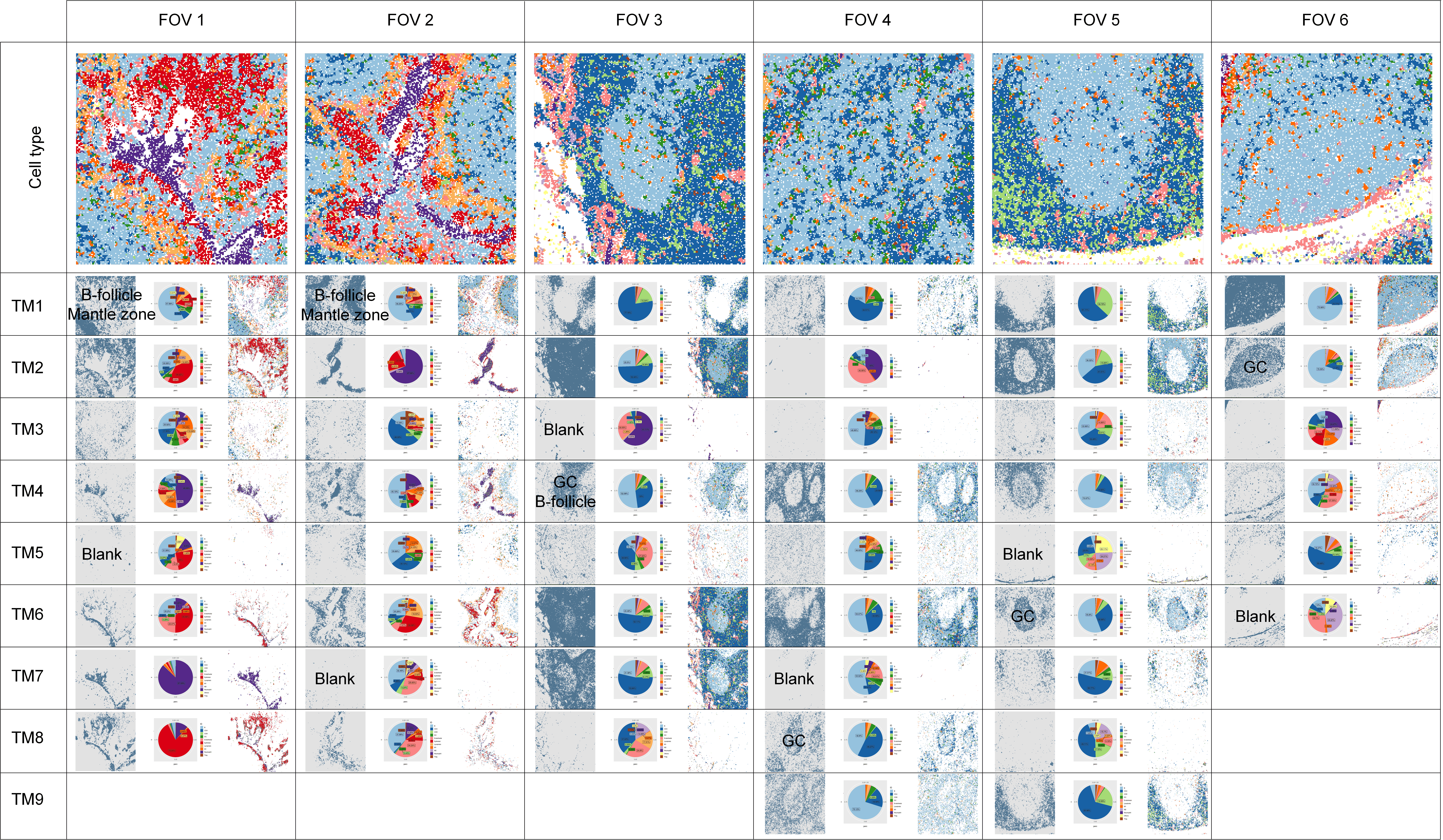

### TM1.tif

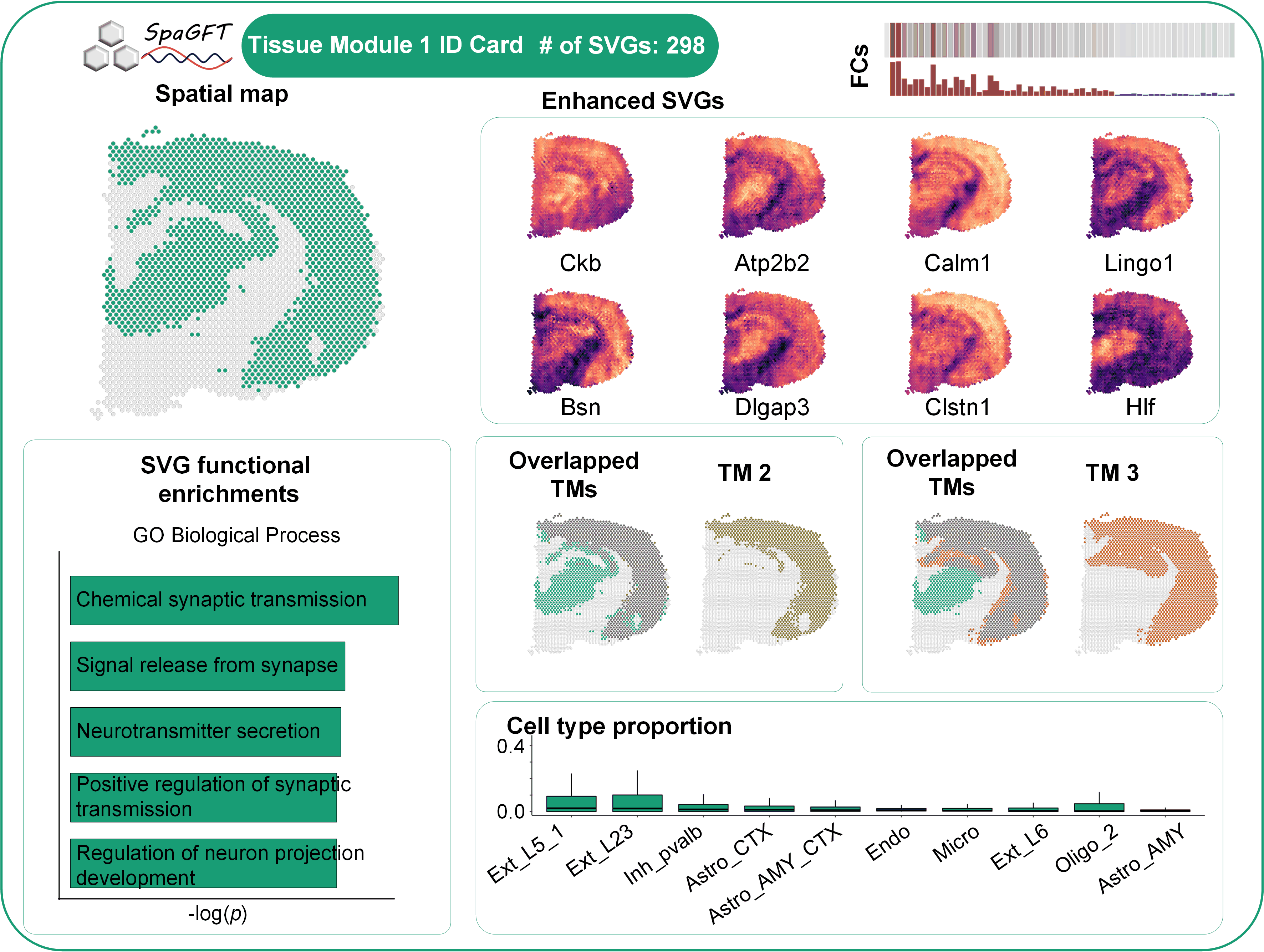

### TM2.tif

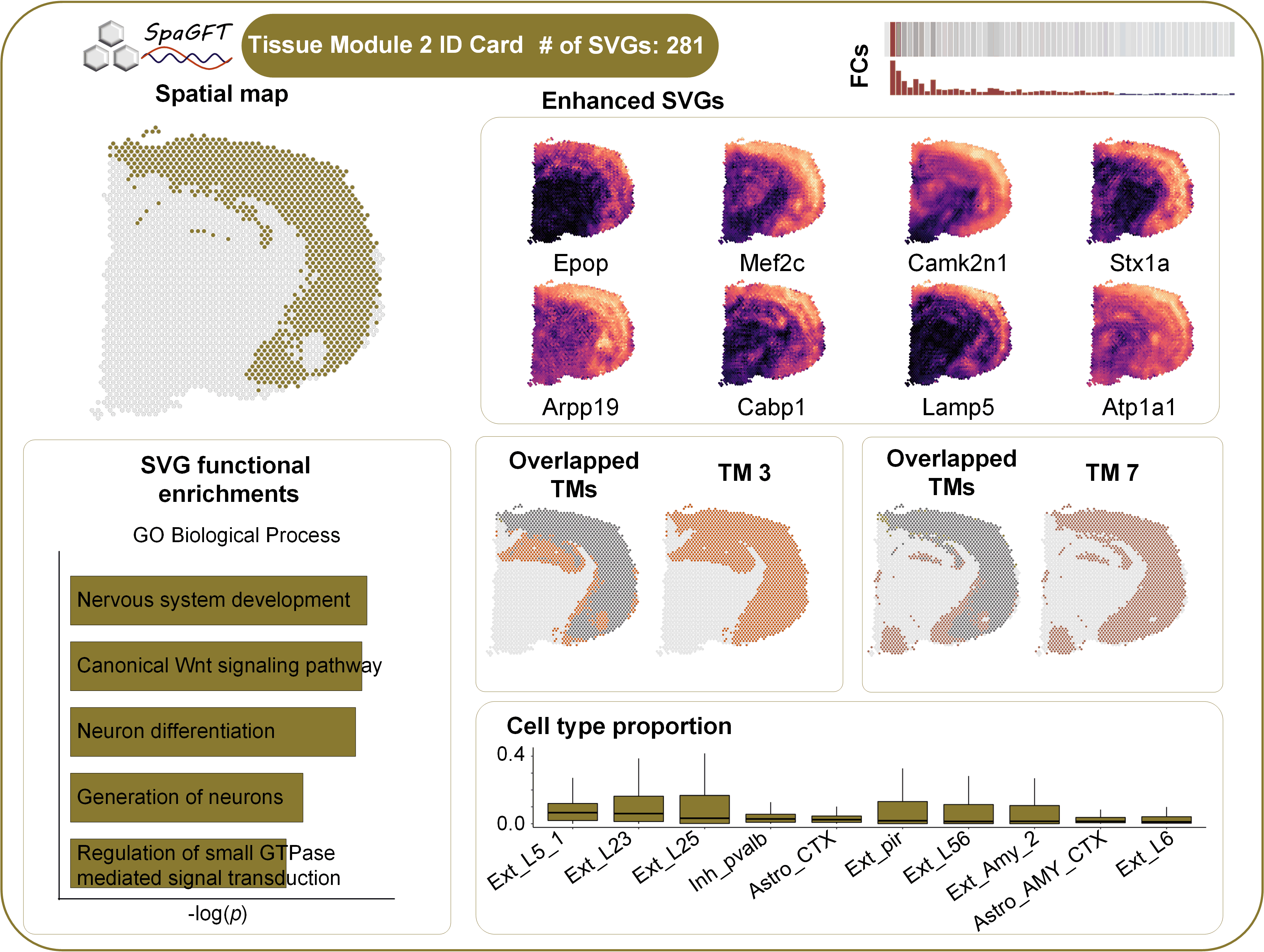

### TM3.tif

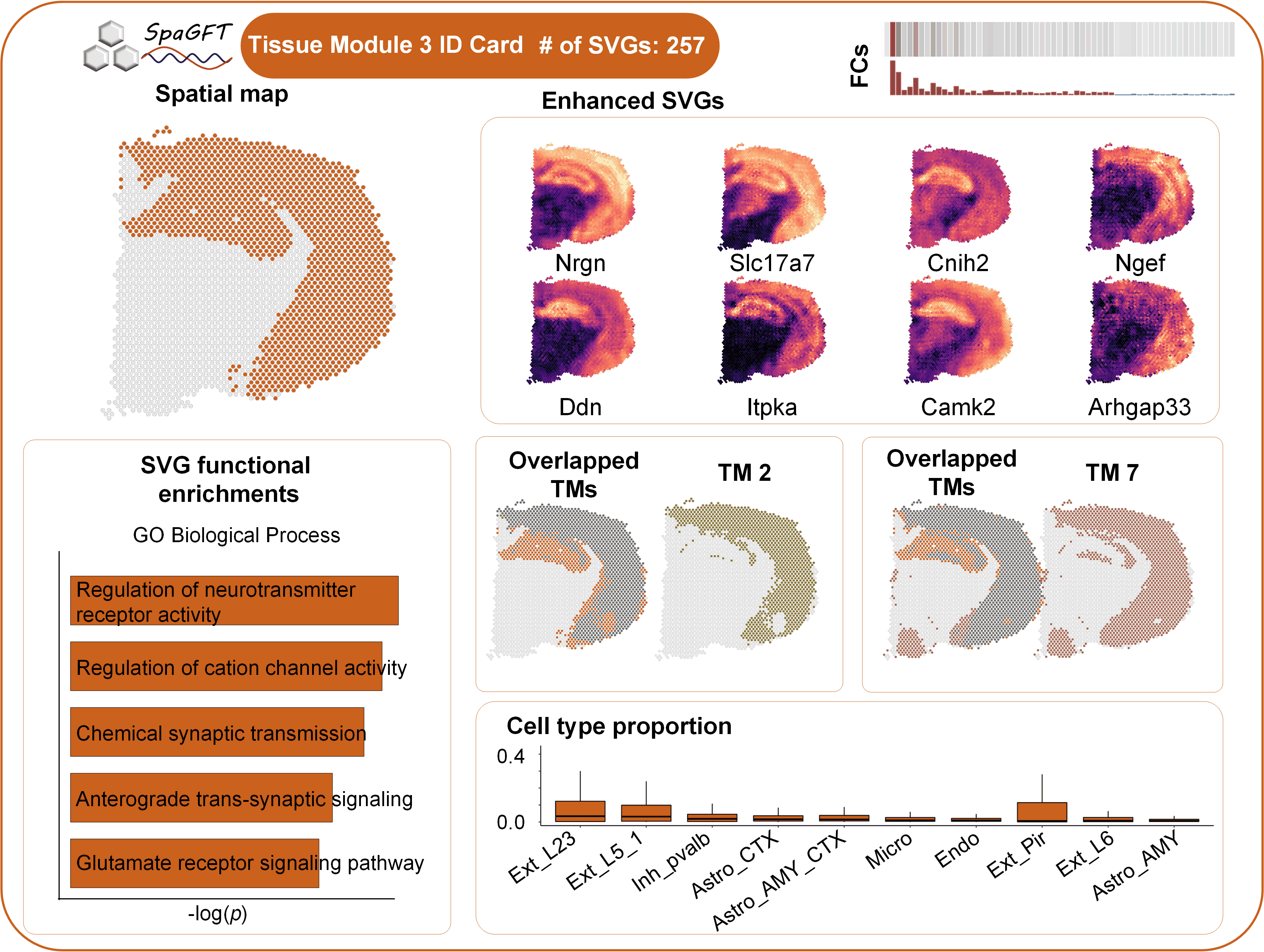

### TM4.tif

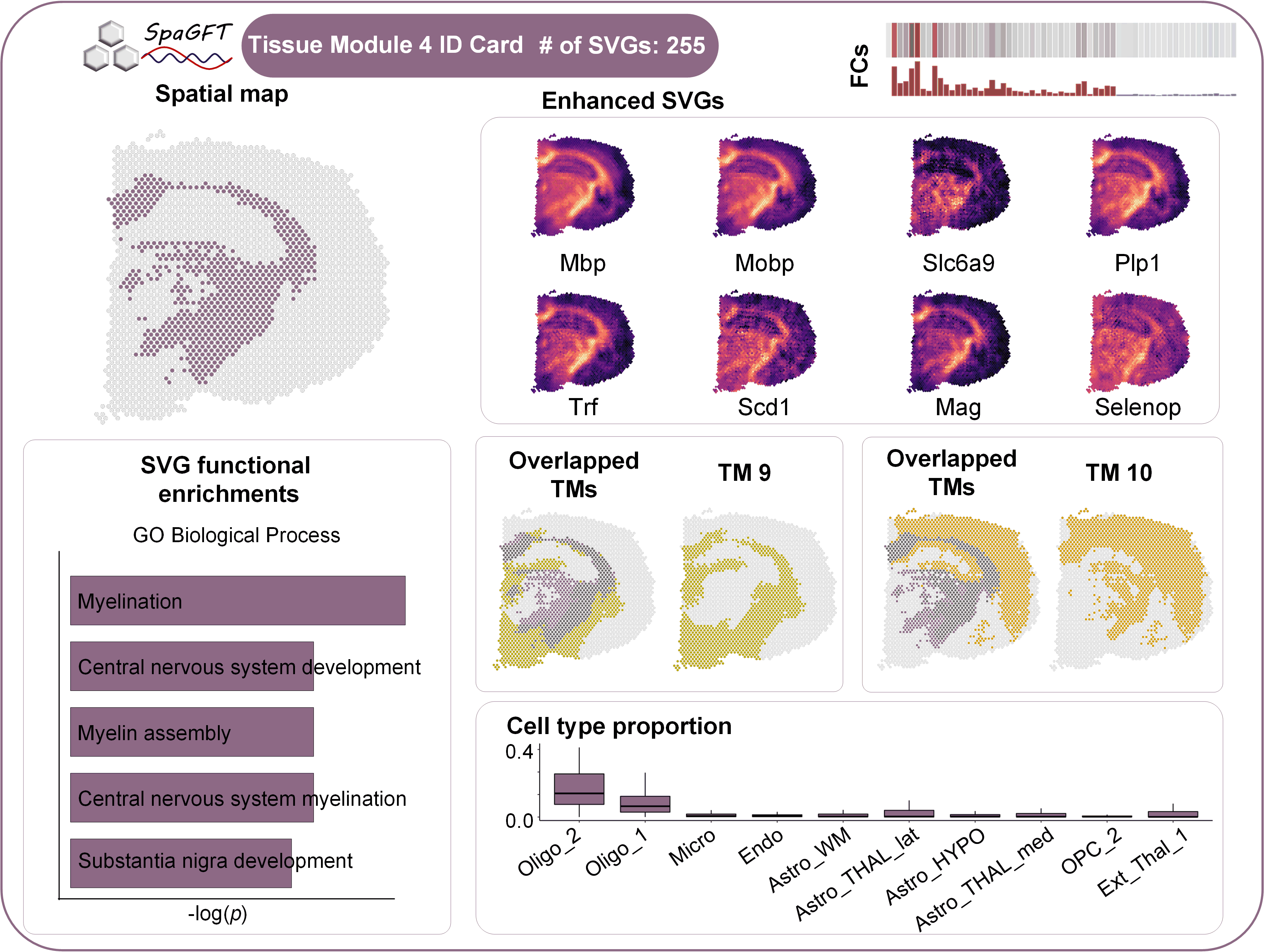

### TM5.tif

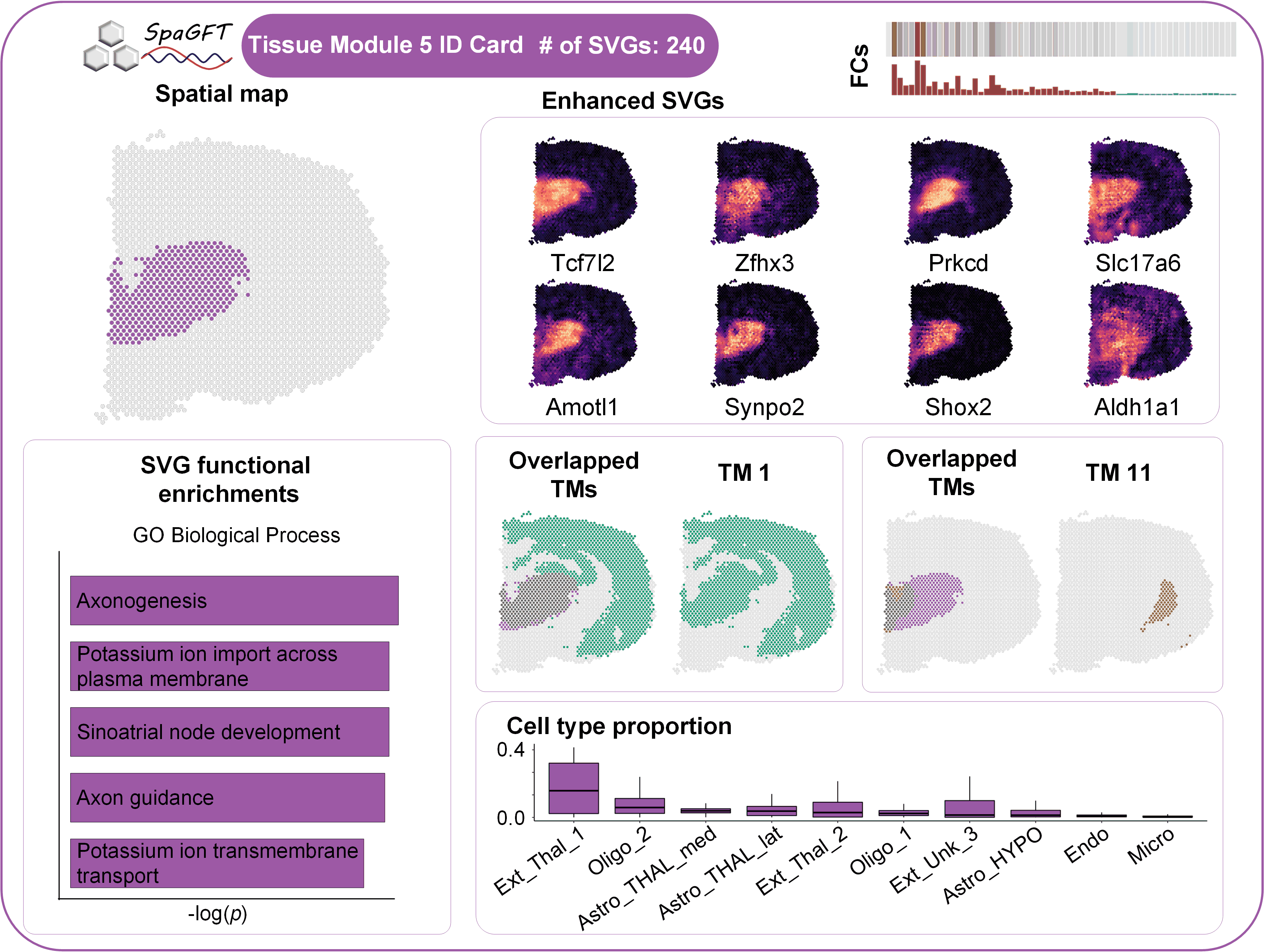

### TM7.tif

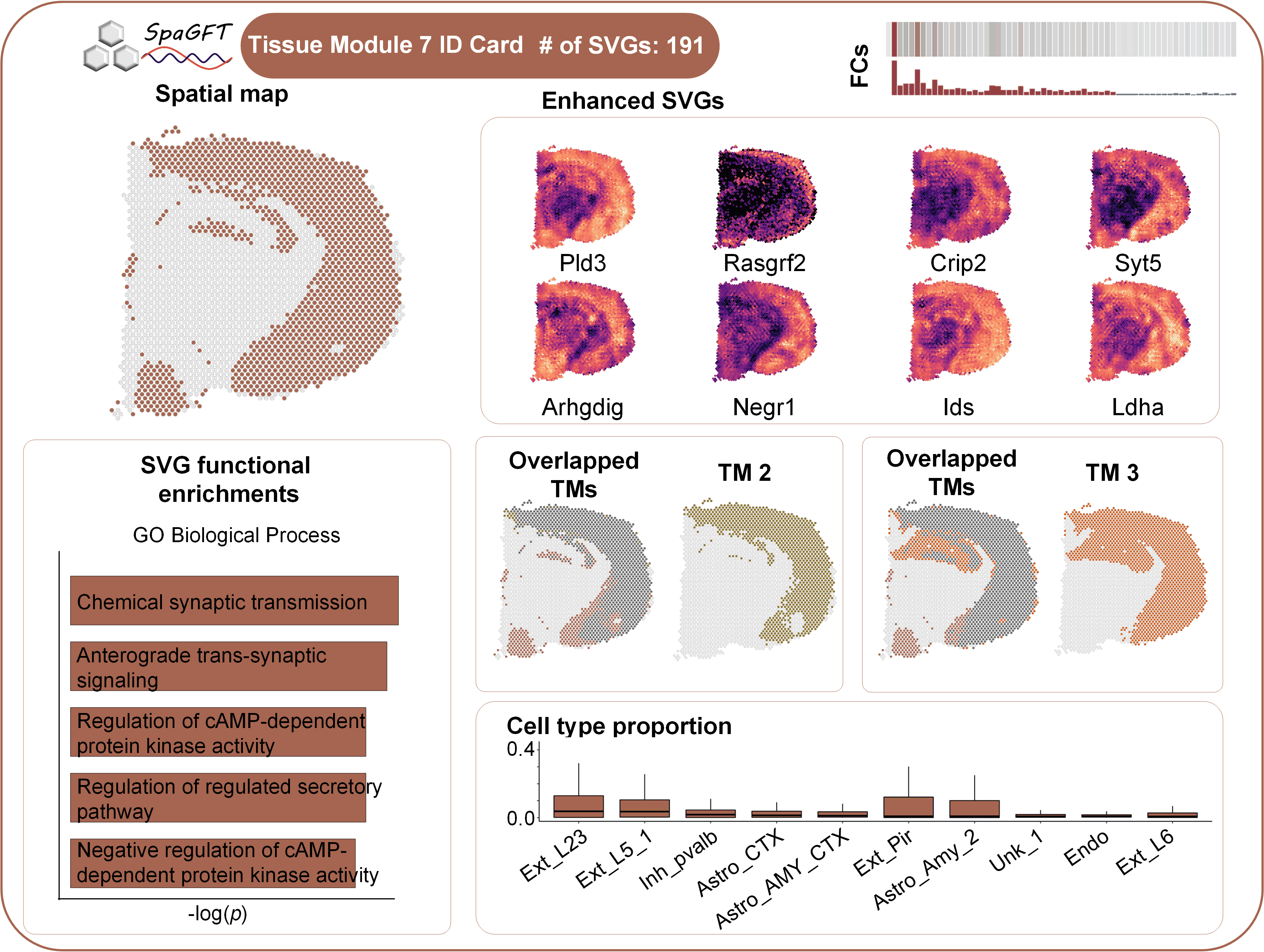

### TM8.tif

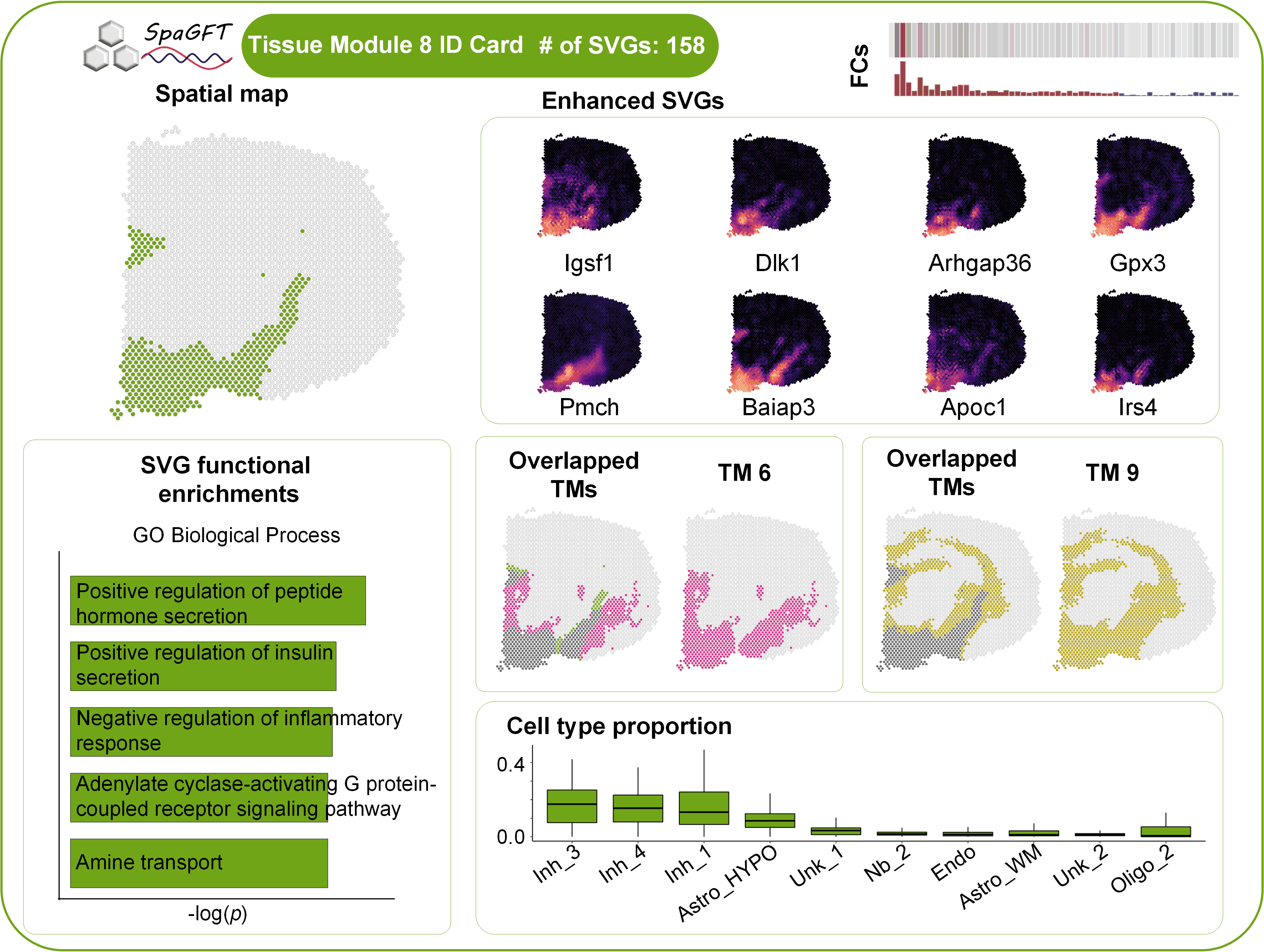

### TM9.tif

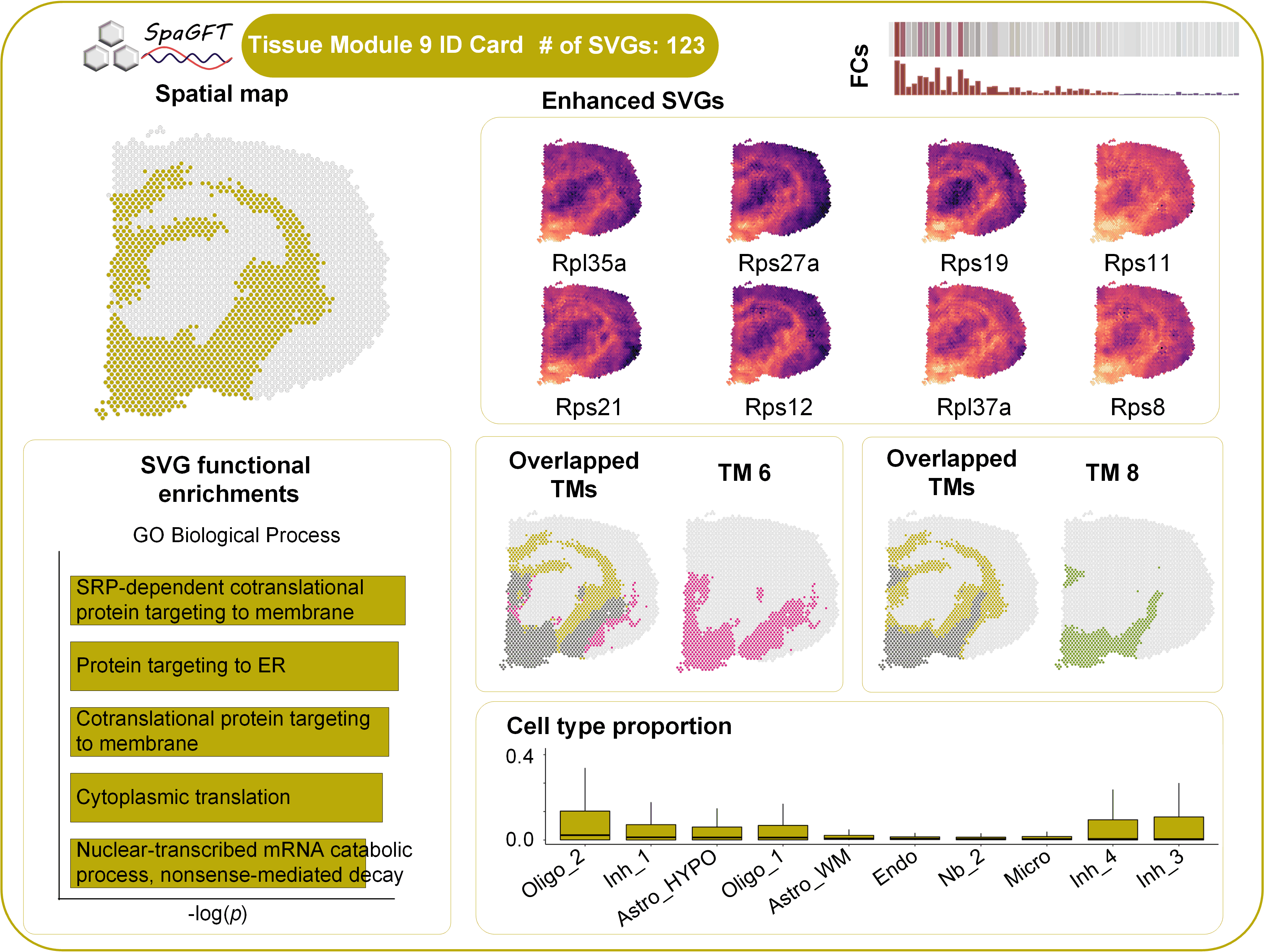

### TM10.tif

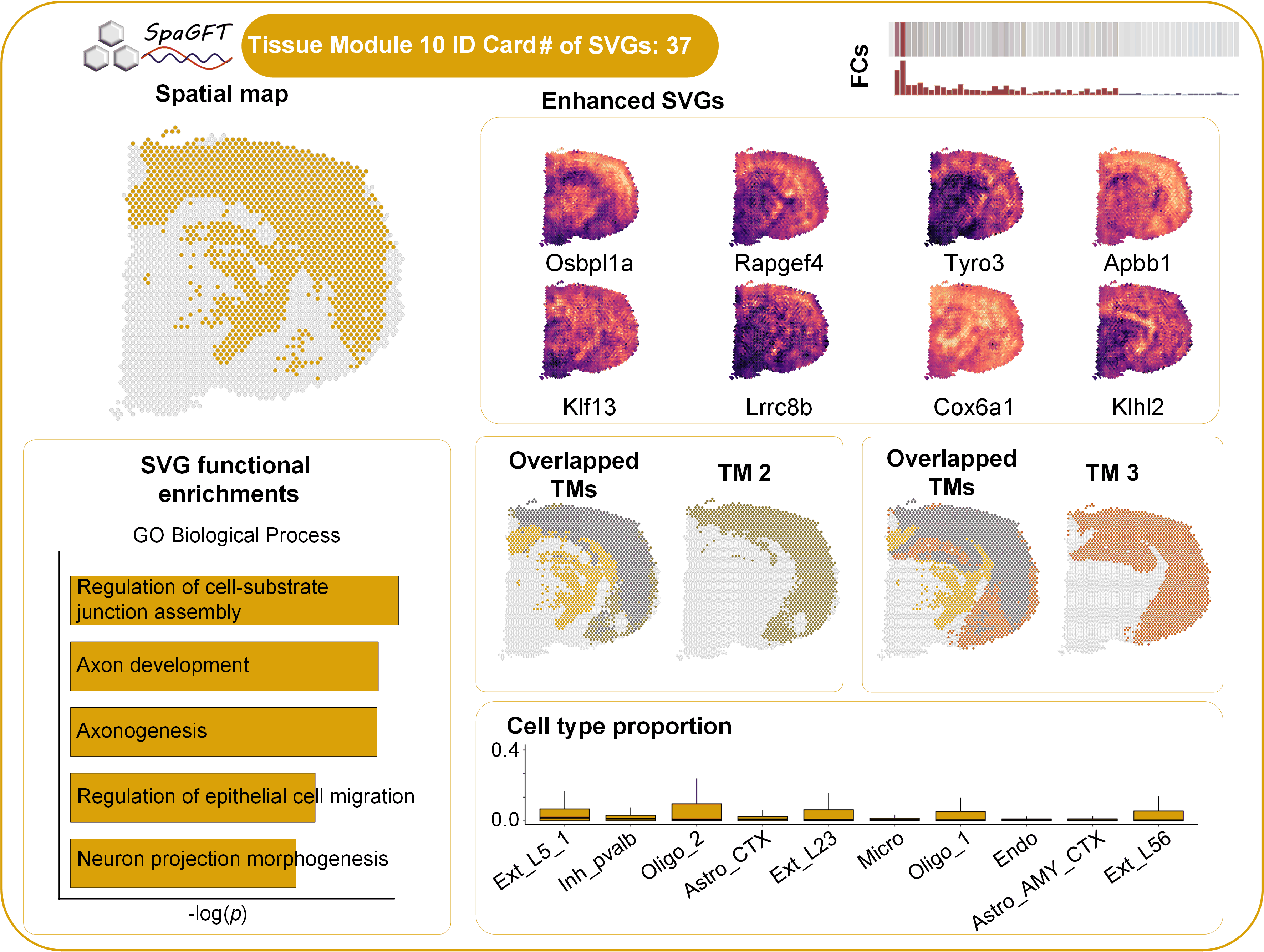

### TM11.tif

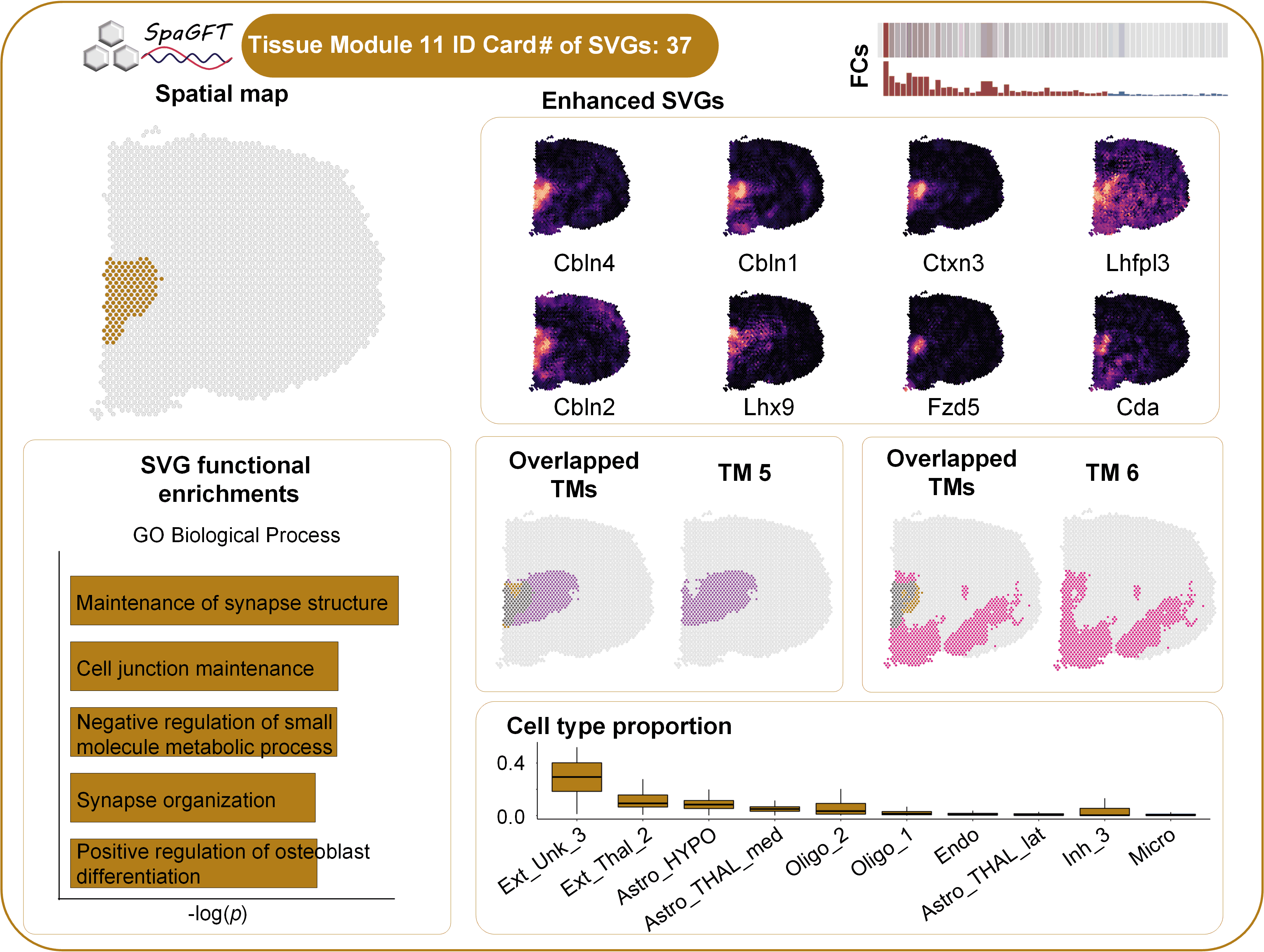

### TM12.tif

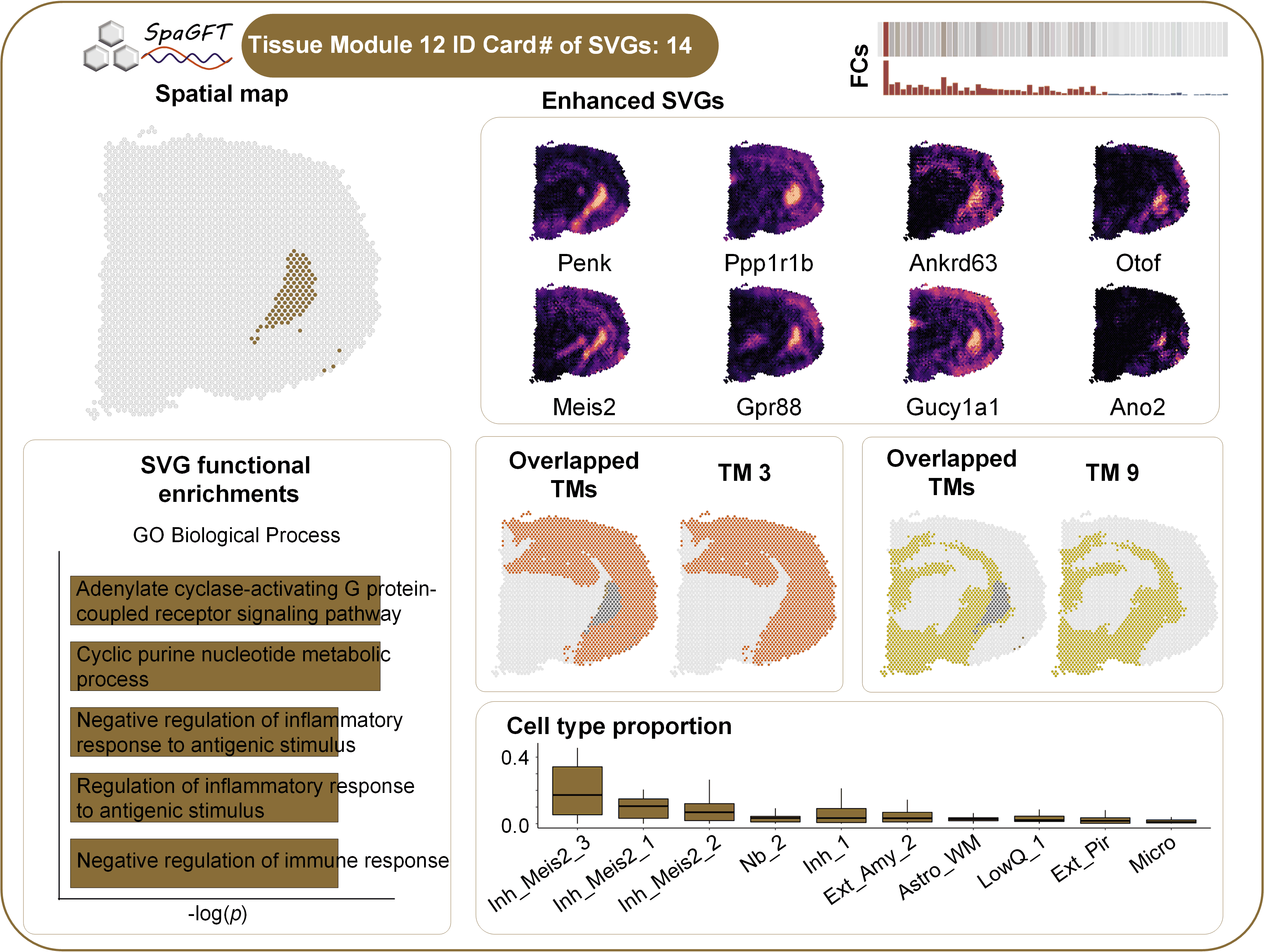
