## Supplementary material for "Spatial omics representation and functional tissue module inference using graph Fourier transform": Supp Data 2: crosstalk.min.js

```
!function o(u,a,l){function s(n,e){if(!a[n]){if(!u[n]){var t="function"==typeof require&&require;if(!e&&t)return t(n,!0);if(f)return f(n,!0);var r=new Error("Cannot find module '"+n+"'");throw r.code="MODULE_NOT_FOUND",r}var i=a[n]={exports:{}};u[n][0].call(i.exports,function(e){var t=u[n][1][e];return s(t||e)},i,i.exports,o,u,a,l)}return a[n].exports}for(var f="function"==typeof require&&require,e=0;e<l.length;e++)s(l[e]);return s}({1:[function(e,t,n){"use strict";Object.defineProperty(n,"__esModule",{value:!0});var r=function(){function r(e,t){for(var n=0;n<t.length;n++){var r=t[n];r.enumerable=r.enumerable||!1,r.configurable=!0,"value"in r&&(r.writable=!0),Object.defineProperty(e,r.key,r)}}return function(e,t,n){return t&&r(e.prototype,t),n&&r(e,n),e}}();var i=function(){function e(){!function(e,t){if(!(e instanceof t))throw new TypeError("Cannot call a class as a function")}(this,e),this._types={},this._seq=0}return r(e,[{key:"on",value:function(e,t){var n=this._types[e];n||(n=this._types[e]={});var r="sub"+this._seq++;return n[r]=t,r}},{key:"off",value:function(e,t){var n=this._types[e];if("function"==typeof t){for(var r in n)if(n.hasOwnProperty(r)&&n[r]===t)return delete n[r],r;return!1}if("string"==typeof t)return!(!n||!n[t])&&(delete n[t],t);throw new Error("Unexpected type for listener")}},{key:"trigger",value:function(e,t,n){var r=this._types[e];for(var i in r)r.hasOwnProperty(i)&&r[i].call(n,t)}}]),e}();n.default=i},{}],2:[function(e,t,n){"use strict";Object.defineProperty(n,"__esModule",{value:!0}),n.FilterHandle=void 0;var r=function(){function r(e,t){for(var n=0;n<t.length;n++){var r=t[n];r.enumerable=r.enumerable||!1,r.configurable=!0,"value"in r&&(r.writable=!0),Object.defineProperty(e,r.key,r)}}return function(e,t,n){return t&&r(e.prototype,t),n&&r(e,n),e}}(),i=l(e("./events")),o=l(e("./filterset")),u=l(e("./group")),a=function(e){{if(e&&e.__esModule)return e;var t={};if(null!=e)for(var n in e)Object.prototype.hasOwnProperty.call(e,n)&&(t[n]=e[n]);return t.default=e,t}}(e("./util"));function l(e){return e&&e.__esModule?e:{default:e}}var s=1;n.FilterHandle=function(){function n(e,t){!function(e,t){if(!(e instanceof t))throw new TypeError("Cannot call a class as a function")}(this,n),this._eventRelay=new i.default,this._emitter=new a.SubscriptionTracker(this._eventRelay),this._group=null,this._filterSet=null,this._filterVar=null,this._varOnChangeSub=null,this._extraInfo=a.extend({sender:this},t),this._id="filter"+s++,this.setGroup(e)}return r(n,[{key:"setGroup",value:function(e){var t,n,r=this;if(this._group!==e&&((this._group||e)&&(this._filterVar&&(this._filterVar.off("change",this._varOnChangeSub),this.clear(),this._varOnChangeSub=null,this._filterVar=null,this._filterSet=null),this._group=e))){e=(0,u.default)(e),this._filterSet=(t=e.var("filterset"),(n=t.get())||(n=new o.default,t.set(n)),n),this._filterVar=(0,u.default)(e).var("filter");var i=this._filterVar.on("change",function(e){r._eventRelay.trigger("change",e,r)});this._varOnChangeSub=i}}},{key:"_mergeExtraInfo",value:function(e){return a.extend({},this._extraInfo?this._extraInfo:null,e||null)}},{key:"close",value:function(){this._emitter.removeAllListeners(),this.clear(),this.setGroup(null)}},{key:"clear",value:function(e){this._filterSet&&(this._filterSet.clear(this._id),this._onChange(e))}},{key:"set",value:function(e,t){this._filterSet&&(this._filterSet.update(this._id,e),this._onChange(t))}},{key:"on",value:function(e,t){return this._emitter.on(e,t)}},{key:"off",value:function(e,t){return this._emitter.off(e,t)}},{key:"_onChange",value:function(e){this._filterSet&&this._filterVar.set(this._filterSet.value,this._mergeExtraInfo(e))}},{key:"filteredKeys",get:function(){return this._filterSet?this._filterSet.value:null}}]),n}()},{"./events":1,"./filterset":3,"./group":4,"./util":11}],3:[function(e,t,n){"use strict";Object.defineProperty(n,"__esModule",{value:!0});var r=function(){function r(e,t){for(var n=0;n<t.length;n++){var r=t[n];r.enumerable=r.enumerable||!1,r.configurable=!0,"value"in r&&(r.writable=!0),Object.defineProperty(e,r.key,r)}}return function(e,t,n){return t&&r(e.prototype,t),n&&r(e,n),e}}(),a=e("./util");function l(e,t){return e===t?0:e<t?-1:t<e?1:void 0}var i=function(){function e(){!function(e,t){if(!(e instanceof t))throw new TypeError("Cannot call a class as a function")}(this,e),this.reset()}return r(e,[{key:"reset",value:function(){this._handles={},this._keys={},this._value=null,this._activeHandles=0}},{key:"update",value:function(e,t){null!==t&&(t=t.slice(0)).sort(l);var n=(0,a.diffSortedLists)(this._handles[e],t),r=n.added,i=n.removed;this._handles[e]=t;for(var o=0;o<r.length;o++)this._keys[r[o]]=(this._keys[r[o]]||0)+1;for(var u=0;u<i.length;u++)this._keys[i[u]]--;this._updateValue(t)}},{key:"_updateValue",value:function(){var e=0<arguments.length&&void 0!==arguments[0]?arguments[0]:this._allKeys,t=Object.keys(this._handles).length;if(0===t)this._value=null;else{this._value=[];for(var n=0;n<e.length;n++){this._keys[e[n]]===t&&this._value.push(e[n])}}}},{key:"clear",value:function(e){if(void 0!==this._handles[e]){var t=this._handles[e];t||(t=[]);for(var n=0;n<t.length;n++)this._keys[t[n]]--;delete this._handles[e],this._updateValue()}}},{key:"value",get:function(){return this._value}},{key:"_allKeys",get:function(){var e=Object.keys(this._keys);return e.sort(l),e}}]),e}();n.default=i},{"./util":11}],4:[function(l,e,s){(function(e){"use strict";Object.defineProperty(s,"__esModule",{value:!0});var n=function(){function r(e,t){for(var n=0;n<t.length;n++){var r=t[n];r.enumerable=r.enumerable||!1,r.configurable=!0,"value"in r&&(r.writable=!0),Object.defineProperty(e,r.key,r)}}return function(e,t,n){return t&&r(e.prototype,t),n&&r(e,n),e}}(),r="function"==typeof Symbol&&"symbol"==typeof Symbol.iterator?function(e){return typeof e}:function(e){return e&&"function"==typeof Symbol&&e.constructor===Symbol&&e!==Symbol.prototype?"symbol":typeof e};s.default=function e(t){{if(t&&"string"==typeof t)return u.hasOwnProperty(t)||(u[t]=new a(t)),u[t];if("object"===(void 0===t?"undefined":r(t))&&t._vars&&t.var)return t;if(Array.isArray(t)&&1==t.length&&"string"==typeof t[0])return e(t[0]);throw new Error("Invalid groupName argument")}};var t,i=l("./var"),o=(t=i)&&t.__esModule?t:{default:t};e.__crosstalk_groups=e.__crosstalk_groups||{};var u=e.__crosstalk_groups;var a=function(){function t(e){!function(e,t){if(!(e instanceof t))throw new TypeError("Cannot call a class as a function")}(this,t),this.name=e,this._vars={}}return n(t,[{key:"var",value:function(e){if(!e||"string"!=typeof e)throw new Error("Invalid var name");return this._vars.hasOwnProperty(e)||(this._vars[e]=new o.default(this,e)),this._vars[e]}},{key:"has",value:function(e){if(!e||"string"!=typeof e)throw new Error("Invalid var name");return this._vars.hasOwnProperty(e)}}]),t}()}).call(this,"undefined"!=typeof global?global:"undefined"!=typeof self?self:"undefined"!=typeof window?window:{})},{"./var":12}],5:[function(f,e,c){(function(e){"use strict";Object.defineProperty(c,"__esModule",{value:!0});var t,n=f("./group"),r=(t=n)&&t.__esModule?t:{default:t},i=f("./selection"),o=f("./filter"),u=f("./input");f("./input_selectize"),f("./input_checkboxgroup"),f("./input_slider");var a=(0,r.default)("default");function l(e){return a.var(e)}e.Shiny&&e.Shiny.addCustomMessageHandler("update-client-value",function(e){"string"==typeof e.group?(0,r.default)(e.group).var(e.name).set(e.value):l(e.name).set(e.value)});var s={group:r.default,var:l,has:function(e){return a.has(e)},SelectionHandle:i.SelectionHandle,FilterHandle:o.FilterHandle,bind:u.bind};c.default=s,e.crosstalk=s}).call(this,"undefined"!=typeof global?global:"undefined"!=typeof self?self:"undefined"!=typeof window?window:{})},{"./filter":2,"./group":4,"./input":6,"./input_checkboxgroup":7,"./input_selectize":8,"./input_slider":9,"./selection":10}],6:[function(e,t,a){(function(t){"use strict";Object.defineProperty(a,"__esModule",{value:!0}),a.register=function(e){r[e.className]=e,t.document&&"complete"!==t.document.readyState?o(function(){n()}):t.document&&setTimeout(n,100)},a.bind=n;var o=t.jQuery,r={};function n(){Object.keys(r).forEach(function(e){var n=r[e];o("."+n.className).not(".crosstalk-input-bound").each(function(e,t){i(n,t)})})}function i(e,t){var n=o(t).find("script[type='application/json'][data-for='"+t.id.replace(/([!"#$%&'()*+,./:;<=>?@[\\\]^`{|}~])/g,"\\$1")+"']"),r=JSON.parse(n[0].innerText),i=e.factory(t,r);o(t).data("crosstalk-instance",i),o(t).addClass("crosstalk-input-bound")}if(t.Shiny){var e=new t.Shiny.InputBinding,u=t.jQuery;u.extend(e,{find:function(e){return u(e).find(".crosstalk-input")},initialize:function(e){var t,n;u(e).hasClass("crosstalk-input-bound")||(n=o(t=e),Object.keys(r).forEach(function(e){n.hasClass(e)&&!n.hasClass("crosstalk-input-bound")&&i(r[e],t)}))},getId:function(e){return e.id},getValue:function(e){},setValue:function(e,t){},receiveMessage:function(e,t){},subscribe:function(e,t){u(e).data("crosstalk-instance").resume()},unsubscribe:function(e){u(e).data("crosstalk-instance").suspend()}}),t.Shiny.inputBindings.register(e,"crosstalk.inputBinding")}}).call(this,"undefined"!=typeof global?global:"undefined"!=typeof self?self:"undefined"!=typeof window?window:{})},{}],7:[function(r,e,t){(function(e){"use strict";var t=function(e){{if(e&&e.__esModule)return e;var t={};if(null!=e)for(var n in e)Object.prototype.hasOwnProperty.call(e,n)&&(t[n]=e[n]);return t.default=e,t}}(r("./input")),n=r("./filter");var a=e.jQuery;t.register({className:"crosstalk-input-checkboxgroup",factory:function(e,r){var i=new n.FilterHandle(r.group),o=void 0,u=a(e);return u.on("change","input[type='checkbox']",function(){var e=u.find("input[type='checkbox']:checked");if(0===e.length)o=null,i.clear();else{var t={};e.each(function(){r.map[this.value].forEach(function(e){t[e]=!0})});var n=Object.keys(t);n.sort(),o=n,i.set(n)}}),{suspend:function(){i.clear()},resume:function(){o&&i.set(o)}}}})}).call(this,"undefined"!=typeof global?global:"undefined"!=typeof self?self:"undefined"!=typeof window?window:{})},{"./filter":2,"./input":6}],8:[function(r,e,t){(function(e){"use strict";var t=n(r("./input")),l=n(r("./util")),s=r("./filter");function n(e){if(e&&e.__esModule)return e;var t={};if(null!=e)for(var n in e)Object.prototype.hasOwnProperty.call(e,n)&&(t[n]=e[n]);return t.default=e,t}var f=e.jQuery;t.register({className:"crosstalk-input-select",factory:function(e,n){var t=l.dataframeToD3(n.items),r={options:[{value:"",label:"(All)"}].concat(t),valueField:"value",labelField:"label",searchField:"label"},i=f(e).find("select")[0],o=f(i).selectize(r)[0].selectize,u=new s.FilterHandle(n.group),a=void 0;return o.on("change",function(){if(0===o.items.length)a=null,u.clear();else{var t={};o.items.forEach(function(e){n.map[e].forEach(function(e){t[e]=!0})});var e=Object.keys(t);e.sort(),a=e,u.set(e)}}),{suspend:function(){u.clear()},resume:function(){a&&u.set(a)}}}})}).call(this,"undefined"!=typeof global?global:"undefined"!=typeof self?self:"undefined"!=typeof window?window:{})},{"./filter":2,"./input":6,"./util":11}],9:[function(n,e,t){(function(e){"use strict";var d=function(e,t){if(Array.isArray(e))return e;if(Symbol.iterator in Object(e))return function(e,t){var n=[],r=!0,i=!1,o=void 0;try{for(var u,a=e[Symbol.iterator]();!(r=(u=a.next()).done)&&(n.push(u.value),!t||n.length!==t);r=!0);}catch(e){i=!0,o=e}finally{try{!r&&a.return&&a.return()}finally{if(i)throw o}}return n}(e,t);throw new TypeError("Invalid attempt to destructure non-iterable instance")},t=function(e){{if(e&&e.__esModule)return e;var t={};if(null!=e)for(var n in e)Object.prototype.hasOwnProperty.call(e,n)&&(t[n]=e[n]);return t.default=e,t}}(n("./input")),a=n("./filter");var v=e.jQuery,p=e.strftime;function y(e,t){for(var n=e.toString();n.length<t;)n="0"+n;return n}t.register({className:"crosstalk-input-slider",factory:function(e,l){var s=new a.FilterHandle(l.group),t={},f=v(e).find("input"),n=f.data("data-type"),r=f.data("time-format"),i=f.data("round"),o=void 0;if("date"===n)o=p.utc(),t.prettify=function(e){return o(r,new Date(e))};else if("datetime"===n){var u=f.data("timezone");o=u?p.timezone(u):p,t.prettify=function(e){return o(r,new Date(e))}}else"number"===n&&void 0!==i&&(t.prettify=function(e){var t=Math.pow(10,i);return Math.round(e*t)/t});function c(){var e=f.data("ionRangeSlider").result,t=void 0,n=f.data("data-type");return t="date"===n?function(e){return(t=new Date(+e))instanceof Date?t.getUTCFullYear()+"-"+y(t.getUTCMonth()+1,2)+"-"+y(t.getUTCDate(),2):null;var t}:"datetime"===n?function(e){return+e/1e3}:function(e){return+e},"double"===f.data("ionRangeSlider").options.type?[t(e.from),t(e.to)]:t(e.from)}f.ionRangeSlider(t);var h=null;return f.on("change.crosstalkSliderInput",function(e){if(!f.data("updating")&&!f.data("animating")){for(var t=c(),n=d(t,2),r=n[0],i=n[1],o=[],u=0;u<l.values.length;u++){var a=l.values[u];r<=a&&a<=i&&o.push(l.keys[u])}o.sort(),s.set(o),h=o}}),{suspend:function(){s.clear()},resume:function(){h&&s.set(h)}}}})}).call(this,"undefined"!=typeof global?global:"undefined"!=typeof self?self:"undefined"!=typeof window?window:{})},{"./filter":2,"./input":6}],10:[function(e,t,n){"use strict";Object.defineProperty(n,"__esModule",{value:!0}),n.SelectionHandle=void 0;var r=function(){function r(e,t){for(var n=0;n<t.length;n++){var r=t[n];r.enumerable=r.enumerable||!1,r.configurable=!0,"value"in r&&(r.writable=!0),Object.defineProperty(e,r.key,r)}}return function(e,t,n){return t&&r(e.prototype,t),n&&r(e,n),e}}(),i=a(e("./events")),o=a(e("./group")),u=function(e){{if(e&&e.__esModule)return e;var t={};if(null!=e)for(var n in e)Object.prototype.hasOwnProperty.call(e,n)&&(t[n]=e[n]);return t.default=e,t}}(e("./util"));function a(e){return e&&e.__esModule?e:{default:e}}n.SelectionHandle=function(){function n(){var e=0<arguments.length&&void 0!==arguments[0]?arguments[0]:null,t=1<arguments.length&&void 0!==arguments[1]?arguments[1]:null;!function(e,t){if(!(e instanceof t))throw new TypeError("Cannot call a class as a function")}(this,n),this._eventRelay=new i.default,this._emitter=new u.SubscriptionTracker(this._eventRelay),this._group=null,this._var=null,this._varOnChangeSub=null,this._extraInfo=u.extend({sender:this},t),this.setGroup(e)}return r(n,[{key:"setGroup",value:function(e){var t=this;if(this._group!==e&&(this._group||e)&&(this._var&&(this._var.off("change",this._varOnChangeSub),this._var=null,this._varOnChangeSub=null),this._group=e)){this._var=(0,o.default)(e).var("selection");var n=this._var.on("change",function(e){t._eventRelay.trigger("change",e,t)});this._varOnChangeSub=n}}},{key:"_mergeExtraInfo",value:function(e){return u.extend({},this._extraInfo?this._extraInfo:null,e||null)}},{key:"set",value:function(e,t){this._var&&this._var.set(e,this._mergeExtraInfo(t))}},{key:"clear",value:function(e){this._var&&this.set(void 0,this._mergeExtraInfo(e))}},{key:"on",value:function(e,t){return this._emitter.on(e,t)}},{key:"off",value:function(e,t){return this._emitter.off(e,t)}},{key:"close",value:function(){this._emitter.removeAllListeners(),this.setGroup(null)}},{key:"value",get:function(){return this._var?this._var.get():null}}]),n}()},{"./events":1,"./group":4,"./util":11}],11:[function(e,t,n){"use strict";Object.defineProperty(n,"__esModule",{value:!0});var r=function(){function r(e,t){for(var n=0;n<t.length;n++){var r=t[n];r.enumerable=r.enumerable||!1,r.configurable=!0,"value"in r&&(r.writable=!0),Object.defineProperty(e,r.key,r)}}return function(e,t,n){return t&&r(e.prototype,t),n&&r(e,n),e}}(),l="function"==typeof Symbol&&"symbol"==typeof Symbol.iterator?function(e){return typeof e}:function(e){return e&&"function"==typeof Symbol&&e.constructor===Symbol&&e!==Symbol.prototype?"symbol":typeof e};function u(e){for(var t=1;t<e.length;t++)if(e[t]<=e[t-1])throw new Error("List is not sorted or contains duplicate")}n.extend=function(e){for(var t=arguments.length,n=Array(1<t?t-1:0),r=1;r<t;r++)n[r-1]=arguments[r];for(var i=0;i<n.length;i++){var o=n[i];if(null!=o)for(var u in o)o.hasOwnProperty(u)&&(e[u]=o[u])}return e},n.checkSorted=u,n.diffSortedLists=function(e,t){var n=0,r=0;e||(e=[]);t||(t=[]);var i=[],o=[];u(e),u(t);for(;n<e.length&&r<t.length;)e[n]===t[r]?(n++,r++):e[n]<t[r]?i.push(e[n++]):o.push(t[r++]);n<e.length&&(i=i.concat(e.slice(n)));r<t.length&&(o=o.concat(t.slice(r)));return{removed:i,added:o}},n.dataframeToD3=function(e){var t=[],n=void 0;for(var r in e){if(e.hasOwnProperty(r)&&t.push(r),"object"!==l(e[r])||void 0===e[r].length)throw new Error("All fields must be arrays");if(void 0!==n&&n!==e[r].length)throw new Error("All fields must be arrays of the same length");n=e[r].length}for(var i=[],o=void 0,u=0;u<n;u++){o={};for(var a=0;a<t.length;a++)o[t[a]]=e[t[a]][u];i.push(o)}return i};n.SubscriptionTracker=function(){function t(e){!function(e,t){if(!(e instanceof t))throw new TypeError("Cannot call a class as a function")}(this,t),this._emitter=e,this._subs={}}return r(t,[{key:"on",value:function(e,t){var n=this._emitter.on(e,t);return this._subs[n]=e,n}},{key:"off",value:function(e,t){var n=this._emitter.off(e,t);return n&&delete this._subs[n],n}},{key:"removeAllListeners",value:function(){var t=this,n=this._subs;this._subs={},Object.keys(n).forEach(function(e){t._emitter.off(n[e],e)})}}]),t}()},{}],12:[function(a,e,l){(function(o){"use strict";Object.defineProperty(l,"__esModule",{value:!0});var e,u="function"==typeof Symbol&&"symbol"==typeof Symbol.iterator?function(e){return typeof e}:function(e){return e&&"function"==typeof Symbol&&e.constructor===Symbol&&e!==Symbol.prototype?"symbol":typeof e},t=function(){function r(e,t){for(var n=0;n<t.length;n++){var r=t[n];r.enumerable=r.enumerable||!1,r.configurable=!0,"value"in r&&(r.writable=!0),Object.defineProperty(e,r.key,r)}}return function(e,t,n){return t&&r(e.prototype,t),n&&r(e,n),e}}(),n=a("./events"),i=(e=n)&&e.__esModule?e:{default:e};var r=function(){function r(e,t,n){!function(e,t){if(!(e instanceof t))throw new TypeError("Cannot call a class as a function")}(this,r),this._group=e,this._name=t,this._value=n,this._events=new i.default}return t(r,[{key:"get",value:function(){return this._value}},{key:"set",value:function(e,t){if(this._value!==e){var n=this._value;this._value=e;var r={};if(t&&"object"===(void 0===t?"undefined":u(t)))for(var i in t)t.hasOwnProperty(i)&&(r[i]=t[i]);r.oldValue=n,r.value=e,this._events.trigger("change",r,this),o.Shiny&&o.Shiny.onInputChange&&o.Shiny.onInputChange(".clientValue-"+(null!==this._group.name?this._group.name+"-":"")+this._name,void 0===e?null:e)}}},{key:"on",value:function(e,t){return this._events.on(e,t)}},{key:"off",value:function(e,t){return this._events.off(e,t)}}]),r}();l.default=r}).call(this,"undefined"!=typeof global?global:"undefined"!=typeof self?self:"undefined"!=typeof window?window:{})},{"./events":1}]},{},[5]);
//# sourceMappingURL=crosstalk.min.js.map
```
