## Supplementary material for "Spatial omics representation and functional tissue module inference using graph Fourier transform": Supp Data 2: plotly-latest.min.js

```
/**
* plotly.js v2.5.1
* Copyright 2012-2021, Plotly, Inc.
* All rights reserved.
* Licensed under the MIT license
*/
!function(t){if("object"==typeof exports&&"undefined"!=typeof module)module.exports=t();else if("function"==typeof define&&define.amd)define([],t);else{("undefined"!=typeof window?window:"undefined"!=typeof global?global:"undefined"!=typeof self?self:this).Plotly=t()}}((function(){return function t(e,r,n){function i(o,s){if(!r[o]){if(!e[o]){var l="function"==typeof require&&require;if(!s&&l)return l(o,!0);if(a)return a(o,!0);var c=new Error("Cannot find module '"+o+"'");throw c.code="MODULE_NOT_FOUND",c}var u=r[o]={exports:{}};e[o][0].call(u.exports,(function(t){return i(e[o][1][t]||t)}),u,u.exports,t,e,r,n)}return r[o].exports}for(var a="function"==typeof require&&require,o=0;o<n.length;o++)i(n[o]);return i}({1:[function(t,e,r){"use strict";var n=t("../src/lib"),i={"X,X div":'direction:ltr;font-family:"Open Sans",verdana,arial,sans-serif;margin:0;padding:0;',"X input,X button":'font-family:"Open Sans",verdana,arial,sans-serif;',"X input:focus,X button:focus":"outline:none;","X a":"text-decoration:none;","X a:hover":"text-decoration:none;","X .crisp":"shape-rendering:crispEdges;","X .user-select-none":"-webkit-user-select:none;-moz-user-select:none;-ms-user-select:none;-o-user-select:none;user-select:none;","X svg":"overflow:hidden;","X svg a":"fill:#447adb;","X svg a:hover":"fill:#3c6dc5;","X .main-svg":"position:absolute;top:0;left:0;pointer-events:none;","X .main-svg .draglayer":"pointer-events:all;","X .cursor-default":"cursor:default;","X .cursor-pointer":"cursor:pointer;","X .cursor-crosshair":"cursor:crosshair;","X .cursor-move":"cursor:move;","X .cursor-col-resize":"cursor:col-resize;","X .cursor-row-resize":"cursor:row-resize;","X .cursor-ns-resize":"cursor:ns-resize;","X .cursor-ew-resize":"cursor:ew-resize;","X .cursor-sw-resize":"cursor:sw-resize;","X .cursor-s-resize":"cursor:s-resize;","X .cursor-se-resize":"cursor:se-resize;","X .cursor-w-resize":"cursor:w-resize;","X .cursor-e-resize":"cursor:e-resize;","X .cursor-nw-resize":"cursor:nw-resize;","X .cursor-n-resize":"cursor:n-resize;","X .cursor-ne-resize":"cursor:ne-resize;","X .cursor-grab":"cursor:-webkit-grab;cursor:grab;","X .modebar":"position:absolute;top:2px;right:2px;","X .ease-bg":"-webkit-transition:background-color .3s ease 0s;-moz-transition:background-color .3s ease 0s;-ms-transition:background-color .3s ease 0s;-o-transition:background-color .3s ease 0s;transition:background-color .3s ease 0s;","X .modebar--hover>:not(.watermark)":"opacity:0;-webkit-transition:opacity .3s ease 0s;-moz-transition:opacity .3s ease 0s;-ms-transition:opacity .3s ease 0s;-o-transition:opacity .3s ease 0s;transition:opacity .3s ease 0s;","X:hover .modebar--hover .modebar-group":"opacity:1;","X .modebar-group":"float:left;display:inline-block;box-sizing:border-box;padding-left:8px;position:relative;vertical-align:middle;white-space:nowrap;","X .modebar-btn":"position:relative;font-size:16px;padding:3px 4px;height:22px;cursor:pointer;line-height:normal;box-sizing:border-box;","X .modebar-btn svg":"position:relative;top:2px;","X .modebar.vertical":"display:flex;flex-direction:column;flex-wrap:wrap;align-content:flex-end;max-height:100%;","X .modebar.vertical svg":"top:-1px;","X .modebar.vertical .modebar-group":"display:block;float:none;padding-left:0px;padding-bottom:8px;","X .modebar.vertical .modebar-group .modebar-btn":"display:block;text-align:center;","X [data-title]:before,X [data-title]:after":"position:absolute;-webkit-transform:translate3d(0, 0, 0);-moz-transform:translate3d(0, 0, 0);-ms-transform:translate3d(0, 0, 0);-o-transform:translate3d(0, 0, 0);transform:translate3d(0, 0, 0);display:none;opacity:0;z-index:1001;pointer-events:none;top:110%;right:50%;","X [data-title]:hover:before,X [data-title]:hover:after":"display:block;opacity:1;","X [data-title]:before":'content:"";position:absolute;background:transparent;border:6px solid transparent;z-index:1002;margin-top:-12px;border-bottom-color:#69738a;margin-right:-6px;',"X [data-title]:after":"content:attr(data-title);background:#69738a;color:#fff;padding:8px 10px;font-size:12px;line-height:12px;white-space:nowrap;margin-right:-18px;border-radius:2px;","X .vertical [data-title]:before,X .vertical [data-title]:after":"top:0%;right:200%;","X .vertical [data-title]:before":"border:6px solid transparent;border-left-color:#69738a;margin-top:8px;margin-right:-30px;","X .select-outline":"fill:none;stroke-width:1;shape-rendering:crispEdges;","X .select-outline-1":"stroke:#fff;","X .select-outline-2":"stroke:#000;stroke-dasharray:2px 2px;",Y:'font-family:"Open Sans",verdana,arial,sans-serif;position:fixed;top:50px;right:20px;z-index:10000;font-size:10pt;max-width:180px;',"Y p":"margin:0;","Y .notifier-note":"min-width:180px;max-width:250px;border:1px solid #fff;z-index:3000;margin:0;background-color:#8c97af;background-color:rgba(140,151,175,.9);color:#fff;padding:10px;overflow-wrap:break-word;word-wrap:break-word;-ms-hyphens:auto;-webkit-hyphens:auto;hyphens:auto;","Y .notifier-close":"color:#fff;opacity:.8;float:right;padding:0 5px;background:none;border:none;font-size:20px;font-weight:bold;line-height:20px;","Y .notifier-close:hover":"color:#444;text-decoration:none;cursor:pointer;"};for(var a in i){var o=a.replace(/^,/," ,").replace(/X/g,".js-plotly-plot .plotly").replace(/Y/g,".plotly-notifier");n.addStyleRule(o,i[a])}},{"../src/lib":776}],2:[function(t,e,r){"use strict";e.exports=t("../src/transforms/aggregate")},{"../src/transforms/aggregate":1372}],3:[function(t,e,r){"use strict";e.exports=t("../src/traces/bar")},{"../src/traces/bar":922}],4:[function(t,e,r){"use strict";e.exports=t("../src/traces/barpolar")},{"../src/traces/barpolar":935}],5:[function(t,e,r){"use strict";e.exports=t("../src/traces/box")},{"../src/traces/box":945}],6:[function(t,e,r){"use strict";e.exports=t("../src/components/calendars")},{"../src/components/calendars":637}],7:[function(t,e,r){"use strict";e.exports=t("../src/traces/candlestick")},{"../src/traces/candlestick":954}],8:[function(t,e,r){"use strict";e.exports=t("../src/traces/carpet")},{"../src/traces/carpet":973}],9:[function(t,e,r){"use strict";e.exports=t("../src/traces/choropleth")},{"../src/traces/choropleth":987}],10:[function(t,e,r){"use strict";e.exports=t("../src/traces/choroplethmapbox")},{"../src/traces/choroplethmapbox":994}],11:[function(t,e,r){"use strict";e.exports=t("../src/traces/cone")},{"../src/traces/cone":1e3}],12:[function(t,e,r){"use strict";e.exports=t("../src/traces/contour")},{"../src/traces/contour":1015}],13:[function(t,e,r){"use strict";e.exports=t("../src/traces/contourcarpet")},{"../src/traces/contourcarpet":1026}],14:[function(t,e,r){"use strict";e.exports=t("../src/core")},{"../src/core":754}],15:[function(t,e,r){"use strict";e.exports=t("../src/traces/densitymapbox")},{"../src/traces/densitymapbox":1034}],16:[function(t,e,r){"use strict";e.exports=t("../src/transforms/filter")},{"../src/transforms/filter":1373}],17:[function(t,e,r){"use strict";e.exports=t("../src/traces/funnel")},{"../src/traces/funnel":1044}],18:[function(t,e,r){"use strict";e.exports=t("../src/traces/funnelarea")},{"../src/traces/funnelarea":1053}],19:[function(t,e,r){"use strict";e.exports=t("../src/transforms/groupby")},{"../src/transforms/groupby":1374}],20:[function(t,e,r){"use strict";e.exports=t("../src/traces/heatmap")},{"../src/traces/heatmap":1066}],21:[function(t,e,r){"use strict";e.exports=t("../src/traces/heatmapgl")},{"../src/traces/heatmapgl":1076}],22:[function(t,e,r){"use strict";e.exports=t("../src/traces/histogram")},{"../src/traces/histogram":1088}],23:[function(t,e,r){"use strict";e.exports=t("../src/traces/histogram2d")},{"../src/traces/histogram2d":1094}],24:[function(t,e,r){"use strict";e.exports=t("../src/traces/histogram2dcontour")},{"../src/traces/histogram2dcontour":1098}],25:[function(t,e,r){"use strict";e.exports=t("../src/traces/icicle")},{"../src/traces/icicle":1104}],26:[function(t,e,r){"use strict";e.exports=t("../src/traces/image")},{"../src/traces/image":1117}],27:[function(t,e,r){"use strict";var n=t("./core");n.register([t("./bar"),t("./box"),t("./heatmap"),t("./histogram"),t("./histogram2d"),t("./histogram2dcontour"),t("./contour"),t("./scatterternary"),t("./violin"),t("./funnel"),t("./waterfall"),t("./image"),t("./pie"),t("./sunburst"),t("./treemap"),t("./icicle"),t("./funnelarea"),t("./scatter3d"),t("./surface"),t("./isosurface"),t("./volume"),t("./mesh3d"),t("./cone"),t("./streamtube"),t("./scattergeo"),t("./choropleth"),t("./scattergl"),t("./splom"),t("./pointcloud"),t("./heatmapgl"),t("./parcoords"),t("./parcats"),t("./scattermapbox"),t("./choroplethmapbox"),t("./densitymapbox"),t("./sankey"),t("./indicator"),t("./table"),t("./carpet"),t("./scattercarpet"),t("./contourcarpet"),t("./ohlc"),t("./candlestick"),t("./scatterpolar"),t("./scatterpolargl"),t("./barpolar"),t("./aggregate"),t("./filter"),t("./groupby"),t("./sort"),t("./calendars")]),e.exports=n},{"./aggregate":2,"./bar":3,"./barpolar":4,"./box":5,"./calendars":6,"./candlestick":7,"./carpet":8,"./choropleth":9,"./choroplethmapbox":10,"./cone":11,"./contour":12,"./contourcarpet":13,"./core":14,"./densitymapbox":15,"./filter":16,"./funnel":17,"./funnelarea":18,"./groupby":19,"./heatmap":20,"./heatmapgl":21,"./histogram":22,"./histogram2d":23,"./histogram2dcontour":24,"./icicle":25,"./image":26,"./indicator":28,"./isosurface":29,"./mesh3d":30,"./ohlc":31,"./parcats":32,"./parcoords":33,"./pie":34,"./pointcloud":35,"./sankey":36,"./scatter3d":37,"./scattercarpet":38,"./scattergeo":39,"./scattergl":40,"./scattermapbox":41,"./scatterpolar":42,"./scatterpolargl":43,"./scatterternary":44,"./sort":45,"./splom":46,"./streamtube":47,"./sunburst":48,"./surface":49,"./table":50,"./treemap":51,"./violin":52,"./volume":53,"./waterfall":54}],28:[function(t,e,r){"use strict";e.exports=t("../src/traces/indicator")},{"../src/traces/indicator":1125}],29:[function(t,e,r){"use strict";e.exports=t("../src/traces/isosurface")},{"../src/traces/isosurface":1131}],30:[function(t,e,r){"use strict";e.exports=t("../src/traces/mesh3d")},{"../src/traces/mesh3d":1136}],31:[function(t,e,r){"use strict";e.exports=t("../src/traces/ohlc")},{"../src/traces/ohlc":1141}],32:[function(t,e,r){"use strict";e.exports=t("../src/traces/parcats")},{"../src/traces/parcats":1150}],33:[function(t,e,r){"use strict";e.exports=t("../src/traces/parcoords")},{"../src/traces/parcoords":1160}],34:[function(t,e,r){"use strict";e.exports=t("../src/traces/pie")},{"../src/traces/pie":1171}],35:[function(t,e,r){"use strict";e.exports=t("../src/traces/pointcloud")},{"../src/traces/pointcloud":1180}],36:[function(t,e,r){"use strict";e.exports=t("../src/traces/sankey")},{"../src/traces/sankey":1186}],37:[function(t,e,r){"use strict";e.exports=t("../src/traces/scatter3d")},{"../src/traces/scatter3d":1224}],38:[function(t,e,r){"use strict";e.exports=t("../src/traces/scattercarpet")},{"../src/traces/scattercarpet":1231}],39:[function(t,e,r){"use strict";e.exports=t("../src/traces/scattergeo")},{"../src/traces/scattergeo":1239}],40:[function(t,e,r){"use strict";e.exports=t("../src/traces/scattergl")},{"../src/traces/scattergl":1252}],41:[function(t,e,r){"use strict";e.exports=t("../src/traces/scattermapbox")},{"../src/traces/scattermapbox":1262}],42:[function(t,e,r){"use strict";e.exports=t("../src/traces/scatterpolar")},{"../src/traces/scatterpolar":1270}],43:[function(t,e,r){"use strict";e.exports=t("../src/traces/scatterpolargl")},{"../src/traces/scatterpolargl":1277}],44:[function(t,e,r){"use strict";e.exports=t("../src/traces/scatterternary")},{"../src/traces/scatterternary":1285}],45:[function(t,e,r){"use strict";e.exports=t("../src/transforms/sort")},{"../src/transforms/sort":1376}],46:[function(t,e,r){"use strict";e.exports=t("../src/traces/splom")},{"../src/traces/splom":1294}],47:[function(t,e,r){"use strict";e.exports=t("../src/traces/streamtube")},{"../src/traces/streamtube":1302}],48:[function(t,e,r){"use strict";e.exports=t("../src/traces/sunburst")},{"../src/traces/sunburst":1310}],49:[function(t,e,r){"use strict";e.exports=t("../src/traces/surface")},{"../src/traces/surface":1319}],50:[function(t,e,r){"use strict";e.exports=t("../src/traces/table")},{"../src/traces/table":1327}],51:[function(t,e,r){"use strict";e.exports=t("../src/traces/treemap")},{"../src/traces/treemap":1338}],52:[function(t,e,r){"use strict";e.exports=t("../src/traces/violin")},{"../src/traces/violin":1351}],53:[function(t,e,r){"use strict";e.exports=t("../src/traces/volume")},{"../src/traces/volume":1359}],54:[function(t,e,r){"use strict";e.exports=t("../src/traces/waterfall")},{"../src/traces/waterfall":1367}],55:[function(t,e,r){"use strict";e.exports=function(t){var e=(t=t||{}).eye||[0,0,1],r=t.center||[0,0,0],s=t.up||[0,1,0],l=t.distanceLimits||[0,1/0],c=t.mode||"turntable",u=n(),f=i(),h=a();return u.setDistanceLimits(l[0],l[1]),u.lookAt(0,e,r,s),f.setDistanceLimits(l[0],l[1]),f.lookAt(0,e,r,s),h.setDistanceLimits(l[0],l[1]),h.lookAt(0,e,r,s),new o({turntable:u,orbit:f,matrix:h},c)};var n=t("turntable-camera-controller"),i=t("orbit-camera-controller"),a=t("matrix-camera-controller");function o(t,e){this._controllerNames=Object.keys(t),this._controllerList=this._controllerNames.map((function(e){return t[e]})),this._mode=e,this._active=t[e],this._active||(this._mode="turntable",this._active=t.turntable),this.modes=this._controllerNames,this.computedMatrix=this._active.computedMatrix,this.computedEye=this._active.computedEye,this.computedUp=this._active.computedUp,this.computedCenter=this._active.computedCenter,this.computedRadius=this._active.computedRadius}var s=o.prototype;s.flush=function(t){for(var e=this._controllerList,r=0;r<e.length;++r)e[r].flush(t)},s.idle=function(t){for(var e=this._controllerList,r=0;r<e.length;++r)e[r].idle(t)},s.lookAt=function(t,e,r,n){for(var i=this._controllerList,a=0;a<i.length;++a)i[a].lookAt(t,e,r,n)},s.rotate=function(t,e,r,n){for(var i=this._controllerList,a=0;a<i.length;++a)i[a].rotate(t,e,r,n)},s.pan=function(t,e,r,n){for(var i=this._controllerList,a=0;a<i.length;++a)i[a].pan(t,e,r,n)},s.translate=function(t,e,r,n){for(var i=this._controllerList,a=0;a<i.length;++a)i[a].translate(t,e,r,n)},s.setMatrix=function(t,e){for(var r=this._controllerList,n=0;n<r.length;++n)r[n].setMatrix(t,e)},s.setDistanceLimits=function(t,e){for(var r=this._controllerList,n=0;n<r.length;++n)r[n].setDistanceLimits(t,e)},s.setDistance=function(t,e){for(var r=this._controllerList,n=0;n<r.length;++n)r[n].setDistance(t,e)},s.recalcMatrix=function(t){this._active.recalcMatrix(t)},s.getDistance=function(t){return this._active.getDistance(t)},s.getDistanceLimits=function(t){return this._active.getDistanceLimits(t)},s.lastT=function(){return this._active.lastT()},s.setMode=function(t){if(t!==this._mode){var e=this._controllerNames.indexOf(t);if(!(e<0)){var r=this._active,n=this._controllerList[e],i=Math.max(r.lastT(),n.lastT());r.recalcMatrix(i),n.setMatrix(i,r.computedMatrix),this._active=n,this._mode=t,this.computedMatrix=this._active.computedMatrix,this.computedEye=this._active.computedEye,this.computedUp=this._active.computedUp,this.computedCenter=this._active.computedCenter,this.computedRadius=this._active.computedRadius}}},s.getMode=function(){return this._mode}},{"matrix-camera-controller":447,"orbit-camera-controller":468,"turntable-camera-controller":576}],56:[function(t,e,r){!function(n,i){"object"==typeof r&&void 0!==e?i(r,t("d3-array"),t("d3-collection"),t("d3-shape"),t("elementary-circuits-directed-graph")):i(n.d3=n.d3||{},n.d3,n.d3,n.d3,null)}(this,(function(t,e,r,n,i){"use strict";function a(t){return t.target.depth}function o(t,e){return t.sourceLinks.length?t.depth:e-1}function s(t){return function(){return t}}i=i&&i.hasOwnProperty("default")?i.default:i;var l="function"==typeof Symbol&&"symbol"==typeof Symbol.iterator?function(t){return typeof t}:function(t){return t&&"function"==typeof Symbol&&t.constructor===Symbol&&t!==Symbol.prototype?"symbol":typeof t};function c(t,e){return f(t.source,e.source)||t.index-e.index}function u(t,e){return f(t.target,e.target)||t.index-e.index}function f(t,e){return t.partOfCycle===e.partOfCycle?t.y0-e.y0:"top"===t.circularLinkType||"bottom"===e.circularLinkType?-1:1}function h(t){return t.value}function p(t){return(t.y0+t.y1)/2}function d(t){return p(t.source)}function m(t){return p(t.target)}function g(t){return t.index}function v(t){return t.nodes}function y(t){return t.links}function x(t,e){var r=t.get(e);if(!r)throw new Error("missing: "+e);return r}function b(t,e){return e(t)}function _(t,e,r){var n=0;if(null===r){for(var a=[],o=0;o<t.links.length;o++){var s=t.links[o],l=s.source.index,c=s.target.index;a[l]||(a[l]=[]),a[c]||(a[c]=[]),-1===a[l].indexOf(c)&&a[l].push(c)}var u=i(a);u.sort((function(t,e){return t.length-e.length}));var f={};for(o=0;o<u.length;o++){var h=u[o].slice(-2);f[h[0]]||(f[h[0]]={}),f[h[0]][h[1]]=!0}t.links.forEach((function(t){var e=t.target.index,r=t.source.index;e===r||f[r]&&f[r][e]?(t.circular=!0,t.circularLinkID=n,n+=1):t.circular=!1}))}else t.links.forEach((function(t){t.source[r]<t.target[r]?t.circular=!1:(t.circular=!0,t.circularLinkID=n,n+=1)}))}function w(t,e){var r=0,n=0;t.links.forEach((function(i){i.circular&&(i.source.circularLinkType||i.target.circularLinkType?i.circularLinkType=i.source.circularLinkType?i.source.circularLinkType:i.target.circularLinkType:i.circularLinkType=r<n?"top":"bottom","top"==i.circularLinkType?r+=1:n+=1,t.nodes.forEach((function(t){b(t,e)!=b(i.source,e)&&b(t,e)!=b(i.target,e)||(t.circularLinkType=i.circularLinkType)})))})),t.links.forEach((function(t){t.circular&&(t.source.circularLinkType==t.target.circularLinkType&&(t.circularLinkType=t.source.circularLinkType),q(t,e)&&(t.circularLinkType=t.source.circularLinkType))}))}function T(t){var e=Math.abs(t.y1-t.y0),r=Math.abs(t.target.x0-t.source.x1);return Math.atan(r/e)}function k(t,e){var r=0;t.sourceLinks.forEach((function(t){r=t.circular&&!q(t,e)?r+1:r}));var n=0;return t.targetLinks.forEach((function(t){n=t.circular&&!q(t,e)?n+1:n})),r+n}function A(t){var e=t.source.sourceLinks,r=0;e.forEach((function(t){r=t.circular?r+1:r}));var n=t.target.targetLinks,i=0;return n.forEach((function(t){i=t.circular?i+1:i})),!(r>1||i>1)}function M(t,e,r){return t.sort(E),t.forEach((function(n,i){var a,o,s=0;if(q(n,r)&&A(n))n.circularPathData.verticalBuffer=s+n.width/2;else{for(var l=0;l<i;l++)if(a=t[i],o=t[l],!(a.source.column<o.target.column||a.target.column>o.source.column)){var c=t[l].circularPathData.verticalBuffer+t[l].width/2+e;s=c>s?c:s}n.circularPathData.verticalBuffer=s+n.width/2}})),t}function S(t,r,i,a){var o=e.min(t.links,(function(t){return t.source.y0}));t.links.forEach((function(t){t.circular&&(t.circularPathData={})})),M(t.links.filter((function(t){return"top"==t.circularLinkType})),r,a),M(t.links.filter((function(t){return"bottom"==t.circularLinkType})),r,a),t.links.forEach((function(e){if(e.circular){if(e.circularPathData.arcRadius=e.width+10,e.circularPathData.leftNodeBuffer=5,e.circularPathData.rightNodeBuffer=5,e.circularPathData.sourceWidth=e.source.x1-e.source.x0,e.circularPathData.sourceX=e.source.x0+e.circularPathData.sourceWidth,e.circularPathData.targetX=e.target.x0,e.circularPathData.sourceY=e.y0,e.circularPathData.targetY=e.y1,q(e,a)&&A(e))e.circularPathData.leftSmallArcRadius=10+e.width/2,e.circularPathData.leftLargeArcRadius=10+e.width/2,e.circularPathData.rightSmallArcRadius=10+e.width/2,e.circularPathData.rightLargeArcRadius=10+e.width/2,"bottom"==e.circularLinkType?(e.circularPathData.verticalFullExtent=e.source.y1+25+e.circularPathData.verticalBuffer,e.circularPathData.verticalLeftInnerExtent=e.circularPathData.verticalFullExtent-e.circularPathData.leftLargeArcRadius,e.circularPathData.verticalRightInnerExtent=e.circularPathData.verticalFullExtent-e.circularPathData.rightLargeArcRadius):(e.circularPathData.verticalFullExtent=e.source.y0-25-e.circularPathData.verticalBuffer,e.circularPathData.verticalLeftInnerExtent=e.circularPathData.verticalFullExtent+e.circularPathData.leftLargeArcRadius,e.circularPathData.verticalRightInnerExtent=e.circularPathData.verticalFullExtent+e.circularPathData.rightLargeArcRadius);else{var s=e.source.column,l=e.circularLinkType,c=t.links.filter((function(t){return t.source.column==s&&t.circularLinkType==l}));"bottom"==e.circularLinkType?c.sort(C):c.sort(L);var u=0;c.forEach((function(t,n){t.circularLinkID==e.circularLinkID&&(e.circularPathData.leftSmallArcRadius=10+e.width/2+u,e.circularPathData.leftLargeArcRadius=10+e.width/2+n*r+u),u+=t.width})),s=e.target.column,c=t.links.filter((function(t){return t.target.column==s&&t.circularLinkType==l})),"bottom"==e.circularLinkType?c.sort(I):c.sort(P),u=0,c.forEach((function(t,n){t.circularLinkID==e.circularLinkID&&(e.circularPathData.rightSmallArcRadius=10+e.width/2+u,e.circularPathData.rightLargeArcRadius=10+e.width/2+n*r+u),u+=t.width})),"bottom"==e.circularLinkType?(e.circularPathData.verticalFullExtent=Math.max(i,e.source.y1,e.target.y1)+25+e.circularPathData.verticalBuffer,e.circularPathData.verticalLeftInnerExtent=e.circularPathData.verticalFullExtent-e.circularPathData.leftLargeArcRadius,e.circularPathData.verticalRightInnerExtent=e.circularPathData.verticalFullExtent-e.circularPathData.rightLargeArcRadius):(e.circularPathData.verticalFullExtent=o-25-e.circularPathData.verticalBuffer,e.circularPathData.verticalLeftInnerExtent=e.circularPathData.verticalFullExtent+e.circularPathData.leftLargeArcRadius,e.circularPathData.verticalRightInnerExtent=e.circularPathData.verticalFullExtent+e.circularPathData.rightLargeArcRadius)}e.circularPathData.leftInnerExtent=e.circularPathData.sourceX+e.circularPathData.leftNodeBuffer,e.circularPathData.rightInnerExtent=e.circularPathData.targetX-e.circularPathData.rightNodeBuffer,e.circularPathData.leftFullExtent=e.circularPathData.sourceX+e.circularPathData.leftLargeArcRadius+e.circularPathData.leftNodeBuffer,e.circularPathData.rightFullExtent=e.circularPathData.targetX-e.circularPathData.rightLargeArcRadius-e.circularPathData.rightNodeBuffer}if(e.circular)e.path=function(t){var e="";e="top"==t.circularLinkType?"M"+t.circularPathData.sourceX+" "+t.circularPathData.sourceY+" L"+t.circularPathData.leftInnerExtent+" "+t.circularPathData.sourceY+" A"+t.circularPathData.leftLargeArcRadius+" "+t.circularPathData.leftSmallArcRadius+" 0 0 0 "+t.circularPathData.leftFullExtent+" "+(t.circularPathData.sourceY-t.circularPathData.leftSmallArcRadius)+" L"+t.circularPathData.leftFullExtent+" "+t.circularPathData.verticalLeftInnerExtent+" A"+t.circularPathData.leftLargeArcRadius+" "+t.circularPathData.leftLargeArcRadius+" 0 0 0 "+t.circularPathData.leftInnerExtent+" "+t.circularPathData.verticalFullExtent+" L"+t.circularPathData.rightInnerExtent+" "+t.circularPathData.verticalFullExtent+" A"+t.circularPathData.rightLargeArcRadius+" "+t.circularPathData.rightLargeArcRadius+" 0 0 0 "+t.circularPathData.rightFullExtent+" "+t.circularPathData.verticalRightInnerExtent+" L"+t.circularPathData.rightFullExtent+" "+(t.circularPathData.targetY-t.circularPathData.rightSmallArcRadius)+" A"+t.circularPathData.rightLargeArcRadius+" "+t.circularPathData.rightSmallArcRadius+" 0 0 0 "+t.circularPathData.rightInnerExtent+" "+t.circularPathData.targetY+" L"+t.circularPathData.targetX+" "+t.circularPathData.targetY:"M"+t.circularPathData.sourceX+" "+t.circularPathData.sourceY+" L"+t.circularPathData.leftInnerExtent+" "+t.circularPathData.sourceY+" A"+t.circularPathData.leftLargeArcRadius+" "+t.circularPathData.leftSmallArcRadius+" 0 0 1 "+t.circularPathData.leftFullExtent+" "+(t.circularPathData.sourceY+t.circularPathData.leftSmallArcRadius)+" L"+t.circularPathData.leftFullExtent+" "+t.circularPathData.verticalLeftInnerExtent+" A"+t.circularPathData.leftLargeArcRadius+" "+t.circularPathData.leftLargeArcRadius+" 0 0 1 "+t.circularPathData.leftInnerExtent+" "+t.circularPathData.verticalFullExtent+" L"+t.circularPathData.rightInnerExtent+" "+t.circularPathData.verticalFullExtent+" A"+t.circularPathData.rightLargeArcRadius+" "+t.circularPathData.rightLargeArcRadius+" 0 0 1 "+t.circularPathData.rightFullExtent+" "+t.circularPathData.verticalRightInnerExtent+" L"+t.circularPathData.rightFullExtent+" "+(t.circularPathData.targetY+t.circularPathData.rightSmallArcRadius)+" A"+t.circularPathData.rightLargeArcRadius+" "+t.circularPathData.rightSmallArcRadius+" 0 0 1 "+t.circularPathData.rightInnerExtent+" "+t.circularPathData.targetY+" L"+t.circularPathData.targetX+" "+t.circularPathData.targetY;return e}(e);else{var f=n.linkHorizontal().source((function(t){return[t.source.x0+(t.source.x1-t.source.x0),t.y0]})).target((function(t){return[t.target.x0,t.y1]}));e.path=f(e)}}))}function E(t,e){return O(t)==O(e)?"bottom"==t.circularLinkType?C(t,e):L(t,e):O(e)-O(t)}function L(t,e){return t.y0-e.y0}function C(t,e){return e.y0-t.y0}function P(t,e){return t.y1-e.y1}function I(t,e){return e.y1-t.y1}function O(t){return t.target.column-t.source.column}function z(t){return t.target.x0-t.source.x1}function D(t,e){var r=T(t),n=z(e)/Math.tan(r);return"up"==H(t)?t.y1+n:t.y1-n}function R(t,e){var r=T(t),n=z(e)/Math.tan(r);return"up"==H(t)?t.y1-n:t.y1+n}function F(t,e,r,n){t.links.forEach((function(i){if(!i.circular&&i.target.column-i.source.column>1){var a=i.source.column+1,o=i.target.column-1,s=1,l=o-a+1;for(s=1;a<=o;a++,s++)t.nodes.forEach((function(o){if(o.column==a){var c,u=s/(l+1),f=Math.pow(1-u,3),h=3*u*Math.pow(1-u,2),p=3*Math.pow(u,2)*(1-u),d=Math.pow(u,3),m=f*i.y0+h*i.y0+p*i.y1+d*i.y1,g=m-i.width/2,v=m+i.width/2;g>o.y0&&g<o.y1?(c=o.y1-g+10,c="bottom"==o.circularLinkType?c:-c,o=N(o,c,e,r),t.nodes.forEach((function(t){b(t,n)!=b(o,n)&&t.column==o.column&&B(o,t)&&N(t,c,e,r)}))):(v>o.y0&&v<o.y1||g<o.y0&&v>o.y1)&&(c=v-o.y0+10,o=N(o,c,e,r),t.nodes.forEach((function(t){b(t,n)!=b(o,n)&&t.column==o.column&&t.y0<o.y1&&t.y1>o.y1&&N(t,c,e,r)})))}}))}}))}function B(t,e){return t.y0>e.y0&&t.y0<e.y1||(t.y1>e.y0&&t.y1<e.y1||t.y0<e.y0&&t.y1>e.y1)}function N(t,e,r,n){return t.y0+e>=r&&t.y1+e<=n&&(t.y0=t.y0+e,t.y1=t.y1+e,t.targetLinks.forEach((function(t){t.y1=t.y1+e})),t.sourceLinks.forEach((function(t){t.y0=t.y0+e}))),t}function j(t,e,r,n){t.nodes.forEach((function(i){n&&i.y+(i.y1-i.y0)>e&&(i.y=i.y-(i.y+(i.y1-i.y0)-e));var a=t.links.filter((function(t){return b(t.source,r)==b(i,r)})),o=a.length;o>1&&a.sort((function(t,e){if(!t.circular&&!e.circular){if(t.target.column==e.target.column)return t.y1-e.y1;if(!V(t,e))return t.y1-e.y1;if(t.target.column>e.target.column){var r=R(e,t);return t.y1-r}if(e.target.column>t.target.column)return R(t,e)-e.y1}return t.circular&&!e.circular?"top"==t.circularLinkType?-1:1:e.circular&&!t.circular?"top"==e.circularLinkType?1:-1:t.circular&&e.circular?t.circularLinkType===e.circularLinkType&&"top"==t.circularLinkType?t.target.column===e.target.column?t.target.y1-e.target.y1:e.target.column-t.target.column:t.circularLinkType===e.circularLinkType&&"bottom"==t.circularLinkType?t.target.column===e.target.column?e.target.y1-t.target.y1:t.target.column-e.target.column:"top"==t.circularLinkType?-1:1:void 0}));var s=i.y0;a.forEach((function(t){t.y0=s+t.width/2,s+=t.width})),a.forEach((function(t,e){if("bottom"==t.circularLinkType){for(var r=e+1,n=0;r<o;r++)n+=a[r].width;t.y0=i.y1-n-t.width/2}}))}))}function U(t,e,r){t.nodes.forEach((function(e){var n=t.links.filter((function(t){return b(t.target,r)==b(e,r)})),i=n.length;i>1&&n.sort((function(t,e){if(!t.circular&&!e.circular){if(t.source.column==e.source.column)return t.y0-e.y0;if(!V(t,e))return t.y0-e.y0;if(e.source.column<t.source.column){var r=D(e,t);return t.y0-r}if(t.source.column<e.source.column)return D(t,e)-e.y0}return t.circular&&!e.circular?"top"==t.circularLinkType?-1:1:e.circular&&!t.circular?"top"==e.circularLinkType?1:-1:t.circular&&e.circular?t.circularLinkType===e.circularLinkType&&"top"==t.circularLinkType?t.source.column===e.source.column?t.source.y1-e.source.y1:t.source.column-e.source.column:t.circularLinkType===e.circularLinkType&&"bottom"==t.circularLinkType?t.source.column===e.source.column?t.source.y1-e.source.y1:e.source.column-t.source.column:"top"==t.circularLinkType?-1:1:void 0}));var a=e.y0;n.forEach((function(t){t.y1=a+t.width/2,a+=t.width})),n.forEach((function(t,r){if("bottom"==t.circularLinkType){for(var a=r+1,o=0;a<i;a++)o+=n[a].width;t.y1=e.y1-o-t.width/2}}))}))}function V(t,e){return H(t)==H(e)}function H(t){return t.y0-t.y1>0?"up":"down"}function q(t,e){return b(t.source,e)==b(t.target,e)}function G(t,r,n){var i=t.nodes,a=t.links,o=!1,s=!1;if(a.forEach((function(t){"top"==t.circularLinkType?o=!0:"bottom"==t.circularLinkType&&(s=!0)})),0==o||0==s){var l=e.min(i,(function(t){return t.y0})),c=(n-r)/(e.max(i,(function(t){return t.y1}))-l);i.forEach((function(t){var e=(t.y1-t.y0)*c;t.y0=(t.y0-l)*c,t.y1=t.y0+e})),a.forEach((function(t){t.y0=(t.y0-l)*c,t.y1=(t.y1-l)*c,t.width=t.width*c}))}}t.sankeyCircular=function(){var t,n,i=0,a=0,b=1,T=1,A=24,M=g,E=o,L=v,C=y,P=32,I=2,O=null;function z(){var t={nodes:L.apply(null,arguments),links:C.apply(null,arguments)};D(t),_(t,M,O),R(t),B(t),w(t,M),N(t,P,M),V(t);for(var e=4,r=0;r<e;r++)j(t,T,M),U(t,T,M),F(t,a,T,M),j(t,T,M),U(t,T,M);return G(t,a,T),S(t,I,T,M),t}function D(t){t.nodes.forEach((function(t,e){t.index=e,t.sourceLinks=[],t.targetLinks=[]}));var e=r.map(t.nodes,M);return t.links.forEach((function(t,r){t.index=r;var n=t.source,i=t.target;"object"!==(void 0===n?"undefined":l(n))&&(n=t.source=x(e,n)),"object"!==(void 0===i?"undefined":l(i))&&(i=t.target=x(e,i)),n.sourceLinks.push(t),i.targetLinks.push(t)})),t}function R(t){t.nodes.forEach((function(t){t.partOfCycle=!1,t.value=Math.max(e.sum(t.sourceLinks,h),e.sum(t.targetLinks,h)),t.sourceLinks.forEach((function(e){e.circular&&(t.partOfCycle=!0,t.circularLinkType=e.circularLinkType)})),t.targetLinks.forEach((function(e){e.circular&&(t.partOfCycle=!0,t.circularLinkType=e.circularLinkType)}))}))}function B(t){var e,r,n;for(e=t.nodes,r=[],n=0;e.length;++n,e=r,r=[])e.forEach((function(t){t.depth=n,t.sourceLinks.forEach((function(t){r.indexOf(t.target)<0&&!t.circular&&r.push(t.target)}))}));for(e=t.nodes,r=[],n=0;e.length;++n,e=r,r=[])e.forEach((function(t){t.height=n,t.targetLinks.forEach((function(t){r.indexOf(t.source)<0&&!t.circular&&r.push(t.source)}))}));t.nodes.forEach((function(t){t.column=Math.floor(E.call(null,t,n))}))}function N(o,s,l){var c=r.nest().key((function(t){return t.column})).sortKeys(e.ascending).entries(o.nodes).map((function(t){return t.values}));!function(r){if(n){var s=1/0;c.forEach((function(t){var e=T*n/(t.length+1);s=e<s?e:s})),t=s}var l=e.min(c,(function(r){return(T-a-(r.length-1)*t)/e.sum(r,h)}));l*=.3,o.links.forEach((function(t){t.width=t.value*l}));var u=function(t){var r=0,n=0,i=0,a=0,o=e.max(t.nodes,(function(t){return t.column}));return t.links.forEach((function(t){t.circular&&("top"==t.circularLinkType?r+=t.width:n+=t.width,0==t.target.column&&(a+=t.width),t.source.column==o&&(i+=t.width))})),{top:r=r>0?r+25+10:r,bottom:n=n>0?n+25+10:n,left:a=a>0?a+25+10:a,right:i=i>0?i+25+10:i}}(o),f=function(t,r){var n=e.max(t.nodes,(function(t){return t.column})),o=b-i,s=T-a,l=o/(o+r.right+r.left),c=s/(s+r.top+r.bottom);return i=i*l+r.left,b=0==r.right?b:b*l,a=a*c+r.top,T*=c,t.nodes.forEach((function(t){t.x0=i+t.column*((b-i-A)/n),t.x1=t.x0+A})),c}(o,u);l*=f,o.links.forEach((function(t){t.width=t.value*l})),c.forEach((function(t){var e=t.length;t.forEach((function(t,n){t.depth==c.length-1&&1==e||0==t.depth&&1==e?(t.y0=T/2-t.value*l,t.y1=t.y0+t.value*l):t.partOfCycle?0==k(t,r)?(t.y0=T/2+n,t.y1=t.y0+t.value*l):"top"==t.circularLinkType?(t.y0=a+n,t.y1=t.y0+t.value*l):(t.y0=T-t.value*l-n,t.y1=t.y0+t.value*l):0==u.top||0==u.bottom?(t.y0=(T-a)/e*n,t.y1=t.y0+t.value*l):(t.y0=(T-a)/2-e/2+n,t.y1=t.y0+t.value*l)}))}))}(l),y();for(var u=1,g=s;g>0;--g)v(u*=.99,l),y();function v(t,r){var n=c.length;c.forEach((function(i){var a=i.length,o=i[0].depth;i.forEach((function(i){var s;if(i.sourceLinks.length||i.targetLinks.length)if(i.partOfCycle&&k(i,r)>0);else if(0==o&&1==a)s=i.y1-i.y0,i.y0=T/2-s/2,i.y1=T/2+s/2;else if(o==n-1&&1==a)s=i.y1-i.y0,i.y0=T/2-s/2,i.y1=T/2+s/2;else{var l=e.mean(i.sourceLinks,m),c=e.mean(i.targetLinks,d),u=((l&&c?(l+c)/2:l||c)-p(i))*t;i.y0+=u,i.y1+=u}}))}))}function y(){c.forEach((function(e){var r,n,i,o=a,s=e.length;for(e.sort(f),i=0;i<s;++i)(n=o-(r=e[i]).y0)>0&&(r.y0+=n,r.y1+=n),o=r.y1+t;if((n=o-t-T)>0)for(o=r.y0-=n,r.y1-=n,i=s-2;i>=0;--i)(n=(r=e[i]).y1+t-o)>0&&(r.y0-=n,r.y1-=n),o=r.y0}))}}function V(t){t.nodes.forEach((function(t){t.sourceLinks.sort(u),t.targetLinks.sort(c)})),t.nodes.forEach((function(t){var e=t.y0,r=e,n=t.y1,i=n;t.sourceLinks.forEach((function(t){t.circular?(t.y0=n-t.width/2,n-=t.width):(t.y0=e+t.width/2,e+=t.width)})),t.targetLinks.forEach((function(t){t.circular?(t.y1=i-t.width/2,i-=t.width):(t.y1=r+t.width/2,r+=t.width)}))}))}return z.nodeId=function(t){return arguments.length?(M="function"==typeof t?t:s(t),z):M},z.nodeAlign=function(t){return arguments.length?(E="function"==typeof t?t:s(t),z):E},z.nodeWidth=function(t){return arguments.length?(A=+t,z):A},z.nodePadding=function(e){return arguments.length?(t=+e,z):t},z.nodes=function(t){return arguments.length?(L="function"==typeof t?t:s(t),z):L},z.links=function(t){return arguments.length?(C="function"==typeof t?t:s(t),z):C},z.size=function(t){return arguments.length?(i=a=0,b=+t[0],T=+t[1],z):[b-i,T-a]},z.extent=function(t){return arguments.length?(i=+t[0][0],b=+t[1][0],a=+t[0][1],T=+t[1][1],z):[[i,a],[b,T]]},z.iterations=function(t){return arguments.length?(P=+t,z):P},z.circularLinkGap=function(t){return arguments.length?(I=+t,z):I},z.nodePaddingRatio=function(t){return arguments.length?(n=+t,z):n},z.sortNodes=function(t){return arguments.length?(O=t,z):O},z.update=function(t){return w(t,M),V(t),t.links.forEach((function(t){t.circular&&(t.circularLinkType=t.y0+t.y1<T?"top":"bottom",t.source.circularLinkType=t.circularLinkType,t.target.circularLinkType=t.circularLinkType)})),j(t,T,M,!1),U(t,T,M),S(t,I,T,M),t},z},t.sankeyCenter=function(t){return t.targetLinks.length?t.depth:t.sourceLinks.length?e.min(t.sourceLinks,a)-1:0},t.sankeyLeft=function(t){return t.depth},t.sankeyRight=function(t,e){return e-1-t.height},t.sankeyJustify=o,Object.defineProperty(t,"__esModule",{value:!0})}))},{"d3-array":155,"d3-collection":156,"d3-shape":167,"elementary-circuits-directed-graph":180}],57:[function(t,e,r){!function(n,i){"object"==typeof r&&void 0!==e?i(r,t("d3-array"),t("d3-collection"),t("d3-shape")):i(n.d3=n.d3||{},n.d3,n.d3,n.d3)}(this,(function(t,e,r,n){"use strict";function i(t){return t.target.depth}function a(t,e){return t.sourceLinks.length?t.depth:e-1}function o(t){return function(){return t}}function s(t,e){return c(t.source,e.source)||t.index-e.index}function l(t,e){return c(t.target,e.target)||t.index-e.index}function c(t,e){return t.y0-e.y0}function u(t){return t.value}function f(t){return(t.y0+t.y1)/2}function h(t){return f(t.source)*t.value}function p(t){return f(t.target)*t.value}function d(t){return t.index}function m(t){return t.nodes}function g(t){return t.links}function v(t,e){var r=t.get(e);if(!r)throw new Error("missing: "+e);return r}function y(t){return[t.source.x1,t.y0]}function x(t){return[t.target.x0,t.y1]}t.sankey=function(){var t=0,n=0,i=1,y=1,x=24,b=8,_=d,w=a,T=m,k=g,A=32;function M(){var t={nodes:T.apply(null,arguments),links:k.apply(null,arguments)};return S(t),E(t),L(t),C(t),P(t),t}function S(t){t.nodes.forEach((function(t,e){t.index=e,t.sourceLinks=[],t.targetLinks=[]}));var e=r.map(t.nodes,_);t.links.forEach((function(t,r){t.index=r;var n=t.source,i=t.target;"object"!=typeof n&&(n=t.source=v(e,n)),"object"!=typeof i&&(i=t.target=v(e,i)),n.sourceLinks.push(t),i.targetLinks.push(t)}))}function E(t){t.nodes.forEach((function(t){t.value=Math.max(e.sum(t.sourceLinks,u),e.sum(t.targetLinks,u))}))}function L(e){var r,n,a;for(r=e.nodes,n=[],a=0;r.length;++a,r=n,n=[])r.forEach((function(t){t.depth=a,t.sourceLinks.forEach((function(t){n.indexOf(t.target)<0&&n.push(t.target)}))}));for(r=e.nodes,n=[],a=0;r.length;++a,r=n,n=[])r.forEach((function(t){t.height=a,t.targetLinks.forEach((function(t){n.indexOf(t.source)<0&&n.push(t.source)}))}));var o=(i-t-x)/(a-1);e.nodes.forEach((function(e){e.x1=(e.x0=t+Math.max(0,Math.min(a-1,Math.floor(w.call(null,e,a))))*o)+x}))}function C(t){var i=r.nest().key((function(t){return t.x0})).sortKeys(e.ascending).entries(t.nodes).map((function(t){return t.values}));!function(){var r=e.max(i,(function(t){return t.length})),a=2/3*(y-n)/(r-1);b>a&&(b=a);var o=e.min(i,(function(t){return(y-n-(t.length-1)*b)/e.sum(t,u)}));i.forEach((function(t){t.forEach((function(t,e){t.y1=(t.y0=e)+t.value*o}))})),t.links.forEach((function(t){t.width=t.value*o}))}(),d();for(var a=1,o=A;o>0;--o)l(a*=.99),d(),s(a),d();function s(t){i.forEach((function(r){r.forEach((function(r){if(r.targetLinks.length){var n=(e.sum(r.targetLinks,h)/e.sum(r.targetLinks,u)-f(r))*t;r.y0+=n,r.y1+=n}}))}))}function l(t){i.slice().reverse().forEach((function(r){r.forEach((function(r){if(r.sourceLinks.length){var n=(e.sum(r.sourceLinks,p)/e.sum(r.sourceLinks,u)-f(r))*t;r.y0+=n,r.y1+=n}}))}))}function d(){i.forEach((function(t){var e,r,i,a=n,o=t.length;for(t.sort(c),i=0;i<o;++i)(r=a-(e=t[i]).y0)>0&&(e.y0+=r,e.y1+=r),a=e.y1+b;if((r=a-b-y)>0)for(a=e.y0-=r,e.y1-=r,i=o-2;i>=0;--i)(r=(e=t[i]).y1+b-a)>0&&(e.y0-=r,e.y1-=r),a=e.y0}))}}function P(t){t.nodes.forEach((function(t){t.sourceLinks.sort(l),t.targetLinks.sort(s)})),t.nodes.forEach((function(t){var e=t.y0,r=e;t.sourceLinks.forEach((function(t){t.y0=e+t.width/2,e+=t.width})),t.targetLinks.forEach((function(t){t.y1=r+t.width/2,r+=t.width}))}))}return M.update=function(t){return P(t),t},M.nodeId=function(t){return arguments.length?(_="function"==typeof t?t:o(t),M):_},M.nodeAlign=function(t){return arguments.length?(w="function"==typeof t?t:o(t),M):w},M.nodeWidth=function(t){return arguments.length?(x=+t,M):x},M.nodePadding=function(t){return arguments.length?(b=+t,M):b},M.nodes=function(t){return arguments.length?(T="function"==typeof t?t:o(t),M):T},M.links=function(t){return arguments.length?(k="function"==typeof t?t:o(t),M):k},M.size=function(e){return arguments.length?(t=n=0,i=+e[0],y=+e[1],M):[i-t,y-n]},M.extent=function(e){return arguments.length?(t=+e[0][0],i=+e[1][0],n=+e[0][1],y=+e[1][1],M):[[t,n],[i,y]]},M.iterations=function(t){return arguments.length?(A=+t,M):A},M},t.sankeyCenter=function(t){return t.targetLinks.length?t.depth:t.sourceLinks.length?e.min(t.sourceLinks,i)-1:0},t.sankeyLeft=function(t){return t.depth},t.sankeyRight=function(t,e){return e-1-t.height},t.sankeyJustify=a,t.sankeyLinkHorizontal=function(){return n.linkHorizontal().source(y).target(x)},Object.defineProperty(t,"__esModule",{value:!0})}))},{"d3-array":155,"d3-collection":156,"d3-shape":167}],58:[function(t,e,r){(function(){var t={version:"3.8.0"},r=[].slice,n=function(t){return r.call(t)},i=self.document;function a(t){return t&&(t.ownerDocument||t.document||t).documentElement}function o(t){return t&&(t.ownerDocument&&t.ownerDocument.defaultView||t.document&&t||t.defaultView)}if(i)try{n(i.documentElement.childNodes)[0].nodeType}catch(t){n=function(t){for(var e=t.length,r=new Array(e);e--;)r[e]=t[e];return r}}if(Date.now||(Date.now=function(){return+new Date}),i)try{i.createElement("DIV").style.setProperty("opacity",0,"")}catch(t){var s=this.Element.prototype,l=s.setAttribute,c=s.setAttributeNS,u=this.CSSStyleDeclaration.prototype,f=u.setProperty;s.setAttribute=function(t,e){l.call(this,t,e+"")},s.setAttributeNS=function(t,e,r){c.call(this,t,e,r+"")},u.setProperty=function(t,e,r){f.call(this,t,e+"",r)}}function h(t,e){return t<e?-1:t>e?1:t>=e?0:NaN}function p(t){return null===t?NaN:+t}function d(t){return!isNaN(t)}function m(t){return{left:function(e,r,n,i){for(arguments.length<3&&(n=0),arguments.length<4&&(i=e.length);n<i;){var a=n+i>>>1;t(e[a],r)<0?n=a+1:i=a}return n},right:function(e,r,n,i){for(arguments.length<3&&(n=0),arguments.length<4&&(i=e.length);n<i;){var a=n+i>>>1;t(e[a],r)>0?i=a:n=a+1}return n}}}t.ascending=h,t.descending=function(t,e){return e<t?-1:e>t?1:e>=t?0:NaN},t.min=function(t,e){var r,n,i=-1,a=t.length;if(1===arguments.length){for(;++i<a;)if(null!=(n=t[i])&&n>=n){r=n;break}for(;++i<a;)null!=(n=t[i])&&r>n&&(r=n)}else{for(;++i<a;)if(null!=(n=e.call(t,t[i],i))&&n>=n){r=n;break}for(;++i<a;)null!=(n=e.call(t,t[i],i))&&r>n&&(r=n)}return r},t.max=function(t,e){var r,n,i=-1,a=t.length;if(1===arguments.length){for(;++i<a;)if(null!=(n=t[i])&&n>=n){r=n;break}for(;++i<a;)null!=(n=t[i])&&n>r&&(r=n)}else{for(;++i<a;)if(null!=(n=e.call(t,t[i],i))&&n>=n){r=n;break}for(;++i<a;)null!=(n=e.call(t,t[i],i))&&n>r&&(r=n)}return r},t.extent=function(t,e){var r,n,i,a=-1,o=t.length;if(1===arguments.length){for(;++a<o;)if(null!=(n=t[a])&&n>=n){r=i=n;break}for(;++a<o;)null!=(n=t[a])&&(r>n&&(r=n),i<n&&(i=n))}else{for(;++a<o;)if(null!=(n=e.call(t,t[a],a))&&n>=n){r=i=n;break}for(;++a<o;)null!=(n=e.call(t,t[a],a))&&(r>n&&(r=n),i<n&&(i=n))}return[r,i]},t.sum=function(t,e){var r,n=0,i=t.length,a=-1;if(1===arguments.length)for(;++a<i;)d(r=+t[a])&&(n+=r);else for(;++a<i;)d(r=+e.call(t,t[a],a))&&(n+=r);return n},t.mean=function(t,e){var r,n=0,i=t.length,a=-1,o=i;if(1===arguments.length)for(;++a<i;)d(r=p(t[a]))?n+=r:--o;else for(;++a<i;)d(r=p(e.call(t,t[a],a)))?n+=r:--o;if(o)return n/o},t.quantile=function(t,e){var r=(t.length-1)*e+1,n=Math.floor(r),i=+t[n-1],a=r-n;return a?i+a*(t[n]-i):i},t.median=function(e,r){var n,i=[],a=e.length,o=-1;if(1===arguments.length)for(;++o<a;)d(n=p(e[o]))&&i.push(n);else for(;++o<a;)d(n=p(r.call(e,e[o],o)))&&i.push(n);if(i.length)return t.quantile(i.sort(h),.5)},t.variance=function(t,e){var r,n,i=t.length,a=0,o=0,s=-1,l=0;if(1===arguments.length)for(;++s<i;)d(r=p(t[s]))&&(o+=(n=r-a)*(r-(a+=n/++l)));else for(;++s<i;)d(r=p(e.call(t,t[s],s)))&&(o+=(n=r-a)*(r-(a+=n/++l)));if(l>1)return o/(l-1)},t.deviation=function(){var e=t.variance.apply(this,arguments);return e?Math.sqrt(e):e};var g=m(h);function v(t){return t.length}t.bisectLeft=g.left,t.bisect=t.bisectRight=g.right,t.bisector=function(t){return m(1===t.length?function(e,r){return h(t(e),r)}:t)},t.shuffle=function(t,e,r){(a=arguments.length)<3&&(r=t.length,a<2&&(e=0));for(var n,i,a=r-e;a;)i=Math.random()*a--|0,n=t[a+e],t[a+e]=t[i+e],t[i+e]=n;return t},t.permute=function(t,e){for(var r=e.length,n=new Array(r);r--;)n[r]=t[e[r]];return n},t.pairs=function(t){for(var e=0,r=t.length-1,n=t[0],i=new Array(r<0?0:r);e<r;)i[e]=[n,n=t[++e]];return i},t.transpose=function(e){if(!(a=e.length))return[];for(var r=-1,n=t.min(e,v),i=new Array(n);++r<n;)for(var a,o=-1,s=i[r]=new Array(a);++o<a;)s[o]=e[o][r];return i},t.zip=function(){return t.transpose(arguments)},t.keys=function(t){var e=[];for(var r in t)e.push(r);return e},t.values=function(t){var e=[];for(var r in t)e.push(t[r]);return e},t.entries=function(t){var e=[];for(var r in t)e.push({key:r,value:t[r]});return e},t.merge=function(t){for(var e,r,n,i=t.length,a=-1,o=0;++a<i;)o+=t[a].length;for(r=new Array(o);--i>=0;)for(e=(n=t[i]).length;--e>=0;)r[--o]=n[e];return r};var y=Math.abs;function x(t){for(var e=1;t*e%1;)e*=10;return e}function b(t,e){for(var r in e)Object.defineProperty(t.prototype,r,{value:e[r],enumerable:!1})}function _(){this._=Object.create(null)}t.range=function(t,e,r){if(arguments.length<3&&(r=1,arguments.length<2&&(e=t,t=0)),(e-t)/r==1/0)throw new Error("infinite range");var n,i=[],a=x(y(r)),o=-1;if(t*=a,e*=a,(r*=a)<0)for(;(n=t+r*++o)>e;)i.push(n/a);else for(;(n=t+r*++o)<e;)i.push(n/a);return i},t.map=function(t,e){var r=new _;if(t instanceof _)t.forEach((function(t,e){r.set(t,e)}));else if(Array.isArray(t)){var n,i=-1,a=t.length;if(1===arguments.length)for(;++i<a;)r.set(i,t[i]);else for(;++i<a;)r.set(e.call(t,n=t[i],i),n)}else for(var o in t)r.set(o,t[o]);return r};function w(t){return"__proto__"==(t+="")||"\0"===t[0]?"\0"+t:t}function T(t){return"\0"===(t+="")[0]?t.slice(1):t}function k(t){return w(t)in this._}function A(t){return(t=w(t))in this._&&delete this._[t]}function M(){var t=[];for(var e in this._)t.push(T(e));return t}function S(){var t=0;for(var e in this._)++t;return t}function E(){for(var t in this._)return!1;return!0}function L(){this._=Object.create(null)}function C(t){return t}function P(t,e,r){return function(){var n=r.apply(e,arguments);return n===e?t:n}}function I(t,e){if(e in t)return e;e=e.charAt(0).toUpperCase()+e.slice(1);for(var r=0,n=O.length;r<n;++r){var i=O[r]+e;if(i in t)return i}}b(_,{has:k,get:function(t){return this._[w(t)]},set:function(t,e){return this._[w(t)]=e},remove:A,keys:M,values:function(){var t=[];for(var e in this._)t.push(this._[e]);return t},entries:function(){var t=[];for(var e in this._)t.push({key:T(e),value:this._[e]});return t},size:S,empty:E,forEach:function(t){for(var e in this._)t.call(this,T(e),this._[e])}}),t.nest=function(){var e,r,n={},i=[],a=[];function o(t,a,s){if(s>=i.length)return r?r.call(n,a):e?a.sort(e):a;for(var l,c,u,f,h=-1,p=a.length,d=i[s++],m=new _;++h<p;)(f=m.get(l=d(c=a[h])))?f.push(c):m.set(l,[c]);return t?(c=t(),u=function(e,r){c.set(e,o(t,r,s))}):(c={},u=function(e,r){c[e]=o(t,r,s)}),m.forEach(u),c}return n.map=function(t,e){return o(e,t,0)},n.entries=function(e){return function t(e,r){if(r>=i.length)return e;var n=[],o=a[r++];return e.forEach((function(e,i){n.push({key:e,values:t(i,r)})})),o?n.sort((function(t,e){return o(t.key,e.key)})):n}(o(t.map,e,0),0)},n.key=function(t){return i.push(t),n},n.sortKeys=function(t){return a[i.length-1]=t,n},n.sortValues=function(t){return e=t,n},n.rollup=function(t){return r=t,n},n},t.set=function(t){var e=new L;if(t)for(var r=0,n=t.length;r<n;++r)e.add(t[r]);return e},b(L,{has:k,add:function(t){return this._[w(t+="")]=!0,t},remove:A,values:M,size:S,empty:E,forEach:function(t){for(var e in this._)t.call(this,T(e))}}),t.behavior={},t.rebind=function(t,e){for(var r,n=1,i=arguments.length;++n<i;)t[r=arguments[n]]=P(t,e,e[r]);return t};var O=["webkit","ms","moz","Moz","o","O"];function z(){}function D(){}function R(t){var e=[],r=new _;function n(){for(var r,n=e,i=-1,a=n.length;++i<a;)(r=n[i].on)&&r.apply(this,arguments);return t}return n.on=function(n,i){var a,o=r.get(n);return arguments.length<2?o&&o.on:(o&&(o.on=null,e=e.slice(0,a=e.indexOf(o)).concat(e.slice(a+1)),r.remove(n)),i&&e.push(r.set(n,{on:i})),t)},n}function F(){t.event.preventDefault()}function B(){for(var e,r=t.event;e=r.sourceEvent;)r=e;return r}function N(e){for(var r=new D,n=0,i=arguments.length;++n<i;)r[arguments[n]]=R(r);return r.of=function(n,i){return function(a){try{var o=a.sourceEvent=t.event;a.target=e,t.event=a,r[a.type].apply(n,i)}finally{t.event=o}}},r}t.dispatch=function(){for(var t=new D,e=-1,r=arguments.length;++e<r;)t[arguments[e]]=R(t);return t},D.prototype.on=function(t,e){var r=t.indexOf("."),n="";if(r>=0&&(n=t.slice(r+1),t=t.slice(0,r)),t)return arguments.length<2?this[t].on(n):this[t].on(n,e);if(2===arguments.length){if(null==e)for(t in this)this.hasOwnProperty(t)&&this[t].on(n,null);return this}},t.event=null,t.requote=function(t){return t.replace(j,"\\$&")};var j=/[\\\^\$\*\+\?\|\[\]\(\)\.\{\}]/g,U={}.__proto__?function(t,e){t.__proto__=e}:function(t,e){for(var r in e)t[r]=e[r]};function V(t){return U(t,Y),t}var H=function(t,e){return e.querySelector(t)},q=function(t,e){return e.querySelectorAll(t)},G=function(t,e){var r=t.matches||t[I(t,"matchesSelector")];return(G=function(t,e){return r.call(t,e)})(t,e)};"function"==typeof Sizzle&&(H=function(t,e){return Sizzle(t,e)[0]||null},q=Sizzle,G=Sizzle.matchesSelector),t.selection=function(){return t.select(i.documentElement)};var Y=t.selection.prototype=[];function W(t){return"function"==typeof t?t:function(){return H(t,this)}}function X(t){return"function"==typeof t?t:function(){return q(t,this)}}Y.select=function(t){var e,r,n,i,a=[];t=W(t);for(var o=-1,s=this.length;++o<s;){a.push(e=[]),e.parentNode=(n=this[o]).parentNode;for(var l=-1,c=n.length;++l<c;)(i=n[l])?(e.push(r=t.call(i,i.__data__,l,o)),r&&"__data__"in i&&(r.__data__=i.__data__)):e.push(null)}return V(a)},Y.selectAll=function(t){var e,r,i=[];t=X(t);for(var a=-1,o=this.length;++a<o;)for(var s=this[a],l=-1,c=s.length;++l<c;)(r=s[l])&&(i.push(e=n(t.call(r,r.__data__,l,a))),e.parentNode=r);return V(i)};var Z="http://www.w3.org/1999/xhtml",J={svg:"http://www.w3.org/2000/svg",xhtml:Z,xlink:"http://www.w3.org/1999/xlink",xml:"http://www.w3.org/XML/1998/namespace",xmlns:"http://www.w3.org/2000/xmlns/"};function K(e,r){return e=t.ns.qualify(e),null==r?e.local?function(){this.removeAttributeNS(e.space,e.local)}:function(){this.removeAttribute(e)}:"function"==typeof r?e.local?function(){var t=r.apply(this,arguments);null==t?this.removeAttributeNS(e.space,e.local):this.setAttributeNS(e.space,e.local,t)}:function(){var t=r.apply(this,arguments);null==t?this.removeAttribute(e):this.setAttribute(e,t)}:e.local?function(){this.setAttributeNS(e.space,e.local,r)}:function(){this.setAttribute(e,r)}}function Q(t){return t.trim().replace(/\s+/g," ")}function $(e){return new RegExp("(?:^|\\s+)"+t.requote(e)+"(?:\\s+|$)","g")}function tt(t){return(t+"").trim().split(/^|\s+/)}function et(t,e){var r=(t=tt(t).map(rt)).length;return"function"==typeof e?function(){for(var n=-1,i=e.apply(this,arguments);++n<r;)t[n](this,i)}:function(){for(var n=-1;++n<r;)t[n](this,e)}}function rt(t){var e=$(t);return function(r,n){if(i=r.classList)return n?i.add(t):i.remove(t);var i=r.getAttribute("class")||"";n?(e.lastIndex=0,e.test(i)||r.setAttribute("class",Q(i+" "+t))):r.setAttribute("class",Q(i.replace(e," ")))}}function nt(t,e,r){return null==e?function(){this.style.removeProperty(t)}:"function"==typeof e?function(){var n=e.apply(this,arguments);null==n?this.style.removeProperty(t):this.style.setProperty(t,n,r)}:function(){this.style.setProperty(t,e,r)}}function it(t,e){return null==e?function(){delete this[t]}:"function"==typeof e?function(){var r=e.apply(this,arguments);null==r?delete this[t]:this[t]=r}:function(){this[t]=e}}function at(e){return"function"==typeof e?e:(e=t.ns.qualify(e)).local?function(){return this.ownerDocument.createElementNS(e.space,e.local)}:function(){var t=this.ownerDocument,r=this.namespaceURI;return r===Z&&t.documentElement.namespaceURI===Z?t.createElement(e):t.createElementNS(r,e)}}function ot(){var t=this.parentNode;t&&t.removeChild(this)}function st(t){return{__data__:t}}function lt(t){return function(){return G(this,t)}}function ct(t){return arguments.length||(t=h),function(e,r){return e&&r?t(e.__data__,r.__data__):!e-!r}}function ut(t,e){for(var r=0,n=t.length;r<n;r++)for(var i,a=t[r],o=0,s=a.length;o<s;o++)(i=a[o])&&e(i,o,r);return t}function ft(t){return U(t,ht),t}t.ns={prefix:J,qualify:function(t){var e=t.indexOf(":"),r=t;return e>=0&&"xmlns"!==(r=t.slice(0,e))&&(t=t.slice(e+1)),J.hasOwnProperty(r)?{space:J[r],local:t}:t}},Y.attr=function(e,r){if(arguments.length<2){if("string"==typeof e){var n=this.node();return(e=t.ns.qualify(e)).local?n.getAttributeNS(e.space,e.local):n.getAttribute(e)}for(r in e)this.each(K(r,e[r]));return this}return this.each(K(e,r))},Y.classed=function(t,e){if(arguments.length<2){if("string"==typeof t){var r=this.node(),n=(t=tt(t)).length,i=-1;if(e=r.classList){for(;++i<n;)if(!e.contains(t[i]))return!1}else for(e=r.getAttribute("class");++i<n;)if(!$(t[i]).test(e))return!1;return!0}for(e in t)this.each(et(e,t[e]));return this}return this.each(et(t,e))},Y.style=function(t,e,r){var n=arguments.length;if(n<3){if("string"!=typeof t){for(r in n<2&&(e=""),t)this.each(nt(r,t[r],e));return this}if(n<2){var i=this.node();return o(i).getComputedStyle(i,null).getPropertyValue(t)}r=""}return this.each(nt(t,e,r))},Y.property=function(t,e){if(arguments.length<2){if("string"==typeof t)return this.node()[t];for(e in t)this.each(it(e,t[e]));return this}return this.each(it(t,e))},Y.text=function(t){return arguments.length?this.each("function"==typeof t?function(){var e=t.apply(this,arguments);this.textContent=null==e?"":e}:null==t?function(){this.textContent=""}:function(){this.textContent=t}):this.node().textContent},Y.html=function(t){return arguments.length?this.each("function"==typeof t?function(){var e=t.apply(this,arguments);this.innerHTML=null==e?"":e}:null==t?function(){this.innerHTML=""}:function(){this.innerHTML=t}):this.node().innerHTML},Y.append=function(t){return t=at(t),this.select((function(){return this.appendChild(t.apply(this,arguments))}))},Y.insert=function(t,e){return t=at(t),e=W(e),this.select((function(){return this.insertBefore(t.apply(this,arguments),e.apply(this,arguments)||null)}))},Y.remove=function(){return this.each(ot)},Y.data=function(t,e){var r,n,i=-1,a=this.length;if(!arguments.length){for(t=new Array(a=(r=this[0]).length);++i<a;)(n=r[i])&&(t[i]=n.__data__);return t}function o(t,r){var n,i,a,o=t.length,u=r.length,f=Math.min(o,u),h=new Array(u),p=new Array(u),d=new Array(o);if(e){var m,g=new _,v=new Array(o);for(n=-1;++n<o;)(i=t[n])&&(g.has(m=e.call(i,i.__data__,n))?d[n]=i:g.set(m,i),v[n]=m);for(n=-1;++n<u;)(i=g.get(m=e.call(r,a=r[n],n)))?!0!==i&&(h[n]=i,i.__data__=a):p[n]=st(a),g.set(m,!0);for(n=-1;++n<o;)n in v&&!0!==g.get(v[n])&&(d[n]=t[n])}else{for(n=-1;++n<f;)i=t[n],a=r[n],i?(i.__data__=a,h[n]=i):p[n]=st(a);for(;n<u;++n)p[n]=st(r[n]);for(;n<o;++n)d[n]=t[n]}p.update=h,p.parentNode=h.parentNode=d.parentNode=t.parentNode,s.push(p),l.push(h),c.push(d)}var s=ft([]),l=V([]),c=V([]);if("function"==typeof t)for(;++i<a;)o(r=this[i],t.call(r,r.parentNode.__data__,i));else for(;++i<a;)o(r=this[i],t);return l.enter=function(){return s},l.exit=function(){return c},l},Y.datum=function(t){return arguments.length?this.property("__data__",t):this.property("__data__")},Y.filter=function(t){var e,r,n,i=[];"function"!=typeof t&&(t=lt(t));for(var a=0,o=this.length;a<o;a++){i.push(e=[]),e.parentNode=(r=this[a]).parentNode;for(var s=0,l=r.length;s<l;s++)(n=r[s])&&t.call(n,n.__data__,s,a)&&e.push(n)}return V(i)},Y.order=function(){for(var t=-1,e=this.length;++t<e;)for(var r,n=this[t],i=n.length-1,a=n[i];--i>=0;)(r=n[i])&&(a&&a!==r.nextSibling&&a.parentNode.insertBefore(r,a),a=r);return this},Y.sort=function(t){t=ct.apply(this,arguments);for(var e=-1,r=this.length;++e<r;)this[e].sort(t);return this.order()},Y.each=function(t){return ut(this,(function(e,r,n){t.call(e,e.__data__,r,n)}))},Y.call=function(t){var e=n(arguments);return t.apply(e[0]=this,e),this},Y.empty=function(){return!this.node()},Y.node=function(){for(var t=0,e=this.length;t<e;t++)for(var r=this[t],n=0,i=r.length;n<i;n++){var a=r[n];if(a)return a}return null},Y.size=function(){var t=0;return ut(this,(function(){++t})),t};var ht=[];function pt(t){var e,r;return function(n,i,a){var o,s=t[a].update,l=s.length;for(a!=r&&(r=a,e=0),i>=e&&(e=i+1);!(o=s[e])&&++e<l;);return o}}function dt(e,r,i){var a="__on"+e,o=e.indexOf("."),s=gt;o>0&&(e=e.slice(0,o));var l=mt.get(e);function c(){var t=this[a];t&&(this.removeEventListener(e,t,t.$),delete this[a])}return l&&(e=l,s=vt),o?r?function(){var t=s(r,n(arguments));c.call(this),this.addEventListener(e,this[a]=t,t.$=i),t._=r}:c:r?z:function(){var r,n=new RegExp("^__on([^.]+)"+t.requote(e)+"$");for(var i in this)if(r=i.match(n)){var a=this[i];this.removeEventListener(r[1],a,a.$),delete this[i]}}}t.selection.enter=ft,t.selection.enter.prototype=ht,ht.append=Y.append,ht.empty=Y.empty,ht.node=Y.node,ht.call=Y.call,ht.size=Y.size,ht.select=function(t){for(var e,r,n,i,a,o=[],s=-1,l=this.length;++s<l;){n=(i=this[s]).update,o.push(e=[]),e.parentNode=i.parentNode;for(var c=-1,u=i.length;++c<u;)(a=i[c])?(e.push(n[c]=r=t.call(i.parentNode,a.__data__,c,s)),r.__data__=a.__data__):e.push(null)}return V(o)},ht.insert=function(t,e){return arguments.length<2&&(e=pt(this)),Y.insert.call(this,t,e)},t.select=function(t){var e;return"string"==typeof t?(e=[H(t,i)]).parentNode=i.documentElement:(e=[t]).parentNode=a(t),V([e])},t.selectAll=function(t){var e;return"string"==typeof t?(e=n(q(t,i))).parentNode=i.documentElement:(e=n(t)).parentNode=null,V([e])},Y.on=function(t,e,r){var n=arguments.length;if(n<3){if("string"!=typeof t){for(r in n<2&&(e=!1),t)this.each(dt(r,t[r],e));return this}if(n<2)return(n=this.node()["__on"+t])&&n._;r=!1}return this.each(dt(t,e,r))};var mt=t.map({mouseenter:"mouseover",mouseleave:"mouseout"});function gt(e,r){return function(n){var i=t.event;t.event=n,r[0]=this.__data__;try{e.apply(this,r)}finally{t.event=i}}}function vt(t,e){var r=gt(t,e);return function(t){var e=t.relatedTarget;e&&(e===this||8&e.compareDocumentPosition(this))||r.call(this,t)}}i&&mt.forEach((function(t){"on"+t in i&&mt.remove(t)}));var yt,xt=0;function bt(e){var r=".dragsuppress-"+ ++xt,n="click"+r,i=t.select(o(e)).on("touchmove"+r,F).on("dragstart"+r,F).on("selectstart"+r,F);if(null==yt&&(yt=!("onselectstart"in e)&&I(e.style,"userSelect")),yt){var s=a(e).style,l=s[yt];s[yt]="none"}return function(t){if(i.on(r,null),yt&&(s[yt]=l),t){var e=function(){i.on(n,null)};i.on(n,(function(){F(),e()}),!0),setTimeout(e,0)}}}t.mouse=function(t){return wt(t,B())};var _t=this.navigator&&/WebKit/.test(this.navigator.userAgent)?-1:0;function wt(e,r){r.changedTouches&&(r=r.changedTouches[0]);var n=e.ownerSVGElement||e;if(n.createSVGPoint){var i=n.createSVGPoint();if(_t<0){var a=o(e);if(a.scrollX||a.scrollY){var s=(n=t.select("body").append("svg").style({position:"absolute",top:0,left:0,margin:0,padding:0,border:"none"},"important"))[0][0].getScreenCTM();_t=!(s.f||s.e),n.remove()}}return _t?(i.x=r.pageX,i.y=r.pageY):(i.x=r.clientX,i.y=r.clientY),[(i=i.matrixTransform(e.getScreenCTM().inverse())).x,i.y]}var l=e.getBoundingClientRect();return[r.clientX-l.left-e.clientLeft,r.clientY-l.top-e.clientTop]}function Tt(){return t.event.changedTouches[0].identifier}t.touch=function(t,e,r){if(arguments.length<3&&(r=e,e=B().changedTouches),e)for(var n,i=0,a=e.length;i<a;++i)if((n=e[i]).identifier===r)return wt(t,n)},t.behavior.drag=function(){var e=N(a,"drag","dragstart","dragend"),r=null,n=s(z,t.mouse,o,"mousemove","mouseup"),i=s(Tt,t.touch,C,"touchmove","touchend");function a(){this.on("mousedown.drag",n).on("touchstart.drag",i)}function s(n,i,a,o,s){return function(){var l,c=this,u=t.event.target.correspondingElement||t.event.target,f=c.parentNode,h=e.of(c,arguments),p=0,d=n(),m=".drag"+(null==d?"":"-"+d),g=t.select(a(u)).on(o+m,x).on(s+m,b),v=bt(u),y=i(f,d);function x(){var t,e,r=i(f,d);r&&(t=r[0]-y[0],e=r[1]-y[1],p|=t|e,y=r,h({type:"drag",x:r[0]+l[0],y:r[1]+l[1],dx:t,dy:e}))}function b(){i(f,d)&&(g.on(o+m,null).on(s+m,null),v(p),h({type:"dragend"}))}l=r?[(l=r.apply(c,arguments)).x-y[0],l.y-y[1]]:[0,0],h({type:"dragstart"})}}return a.origin=function(t){return arguments.length?(r=t,a):r},t.rebind(a,e,"on")},t.touches=function(t,e){return arguments.length<2&&(e=B().touches),e?n(e).map((function(e){var r=wt(t,e);return r.identifier=e.identifier,r})):[]};var kt=1e-6,At=Math.PI,Mt=2*At,St=Mt-kt,Et=At/2,Lt=At/180,Ct=180/At;function Pt(t){return t>1?Et:t<-1?-Et:Math.asin(t)}function It(t){return((t=Math.exp(t))+1/t)/2}var Ot=Math.SQRT2;t.interpolateZoom=function(t,e){var r,n,i=t[0],a=t[1],o=t[2],s=e[0],l=e[1],c=e[2],u=s-i,f=l-a,h=u*u+f*f;if(h<1e-12)n=Math.log(c/o)/Ot,r=function(t){return[i+t*u,a+t*f,o*Math.exp(Ot*t*n)]};else{var p=Math.sqrt(h),d=(c*c-o*o+4*h)/(2*o*2*p),m=(c*c-o*o-4*h)/(2*c*2*p),g=Math.log(Math.sqrt(d*d+1)-d),v=Math.log(Math.sqrt(m*m+1)-m);n=(v-g)/Ot,r=function(t){var e,r=t*n,s=It(g),l=o/(2*p)*(s*(e=Ot*r+g,((e=Math.exp(2*e))-1)/(e+1))-function(t){return((t=Math.exp(t))-1/t)/2}(g));return[i+l*u,a+l*f,o*s/It(Ot*r+g)]}}return r.duration=1e3*n,r},t.behavior.zoom=function(){var e,r,n,a,s,l,c,u,f,h={x:0,y:0,k:1},p=[960,500],d=Rt,m=250,g=0,v="mousedown.zoom",y="mousemove.zoom",x="mouseup.zoom",b="touchstart.zoom",_=N(w,"zoomstart","zoom","zoomend");function w(t){t.on(v,P).on(Dt+".zoom",O).on("dblclick.zoom",z).on(b,I)}function T(t){return[(t[0]-h.x)/h.k,(t[1]-h.y)/h.k]}function k(t){h.k=Math.max(d[0],Math.min(d[1],t))}function A(t,e){e=function(t){return[t[0]*h.k+h.x,t[1]*h.k+h.y]}(e),h.x+=t[0]-e[0],h.y+=t[1]-e[1]}function M(e,n,i,a){e.__chart__={x:h.x,y:h.y,k:h.k},k(Math.pow(2,a)),A(r=n,i),e=t.select(e),m>0&&(e=e.transition().duration(m)),e.call(w.event)}function S(){c&&c.domain(l.range().map((function(t){return(t-h.x)/h.k})).map(l.invert)),f&&f.domain(u.range().map((function(t){return(t-h.y)/h.k})).map(u.invert))}function E(t){g++||t({type:"zoomstart"})}function L(t){S(),t({type:"zoom",scale:h.k,translate:[h.x,h.y]})}function C(t){--g||(t({type:"zoomend"}),r=null)}function P(){var e=this,r=_.of(e,arguments),n=0,i=t.select(o(e)).on(y,l).on(x,c),a=T(t.mouse(e)),s=bt(e);function l(){n=1,A(t.mouse(e),a),L(r)}function c(){i.on(y,null).on(x,null),s(n),C(r)}Di.call(e),E(r)}function I(){var e,r=this,n=_.of(r,arguments),i={},a=0,o=".zoom-"+t.event.changedTouches[0].identifier,l="touchmove"+o,c="touchend"+o,u=[],f=t.select(r),p=bt(r);function d(){var n=t.touches(r);return e=h.k,n.forEach((function(t){t.identifier in i&&(i[t.identifier]=T(t))})),n}function m(){var e=t.event.target;t.select(e).on(l,g).on(c,y),u.push(e);for(var n=t.event.changedTouches,o=0,f=n.length;o<f;++o)i[n[o].identifier]=null;var p=d(),m=Date.now();if(1===p.length){if(m-s<500){var v=p[0];M(r,v,i[v.identifier],Math.floor(Math.log(h.k)/Math.LN2)+1),F()}s=m}else if(p.length>1){v=p[0];var x=p[1],b=v[0]-x[0],_=v[1]-x[1];a=b*b+_*_}}function g(){var o,l,c,u,f=t.touches(r);Di.call(r);for(var h=0,p=f.length;h<p;++h,u=null)if(c=f[h],u=i[c.identifier]){if(l)break;o=c,l=u}if(u){var d=(d=c[0]-o[0])*d+(d=c[1]-o[1])*d,m=a&&Math.sqrt(d/a);o=[(o[0]+c[0])/2,(o[1]+c[1])/2],l=[(l[0]+u[0])/2,(l[1]+u[1])/2],k(m*e)}s=null,A(o,l),L(n)}function y(){if(t.event.touches.length){for(var e=t.event.changedTouches,r=0,a=e.length;r<a;++r)delete i[e[r].identifier];for(var s in i)return void d()}t.selectAll(u).on(o,null),f.on(v,P).on(b,I),p(),C(n)}m(),E(n),f.on(v,null).on(b,m)}function O(){var i=_.of(this,arguments);a?clearTimeout(a):(Di.call(this),e=T(r=n||t.mouse(this)),E(i)),a=setTimeout((function(){a=null,C(i)}),50),F(),k(Math.pow(2,.002*zt())*h.k),A(r,e),L(i)}function z(){var e=t.mouse(this),r=Math.log(h.k)/Math.LN2;M(this,e,T(e),t.event.shiftKey?Math.ceil(r)-1:Math.floor(r)+1)}return Dt||(Dt="onwheel"in i?(zt=function(){return-t.event.deltaY*(t.event.deltaMode?120:1)},"wheel"):"onmousewheel"in i?(zt=function(){return t.event.wheelDelta},"mousewheel"):(zt=function(){return-t.event.detail},"MozMousePixelScroll")),w.event=function(e){e.each((function(){var e=_.of(this,arguments),n=h;Bi?t.select(this).transition().each("start.zoom",(function(){h=this.__chart__||{x:0,y:0,k:1},E(e)})).tween("zoom:zoom",(function(){var i=p[0],a=p[1],o=r?r[0]:i/2,s=r?r[1]:a/2,l=t.interpolateZoom([(o-h.x)/h.k,(s-h.y)/h.k,i/h.k],[(o-n.x)/n.k,(s-n.y)/n.k,i/n.k]);return function(t){var r=l(t),n=i/r[2];this.__chart__=h={x:o-r[0]*n,y:s-r[1]*n,k:n},L(e)}})).each("interrupt.zoom",(function(){C(e)})).each("end.zoom",(function(){C(e)})):(this.__chart__=h,E(e),L(e),C(e))}))},w.translate=function(t){return arguments.length?(h={x:+t[0],y:+t[1],k:h.k},S(),w):[h.x,h.y]},w.scale=function(t){return arguments.length?(h={x:h.x,y:h.y,k:null},k(+t),S(),w):h.k},w.scaleExtent=function(t){return arguments.length?(d=null==t?Rt:[+t[0],+t[1]],w):d},w.center=function(t){return arguments.length?(n=t&&[+t[0],+t[1]],w):n},w.size=function(t){return arguments.length?(p=t&&[+t[0],+t[1]],w):p},w.duration=function(t){return arguments.length?(m=+t,w):m},w.x=function(t){return arguments.length?(c=t,l=t.copy(),h={x:0,y:0,k:1},w):c},w.y=function(t){return arguments.length?(f=t,u=t.copy(),h={x:0,y:0,k:1},w):f},t.rebind(w,_,"on")};var zt,Dt,Rt=[0,1/0];function Ft(){}function Bt(t,e,r){return this instanceof Bt?(this.h=+t,this.s=+e,void(this.l=+r)):arguments.length<2?t instanceof Bt?new Bt(t.h,t.s,t.l):ne(""+t,ie,Bt):new Bt(t,e,r)}t.color=Ft,Ft.prototype.toString=function(){return this.rgb()+""},t.hsl=Bt;var Nt=Bt.prototype=new Ft;function jt(t,e,r){var n,i;function a(t){return Math.round(255*function(t){return t>360?t-=360:t<0&&(t+=360),t<60?n+(i-n)*t/60:t<180?i:t<240?n+(i-n)*(240-t)/60:n}(t))}return t=isNaN(t)?0:(t%=360)<0?t+360:t,e=isNaN(e)||e<0?0:e>1?1:e,n=2*(r=r<0?0:r>1?1:r)-(i=r<=.5?r*(1+e):r+e-r*e),new Qt(a(t+120),a(t),a(t-120))}function Ut(e,r,n){return this instanceof Ut?(this.h=+e,this.c=+r,void(this.l=+n)):arguments.length<2?e instanceof Ut?new Ut(e.h,e.c,e.l):Xt(e instanceof qt?e.l:(e=ae((e=t.rgb(e)).r,e.g,e.b)).l,e.a,e.b):new Ut(e,r,n)}Nt.brighter=function(t){return t=Math.pow(.7,arguments.length?t:1),new Bt(this.h,this.s,this.l/t)},Nt.darker=function(t){return t=Math.pow(.7,arguments.length?t:1),new Bt(this.h,this.s,t*this.l)},Nt.rgb=function(){return jt(this.h,this.s,this.l)},t.hcl=Ut;var Vt=Ut.prototype=new Ft;function Ht(t,e,r){return isNaN(t)&&(t=0),isNaN(e)&&(e=0),new qt(r,Math.cos(t*=Lt)*e,Math.sin(t)*e)}function qt(t,e,r){return this instanceof qt?(this.l=+t,this.a=+e,void(this.b=+r)):arguments.length<2?t instanceof qt?new qt(t.l,t.a,t.b):t instanceof Ut?Ht(t.h,t.c,t.l):ae((t=Qt(t)).r,t.g,t.b):new qt(t,e,r)}Vt.brighter=function(t){return new Ut(this.h,this.c,Math.min(100,this.l+Gt*(arguments.length?t:1)))},Vt.darker=function(t){return new Ut(this.h,this.c,Math.max(0,this.l-Gt*(arguments.length?t:1)))},Vt.rgb=function(){return Ht(this.h,this.c,this.l).rgb()},t.lab=qt;var Gt=18,Yt=qt.prototype=new Ft;function Wt(t,e,r){var n=(t+16)/116,i=n+e/500,a=n-r/200;return new Qt(Kt(3.2404542*(i=.95047*Zt(i))-1.5371385*(n=1*Zt(n))-.4985314*(a=1.08883*Zt(a))),Kt(-.969266*i+1.8760108*n+.041556*a),Kt(.0556434*i-.2040259*n+1.0572252*a))}function Xt(t,e,r){return t>0?new Ut(Math.atan2(r,e)*Ct,Math.sqrt(e*e+r*r),t):new Ut(NaN,NaN,t)}function Zt(t){return t>.206893034?t*t*t:(t-4/29)/7.787037}function Jt(t){return t>.008856?Math.pow(t,1/3):7.787037*t+4/29}function Kt(t){return Math.round(255*(t<=.00304?12.92*t:1.055*Math.pow(t,1/2.4)-.055))}function Qt(t,e,r){return this instanceof Qt?(this.r=~~t,this.g=~~e,void(this.b=~~r)):arguments.length<2?t instanceof Qt?new Qt(t.r,t.g,t.b):ne(""+t,Qt,jt):new Qt(t,e,r)}function $t(t){return new Qt(t>>16,t>>8&255,255&t)}function te(t){return $t(t)+""}Yt.brighter=function(t){return new qt(Math.min(100,this.l+Gt*(arguments.length?t:1)),this.a,this.b)},Yt.darker=function(t){return new qt(Math.max(0,this.l-Gt*(arguments.length?t:1)),this.a,this.b)},Yt.rgb=function(){return Wt(this.l,this.a,this.b)},t.rgb=Qt;var ee=Qt.prototype=new Ft;function re(t){return t<16?"0"+Math.max(0,t).toString(16):Math.min(255,t).toString(16)}function ne(t,e,r){var n,i,a,o=0,s=0,l=0;if(n=/([a-z]+)\((.*)\)/.exec(t=t.toLowerCase()))switch(i=n[2].split(","),n[1]){case"hsl":return r(parseFloat(i[0]),parseFloat(i[1])/100,parseFloat(i[2])/100);case"rgb":return e(se(i[0]),se(i[1]),se(i[2]))}return(a=le.get(t))?e(a.r,a.g,a.b):(null==t||"#"!==t.charAt(0)||isNaN(a=parseInt(t.slice(1),16))||(4===t.length?(o=(3840&a)>>4,o|=o>>4,s=240&a,s|=s>>4,l=15&a,l|=l<<4):7===t.length&&(o=(16711680&a)>>16,s=(65280&a)>>8,l=255&a)),e(o,s,l))}function ie(t,e,r){var n,i,a=Math.min(t/=255,e/=255,r/=255),o=Math.max(t,e,r),s=o-a,l=(o+a)/2;return s?(i=l<.5?s/(o+a):s/(2-o-a),n=t==o?(e-r)/s+(e<r?6:0):e==o?(r-t)/s+2:(t-e)/s+4,n*=60):(n=NaN,i=l>0&&l<1?0:n),new Bt(n,i,l)}function ae(t,e,r){var n=Jt((.4124564*(t=oe(t))+.3575761*(e=oe(e))+.1804375*(r=oe(r)))/.95047),i=Jt((.2126729*t+.7151522*e+.072175*r)/1);return qt(116*i-16,500*(n-i),200*(i-Jt((.0193339*t+.119192*e+.9503041*r)/1.08883)))}function oe(t){return(t/=255)<=.04045?t/12.92:Math.pow((t+.055)/1.055,2.4)}function se(t){var e=parseFloat(t);return"%"===t.charAt(t.length-1)?Math.round(2.55*e):e}ee.brighter=function(t){t=Math.pow(.7,arguments.length?t:1);var e=this.r,r=this.g,n=this.b,i=30;return e||r||n?(e&&e<i&&(e=i),r&&r<i&&(r=i),n&&n<i&&(n=i),new Qt(Math.min(255,e/t),Math.min(255,r/t),Math.min(255,n/t))):new Qt(i,i,i)},ee.darker=function(t){return new Qt((t=Math.pow(.7,arguments.length?t:1))*this.r,t*this.g,t*this.b)},ee.hsl=function(){return ie(this.r,this.g,this.b)},ee.toString=function(){return"#"+re(this.r)+re(this.g)+re(this.b)};var le=t.map({aliceblue:15792383,antiquewhite:16444375,aqua:65535,aquamarine:8388564,azure:15794175,beige:16119260,bisque:16770244,black:0,blanchedalmond:16772045,blue:255,blueviolet:9055202,brown:10824234,burlywood:14596231,cadetblue:6266528,chartreuse:8388352,chocolate:13789470,coral:16744272,cornflowerblue:6591981,cornsilk:16775388,crimson:14423100,cyan:65535,darkblue:139,darkcyan:35723,darkgoldenrod:12092939,darkgray:11119017,darkgreen:25600,darkgrey:11119017,darkkhaki:12433259,darkmagenta:9109643,darkolivegreen:5597999,darkorange:16747520,darkorchid:10040012,darkred:9109504,darksalmon:15308410,darkseagreen:9419919,darkslateblue:4734347,darkslategray:3100495,darkslategrey:3100495,darkturquoise:52945,darkviolet:9699539,deeppink:16716947,deepskyblue:49151,dimgray:6908265,dimgrey:6908265,dodgerblue:2003199,firebrick:11674146,floralwhite:16775920,forestgreen:2263842,fuchsia:16711935,gainsboro:14474460,ghostwhite:16316671,gold:16766720,goldenrod:14329120,gray:8421504,green:32768,greenyellow:11403055,grey:8421504,honeydew:15794160,hotpink:16738740,indianred:13458524,indigo:4915330,ivory:16777200,khaki:15787660,lavender:15132410,lavenderblush:16773365,lawngreen:8190976,lemonchiffon:16775885,lightblue:11393254,lightcoral:15761536,lightcyan:14745599,lightgoldenrodyellow:16448210,lightgray:13882323,lightgreen:9498256,lightgrey:13882323,lightpink:16758465,lightsalmon:16752762,lightseagreen:2142890,lightskyblue:8900346,lightslategray:7833753,lightslategrey:7833753,lightsteelblue:11584734,lightyellow:16777184,lime:65280,limegreen:3329330,linen:16445670,magenta:16711935,maroon:8388608,mediumaquamarine:6737322,mediumblue:205,mediumorchid:12211667,mediumpurple:9662683,mediumseagreen:3978097,mediumslateblue:8087790,mediumspringgreen:64154,mediumturquoise:4772300,mediumvioletred:13047173,midnightblue:1644912,mintcream:16121850,mistyrose:16770273,moccasin:16770229,navajowhite:16768685,navy:128,oldlace:16643558,olive:8421376,olivedrab:7048739,orange:16753920,orangered:16729344,orchid:14315734,palegoldenrod:15657130,palegreen:10025880,paleturquoise:11529966,palevioletred:14381203,papayawhip:16773077,peachpuff:16767673,peru:13468991,pink:16761035,plum:14524637,powderblue:11591910,purple:8388736,rebeccapurple:6697881,red:16711680,rosybrown:12357519,royalblue:4286945,saddlebrown:9127187,salmon:16416882,sandybrown:16032864,seagreen:3050327,seashell:16774638,sienna:10506797,silver:12632256,skyblue:8900331,slateblue:6970061,slategray:7372944,slategrey:7372944,snow:16775930,springgreen:65407,steelblue:4620980,tan:13808780,teal:32896,thistle:14204888,tomato:16737095,turquoise:4251856,violet:15631086,wheat:16113331,white:16777215,whitesmoke:16119285,yellow:16776960,yellowgreen:10145074});function ce(t){return"function"==typeof t?t:function(){return t}}function ue(t){return function(e,r,n){return 2===arguments.length&&"function"==typeof r&&(n=r,r=null),fe(e,r,t,n)}}function fe(e,r,i,a){var o={},s=t.dispatch("beforesend","progress","load","error"),l={},c=new XMLHttpRequest,u=null;function f(){var t,e=c.status;if(!e&&function(t){var e=t.responseType;return e&&"text"!==e?t.response:t.responseText}(c)||e>=200&&e<300||304===e){try{t=i.call(o,c)}catch(t){return void s.error.call(o,t)}s.load.call(o,t)}else s.error.call(o,c)}return self.XDomainRequest&&!("withCredentials"in c)&&/^(http(s)?:)?\/\//.test(e)&&(c=new XDomainRequest),"onload"in c?c.onload=c.onerror=f:c.onreadystatechange=function(){c.readyState>3&&f()},c.onprogress=function(e){var r=t.event;t.event=e;try{s.progress.call(o,c)}finally{t.event=r}},o.header=function(t,e){return t=(t+"").toLowerCase(),arguments.length<2?l[t]:(null==e?delete l[t]:l[t]=e+"",o)},o.mimeType=function(t){return arguments.length?(r=null==t?null:t+"",o):r},o.responseType=function(t){return arguments.length?(u=t,o):u},o.response=function(t){return i=t,o},["get","post"].forEach((function(t){o[t]=function(){return o.send.apply(o,[t].concat(n(arguments)))}})),o.send=function(t,n,i){if(2===arguments.length&&"function"==typeof n&&(i=n,n=null),c.open(t,e,!0),null==r||"accept"in l||(l.accept=r+",*/*"),c.setRequestHeader)for(var a in l)c.setRequestHeader(a,l[a]);return null!=r&&c.overrideMimeType&&c.overrideMimeType(r),null!=u&&(c.responseType=u),null!=i&&o.on("error",i).on("load",(function(t){i(null,t)})),s.beforesend.call(o,c),c.send(null==n?null:n),o},o.abort=function(){return c.abort(),o},t.rebind(o,s,"on"),null==a?o:o.get(function(t){return 1===t.length?function(e,r){t(null==e?r:null)}:t}(a))}le.forEach((function(t,e){le.set(t,$t(e))})),t.functor=ce,t.xhr=ue(C),t.dsv=function(t,e){var r=new RegExp('["'+t+"\n]"),n=t.charCodeAt(0);function i(t,r,n){arguments.length<3&&(n=r,r=null);var i=fe(t,e,null==r?a:o(r),n);return i.row=function(t){return arguments.length?i.response(null==(r=t)?a:o(t)):r},i}function a(t){return i.parse(t.responseText)}function o(t){return function(e){return i.parse(e.responseText,t)}}function s(e){return e.map(l).join(t)}function l(t){return r.test(t)?'"'+t.replace(/\"/g,'""')+'"':t}return i.parse=function(t,e){var r;return i.parseRows(t,(function(t,n){if(r)return r(t,n-1);var i=function(e){for(var r={},n=t.length,i=0;i<n;++i)r[t[i]]=e[i];return r};r=e?function(t,r){return e(i(t),r)}:i}))},i.parseRows=function(t,e){var r,i,a={},o={},s=[],l=t.length,c=0,u=0;function f(){if(c>=l)return o;if(i)return i=!1,a;var e=c;if(34===t.charCodeAt(e)){for(var r=e;r++<l;)if(34===t.charCodeAt(r)){if(34!==t.charCodeAt(r+1))break;++r}return c=r+2,13===(s=t.charCodeAt(r+1))?(i=!0,10===t.charCodeAt(r+2)&&++c):10===s&&(i=!0),t.slice(e+1,r).replace(/""/g,'"')}for(;c<l;){var s,u=1;if(10===(s=t.charCodeAt(c++)))i=!0;else if(13===s)i=!0,10===t.charCodeAt(c)&&(++c,++u);else if(s!==n)continue;return t.slice(e,c-u)}return t.slice(e)}for(;(r=f())!==o;){for(var h=[];r!==a&&r!==o;)h.push(r),r=f();e&&null==(h=e(h,u++))||s.push(h)}return s},i.format=function(e){if(Array.isArray(e[0]))return i.formatRows(e);var r=new L,n=[];return e.forEach((function(t){for(var e in t)r.has(e)||n.push(r.add(e))})),[n.map(l).join(t)].concat(e.map((function(e){return n.map((function(t){return l(e[t])})).join(t)}))).join("\n")},i.formatRows=function(t){return t.map(s).join("\n")},i},t.csv=t.dsv(",","text/csv"),t.tsv=t.dsv("\t","text/tab-separated-values");var he,pe,de,me,ge=this[I(this,"requestAnimationFrame")]||function(t){setTimeout(t,17)};function ve(t,e,r){var n=arguments.length;n<2&&(e=0),n<3&&(r=Date.now());var i=r+e,a={c:t,t:i,n:null};return pe?pe.n=a:he=a,pe=a,de||(me=clearTimeout(me),de=1,ge(ye)),a}function ye(){var t=xe(),e=be()-t;e>24?(isFinite(e)&&(clearTimeout(me),me=setTimeout(ye,e)),de=0):(de=1,ge(ye))}function xe(){for(var t=Date.now(),e=he;e;)t>=e.t&&e.c(t-e.t)&&(e.c=null),e=e.n;return t}function be(){for(var t,e=he,r=1/0;e;)e.c?(e.t<r&&(r=e.t),e=(t=e).n):e=t?t.n=e.n:he=e.n;return pe=t,r}function _e(t){return t[0]}function we(t){return t[1]}function Te(t){for(var e,r,n,i=t.length,a=[0,1],o=2,s=2;s<i;s++){for(;o>1&&(e=t[a[o-2]],r=t[a[o-1]],n=t[s],(r[0]-e[0])*(n[1]-e[1])-(r[1]-e[1])*(n[0]-e[0])<=0);)--o;a[o++]=s}return a.slice(0,o)}function ke(t,e){return t[0]-e[0]||t[1]-e[1]}t.timer=function(){ve.apply(this,arguments)},t.timer.flush=function(){xe(),be()},t.round=function(t,e){return e?Math.round(t*(e=Math.pow(10,e)))/e:Math.round(t)},t.geom={},t.geom.hull=function(t){var e=_e,r=we;if(arguments.length)return n(t);function n(t){if(t.length<3)return[];var n,i=ce(e),a=ce(r),o=t.length,s=[],l=[];for(n=0;n<o;n++)s.push([+i.call(this,t[n],n),+a.call(this,t[n],n),n]);for(s.sort(ke),n=0;n<o;n++)l.push([s[n][0],-s[n][1]]);var c=Te(s),u=Te(l),f=u[0]===c[0],h=u[u.length-1]===c[c.length-1],p=[];for(n=c.length-1;n>=0;--n)p.push(t[s[c[n]][2]]);for(n=+f;n<u.length-h;++n)p.push(t[s[u[n]][2]]);return p}return n.x=function(t){return arguments.length?(e=t,n):e},n.y=function(t){return arguments.length?(r=t,n):r},n},t.geom.polygon=function(t){return U(t,Ae),t};var Ae=t.geom.polygon.prototype=[];function Me(t,e,r){return(r[0]-e[0])*(t[1]-e[1])<(r[1]-e[1])*(t[0]-e[0])}function Se(t,e,r,n){var i=t[0],a=r[0],o=e[0]-i,s=n[0]-a,l=t[1],c=r[1],u=e[1]-l,f=n[1]-c,h=(s*(l-c)-f*(i-a))/(f*o-s*u);return[i+h*o,l+h*u]}function Ee(t){var e=t[0],r=t[t.length-1];return!(e[0]-r[0]||e[1]-r[1])}Ae.area=function(){for(var t,e=-1,r=this.length,n=this[r-1],i=0;++e<r;)t=n,n=this[e],i+=t[1]*n[0]-t[0]*n[1];return.5*i},Ae.centroid=function(t){var e,r,n=-1,i=this.length,a=0,o=0,s=this[i-1];for(arguments.length||(t=-1/(6*this.area()));++n<i;)e=s,s=this[n],r=e[0]*s[1]-s[0]*e[1],a+=(e[0]+s[0])*r,o+=(e[1]+s[1])*r;return[a*t,o*t]},Ae.clip=function(t){for(var e,r,n,i,a,o,s=Ee(t),l=-1,c=this.length-Ee(this),u=this[c-1];++l<c;){for(e=t.slice(),t.length=0,i=this[l],a=e[(n=e.length-s)-1],r=-1;++r<n;)Me(o=e[r],u,i)?(Me(a,u,i)||t.push(Se(a,o,u,i)),t.push(o)):Me(a,u,i)&&t.push(Se(a,o,u,i)),a=o;s&&t.push(t[0]),u=i}return t};var Le,Ce,Pe,Ie,Oe,ze=[],De=[];function Re(){er(this),this.edge=this.site=this.circle=null}function Fe(t){var e=ze.pop()||new Re;return e.site=t,e}function Be(t){We(t),Pe.remove(t),ze.push(t),er(t)}function Ne(t){var e=t.circle,r=e.x,n=e.cy,i={x:r,y:n},a=t.P,o=t.N,s=[t];Be(t);for(var l=a;l.circle&&y(r-l.circle.x)<kt&&y(n-l.circle.cy)<kt;)a=l.P,s.unshift(l),Be(l),l=a;s.unshift(l),We(l);for(var c=o;c.circle&&y(r-c.circle.x)<kt&&y(n-c.circle.cy)<kt;)o=c.N,s.push(c),Be(c),c=o;s.push(c),We(c);var u,f=s.length;for(u=1;u<f;++u)c=s[u],l=s[u-1],Qe(c.edge,l.site,c.site,i);l=s[0],(c=s[f-1]).edge=Je(l.site,c.site,null,i),Ye(l),Ye(c)}function je(t){for(var e,r,n,i,a=t.x,o=t.y,s=Pe._;s;)if((n=Ue(s,o)-a)>kt)s=s.L;else{if(!((i=a-Ve(s,o))>kt)){n>-kt?(e=s.P,r=s):i>-kt?(e=s,r=s.N):e=r=s;break}if(!s.R){e=s;break}s=s.R}var l=Fe(t);if(Pe.insert(e,l),e||r){if(e===r)return We(e),r=Fe(e.site),Pe.insert(l,r),l.edge=r.edge=Je(e.site,l.site),Ye(e),void Ye(r);if(r){We(e),We(r);var c=e.site,u=c.x,f=c.y,h=t.x-u,p=t.y-f,d=r.site,m=d.x-u,g=d.y-f,v=2*(h*g-p*m),y=h*h+p*p,x=m*m+g*g,b={x:(g*y-p*x)/v+u,y:(h*x-m*y)/v+f};Qe(r.edge,c,d,b),l.edge=Je(c,t,null,b),r.edge=Je(t,d,null,b),Ye(e),Ye(r)}else l.edge=Je(e.site,l.site)}}function Ue(t,e){var r=t.site,n=r.x,i=r.y,a=i-e;if(!a)return n;var o=t.P;if(!o)return-1/0;var s=(r=o.site).x,l=r.y,c=l-e;if(!c)return s;var u=s-n,f=1/a-1/c,h=u/c;return f?(-h+Math.sqrt(h*h-2*f*(u*u/(-2*c)-l+c/2+i-a/2)))/f+n:(n+s)/2}function Ve(t,e){var r=t.N;if(r)return Ue(r,e);var n=t.site;return n.y===e?n.x:1/0}function He(t){this.site=t,this.edges=[]}function qe(t,e){return e.angle-t.angle}function Ge(){er(this),this.x=this.y=this.arc=this.site=this.cy=null}function Ye(t){var e=t.P,r=t.N;if(e&&r){var n=e.site,i=t.site,a=r.site;if(n!==a){var o=i.x,s=i.y,l=n.x-o,c=n.y-s,u=a.x-o,f=2*(l*(g=a.y-s)-c*u);if(!(f>=-1e-12)){var h=l*l+c*c,p=u*u+g*g,d=(g*h-c*p)/f,m=(l*p-u*h)/f,g=m+s,v=De.pop()||new Ge;v.arc=t,v.site=i,v.x=d+o,v.y=g+Math.sqrt(d*d+m*m),v.cy=g,t.circle=v;for(var y=null,x=Oe._;x;)if(v.y<x.y||v.y===x.y&&v.x<=x.x){if(!x.L){y=x.P;break}x=x.L}else{if(!x.R){y=x;break}x=x.R}Oe.insert(y,v),y||(Ie=v)}}}}function We(t){var e=t.circle;e&&(e.P||(Ie=e.N),Oe.remove(e),De.push(e),er(e),t.circle=null)}function Xe(t,e){var r=t.b;if(r)return!0;var n,i,a=t.a,o=e[0][0],s=e[1][0],l=e[0][1],c=e[1][1],u=t.l,f=t.r,h=u.x,p=u.y,d=f.x,m=f.y,g=(h+d)/2,v=(p+m)/2;if(m===p){if(g<o||g>=s)return;if(h>d){if(a){if(a.y>=c)return}else a={x:g,y:l};r={x:g,y:c}}else{if(a){if(a.y<l)return}else a={x:g,y:c};r={x:g,y:l}}}else if(i=v-(n=(h-d)/(m-p))*g,n<-1||n>1)if(h>d){if(a){if(a.y>=c)return}else a={x:(l-i)/n,y:l};r={x:(c-i)/n,y:c}}else{if(a){if(a.y<l)return}else a={x:(c-i)/n,y:c};r={x:(l-i)/n,y:l}}else if(p<m){if(a){if(a.x>=s)return}else a={x:o,y:n*o+i};r={x:s,y:n*s+i}}else{if(a){if(a.x<o)return}else a={x:s,y:n*s+i};r={x:o,y:n*o+i}}return t.a=a,t.b=r,!0}function Ze(t,e){this.l=t,this.r=e,this.a=this.b=null}function Je(t,e,r,n){var i=new Ze(t,e);return Le.push(i),r&&Qe(i,t,e,r),n&&Qe(i,e,t,n),Ce[t.i].edges.push(new $e(i,t,e)),Ce[e.i].edges.push(new $e(i,e,t)),i}function Ke(t,e,r){var n=new Ze(t,null);return n.a=e,n.b=r,Le.push(n),n}function Qe(t,e,r,n){t.a||t.b?t.l===r?t.b=n:t.a=n:(t.a=n,t.l=e,t.r=r)}function $e(t,e,r){var n=t.a,i=t.b;this.edge=t,this.site=e,this.angle=r?Math.atan2(r.y-e.y,r.x-e.x):t.l===e?Math.atan2(i.x-n.x,n.y-i.y):Math.atan2(n.x-i.x,i.y-n.y)}function tr(){this._=null}function er(t){t.U=t.C=t.L=t.R=t.P=t.N=null}function rr(t,e){var r=e,n=e.R,i=r.U;i?i.L===r?i.L=n:i.R=n:t._=n,n.U=i,r.U=n,r.R=n.L,r.R&&(r.R.U=r),n.L=r}function nr(t,e){var r=e,n=e.L,i=r.U;i?i.L===r?i.L=n:i.R=n:t._=n,n.U=i,r.U=n,r.L=n.R,r.L&&(r.L.U=r),n.R=r}function ir(t){for(;t.L;)t=t.L;return t}function ar(t,e){var r,n,i,a=t.sort(or).pop();for(Le=[],Ce=new Array(t.length),Pe=new tr,Oe=new tr;;)if(i=Ie,a&&(!i||a.y<i.y||a.y===i.y&&a.x<i.x))a.x===r&&a.y===n||(Ce[a.i]=new He(a),je(a),r=a.x,n=a.y),a=t.pop();else{if(!i)break;Ne(i.arc)}e&&(function(t){for(var e,r,n,i,a,o=Le,s=(r=t[0][0],n=t[0][1],i=t[1][0],a=t[1][1],function(t){var e,o=t.a,s=t.b,l=o.x,c=o.y,u=0,f=1,h=s.x-l,p=s.y-c;if(e=r-l,h||!(e>0)){if(e/=h,h<0){if(e<u)return;e<f&&(f=e)}else if(h>0){if(e>f)return;e>u&&(u=e)}if(e=i-l,h||!(e<0)){if(e/=h,h<0){if(e>f)return;e>u&&(u=e)}else if(h>0){if(e<u)return;e<f&&(f=e)}if(e=n-c,p||!(e>0)){if(e/=p,p<0){if(e<u)return;e<f&&(f=e)}else if(p>0){if(e>f)return;e>u&&(u=e)}if(e=a-c,p||!(e<0)){if(e/=p,p<0){if(e>f)return;e>u&&(u=e)}else if(p>0){if(e<u)return;e<f&&(f=e)}return u>0&&(t.a={x:l+u*h,y:c+u*p}),f<1&&(t.b={x:l+f*h,y:c+f*p}),t}}}}}),l=o.length;l--;)(!Xe(e=o[l],t)||!s(e)||y(e.a.x-e.b.x)<kt&&y(e.a.y-e.b.y)<kt)&&(e.a=e.b=null,o.splice(l,1))}(e),function(t){for(var e,r,n,i,a,o,s,l,c,u,f=t[0][0],h=t[1][0],p=t[0][1],d=t[1][1],m=Ce,g=m.length;g--;)if((a=m[g])&&a.prepare())for(l=(s=a.edges).length,o=0;o<l;)n=(u=s[o].end()).x,i=u.y,e=(c=s[++o%l].start()).x,r=c.y,(y(n-e)>kt||y(i-r)>kt)&&(s.splice(o,0,new $e(Ke(a.site,u,y(n-f)<kt&&d-i>kt?{x:f,y:y(e-f)<kt?r:d}:y(i-d)<kt&&h-n>kt?{x:y(r-d)<kt?e:h,y:d}:y(n-h)<kt&&i-p>kt?{x:h,y:y(e-h)<kt?r:p}:y(i-p)<kt&&n-f>kt?{x:y(r-p)<kt?e:f,y:p}:null),a.site,null)),++l)}(e));var o={cells:Ce,edges:Le};return Pe=Oe=Le=Ce=null,o}function or(t,e){return e.y-t.y||e.x-t.x}He.prototype.prepare=function(){for(var t,e=this.edges,r=e.length;r--;)(t=e[r].edge).b&&t.a||e.splice(r,1);return e.sort(qe),e.length},$e.prototype={start:function(){return this.edge.l===this.site?this.edge.a:this.edge.b},end:function(){return this.edge.l===this.site?this.edge.b:this.edge.a}},tr.prototype={insert:function(t,e){var r,n,i;if(t){if(e.P=t,e.N=t.N,t.N&&(t.N.P=e),t.N=e,t.R){for(t=t.R;t.L;)t=t.L;t.L=e}else t.R=e;r=t}else this._?(t=ir(this._),e.P=null,e.N=t,t.P=t.L=e,r=t):(e.P=e.N=null,this._=e,r=null);for(e.L=e.R=null,e.U=r,e.C=!0,t=e;r&&r.C;)r===(n=r.U).L?(i=n.R)&&i.C?(r.C=i.C=!1,n.C=!0,t=n):(t===r.R&&(rr(this,r),r=(t=r).U),r.C=!1,n.C=!0,nr(this,n)):(i=n.L)&&i.C?(r.C=i.C=!1,n.C=!0,t=n):(t===r.L&&(nr(this,r),r=(t=r).U),r.C=!1,n.C=!0,rr(this,n)),r=t.U;this._.C=!1},remove:function(t){t.N&&(t.N.P=t.P),t.P&&(t.P.N=t.N),t.N=t.P=null;var e,r,n,i=t.U,a=t.L,o=t.R;if(r=a?o?ir(o):a:o,i?i.L===t?i.L=r:i.R=r:this._=r,a&&o?(n=r.C,r.C=t.C,r.L=a,a.U=r,r!==o?(i=r.U,r.U=t.U,t=r.R,i.L=t,r.R=o,o.U=r):(r.U=i,i=r,t=r.R)):(n=t.C,t=r),t&&(t.U=i),!n)if(t&&t.C)t.C=!1;else{do{if(t===this._)break;if(t===i.L){if((e=i.R).C&&(e.C=!1,i.C=!0,rr(this,i),e=i.R),e.L&&e.L.C||e.R&&e.R.C){e.R&&e.R.C||(e.L.C=!1,e.C=!0,nr(this,e),e=i.R),e.C=i.C,i.C=e.R.C=!1,rr(this,i),t=this._;break}}else if((e=i.L).C&&(e.C=!1,i.C=!0,nr(this,i),e=i.L),e.L&&e.L.C||e.R&&e.R.C){e.L&&e.L.C||(e.R.C=!1,e.C=!0,rr(this,e),e=i.L),e.C=i.C,i.C=e.L.C=!1,nr(this,i),t=this._;break}e.C=!0,t=i,i=i.U}while(!t.C);t&&(t.C=!1)}}},t.geom.voronoi=function(t){var e=_e,r=we,n=e,i=r,a=sr;if(t)return o(t);function o(t){var e=new Array(t.length),r=a[0][0],n=a[0][1],i=a[1][0],o=a[1][1];return ar(s(t),a).cells.forEach((function(a,s){var l=a.edges,c=a.site;(e[s]=l.length?l.map((function(t){var e=t.start();return[e.x,e.y]})):c.x>=r&&c.x<=i&&c.y>=n&&c.y<=o?[[r,o],[i,o],[i,n],[r,n]]:[]).point=t[s]})),e}function s(t){return t.map((function(t,e){return{x:Math.round(n(t,e)/kt)*kt,y:Math.round(i(t,e)/kt)*kt,i:e}}))}return o.links=function(t){return ar(s(t)).edges.filter((function(t){return t.l&&t.r})).map((function(e){return{source:t[e.l.i],target:t[e.r.i]}}))},o.triangles=function(t){var e=[];return ar(s(t)).cells.forEach((function(r,n){for(var i,a,o,s,l=r.site,c=r.edges.sort(qe),u=-1,f=c.length,h=c[f-1].edge,p=h.l===l?h.r:h.l;++u<f;)h,i=p,p=(h=c[u].edge).l===l?h.r:h.l,n<i.i&&n<p.i&&(o=i,s=p,((a=l).x-s.x)*(o.y-a.y)-(a.x-o.x)*(s.y-a.y)<0)&&e.push([t[n],t[i.i],t[p.i]])})),e},o.x=function(t){return arguments.length?(n=ce(e=t),o):e},o.y=function(t){return arguments.length?(i=ce(r=t),o):r},o.clipExtent=function(t){return arguments.length?(a=null==t?sr:t,o):a===sr?null:a},o.size=function(t){return arguments.length?o.clipExtent(t&&[[0,0],t]):a===sr?null:a&&a[1]},o};var sr=[[-1e6,-1e6],[1e6,1e6]];function lr(t){return t.x}function cr(t){return t.y}function ur(t,e,r,n,i,a){if(!t(e,r,n,i,a)){var o=.5*(r+i),s=.5*(n+a),l=e.nodes;l[0]&&ur(t,l[0],r,n,o,s),l[1]&&ur(t,l[1],o,n,i,s),l[2]&&ur(t,l[2],r,s,o,a),l[3]&&ur(t,l[3],o,s,i,a)}}function fr(t,e,r,n,i,a,o){var s,l=1/0;return function t(c,u,f,h,p){if(!(u>a||f>o||h<n||p<i)){if(d=c.point){var d,m=e-c.x,g=r-c.y,v=m*m+g*g;if(v<l){var y=Math.sqrt(l=v);n=e-y,i=r-y,a=e+y,o=r+y,s=d}}for(var x=c.nodes,b=.5*(u+h),_=.5*(f+p),w=(r>=_)<<1|e>=b,T=w+4;w<T;++w)if(c=x[3&w])switch(3&w){case 0:t(c,u,f,b,_);break;case 1:t(c,b,f,h,_);break;case 2:t(c,u,_,b,p);break;case 3:t(c,b,_,h,p)}}}(t,n,i,a,o),s}function hr(e,r){e=t.rgb(e),r=t.rgb(r);var n=e.r,i=e.g,a=e.b,o=r.r-n,s=r.g-i,l=r.b-a;return function(t){return"#"+re(Math.round(n+o*t))+re(Math.round(i+s*t))+re(Math.round(a+l*t))}}function pr(t,e){var r,n={},i={};for(r in t)r in e?n[r]=yr(t[r],e[r]):i[r]=t[r];for(r in e)r in t||(i[r]=e[r]);return function(t){for(r in n)i[r]=n[r](t);return i}}function dr(t,e){return t=+t,e=+e,function(r){return t*(1-r)+e*r}}function mr(t,e){var r,n,i,a=gr.lastIndex=vr.lastIndex=0,o=-1,s=[],l=[];for(t+="",e+="";(r=gr.exec(t))&&(n=vr.exec(e));)(i=n.index)>a&&(i=e.slice(a,i),s[o]?s[o]+=i:s[++o]=i),(r=r[0])===(n=n[0])?s[o]?s[o]+=n:s[++o]=n:(s[++o]=null,l.push({i:o,x:dr(r,n)})),a=vr.lastIndex;return a<e.length&&(i=e.slice(a),s[o]?s[o]+=i:s[++o]=i),s.length<2?l[0]?(e=l[0].x,function(t){return e(t)+""}):function(){return e}:(e=l.length,function(t){for(var r,n=0;n<e;++n)s[(r=l[n]).i]=r.x(t);return s.join("")})}t.geom.delaunay=function(e){return t.geom.voronoi().triangles(e)},t.geom.quadtree=function(t,e,r,n,i){var a,o=_e,s=we;if(a=arguments.length)return o=lr,s=cr,3===a&&(i=r,n=e,r=e=0),l(t);function l(t){var l,c,u,f,h,p,d,m,g,v=ce(o),x=ce(s);if(null!=e)p=e,d=r,m=n,g=i;else if(m=g=-(p=d=1/0),c=[],u=[],h=t.length,a)for(f=0;f<h;++f)(l=t[f]).x<p&&(p=l.x),l.y<d&&(d=l.y),l.x>m&&(m=l.x),l.y>g&&(g=l.y),c.push(l.x),u.push(l.y);else for(f=0;f<h;++f){var b=+v(l=t[f],f),_=+x(l,f);b<p&&(p=b),_<d&&(d=_),b>m&&(m=b),_>g&&(g=_),c.push(b),u.push(_)}var w=m-p,T=g-d;function k(t,e,r,n,i,a,o,s){if(!isNaN(r)&&!isNaN(n))if(t.leaf){var l=t.x,c=t.y;if(null!=l)if(y(l-r)+y(c-n)<.01)A(t,e,r,n,i,a,o,s);else{var u=t.point;t.x=t.y=t.point=null,A(t,u,l,c,i,a,o,s),A(t,e,r,n,i,a,o,s)}else t.x=r,t.y=n,t.point=e}else A(t,e,r,n,i,a,o,s)}function A(t,e,r,n,i,a,o,s){var l=.5*(i+o),c=.5*(a+s),u=r>=l,f=n>=c,h=f<<1|u;t.leaf=!1,u?i=l:o=l,f?a=c:s=c,k(t=t.nodes[h]||(t.nodes[h]={leaf:!0,nodes:[],point:null,x:null,y:null}),e,r,n,i,a,o,s)}w>T?g=d+w:m=p+T;var M={leaf:!0,nodes:[],point:null,x:null,y:null,add:function(t){k(M,t,+v(t,++f),+x(t,f),p,d,m,g)},visit:function(t){ur(t,M,p,d,m,g)},find:function(t){return fr(M,t[0],t[1],p,d,m,g)}};if(f=-1,null==e){for(;++f<h;)k(M,t[f],c[f],u[f],p,d,m,g);--f}else t.forEach(M.add);return c=u=t=l=null,M}return l.x=function(t){return arguments.length?(o=t,l):o},l.y=function(t){return arguments.length?(s=t,l):s},l.extent=function(t){return arguments.length?(null==t?e=r=n=i=null:(e=+t[0][0],r=+t[0][1],n=+t[1][0],i=+t[1][1]),l):null==e?null:[[e,r],[n,i]]},l.size=function(t){return arguments.length?(null==t?e=r=n=i=null:(e=r=0,n=+t[0],i=+t[1]),l):null==e?null:[n-e,i-r]},l},t.interpolateRgb=hr,t.interpolateObject=pr,t.interpolateNumber=dr,t.interpolateString=mr;var gr=/[-+]?(?:\d+\.?\d*|\.?\d+)(?:[eE][-+]?\d+)?/g,vr=new RegExp(gr.source,"g");function yr(e,r){for(var n,i=t.interpolators.length;--i>=0&&!(n=t.interpolators[i](e,r)););return n}function xr(t,e){var r,n=[],i=[],a=t.length,o=e.length,s=Math.min(t.length,e.length);for(r=0;r<s;++r)n.push(yr(t[r],e[r]));for(;r<a;++r)i[r]=t[r];for(;r<o;++r)i[r]=e[r];return function(t){for(r=0;r<s;++r)i[r]=n[r](t);return i}}t.interpolate=yr,t.interpolators=[function(t,e){var r=typeof e;return("string"===r?le.has(e.toLowerCase())||/^(#|rgb\(|hsl\()/i.test(e)?hr:mr:e instanceof Ft?hr:Array.isArray(e)?xr:"object"===r&&isNaN(e)?pr:dr)(t,e)}],t.interpolateArray=xr;var br=function(){return C},_r=t.map({linear:br,poly:function(t){return function(e){return Math.pow(e,t)}},quad:function(){return Mr},cubic:function(){return Sr},sin:function(){return Lr},exp:function(){return Cr},circle:function(){return Pr},elastic:function(t,e){var r;arguments.length<2&&(e=.45);arguments.length?r=e/Mt*Math.asin(1/t):(t=1,r=e/4);return function(n){return 1+t*Math.pow(2,-10*n)*Math.sin((n-r)*Mt/e)}},back:function(t){t||(t=1.70158);return function(e){return e*e*((t+1)*e-t)}},bounce:function(){return Ir}}),wr=t.map({in:C,out:kr,"in-out":Ar,"out-in":function(t){return Ar(kr(t))}});function Tr(t){return function(e){return e<=0?0:e>=1?1:t(e)}}function kr(t){return function(e){return 1-t(1-e)}}function Ar(t){return function(e){return.5*(e<.5?t(2*e):2-t(2-2*e))}}function Mr(t){return t*t}function Sr(t){return t*t*t}function Er(t){if(t<=0)return 0;if(t>=1)return 1;var e=t*t,r=e*t;return 4*(t<.5?r:3*(t-e)+r-.75)}function Lr(t){return 1-Math.cos(t*Et)}function Cr(t){return Math.pow(2,10*(t-1))}function Pr(t){return 1-Math.sqrt(1-t*t)}function Ir(t){return t<1/2.75?7.5625*t*t:t<2/2.75?7.5625*(t-=1.5/2.75)*t+.75:t<2.5/2.75?7.5625*(t-=2.25/2.75)*t+.9375:7.5625*(t-=2.625/2.75)*t+.984375}function Or(t,e){return e-=t,function(r){return Math.round(t+e*r)}}function zr(t){var e,r,n,i=[t.a,t.b],a=[t.c,t.d],o=Rr(i),s=Dr(i,a),l=Rr(((e=a)[0]+=(n=-s)*(r=i)[0],e[1]+=n*r[1],e))||0;i[0]*a[1]<a[0]*i[1]&&(i[0]*=-1,i[1]*=-1,o*=-1,s*=-1),this.rotate=(o?Math.atan2(i[1],i[0]):Math.atan2(-a[0],a[1]))*Ct,this.translate=[t.e,t.f],this.scale=[o,l],this.skew=l?Math.atan2(s,l)*Ct:0}function Dr(t,e){return t[0]*e[0]+t[1]*e[1]}function Rr(t){var e=Math.sqrt(Dr(t,t));return e&&(t[0]/=e,t[1]/=e),e}t.ease=function(t){var e=t.indexOf("-"),n=e>=0?t.slice(0,e):t,i=e>=0?t.slice(e+1):"in";return n=_r.get(n)||br,Tr((i=wr.get(i)||C)(n.apply(null,r.call(arguments,1))))},t.interpolateHcl=function(e,r){e=t.hcl(e),r=t.hcl(r);var n=e.h,i=e.c,a=e.l,o=r.h-n,s=r.c-i,l=r.l-a;isNaN(s)&&(s=0,i=isNaN(i)?r.c:i);isNaN(o)?(o=0,n=isNaN(n)?r.h:n):o>180?o-=360:o<-180&&(o+=360);return function(t){return Ht(n+o*t,i+s*t,a+l*t)+""}},t.interpolateHsl=function(e,r){e=t.hsl(e),r=t.hsl(r);var n=e.h,i=e.s,a=e.l,o=r.h-n,s=r.s-i,l=r.l-a;isNaN(s)&&(s=0,i=isNaN(i)?r.s:i);isNaN(o)?(o=0,n=isNaN(n)?r.h:n):o>180?o-=360:o<-180&&(o+=360);return function(t){return jt(n+o*t,i+s*t,a+l*t)+""}},t.interpolateLab=function(e,r){e=t.lab(e),r=t.lab(r);var n=e.l,i=e.a,a=e.b,o=r.l-n,s=r.a-i,l=r.b-a;return function(t){return Wt(n+o*t,i+s*t,a+l*t)+""}},t.interpolateRound=Or,t.transform=function(e){var r=i.createElementNS(t.ns.prefix.svg,"g");return(t.transform=function(t){if(null!=t){r.setAttribute("transform",t);var e=r.transform.baseVal.consolidate()}return new zr(e?e.matrix:Fr)})(e)},zr.prototype.toString=function(){return"translate("+this.translate+")rotate("+this.rotate+")skewX("+this.skew+")scale("+this.scale+")"};var Fr={a:1,b:0,c:0,d:1,e:0,f:0};function Br(t){return t.length?t.pop()+",":""}function Nr(e,r){var n=[],i=[];return e=t.transform(e),r=t.transform(r),function(t,e,r,n){if(t[0]!==e[0]||t[1]!==e[1]){var i=r.push("translate(",null,",",null,")");n.push({i:i-4,x:dr(t[0],e[0])},{i:i-2,x:dr(t[1],e[1])})}else(e[0]||e[1])&&r.push("translate("+e+")")}(e.translate,r.translate,n,i),function(t,e,r,n){t!==e?(t-e>180?e+=360:e-t>180&&(t+=360),n.push({i:r.push(Br(r)+"rotate(",null,")")-2,x:dr(t,e)})):e&&r.push(Br(r)+"rotate("+e+")")}(e.rotate,r.rotate,n,i),function(t,e,r,n){t!==e?n.push({i:r.push(Br(r)+"skewX(",null,")")-2,x:dr(t,e)}):e&&r.push(Br(r)+"skewX("+e+")")}(e.skew,r.skew,n,i),function(t,e,r,n){if(t[0]!==e[0]||t[1]!==e[1]){var i=r.push(Br(r)+"scale(",null,",",null,")");n.push({i:i-4,x:dr(t[0],e[0])},{i:i-2,x:dr(t[1],e[1])})}else 1===e[0]&&1===e[1]||r.push(Br(r)+"scale("+e+")")}(e.scale,r.scale,n,i),e=r=null,function(t){for(var e,r=-1,a=i.length;++r<a;)n[(e=i[r]).i]=e.x(t);return n.join("")}}function jr(t,e){return e=(e-=t=+t)||1/e,function(r){return(r-t)/e}}function Ur(t,e){return e=(e-=t=+t)||1/e,function(r){return Math.max(0,Math.min(1,(r-t)/e))}}function Vr(t){for(var e=t.source,r=t.target,n=function(t,e){if(t===e)return t;var r=Hr(t),n=Hr(e),i=r.pop(),a=n.pop(),o=null;for(;i===a;)o=i,i=r.pop(),a=n.pop();return o}(e,r),i=[e];e!==n;)e=e.parent,i.push(e);for(var a=i.length;r!==n;)i.splice(a,0,r),r=r.parent;return i}function Hr(t){for(var e=[],r=t.parent;null!=r;)e.push(t),t=r,r=r.parent;return e.push(t),e}function qr(t){t.fixed|=2}function Gr(t){t.fixed&=-7}function Yr(t){t.fixed|=4,t.px=t.x,t.py=t.y}function Wr(t){t.fixed&=-5}t.interpolateTransform=Nr,t.layout={},t.layout.bundle=function(){return function(t){for(var e=[],r=-1,n=t.length;++r<n;)e.push(Vr(t[r]));return e}},t.layout.chord=function(){var e,r,n,i,a,o,s,l={},c=0;function u(){var l,u,h,p,d,m={},g=[],v=t.range(i),y=[];for(e=[],r=[],l=0,p=-1;++p<i;){for(u=0,d=-1;++d<i;)u+=n[p][d];g.push(u),y.push(t.range(i)),l+=u}for(a&&v.sort((function(t,e){return a(g[t],g[e])})),o&&y.forEach((function(t,e){t.sort((function(t,r){return o(n[e][t],n[e][r])}))})),l=(Mt-c*i)/l,u=0,p=-1;++p<i;){for(h=u,d=-1;++d<i;){var x=v[p],b=y[x][d],_=n[x][b],w=u,T=u+=_*l;m[x+"-"+b]={index:x,subindex:b,startAngle:w,endAngle:T,value:_}}r[x]={index:x,startAngle:h,endAngle:u,value:g[x]},u+=c}for(p=-1;++p<i;)for(d=p-1;++d<i;){var k=m[p+"-"+d],A=m[d+"-"+p];(k.value||A.value)&&e.push(k.value<A.value?{source:A,target:k}:{source:k,target:A})}s&&f()}function f(){e.sort((function(t,e){return s((t.source.value+t.target.value)/2,(e.source.value+e.target.value)/2)}))}return l.matrix=function(t){return arguments.length?(i=(n=t)&&n.length,e=r=null,l):n},l.padding=function(t){return arguments.length?(c=t,e=r=null,l):c},l.sortGroups=function(t){return arguments.length?(a=t,e=r=null,l):a},l.sortSubgroups=function(t){return arguments.length?(o=t,e=null,l):o},l.sortChords=function(t){return arguments.length?(s=t,e&&f(),l):s},l.chords=function(){return e||u(),e},l.groups=function(){return r||u(),r},l},t.layout.force=function(){var e,r,n,i,a,o,s={},l=t.dispatch("start","tick","end"),c=[1,1],u=.9,f=Xr,h=Zr,p=-30,d=Jr,m=.1,g=.64,v=[],y=[];function x(t){return function(e,r,n,i){if(e.point!==t){var a=e.cx-t.x,o=e.cy-t.y,s=i-r,l=a*a+o*o;if(s*s/g<l){if(l<d){var c=e.charge/l;t.px-=a*c,t.py-=o*c}return!0}if(e.point&&l&&l<d){c=e.pointCharge/l;t.px-=a*c,t.py-=o*c}}return!e.charge}}function b(e){e.px=t.event.x,e.py=t.event.y,s.resume()}return s.tick=function(){if((n*=.99)<.005)return e=null,l.end({type:"end",alpha:n=0}),!0;var r,s,f,h,d,g,b,_,w,T=v.length,k=y.length;for(s=0;s<k;++s)h=(f=y[s]).source,(g=(_=(d=f.target).x-h.x)*_+(w=d.y-h.y)*w)&&(_*=g=n*a[s]*((g=Math.sqrt(g))-i[s])/g,w*=g,d.x-=_*(b=h.weight+d.weight?h.weight/(h.weight+d.weight):.5),d.y-=w*b,h.x+=_*(b=1-b),h.y+=w*b);if((b=n*m)&&(_=c[0]/2,w=c[1]/2,s=-1,b))for(;++s<T;)(f=v[s]).x+=(_-f.x)*b,f.y+=(w-f.y)*b;if(p)for(!function t(e,r,n){var i=0,a=0;if(e.charge=0,!e.leaf)for(var o,s=e.nodes,l=s.length,c=-1;++c<l;)null!=(o=s[c])&&(t(o,r,n),e.charge+=o.charge,i+=o.charge*o.cx,a+=o.charge*o.cy);if(e.point){e.leaf||(e.point.x+=Math.random()-.5,e.point.y+=Math.random()-.5);var u=r*n[e.point.index];e.charge+=e.pointCharge=u,i+=u*e.point.x,a+=u*e.point.y}e.cx=i/e.charge,e.cy=a/e.charge}(r=t.geom.quadtree(v),n,o),s=-1;++s<T;)(f=v[s]).fixed||r.visit(x(f));for(s=-1;++s<T;)(f=v[s]).fixed?(f.x=f.px,f.y=f.py):(f.x-=(f.px-(f.px=f.x))*u,f.y-=(f.py-(f.py=f.y))*u);l.tick({type:"tick",alpha:n})},s.nodes=function(t){return arguments.length?(v=t,s):v},s.links=function(t){return arguments.length?(y=t,s):y},s.size=function(t){return arguments.length?(c=t,s):c},s.linkDistance=function(t){return arguments.length?(f="function"==typeof t?t:+t,s):f},s.distance=s.linkDistance,s.linkStrength=function(t){return arguments.length?(h="function"==typeof t?t:+t,s):h},s.friction=function(t){return arguments.length?(u=+t,s):u},s.charge=function(t){return arguments.length?(p="function"==typeof t?t:+t,s):p},s.chargeDistance=function(t){return arguments.length?(d=t*t,s):Math.sqrt(d)},s.gravity=function(t){return arguments.length?(m=+t,s):m},s.theta=function(t){return arguments.length?(g=t*t,s):Math.sqrt(g)},s.alpha=function(t){return arguments.length?(t=+t,n?t>0?n=t:(e.c=null,e.t=NaN,e=null,l.end({type:"end",alpha:n=0})):t>0&&(l.start({type:"start",alpha:n=t}),e=ve(s.tick)),s):n},s.start=function(){var t,e,r,n=v.length,l=y.length,u=c[0],d=c[1];for(t=0;t<n;++t)(r=v[t]).index=t,r.weight=0;for(t=0;t<l;++t)"number"==typeof(r=y[t]).source&&(r.source=v[r.source]),"number"==typeof r.target&&(r.target=v[r.target]),++r.source.weight,++r.target.weight;for(t=0;t<n;++t)r=v[t],isNaN(r.x)&&(r.x=m("x",u)),isNaN(r.y)&&(r.y=m("y",d)),isNaN(r.px)&&(r.px=r.x),isNaN(r.py)&&(r.py=r.y);if(i=[],"function"==typeof f)for(t=0;t<l;++t)i[t]=+f.call(this,y[t],t);else for(t=0;t<l;++t)i[t]=f;if(a=[],"function"==typeof h)for(t=0;t<l;++t)a[t]=+h.call(this,y[t],t);else for(t=0;t<l;++t)a[t]=h;if(o=[],"function"==typeof p)for(t=0;t<n;++t)o[t]=+p.call(this,v[t],t);else for(t=0;t<n;++t)o[t]=p;function m(r,i){if(!e){for(e=new Array(n),c=0;c<n;++c)e[c]=[];for(c=0;c<l;++c){var a=y[c];e[a.source.index].push(a.target),e[a.target.index].push(a.source)}}for(var o,s=e[t],c=-1,u=s.length;++c<u;)if(!isNaN(o=s[c][r]))return o;return Math.random()*i}return s.resume()},s.resume=function(){return s.alpha(.1)},s.stop=function(){return s.alpha(0)},s.drag=function(){if(r||(r=t.behavior.drag().origin(C).on("dragstart.force",qr).on("drag.force",b).on("dragend.force",Gr)),!arguments.length)return r;this.on("mouseover.force",Yr).on("mouseout.force",Wr).call(r)},t.rebind(s,l,"on")};var Xr=20,Zr=1,Jr=1/0;function Kr(e,r){return t.rebind(e,r,"sort","children","value"),e.nodes=e,e.links=nn,e}function Qr(t,e){for(var r=[t];null!=(t=r.pop());)if(e(t),(i=t.children)&&(n=i.length))for(var n,i;--n>=0;)r.push(i[n])}function $r(t,e){for(var r=[t],n=[];null!=(t=r.pop());)if(n.push(t),(a=t.children)&&(i=a.length))for(var i,a,o=-1;++o<i;)r.push(a[o]);for(;null!=(t=n.pop());)e(t)}function tn(t){return t.children}function en(t){return t.value}function rn(t,e){return e.value-t.value}function nn(e){return t.merge(e.map((function(t){return(t.children||[]).map((function(e){return{source:t,target:e}}))})))}t.layout.hierarchy=function(){var t=rn,e=tn,r=en;function n(i){var a,o=[i],s=[];for(i.depth=0;null!=(a=o.pop());)if(s.push(a),(c=e.call(n,a,a.depth))&&(l=c.length)){for(var l,c,u;--l>=0;)o.push(u=c[l]),u.parent=a,u.depth=a.depth+1;r&&(a.value=0),a.children=c}else r&&(a.value=+r.call(n,a,a.depth)||0),delete a.children;return $r(i,(function(e){var n,i;t&&(n=e.children)&&n.sort(t),r&&(i=e.parent)&&(i.value+=e.value)})),s}return n.sort=function(e){return arguments.length?(t=e,n):t},n.children=function(t){return arguments.length?(e=t,n):e},n.value=function(t){return arguments.length?(r=t,n):r},n.revalue=function(t){return r&&(Qr(t,(function(t){t.children&&(t.value=0)})),$r(t,(function(t){var e;t.children||(t.value=+r.call(n,t,t.depth)||0),(e=t.parent)&&(e.value+=t.value)}))),t},n},t.layout.partition=function(){var e=t.layout.hierarchy(),r=[1,1];function n(t,n){var i=e.call(this,t,n);return function t(e,r,n,i){var a=e.children;if(e.x=r,e.y=e.depth*i,e.dx=n,e.dy=i,a&&(o=a.length)){var o,s,l,c=-1;for(n=e.value?n/e.value:0;++c<o;)t(s=a[c],r,l=s.value*n,i),r+=l}}(i[0],0,r[0],r[1]/function t(e){var r=e.children,n=0;if(r&&(i=r.length))for(var i,a=-1;++a<i;)n=Math.max(n,t(r[a]));return 1+n}(i[0])),i}return n.size=function(t){return arguments.length?(r=t,n):r},Kr(n,e)},t.layout.pie=function(){var e=Number,r=an,n=0,i=Mt,a=0;function o(s){var l,c=s.length,u=s.map((function(t,r){return+e.call(o,t,r)})),f=+("function"==typeof n?n.apply(this,arguments):n),h=("function"==typeof i?i.apply(this,arguments):i)-f,p=Math.min(Math.abs(h)/c,+("function"==typeof a?a.apply(this,arguments):a)),d=p*(h<0?-1:1),m=t.sum(u),g=m?(h-c*d)/m:0,v=t.range(c),y=[];return null!=r&&v.sort(r===an?function(t,e){return u[e]-u[t]}:function(t,e){return r(s[t],s[e])}),v.forEach((function(t){y[t]={data:s[t],value:l=u[t],startAngle:f,endAngle:f+=l*g+d,padAngle:p}})),y}return o.value=function(t){return arguments.length?(e=t,o):e},o.sort=function(t){return arguments.length?(r=t,o):r},o.startAngle=function(t){return arguments.length?(n=t,o):n},o.endAngle=function(t){return arguments.length?(i=t,o):i},o.padAngle=function(t){return arguments.length?(a=t,o):a},o};var an={};function on(t){return t.x}function sn(t){return t.y}function ln(t,e,r){t.y0=e,t.y=r}t.layout.stack=function(){var e=C,r=fn,n=hn,i=ln,a=on,o=sn;function s(l,c){if(!(p=l.length))return l;var u=l.map((function(t,r){return e.call(s,t,r)})),f=u.map((function(t){return t.map((function(t,e){return[a.call(s,t,e),o.call(s,t,e)]}))})),h=r.call(s,f,c);u=t.permute(u,h),f=t.permute(f,h);var p,d,m,g,v=n.call(s,f,c),y=u[0].length;for(m=0;m<y;++m)for(i.call(s,u[0][m],g=v[m],f[0][m][1]),d=1;d<p;++d)i.call(s,u[d][m],g+=f[d-1][m][1],f[d][m][1]);return l}return s.values=function(t){return arguments.length?(e=t,s):e},s.order=function(t){return arguments.length?(r="function"==typeof t?t:cn.get(t)||fn,s):r},s.offset=function(t){return arguments.length?(n="function"==typeof t?t:un.get(t)||hn,s):n},s.x=function(t){return arguments.length?(a=t,s):a},s.y=function(t){return arguments.length?(o=t,s):o},s.out=function(t){return arguments.length?(i=t,s):i},s};var cn=t.map({"inside-out":function(e){var r,n,i=e.length,a=e.map(pn),o=e.map(dn),s=t.range(i).sort((function(t,e){return a[t]-a[e]})),l=0,c=0,u=[],f=[];for(r=0;r<i;++r)n=s[r],l<c?(l+=o[n],u.push(n)):(c+=o[n],f.push(n));return f.reverse().concat(u)},reverse:function(e){return t.range(e.length).reverse()},default:fn}),un=t.map({silhouette:function(t){var e,r,n,i=t.length,a=t[0].length,o=[],s=0,l=[];for(r=0;r<a;++r){for(e=0,n=0;e<i;e++)n+=t[e][r][1];n>s&&(s=n),o.push(n)}for(r=0;r<a;++r)l[r]=(s-o[r])/2;return l},wiggle:function(t){var e,r,n,i,a,o,s,l,c,u=t.length,f=t[0],h=f.length,p=[];for(p[0]=l=c=0,r=1;r<h;++r){for(e=0,i=0;e<u;++e)i+=t[e][r][1];for(e=0,a=0,s=f[r][0]-f[r-1][0];e<u;++e){for(n=0,o=(t[e][r][1]-t[e][r-1][1])/(2*s);n<e;++n)o+=(t[n][r][1]-t[n][r-1][1])/s;a+=o*t[e][r][1]}p[r]=l-=i?a/i*s:0,l<c&&(c=l)}for(r=0;r<h;++r)p[r]-=c;return p},expand:function(t){var e,r,n,i=t.length,a=t[0].length,o=1/i,s=[];for(r=0;r<a;++r){for(e=0,n=0;e<i;e++)n+=t[e][r][1];if(n)for(e=0;e<i;e++)t[e][r][1]/=n;else for(e=0;e<i;e++)t[e][r][1]=o}for(r=0;r<a;++r)s[r]=0;return s},zero:hn});function fn(e){return t.range(e.length)}function hn(t){for(var e=-1,r=t[0].length,n=[];++e<r;)n[e]=0;return n}function pn(t){for(var e,r=1,n=0,i=t[0][1],a=t.length;r<a;++r)(e=t[r][1])>i&&(n=r,i=e);return n}function dn(t){return t.reduce(mn,0)}function mn(t,e){return t+e[1]}function gn(t,e){return vn(t,Math.ceil(Math.log(e.length)/Math.LN2+1))}function vn(t,e){for(var r=-1,n=+t[0],i=(t[1]-n)/e,a=[];++r<=e;)a[r]=i*r+n;return a}function yn(e){return[t.min(e),t.max(e)]}function xn(t,e){return t.value-e.value}function bn(t,e){var r=t._pack_next;t._pack_next=e,e._pack_prev=t,e._pack_next=r,r._pack_prev=e}function _n(t,e){t._pack_next=e,e._pack_prev=t}function wn(t,e){var r=e.x-t.x,n=e.y-t.y,i=t.r+e.r;return.999*i*i>r*r+n*n}function Tn(t){if((e=t.children)&&(l=e.length)){var e,r,n,i,a,o,s,l,c=1/0,u=-1/0,f=1/0,h=-1/0;if(e.forEach(kn),(r=e[0]).x=-r.r,r.y=0,x(r),l>1&&((n=e[1]).x=n.r,n.y=0,x(n),l>2))for(Mn(r,n,i=e[2]),x(i),bn(r,i),r._pack_prev=i,bn(i,n),n=r._pack_next,a=3;a<l;a++){Mn(r,n,i=e[a]);var p=0,d=1,m=1;for(o=n._pack_next;o!==n;o=o._pack_next,d++)if(wn(o,i)){p=1;break}if(1==p)for(s=r._pack_prev;s!==o._pack_prev&&!wn(s,i);s=s._pack_prev,m++);p?(d<m||d==m&&n.r<r.r?_n(r,n=o):_n(r=s,n),a--):(bn(r,i),n=i,x(i))}var g=(c+u)/2,v=(f+h)/2,y=0;for(a=0;a<l;a++)(i=e[a]).x-=g,i.y-=v,y=Math.max(y,i.r+Math.sqrt(i.x*i.x+i.y*i.y));t.r=y,e.forEach(An)}function x(t){c=Math.min(t.x-t.r,c),u=Math.max(t.x+t.r,u),f=Math.min(t.y-t.r,f),h=Math.max(t.y+t.r,h)}}function kn(t){t._pack_next=t._pack_prev=t}function An(t){delete t._pack_next,delete t._pack_prev}function Mn(t,e,r){var n=t.r+r.r,i=e.x-t.x,a=e.y-t.y;if(n&&(i||a)){var o=e.r+r.r,s=i*i+a*a,l=.5+((n*=n)-(o*=o))/(2*s),c=Math.sqrt(Math.max(0,2*o*(n+s)-(n-=s)*n-o*o))/(2*s);r.x=t.x+l*i+c*a,r.y=t.y+l*a-c*i}else r.x=t.x+n,r.y=t.y}function Sn(t,e){return t.parent==e.parent?1:2}function En(t){var e=t.children;return e.length?e[0]:t.t}function Ln(t){var e,r=t.children;return(e=r.length)?r[e-1]:t.t}function Cn(t,e,r){var n=r/(e.i-t.i);e.c-=n,e.s+=r,t.c+=n,e.z+=r,e.m+=r}function Pn(t,e,r){return t.a.parent===e.parent?t.a:r}function In(t){return{x:t.x,y:t.y,dx:t.dx,dy:t.dy}}function On(t,e){var r=t.x+e[3],n=t.y+e[0],i=t.dx-e[1]-e[3],a=t.dy-e[0]-e[2];return i<0&&(r+=i/2,i=0),a<0&&(n+=a/2,a=0),{x:r,y:n,dx:i,dy:a}}function zn(t){var e=t[0],r=t[t.length-1];return e<r?[e,r]:[r,e]}function Dn(t){return t.rangeExtent?t.rangeExtent():zn(t.range())}function Rn(t,e,r,n){var i=r(t[0],t[1]),a=n(e[0],e[1]);return function(t){return a(i(t))}}function Fn(t,e){var r,n=0,i=t.length-1,a=t[n],o=t[i];return o<a&&(r=n,n=i,i=r,r=a,a=o,o=r),t[n]=e.floor(a),t[i]=e.ceil(o),t}function Bn(t){return t?{floor:function(e){return Math.floor(e/t)*t},ceil:function(e){return Math.ceil(e/t)*t}}:Nn}t.layout.histogram=function(){var e=!0,r=Number,n=yn,i=gn;function a(a,o){for(var s,l,c=[],u=a.map(r,this),f=n.call(this,u,o),h=i.call(this,f,u,o),p=(o=-1,u.length),d=h.length-1,m=e?1:1/p;++o<d;)(s=c[o]=[]).dx=h[o+1]-(s.x=h[o]),s.y=0;if(d>0)for(o=-1;++o<p;)(l=u[o])>=f[0]&&l<=f[1]&&((s=c[t.bisect(h,l,1,d)-1]).y+=m,s.push(a[o]));return c}return a.value=function(t){return arguments.length?(r=t,a):r},a.range=function(t){return arguments.length?(n=ce(t),a):n},a.bins=function(t){return arguments.length?(i="number"==typeof t?function(e){return vn(e,t)}:ce(t),a):i},a.frequency=function(t){return arguments.length?(e=!!t,a):e},a},t.layout.pack=function(){var e,r=t.layout.hierarchy().sort(xn),n=0,i=[1,1];function a(t,a){var o=r.call(this,t,a),s=o[0],l=i[0],c=i[1],u=null==e?Math.sqrt:"function"==typeof e?e:function(){return e};if(s.x=s.y=0,$r(s,(function(t){t.r=+u(t.value)})),$r(s,Tn),n){var f=n*(e?1:Math.max(2*s.r/l,2*s.r/c))/2;$r(s,(function(t){t.r+=f})),$r(s,Tn),$r(s,(function(t){t.r-=f}))}return function t(e,r,n,i){var a=e.children;if(e.x=r+=i*e.x,e.y=n+=i*e.y,e.r*=i,a)for(var o=-1,s=a.length;++o<s;)t(a[o],r,n,i)}(s,l/2,c/2,e?1:1/Math.max(2*s.r/l,2*s.r/c)),o}return a.size=function(t){return arguments.length?(i=t,a):i},a.radius=function(t){return arguments.length?(e=null==t||"function"==typeof t?t:+t,a):e},a.padding=function(t){return arguments.length?(n=+t,a):n},Kr(a,r)},t.layout.tree=function(){var e=t.layout.hierarchy().sort(null).value(null),r=Sn,n=[1,1],i=null;function a(t,a){var c=e.call(this,t,a),u=c[0],f=function(t){var e,r={A:null,children:[t]},n=[r];for(;null!=(e=n.pop());)for(var i,a=e.children,o=0,s=a.length;o<s;++o)n.push((a[o]=i={_:a[o],parent:e,children:(i=a[o].children)&&i.slice()||[],A:null,a:null,z:0,m:0,c:0,s:0,t:null,i:o}).a=i);return r.children[0]}(u);if($r(f,o),f.parent.m=-f.z,Qr(f,s),i)Qr(u,l);else{var h=u,p=u,d=u;Qr(u,(function(t){t.x<h.x&&(h=t),t.x>p.x&&(p=t),t.depth>d.depth&&(d=t)}));var m=r(h,p)/2-h.x,g=n[0]/(p.x+r(p,h)/2+m),v=n[1]/(d.depth||1);Qr(u,(function(t){t.x=(t.x+m)*g,t.y=t.depth*v}))}return c}function o(t){var e=t.children,n=t.parent.children,i=t.i?n[t.i-1]:null;if(e.length){!function(t){var e,r=0,n=0,i=t.children,a=i.length;for(;--a>=0;)(e=i[a]).z+=r,e.m+=r,r+=e.s+(n+=e.c)}(t);var a=(e[0].z+e[e.length-1].z)/2;i?(t.z=i.z+r(t._,i._),t.m=t.z-a):t.z=a}else i&&(t.z=i.z+r(t._,i._));t.parent.A=function(t,e,n){if(e){for(var i,a=t,o=t,s=e,l=a.parent.children[0],c=a.m,u=o.m,f=s.m,h=l.m;s=Ln(s),a=En(a),s&&a;)l=En(l),(o=Ln(o)).a=t,(i=s.z+f-a.z-c+r(s._,a._))>0&&(Cn(Pn(s,t,n),t,i),c+=i,u+=i),f+=s.m,c+=a.m,h+=l.m,u+=o.m;s&&!Ln(o)&&(o.t=s,o.m+=f-u),a&&!En(l)&&(l.t=a,l.m+=c-h,n=t)}return n}(t,i,t.parent.A||n[0])}function s(t){t._.x=t.z+t.parent.m,t.m+=t.parent.m}function l(t){t.x*=n[0],t.y=t.depth*n[1]}return a.separation=function(t){return arguments.length?(r=t,a):r},a.size=function(t){return arguments.length?(i=null==(n=t)?l:null,a):i?null:n},a.nodeSize=function(t){return arguments.length?(i=null==(n=t)?null:l,a):i?n:null},Kr(a,e)},t.layout.cluster=function(){var e=t.layout.hierarchy().sort(null).value(null),r=Sn,n=[1,1],i=!1;function a(a,o){var s,l=e.call(this,a,o),c=l[0],u=0;$r(c,(function(e){var n=e.children;n&&n.length?(e.x=function(t){return t.reduce((function(t,e){return t+e.x}),0)/t.length}(n),e.y=function(e){return 1+t.max(e,(function(t){return t.y}))}(n)):(e.x=s?u+=r(e,s):0,e.y=0,s=e)}));var f=function t(e){var r=e.children;return r&&r.length?t(r[0]):e}(c),h=function t(e){var r,n=e.children;return n&&(r=n.length)?t(n[r-1]):e}(c),p=f.x-r(f,h)/2,d=h.x+r(h,f)/2;return $r(c,i?function(t){t.x=(t.x-c.x)*n[0],t.y=(c.y-t.y)*n[1]}:function(t){t.x=(t.x-p)/(d-p)*n[0],t.y=(1-(c.y?t.y/c.y:1))*n[1]}),l}return a.separation=function(t){return arguments.length?(r=t,a):r},a.size=function(t){return arguments.length?(i=null==(n=t),a):i?null:n},a.nodeSize=function(t){return arguments.length?(i=null!=(n=t),a):i?n:null},Kr(a,e)},t.layout.treemap=function(){var e,r=t.layout.hierarchy(),n=Math.round,i=[1,1],a=null,o=In,s=!1,l="squarify",c=.5*(1+Math.sqrt(5));function u(t,e){for(var r,n,i=-1,a=t.length;++i<a;)n=(r=t[i]).value*(e<0?0:e),r.area=isNaN(n)||n<=0?0:n}function f(t){var e=t.children;if(e&&e.length){var r,n,i,a=o(t),s=[],c=e.slice(),h=1/0,m="slice"===l?a.dx:"dice"===l?a.dy:"slice-dice"===l?1&t.depth?a.dy:a.dx:Math.min(a.dx,a.dy);for(u(c,a.dx*a.dy/t.value),s.area=0;(i=c.length)>0;)s.push(r=c[i-1]),s.area+=r.area,"squarify"!==l||(n=p(s,m))<=h?(c.pop(),h=n):(s.area-=s.pop().area,d(s,m,a,!1),m=Math.min(a.dx,a.dy),s.length=s.area=0,h=1/0);s.length&&(d(s,m,a,!0),s.length=s.area=0),e.forEach(f)}}function h(t){var e=t.children;if(e&&e.length){var r,n=o(t),i=e.slice(),a=[];for(u(i,n.dx*n.dy/t.value),a.area=0;r=i.pop();)a.push(r),a.area+=r.area,null!=r.z&&(d(a,r.z?n.dx:n.dy,n,!i.length),a.length=a.area=0);e.forEach(h)}}function p(t,e){for(var r,n=t.area,i=0,a=1/0,o=-1,s=t.length;++o<s;)(r=t[o].area)&&(r<a&&(a=r),r>i&&(i=r));return e*=e,(n*=n)?Math.max(e*i*c/n,n/(e*a*c)):1/0}function d(t,e,r,i){var a,o=-1,s=t.length,l=r.x,c=r.y,u=e?n(t.area/e):0;if(e==r.dx){for((i||u>r.dy)&&(u=r.dy);++o<s;)(a=t[o]).x=l,a.y=c,a.dy=u,l+=a.dx=Math.min(r.x+r.dx-l,u?n(a.area/u):0);a.z=!0,a.dx+=r.x+r.dx-l,r.y+=u,r.dy-=u}else{for((i||u>r.dx)&&(u=r.dx);++o<s;)(a=t[o]).x=l,a.y=c,a.dx=u,c+=a.dy=Math.min(r.y+r.dy-c,u?n(a.area/u):0);a.z=!1,a.dy+=r.y+r.dy-c,r.x+=u,r.dx-=u}}function m(t){var n=e||r(t),a=n[0];return a.x=a.y=0,a.value?(a.dx=i[0],a.dy=i[1]):a.dx=a.dy=0,e&&r.revalue(a),u([a],a.dx*a.dy/a.value),(e?h:f)(a),s&&(e=n),n}return m.size=function(t){return arguments.length?(i=t,m):i},m.padding=function(t){if(!arguments.length)return a;function e(e){var r=t.call(m,e,e.depth);return null==r?In(e):On(e,"number"==typeof r?[r,r,r,r]:r)}function r(e){return On(e,t)}var n;return o=null==(a=t)?In:"function"==(n=typeof t)?e:"number"===n?(t=[t,t,t,t],r):r,m},m.round=function(t){return arguments.length?(n=t?Math.round:Number,m):n!=Number},m.sticky=function(t){return arguments.length?(s=t,e=null,m):s},m.ratio=function(t){return arguments.length?(c=t,m):c},m.mode=function(t){return arguments.length?(l=t+"",m):l},Kr(m,r)},t.random={normal:function(t,e){var r=arguments.length;return r<2&&(e=1),r<1&&(t=0),function(){var r,n,i;do{i=(r=2*Math.random()-1)*r+(n=2*Math.random()-1)*n}while(!i||i>1);return t+e*r*Math.sqrt(-2*Math.log(i)/i)}},logNormal:function(){var e=t.random.normal.apply(t,arguments);return function(){return Math.exp(e())}},bates:function(e){var r=t.random.irwinHall(e);return function(){return r()/e}},irwinHall:function(t){return function(){for(var e=0,r=0;r<t;r++)e+=Math.random();return e}}},t.scale={};var Nn={floor:C,ceil:C};function jn(e,r,n,i){var a=[],o=[],s=0,l=Math.min(e.length,r.length)-1;for(e[l]<e[0]&&(e=e.slice().reverse(),r=r.slice().reverse());++s<=l;)a.push(n(e[s-1],e[s])),o.push(i(r[s-1],r[s]));return function(r){var n=t.bisect(e,r,1,l)-1;return o[n](a[n](r))}}function Un(e,r){return t.rebind(e,r,"range","rangeRound","interpolate","clamp")}function Vn(t,e){return Fn(t,Bn(Hn(t,e)[2])),Fn(t,Bn(Hn(t,e)[2])),t}function Hn(t,e){null==e&&(e=10);var r=zn(t),n=r[1]-r[0],i=Math.pow(10,Math.floor(Math.log(n/e)/Math.LN10)),a=e/n*i;return a<=.15?i*=10:a<=.35?i*=5:a<=.75&&(i*=2),r[0]=Math.ceil(r[0]/i)*i,r[1]=Math.floor(r[1]/i)*i+.5*i,r[2]=i,r}function qn(e,r){return t.range.apply(t,Hn(e,r))}t.scale.linear=function(){return function t(e,r,n,i){var a,o;function s(){var t=Math.min(e.length,r.length)>2?jn:Rn,s=i?Ur:jr;return a=t(e,r,s,n),o=t(r,e,s,yr),l}function l(t){return a(t)}return l.invert=function(t){return o(t)},l.domain=function(t){return arguments.length?(e=t.map(Number),s()):e},l.range=function(t){return arguments.length?(r=t,s()):r},l.rangeRound=function(t){return l.range(t).interpolate(Or)},l.clamp=function(t){return arguments.length?(i=t,s()):i},l.interpolate=function(t){return arguments.length?(n=t,s()):n},l.ticks=function(t){return qn(e,t)},l.tickFormat=function(t,r){return d3_scale_linearTickFormat(e,t,r)},l.nice=function(t){return Vn(e,t),s()},l.copy=function(){return t(e,r,n,i)},s()}([0,1],[0,1],yr,!1)};t.scale.log=function(){return function t(e,r,n,i){function a(t){return(n?Math.log(t<0?0:t):-Math.log(t>0?0:-t))/Math.log(r)}function o(t){return n?Math.pow(r,t):-Math.pow(r,-t)}function s(t){return e(a(t))}return s.invert=function(t){return o(e.invert(t))},s.domain=function(t){return arguments.length?(n=t[0]>=0,e.domain((i=t.map(Number)).map(a)),s):i},s.base=function(t){return arguments.length?(r=+t,e.domain(i.map(a)),s):r},s.nice=function(){var t=Fn(i.map(a),n?Math:Gn);return e.domain(t),i=t.map(o),s},s.ticks=function(){var t=zn(i),e=[],s=t[0],l=t[1],c=Math.floor(a(s)),u=Math.ceil(a(l)),f=r%1?2:r;if(isFinite(u-c)){if(n){for(;c<u;c++)for(var h=1;h<f;h++)e.push(o(c)*h);e.push(o(c))}else for(e.push(o(c));c++<u;)for(h=f-1;h>0;h--)e.push(o(c)*h);for(c=0;e[c]<s;c++);for(u=e.length;e[u-1]>l;u--);e=e.slice(c,u)}return e},s.copy=function(){return t(e.copy(),r,n,i)},Un(s,e)}(t.scale.linear().domain([0,1]),10,!0,[1,10])};var Gn={floor:function(t){return-Math.ceil(-t)},ceil:function(t){return-Math.floor(-t)}};function Yn(t){return function(e){return e<0?-Math.pow(-e,t):Math.pow(e,t)}}t.scale.pow=function(){return function t(e,r,n){var i=Yn(r),a=Yn(1/r);function o(t){return e(i(t))}return o.invert=function(t){return a(e.invert(t))},o.domain=function(t){return arguments.length?(e.domain((n=t.map(Number)).map(i)),o):n},o.ticks=function(t){return qn(n,t)},o.tickFormat=function(t,e){return d3_scale_linearTickFormat(n,t,e)},o.nice=function(t){return o.domain(Vn(n,t))},o.exponent=function(t){return arguments.length?(i=Yn(r=t),a=Yn(1/r),e.domain(n.map(i)),o):r},o.copy=function(){return t(e.copy(),r,n)},Un(o,e)}(t.scale.linear(),1,[0,1])},t.scale.sqrt=function(){return t.scale.pow().exponent(.5)},t.scale.ordinal=function(){return function e(r,n){var i,a,o;function s(t){return a[((i.get(t)||("range"===n.t?i.set(t,r.push(t)):NaN))-1)%a.length]}function l(e,n){return t.range(r.length).map((function(t){return e+n*t}))}return s.domain=function(t){if(!arguments.length)return r;r=[],i=new _;for(var e,a=-1,o=t.length;++a<o;)i.has(e=t[a])||i.set(e,r.push(e));return s[n.t].apply(s,n.a)},s.range=function(t){return arguments.length?(a=t,o=0,n={t:"range",a:arguments},s):a},s.rangePoints=function(t,e){arguments.length<2&&(e=0);var i=t[0],c=t[1],u=r.length<2?(i=(i+c)/2,0):(c-i)/(r.length-1+e);return a=l(i+u*e/2,u),o=0,n={t:"rangePoints",a:arguments},s},s.rangeRoundPoints=function(t,e){arguments.length<2&&(e=0);var i=t[0],c=t[1],u=r.length<2?(i=c=Math.round((i+c)/2),0):(c-i)/(r.length-1+e)|0;return a=l(i+Math.round(u*e/2+(c-i-(r.length-1+e)*u)/2),u),o=0,n={t:"rangeRoundPoints",a:arguments},s},s.rangeBands=function(t,e,i){arguments.length<2&&(e=0),arguments.length<3&&(i=e);var c=t[1]<t[0],u=t[c-0],f=t[1-c],h=(f-u)/(r.length-e+2*i);return a=l(u+h*i,h),c&&a.reverse(),o=h*(1-e),n={t:"rangeBands",a:arguments},s},s.rangeRoundBands=function(t,e,i){arguments.length<2&&(e=0),arguments.length<3&&(i=e);var c=t[1]<t[0],u=t[c-0],f=t[1-c],h=Math.floor((f-u)/(r.length-e+2*i));return a=l(u+Math.round((f-u-(r.length-e)*h)/2),h),c&&a.reverse(),o=Math.round(h*(1-e)),n={t:"rangeRoundBands",a:arguments},s},s.rangeBand=function(){return o},s.rangeExtent=function(){return zn(n.a[0])},s.copy=function(){return e(r,n)},s.domain(r)}([],{t:"range",a:[[]]})},t.scale.category10=function(){return t.scale.ordinal().range(Wn)},t.scale.category20=function(){return t.scale.ordinal().range(Xn)},t.scale.category20b=function(){return t.scale.ordinal().range(Zn)},t.scale.category20c=function(){return t.scale.ordinal().range(Jn)};var Wn=[2062260,16744206,2924588,14034728,9725885,9197131,14907330,8355711,12369186,1556175].map(te),Xn=[2062260,11454440,16744206,16759672,2924588,10018698,14034728,16750742,9725885,12955861,9197131,12885140,14907330,16234194,8355711,13092807,12369186,14408589,1556175,10410725].map(te),Zn=[3750777,5395619,7040719,10264286,6519097,9216594,11915115,13556636,9202993,12426809,15186514,15190932,8666169,11356490,14049643,15177372,8077683,10834324,13528509,14589654].map(te),Jn=[3244733,7057110,10406625,13032431,15095053,16616764,16625259,16634018,3253076,7652470,10607003,13101504,7695281,10394312,12369372,14342891,6513507,9868950,12434877,14277081].map(te);function Kn(){return 0}t.scale.quantile=function(){return function e(r,n){var i;function a(){var e=0,a=n.length;for(i=[];++e<a;)i[e-1]=t.quantile(r,e/a);return o}function o(e){if(!isNaN(e=+e))return n[t.bisect(i,e)]}return o.domain=function(t){return arguments.length?(r=t.map(p).filter(d).sort(h),a()):r},o.range=function(t){return arguments.length?(n=t,a()):n},o.quantiles=function(){return i},o.invertExtent=function(t){return(t=n.indexOf(t))<0?[NaN,NaN]:[t>0?i[t-1]:r[0],t<i.length?i[t]:r[r.length-1]]},o.copy=function(){return e(r,n)},a()}([],[])},t.scale.quantize=function(){return function t(e,r,n){var i,a;function o(t){return n[Math.max(0,Math.min(a,Math.floor(i*(t-e))))]}function s(){return i=n.length/(r-e),a=n.length-1,o}return o.domain=function(t){return arguments.length?(e=+t[0],r=+t[t.length-1],s()):[e,r]},o.range=function(t){return arguments.length?(n=t,s()):n},o.invertExtent=function(t){return[t=(t=n.indexOf(t))<0?NaN:t/i+e,t+1/i]},o.copy=function(){return t(e,r,n)},s()}(0,1,[0,1])},t.scale.threshold=function(){return function e(r,n){function i(e){if(e<=e)return n[t.bisect(r,e)]}return i.domain=function(t){return arguments.length?(r=t,i):r},i.range=function(t){return arguments.length?(n=t,i):n},i.invertExtent=function(t){return t=n.indexOf(t),[r[t-1],r[t]]},i.copy=function(){return e(r,n)},i}([.5],[0,1])},t.scale.identity=function(){return function t(e){function r(t){return+t}return r.invert=r,r.domain=r.range=function(t){return arguments.length?(e=t.map(r),r):e},r.ticks=function(t){return qn(e,t)},r.tickFormat=function(t,r){return d3_scale_linearTickFormat(e,t,r)},r.copy=function(){return t(e)},r}([0,1])},t.svg={},t.svg.arc=function(){var t=$n,e=ti,r=Kn,n=Qn,i=ei,a=ri,o=ni;function s(){var s=Math.max(0,+t.apply(this,arguments)),c=Math.max(0,+e.apply(this,arguments)),u=i.apply(this,arguments)-Et,f=a.apply(this,arguments)-Et,h=Math.abs(f-u),p=u>f?0:1;if(c<s&&(d=c,c=s,s=d),h>=St)return l(c,p)+(s?l(s,1-p):"")+"Z";var d,m,g,v,y,x,b,_,w,T,k,A,M=0,S=0,E=[];if((v=(+o.apply(this,arguments)||0)/2)&&(g=n===Qn?Math.sqrt(s*s+c*c):+n.apply(this,arguments),p||(S*=-1),c&&(S=Pt(g/c*Math.sin(v))),s&&(M=Pt(g/s*Math.sin(v)))),c){y=c*Math.cos(u+S),x=c*Math.sin(u+S),b=c*Math.cos(f-S),_=c*Math.sin(f-S);var L=Math.abs(f-u-2*S)<=At?0:1;if(S&&ii(y,x,b,_)===p^L){var C=(u+f)/2;y=c*Math.cos(C),x=c*Math.sin(C),b=_=null}}else y=x=0;if(s){w=s*Math.cos(f-M),T=s*Math.sin(f-M),k=s*Math.cos(u+M),A=s*Math.sin(u+M);var P=Math.abs(u-f+2*M)<=At?0:1;if(M&&ii(w,T,k,A)===1-p^P){var I=(u+f)/2;w=s*Math.cos(I),T=s*Math.sin(I),k=A=null}}else w=T=0;if(h>kt&&(d=Math.min(Math.abs(c-s)/2,+r.apply(this,arguments)))>.001){m=s<c^p?0:1;var O=d,z=d;if(h<At){var D=null==k?[w,T]:null==b?[y,x]:Se([y,x],[k,A],[b,_],[w,T]),R=y-D[0],F=x-D[1],B=b-D[0],N=_-D[1],j=1/Math.sin(Math.acos((R*B+F*N)/(Math.sqrt(R*R+F*F)*Math.sqrt(B*B+N*N)))/2),U=Math.sqrt(D[0]*D[0]+D[1]*D[1]);z=Math.min(d,(s-U)/(j-1)),O=Math.min(d,(c-U)/(j+1))}if(null!=b){var V=ai(null==k?[w,T]:[k,A],[y,x],c,O,p),H=ai([b,_],[w,T],c,O,p);d===O?E.push("M",V[0],"A",O,",",O," 0 0,",m," ",V[1],"A",c,",",c," 0 ",1-p^ii(V[1][0],V[1][1],H[1][0],H[1][1]),",",p," ",H[1],"A",O,",",O," 0 0,",m," ",H[0]):E.push("M",V[0],"A",O,",",O," 0 1,",m," ",H[0])}else E.push("M",y,",",x);if(null!=k){var q=ai([y,x],[k,A],s,-z,p),G=ai([w,T],null==b?[y,x]:[b,_],s,-z,p);d===z?E.push("L",G[0],"A",z,",",z," 0 0,",m," ",G[1],"A",s,",",s," 0 ",p^ii(G[1][0],G[1][1],q[1][0],q[1][1]),",",1-p," ",q[1],"A",z,",",z," 0 0,",m," ",q[0]):E.push("L",G[0],"A",z,",",z," 0 0,",m," ",q[0])}else E.push("L",w,",",T)}else E.push("M",y,",",x),null!=b&&E.push("A",c,",",c," 0 ",L,",",p," ",b,",",_),E.push("L",w,",",T),null!=k&&E.push("A",s,",",s," 0 ",P,",",1-p," ",k,",",A);return E.push("Z"),E.join("")}function l(t,e){return"M0,"+t+"A"+t+","+t+" 0 1,"+e+" 0,"+-t+"A"+t+","+t+" 0 1,"+e+" 0,"+t}return s.innerRadius=function(e){return arguments.length?(t=ce(e),s):t},s.outerRadius=function(t){return arguments.length?(e=ce(t),s):e},s.cornerRadius=function(t){return arguments.length?(r=ce(t),s):r},s.padRadius=function(t){return arguments.length?(n=t==Qn?Qn:ce(t),s):n},s.startAngle=function(t){return arguments.length?(i=ce(t),s):i},s.endAngle=function(t){return arguments.length?(a=ce(t),s):a},s.padAngle=function(t){return arguments.length?(o=ce(t),s):o},s.centroid=function(){var r=(+t.apply(this,arguments)+ +e.apply(this,arguments))/2,n=(+i.apply(this,arguments)+ +a.apply(this,arguments))/2-Et;return[Math.cos(n)*r,Math.sin(n)*r]},s};var Qn="auto";function $n(t){return t.innerRadius}function ti(t){return t.outerRadius}function ei(t){return t.startAngle}function ri(t){return t.endAngle}function ni(t){return t&&t.padAngle}function ii(t,e,r,n){return(t-r)*e-(e-n)*t>0?0:1}function ai(t,e,r,n,i){var a=t[0]-e[0],o=t[1]-e[1],s=(i?n:-n)/Math.sqrt(a*a+o*o),l=s*o,c=-s*a,u=t[0]+l,f=t[1]+c,h=e[0]+l,p=e[1]+c,d=(u+h)/2,m=(f+p)/2,g=h-u,v=p-f,y=g*g+v*v,x=r-n,b=u*p-h*f,_=(v<0?-1:1)*Math.sqrt(Math.max(0,x*x*y-b*b)),w=(b*v-g*_)/y,T=(-b*g-v*_)/y,k=(b*v+g*_)/y,A=(-b*g+v*_)/y,M=w-d,S=T-m,E=k-d,L=A-m;return M*M+S*S>E*E+L*L&&(w=k,T=A),[[w-l,T-c],[w*r/x,T*r/x]]}function oi(){return!0}function si(t){var e=_e,r=we,n=oi,i=ci,a=i.key,o=.7;function s(a){var s,l=[],c=[],u=-1,f=a.length,h=ce(e),p=ce(r);function d(){l.push("M",i(t(c),o))}for(;++u<f;)n.call(this,s=a[u],u)?c.push([+h.call(this,s,u),+p.call(this,s,u)]):c.length&&(d(),c=[]);return c.length&&d(),l.length?l.join(""):null}return s.x=function(t){return arguments.length?(e=t,s):e},s.y=function(t){return arguments.length?(r=t,s):r},s.defined=function(t){return arguments.length?(n=t,s):n},s.interpolate=function(t){return arguments.length?(a="function"==typeof t?i=t:(i=li.get(t)||ci).key,s):a},s.tension=function(t){return arguments.length?(o=t,s):o},s}t.svg.line=function(){return si(C)};var li=t.map({linear:ci,"linear-closed":ui,step:function(t){var e=0,r=t.length,n=t[0],i=[n[0],",",n[1]];for(;++e<r;)i.push("H",(n[0]+(n=t[e])[0])/2,"V",n[1]);r>1&&i.push("H",n[0]);return i.join("")},"step-before":fi,"step-after":hi,basis:mi,"basis-open":function(t){if(t.length<4)return ci(t);var e,r=[],n=-1,i=t.length,a=[0],o=[0];for(;++n<3;)e=t[n],a.push(e[0]),o.push(e[1]);r.push(gi(xi,a)+","+gi(xi,o)),--n;for(;++n<i;)e=t[n],a.shift(),a.push(e[0]),o.shift(),o.push(e[1]),bi(r,a,o);return r.join("")},"basis-closed":function(t){var e,r,n=-1,i=t.length,a=i+4,o=[],s=[];for(;++n<4;)r=t[n%i],o.push(r[0]),s.push(r[1]);e=[gi(xi,o),",",gi(xi,s)],--n;for(;++n<a;)r=t[n%i],o.shift(),o.push(r[0]),s.shift(),s.push(r[1]),bi(e,o,s);return e.join("")},bundle:function(t,e){var r=t.length-1;if(r)for(var n,i,a=t[0][0],o=t[0][1],s=t[r][0]-a,l=t[r][1]-o,c=-1;++c<=r;)n=t[c],i=c/r,n[0]=e*n[0]+(1-e)*(a+i*s),n[1]=e*n[1]+(1-e)*(o+i*l);return mi(t)},cardinal:function(t,e){return t.length<3?ci(t):t[0]+pi(t,di(t,e))},"cardinal-open":function(t,e){return t.length<4?ci(t):t[1]+pi(t.slice(1,-1),di(t,e))},"cardinal-closed":function(t,e){return t.length<3?ui(t):t[0]+pi((t.push(t[0]),t),di([t[t.length-2]].concat(t,[t[1]]),e))},monotone:function(t){return t.length<3?ci(t):t[0]+pi(t,function(t){var e,r,n,i,a=[],o=function(t){var e=0,r=t.length-1,n=[],i=t[0],a=t[1],o=n[0]=_i(i,a);for(;++e<r;)n[e]=(o+(o=_i(i=a,a=t[e+1])))/2;return n[e]=o,n}(t),s=-1,l=t.length-1;for(;++s<l;)e=_i(t[s],t[s+1]),y(e)<kt?o[s]=o[s+1]=0:(r=o[s]/e,n=o[s+1]/e,(i=r*r+n*n)>9&&(i=3*e/Math.sqrt(i),o[s]=i*r,o[s+1]=i*n));s=-1;for(;++s<=l;)i=(t[Math.min(l,s+1)][0]-t[Math.max(0,s-1)][0])/(6*(1+o[s]*o[s])),a.push([i||0,o[s]*i||0]);return a}(t))}});function ci(t){return t.length>1?t.join("L"):t+"Z"}function ui(t){return t.join("L")+"Z"}function fi(t){for(var e=0,r=t.length,n=t[0],i=[n[0],",",n[1]];++e<r;)i.push("V",(n=t[e])[1],"H",n[0]);return i.join("")}function hi(t){for(var e=0,r=t.length,n=t[0],i=[n[0],",",n[1]];++e<r;)i.push("H",(n=t[e])[0],"V",n[1]);return i.join("")}function pi(t,e){if(e.length<1||t.length!=e.length&&t.length!=e.length+2)return ci(t);var r=t.length!=e.length,n="",i=t[0],a=t[1],o=e[0],s=o,l=1;if(r&&(n+="Q"+(a[0]-2*o[0]/3)+","+(a[1]-2*o[1]/3)+","+a[0]+","+a[1],i=t[1],l=2),e.length>1){s=e[1],a=t[l],l++,n+="C"+(i[0]+o[0])+","+(i[1]+o[1])+","+(a[0]-s[0])+","+(a[1]-s[1])+","+a[0]+","+a[1];for(var c=2;c<e.length;c++,l++)a=t[l],s=e[c],n+="S"+(a[0]-s[0])+","+(a[1]-s[1])+","+a[0]+","+a[1]}if(r){var u=t[l];n+="Q"+(a[0]+2*s[0]/3)+","+(a[1]+2*s[1]/3)+","+u[0]+","+u[1]}return n}function di(t,e){for(var r,n=[],i=(1-e)/2,a=t[0],o=t[1],s=1,l=t.length;++s<l;)r=a,a=o,o=t[s],n.push([i*(o[0]-r[0]),i*(o[1]-r[1])]);return n}function mi(t){if(t.length<3)return ci(t);var e=1,r=t.length,n=t[0],i=n[0],a=n[1],o=[i,i,i,(n=t[1])[0]],s=[a,a,a,n[1]],l=[i,",",a,"L",gi(xi,o),",",gi(xi,s)];for(t.push(t[r-1]);++e<=r;)n=t[e],o.shift(),o.push(n[0]),s.shift(),s.push(n[1]),bi(l,o,s);return t.pop(),l.push("L",n),l.join("")}function gi(t,e){return t[0]*e[0]+t[1]*e[1]+t[2]*e[2]+t[3]*e[3]}li.forEach((function(t,e){e.key=t,e.closed=/-closed$/.test(t)}));var vi=[0,2/3,1/3,0],yi=[0,1/3,2/3,0],xi=[0,1/6,2/3,1/6];function bi(t,e,r){t.push("C",gi(vi,e),",",gi(vi,r),",",gi(yi,e),",",gi(yi,r),",",gi(xi,e),",",gi(xi,r))}function _i(t,e){return(e[1]-t[1])/(e[0]-t[0])}function wi(t){for(var e,r,n,i=-1,a=t.length;++i<a;)r=(e=t[i])[0],n=e[1]-Et,e[0]=r*Math.cos(n),e[1]=r*Math.sin(n);return t}function Ti(t){var e=_e,r=_e,n=0,i=we,a=oi,o=ci,s=o.key,l=o,c="L",u=.7;function f(s){var f,h,p,d=[],m=[],g=[],v=-1,y=s.length,x=ce(e),b=ce(n),_=e===r?function(){return h}:ce(r),w=n===i?function(){return p}:ce(i);function T(){d.push("M",o(t(g),u),c,l(t(m.reverse()),u),"Z")}for(;++v<y;)a.call(this,f=s[v],v)?(m.push([h=+x.call(this,f,v),p=+b.call(this,f,v)]),g.push([+_.call(this,f,v),+w.call(this,f,v)])):m.length&&(T(),m=[],g=[]);return m.length&&T(),d.length?d.join(""):null}return f.x=function(t){return arguments.length?(e=r=t,f):r},f.x0=function(t){return arguments.length?(e=t,f):e},f.x1=function(t){return arguments.length?(r=t,f):r},f.y=function(t){return arguments.length?(n=i=t,f):i},f.y0=function(t){return arguments.length?(n=t,f):n},f.y1=function(t){return arguments.length?(i=t,f):i},f.defined=function(t){return arguments.length?(a=t,f):a},f.interpolate=function(t){return arguments.length?(s="function"==typeof t?o=t:(o=li.get(t)||ci).key,l=o.reverse||o,c=o.closed?"M":"L",f):s},f.tension=function(t){return arguments.length?(u=t,f):u},f}function ki(t){return t.source}function Ai(t){return t.target}function Mi(t){return t.radius}function Si(t){return[t.x,t.y]}function Ei(t){return function(){var e=t.apply(this,arguments),r=e[0],n=e[1]-Et;return[r*Math.cos(n),r*Math.sin(n)]}}function Li(){return 64}function Ci(){return"circle"}function Pi(t){var e=Math.sqrt(t/At);return"M0,"+e+"A"+e+","+e+" 0 1,1 0,"+-e+"A"+e+","+e+" 0 1,1 0,"+e+"Z"}t.svg.line.radial=function(){var t=si(wi);return t.radius=t.x,delete t.x,t.angle=t.y,delete t.y,t},fi.reverse=hi,hi.reverse=fi,t.svg.area=function(){return Ti(C)},t.svg.area.radial=function(){var t=Ti(wi);return t.radius=t.x,delete t.x,t.innerRadius=t.x0,delete t.x0,t.outerRadius=t.x1,delete t.x1,t.angle=t.y,delete t.y,t.startAngle=t.y0,delete t.y0,t.endAngle=t.y1,delete t.y1,t},t.svg.chord=function(){var t=ki,e=Ai,r=Mi,n=ei,i=ri;function a(r,n){var i,a,c=o(this,t,r,n),u=o(this,e,r,n);return"M"+c.p0+s(c.r,c.p1,c.a1-c.a0)+(a=u,((i=c).a0==a.a0&&i.a1==a.a1?l(c.r,c.p1,c.r,c.p0):l(c.r,c.p1,u.r,u.p0)+s(u.r,u.p1,u.a1-u.a0)+l(u.r,u.p1,c.r,c.p0))+"Z")}function o(t,e,a,o){var s=e.call(t,a,o),l=r.call(t,s,o),c=n.call(t,s,o)-Et,u=i.call(t,s,o)-Et;return{r:l,a0:c,a1:u,p0:[l*Math.cos(c),l*Math.sin(c)],p1:[l*Math.cos(u),l*Math.sin(u)]}}function s(t,e,r){return"A"+t+","+t+" 0 "+ +(r>At)+",1 "+e}function l(t,e,r,n){return"Q 0,0 "+n}return a.radius=function(t){return arguments.length?(r=ce(t),a):r},a.source=function(e){return arguments.length?(t=ce(e),a):t},a.target=function(t){return arguments.length?(e=ce(t),a):e},a.startAngle=function(t){return arguments.length?(n=ce(t),a):n},a.endAngle=function(t){return arguments.length?(i=ce(t),a):i},a},t.svg.diagonal=function(){var t=ki,e=Ai,r=Si;function n(n,i){var a=t.call(this,n,i),o=e.call(this,n,i),s=(a.y+o.y)/2,l=[a,{x:a.x,y:s},{x:o.x,y:s},o];return"M"+(l=l.map(r))[0]+"C"+l[1]+" "+l[2]+" "+l[3]}return n.source=function(e){return arguments.length?(t=ce(e),n):t},n.target=function(t){return arguments.length?(e=ce(t),n):e},n.projection=function(t){return arguments.length?(r=t,n):r},n},t.svg.diagonal.radial=function(){var e=t.svg.diagonal(),r=Si,n=e.projection;return e.projection=function(t){return arguments.length?n(Ei(r=t)):r},e},t.svg.symbol=function(){var t=Ci,e=Li;function r(r,n){return(Ii.get(t.call(this,r,n))||Pi)(e.call(this,r,n))}return r.type=function(e){return arguments.length?(t=ce(e),r):t},r.size=function(t){return arguments.length?(e=ce(t),r):e},r};var Ii=t.map({circle:Pi,cross:function(t){var e=Math.sqrt(t/5)/2;return"M"+-3*e+","+-e+"H"+-e+"V"+-3*e+"H"+e+"V"+-e+"H"+3*e+"V"+e+"H"+e+"V"+3*e+"H"+-e+"V"+e+"H"+-3*e+"Z"},diamond:function(t){var e=Math.sqrt(t/(2*zi)),r=e*zi;return"M0,"+-e+"L"+r+",0 0,"+e+" "+-r+",0Z"},square:function(t){var e=Math.sqrt(t)/2;return"M"+-e+","+-e+"L"+e+","+-e+" "+e+","+e+" "+-e+","+e+"Z"},"triangle-down":function(t){var e=Math.sqrt(t/Oi),r=e*Oi/2;return"M0,"+r+"L"+e+","+-r+" "+-e+","+-r+"Z"},"triangle-up":function(t){var e=Math.sqrt(t/Oi),r=e*Oi/2;return"M0,"+-r+"L"+e+","+r+" "+-e+","+r+"Z"}});t.svg.symbolTypes=Ii.keys();var Oi=Math.sqrt(3),zi=Math.tan(30*Lt);Y.transition=function(t){for(var e,r,n=Bi||++Ui,i=qi(t),a=[],o=Ni||{time:Date.now(),ease:Er,delay:0,duration:250},s=-1,l=this.length;++s<l;){a.push(e=[]);for(var c=this[s],u=-1,f=c.length;++u<f;)(r=c[u])&&Gi(r,u,i,n,o),e.push(r)}return Fi(a,i,n)},Y.interrupt=function(t){return this.each(null==t?Di:Ri(qi(t)))};var Di=Ri(qi());function Ri(t){return function(){var e,r,n;(e=this[t])&&(n=e[r=e.active])&&(n.timer.c=null,n.timer.t=NaN,--e.count?delete e[r]:delete this[t],e.active+=.5,n.event&&n.event.interrupt.call(this,this.__data__,n.index))}}function Fi(t,e,r){return U(t,ji),t.namespace=e,t.id=r,t}var Bi,Ni,ji=[],Ui=0;function Vi(t,e,r,n){var i=t.id,a=t.namespace;return ut(t,"function"==typeof r?function(t,o,s){t[a][i].tween.set(e,n(r.call(t,t.__data__,o,s)))}:(r=n(r),function(t){t[a][i].tween.set(e,r)}))}function Hi(t){return null==t&&(t=""),function(){this.textContent=t}}function qi(t){return null==t?"__transition__":"__transition_"+t+"__"}function Gi(t,e,r,n,i){var a,o,s,l,c,u=t[r]||(t[r]={active:0,count:0}),f=u[n];function h(r){var i=u.active,h=u[i];for(var d in h&&(h.timer.c=null,h.timer.t=NaN,--u.count,delete u[i],h.event&&h.event.interrupt.call(t,t.__data__,h.index)),u)if(+d<n){var m=u[d];m.timer.c=null,m.timer.t=NaN,--u.count,delete u[d]}o.c=p,ve((function(){return o.c&&p(r||1)&&(o.c=null,o.t=NaN),1}),0,a),u.active=n,f.event&&f.event.start.call(t,t.__data__,e),c=[],f.tween.forEach((function(r,n){(n=n.call(t,t.__data__,e))&&c.push(n)})),l=f.ease,s=f.duration}function p(i){for(var a=i/s,o=l(a),h=c.length;h>0;)c[--h].call(t,o);if(a>=1)return f.event&&f.event.end.call(t,t.__data__,e),--u.count?delete u[n]:delete t[r],1}f||(a=i.time,o=ve((function(t){var e=f.delay;if(o.t=e+a,e<=t)return h(t-e);o.c=h}),0,a),f=u[n]={tween:new _,time:a,timer:o,delay:i.delay,duration:i.duration,ease:i.ease,index:e},i=null,++u.count)}ji.call=Y.call,ji.empty=Y.empty,ji.node=Y.node,ji.size=Y.size,t.transition=function(e,r){return e&&e.transition?Bi?e.transition(r):e:t.selection().transition(e)},t.transition.prototype=ji,ji.select=function(t){var e,r,n,i=this.id,a=this.namespace,o=[];t=W(t);for(var s=-1,l=this.length;++s<l;){o.push(e=[]);for(var c=this[s],u=-1,f=c.length;++u<f;)(n=c[u])&&(r=t.call(n,n.__data__,u,s))?("__data__"in n&&(r.__data__=n.__data__),Gi(r,u,a,i,n[a][i]),e.push(r)):e.push(null)}return Fi(o,a,i)},ji.selectAll=function(t){var e,r,n,i,a,o=this.id,s=this.namespace,l=[];t=X(t);for(var c=-1,u=this.length;++c<u;)for(var f=this[c],h=-1,p=f.length;++h<p;)if(n=f[h]){a=n[s][o],r=t.call(n,n.__data__,h,c),l.push(e=[]);for(var d=-1,m=r.length;++d<m;)(i=r[d])&&Gi(i,d,s,o,a),e.push(i)}return Fi(l,s,o)},ji.filter=function(t){var e,r,n=[];"function"!=typeof t&&(t=lt(t));for(var i=0,a=this.length;i<a;i++){n.push(e=[]);for(var o,s=0,l=(o=this[i]).length;s<l;s++)(r=o[s])&&t.call(r,r.__data__,s,i)&&e.push(r)}return Fi(n,this.namespace,this.id)},ji.tween=function(t,e){var r=this.id,n=this.namespace;return arguments.length<2?this.node()[n][r].tween.get(t):ut(this,null==e?function(e){e[n][r].tween.remove(t)}:function(i){i[n][r].tween.set(t,e)})},ji.attr=function(e,r){if(arguments.length<2){for(r in e)this.attr(r,e[r]);return this}var n="transform"==e?Nr:yr,i=t.ns.qualify(e);function a(){this.removeAttribute(i)}function o(){this.removeAttributeNS(i.space,i.local)}function s(t){return null==t?a:(t+="",function(){var e,r=this.getAttribute(i);return r!==t&&(e=n(r,t),function(t){this.setAttribute(i,e(t))})})}function l(t){return null==t?o:(t+="",function(){var e,r=this.getAttributeNS(i.space,i.local);return r!==t&&(e=n(r,t),function(t){this.setAttributeNS(i.space,i.local,e(t))})})}return Vi(this,"attr."+e,r,i.local?l:s)},ji.attrTween=function(e,r){var n=t.ns.qualify(e);return this.tween("attr."+e,n.local?function(t,e){var i=r.call(this,t,e,this.getAttributeNS(n.space,n.local));return i&&function(t){this.setAttributeNS(n.space,n.local,i(t))}}:function(t,e){var i=r.call(this,t,e,this.getAttribute(n));return i&&function(t){this.setAttribute(n,i(t))}})},ji.style=function(t,e,r){var n=arguments.length;if(n<3){if("string"!=typeof t){for(r in n<2&&(e=""),t)this.style(r,t[r],e);return this}r=""}function i(){this.style.removeProperty(t)}function a(e){return null==e?i:(e+="",function(){var n,i=o(this).getComputedStyle(this,null).getPropertyValue(t);return i!==e&&(n=yr(i,e),function(e){this.style.setProperty(t,n(e),r)})})}return Vi(this,"style."+t,e,a)},ji.styleTween=function(t,e,r){function n(n,i){var a=e.call(this,n,i,o(this).getComputedStyle(this,null).getPropertyValue(t));return a&&function(e){this.style.setProperty(t,a(e),r)}}return arguments.length<3&&(r=""),this.tween("style."+t,n)},ji.text=function(t){return Vi(this,"text",t,Hi)},ji.remove=function(){var t=this.namespace;return this.each("end.transition",(function(){var e;this[t].count<2&&(e=this.parentNode)&&e.removeChild(this)}))},ji.ease=function(e){var r=this.id,n=this.namespace;return arguments.length<1?this.node()[n][r].ease:("function"!=typeof e&&(e=t.ease.apply(t,arguments)),ut(this,(function(t){t[n][r].ease=e})))},ji.delay=function(t){var e=this.id,r=this.namespace;return arguments.length<1?this.node()[r][e].delay:ut(this,"function"==typeof t?function(n,i,a){n[r][e].delay=+t.call(n,n.__data__,i,a)}:(t=+t,function(n){n[r][e].delay=t}))},ji.duration=function(t){var e=this.id,r=this.namespace;return arguments.length<1?this.node()[r][e].duration:ut(this,"function"==typeof t?function(n,i,a){n[r][e].duration=Math.max(1,t.call(n,n.__data__,i,a))}:(t=Math.max(1,t),function(n){n[r][e].duration=t}))},ji.each=function(e,r){var n=this.id,i=this.namespace;if(arguments.length<2){var a=Ni,o=Bi;try{Bi=n,ut(this,(function(t,r,a){Ni=t[i][n],e.call(t,t.__data__,r,a)}))}finally{Ni=a,Bi=o}}else ut(this,(function(a){var o=a[i][n];(o.event||(o.event=t.dispatch("start","end","interrupt"))).on(e,r)}));return this},ji.transition=function(){for(var t,e,r,n=this.id,i=++Ui,a=this.namespace,o=[],s=0,l=this.length;s<l;s++){o.push(t=[]);for(var c,u=0,f=(c=this[s]).length;u<f;u++)(e=c[u])&&Gi(e,u,a,i,{time:(r=e[a][n]).time,ease:r.ease,delay:r.delay+r.duration,duration:r.duration}),t.push(e)}return Fi(o,a,i)},t.svg.axis=function(){var e,r=t.scale.linear(),i=Yi,a=6,o=6,s=3,l=[10],c=null;function u(n){n.each((function(){var n,u=t.select(this),f=this.__chart__||r,h=this.__chart__=r.copy(),p=null==c?h.ticks?h.ticks.apply(h,l):h.domain():c,d=null==e?h.tickFormat?h.tickFormat.apply(h,l):C:e,m=u.selectAll(".tick").data(p,h),g=m.enter().insert("g",".domain").attr("class","tick").style("opacity",kt),v=t.transition(m.exit()).style("opacity",kt).remove(),y=t.transition(m.order()).style("opacity",1),x=Math.max(a,0)+s,b=Dn(h),_=u.selectAll(".domain").data([0]),w=(_.enter().append("path").attr("class","domain"),t.transition(_));g.append("line"),g.append("text");var T,k,A,M,S=g.select("line"),E=y.select("line"),L=m.select("text").text(d),P=g.select("text"),I=y.select("text"),O="top"===i||"left"===i?-1:1;if("bottom"===i||"top"===i?(n=Xi,T="x",A="y",k="x2",M="y2",L.attr("dy",O<0?"0em":".71em").style("text-anchor","middle"),w.attr("d","M"+b[0]+","+O*o+"V0H"+b[1]+"V"+O*o)):(n=Zi,T="y",A="x",k="y2",M="x2",L.attr("dy",".32em").style("text-anchor",O<0?"end":"start"),w.attr("d","M"+O*o+","+b[0]+"H0V"+b[1]+"H"+O*o)),S.attr(M,O*a),P.attr(A,O*x),E.attr(k,0).attr(M,O*a),I.attr(T,0).attr(A,O*x),h.rangeBand){var z=h,D=z.rangeBand()/2;f=h=function(t){return z(t)+D}}else f.rangeBand?f=h:v.call(n,h,f);g.call(n,f,h),y.call(n,h,h)}))}return u.scale=function(t){return arguments.length?(r=t,u):r},u.orient=function(t){return arguments.length?(i=t in Wi?t+"":Yi,u):i},u.ticks=function(){return arguments.length?(l=n(arguments),u):l},u.tickValues=function(t){return arguments.length?(c=t,u):c},u.tickFormat=function(t){return arguments.length?(e=t,u):e},u.tickSize=function(t){var e=arguments.length;return e?(a=+t,o=+arguments[e-1],u):a},u.innerTickSize=function(t){return arguments.length?(a=+t,u):a},u.outerTickSize=function(t){return arguments.length?(o=+t,u):o},u.tickPadding=function(t){return arguments.length?(s=+t,u):s},u.tickSubdivide=function(){return arguments.length&&u},u};var Yi="bottom",Wi={top:1,right:1,bottom:1,left:1};function Xi(t,e,r){t.attr("transform",(function(t){var n=e(t);return"translate("+(isFinite(n)?n:r(t))+",0)"}))}function Zi(t,e,r){t.attr("transform",(function(t){var n=e(t);return"translate(0,"+(isFinite(n)?n:r(t))+")"}))}t.svg.brush=function(){var e,r,n=N(h,"brushstart","brush","brushend"),i=null,a=null,s=[0,0],l=[0,0],c=!0,u=!0,f=Ki[0];function h(e){e.each((function(){var e=t.select(this).style("pointer-events","all").style("-webkit-tap-highlight-color","rgba(0,0,0,0)").on("mousedown.brush",g).on("touchstart.brush",g),r=e.selectAll(".background").data([0]);r.enter().append("rect").attr("class","background").style("visibility","hidden").style("cursor","crosshair"),e.selectAll(".extent").data([0]).enter().append("rect").attr("class","extent").style("cursor","move");var n=e.selectAll(".resize").data(f,C);n.exit().remove(),n.enter().append("g").attr("class",(function(t){return"resize "+t})).style("cursor",(function(t){return Ji[t]})).append("rect").attr("x",(function(t){return/[ew]$/.test(t)?-3:null})).attr("y",(function(t){return/^[ns]/.test(t)?-3:null})).attr("width",6).attr("height",6).style("visibility","hidden"),n.style("display",h.empty()?"none":null);var o,s=t.transition(e),l=t.transition(r);i&&(o=Dn(i),l.attr("x",o[0]).attr("width",o[1]-o[0]),d(s)),a&&(o=Dn(a),l.attr("y",o[0]).attr("height",o[1]-o[0]),m(s)),p(s)}))}function p(t){t.selectAll(".resize").attr("transform",(function(t){return"translate("+s[+/e$/.test(t)]+","+l[+/^s/.test(t)]+")"}))}function d(t){t.select(".extent").attr("x",s[0]),t.selectAll(".extent,.n>rect,.s>rect").attr("width",s[1]-s[0])}function m(t){t.select(".extent").attr("y",l[0]),t.selectAll(".extent,.e>rect,.w>rect").attr("height",l[1]-l[0])}function g(){var f,g,v=this,y=t.select(t.event.target),x=n.of(v,arguments),b=t.select(v),_=y.datum(),w=!/^(n|s)$/.test(_)&&i,T=!/^(e|w)$/.test(_)&&a,k=y.classed("extent"),A=bt(v),M=t.mouse(v),S=t.select(o(v)).on("keydown.brush",C).on("keyup.brush",P);if(t.event.changedTouches?S.on("touchmove.brush",I).on("touchend.brush",z):S.on("mousemove.brush",I).on("mouseup.brush",z),b.interrupt().selectAll("*").interrupt(),k)M[0]=s[0]-M[0],M[1]=l[0]-M[1];else if(_){var E=+/w$/.test(_),L=+/^n/.test(_);g=[s[1-E]-M[0],l[1-L]-M[1]],M[0]=s[E],M[1]=l[L]}else t.event.altKey&&(f=M.slice());function C(){32==t.event.keyCode&&(k||(f=null,M[0]-=s[1],M[1]-=l[1],k=2),F())}function P(){32==t.event.keyCode&&2==k&&(M[0]+=s[1],M[1]+=l[1],k=0,F())}function I(){var e=t.mouse(v),r=!1;g&&(e[0]+=g[0],e[1]+=g[1]),k||(t.event.altKey?(f||(f=[(s[0]+s[1])/2,(l[0]+l[1])/2]),M[0]=s[+(e[0]<f[0])],M[1]=l[+(e[1]<f[1])]):f=null),w&&O(e,i,0)&&(d(b),r=!0),T&&O(e,a,1)&&(m(b),r=!0),r&&(p(b),x({type:"brush",mode:k?"move":"resize"}))}function O(t,n,i){var a,o,h=Dn(n),p=h[0],d=h[1],m=M[i],g=i?l:s,v=g[1]-g[0];if(k&&(p-=m,d-=v+m),a=(i?u:c)?Math.max(p,Math.min(d,t[i])):t[i],k?o=(a+=m)+v:(f&&(m=Math.max(p,Math.min(d,2*f[i]-a))),m<a?(o=a,a=m):o=m),g[0]!=a||g[1]!=o)return i?r=null:e=null,g[0]=a,g[1]=o,!0}function z(){I(),b.style("pointer-events","all").selectAll(".resize").style("display",h.empty()?"none":null),t.select("body").style("cursor",null),S.on("mousemove.brush",null).on("mouseup.brush",null).on("touchmove.brush",null).on("touchend.brush",null).on("keydown.brush",null).on("keyup.brush",null),A(),x({type:"brushend"})}b.style("pointer-events","none").selectAll(".resize").style("display",null),t.select("body").style("cursor",y.style("cursor")),x({type:"brushstart"}),I()}return h.event=function(i){i.each((function(){var i=n.of(this,arguments),a={x:s,y:l,i:e,j:r},o=this.__chart__||a;this.__chart__=a,Bi?t.select(this).transition().each("start.brush",(function(){e=o.i,r=o.j,s=o.x,l=o.y,i({type:"brushstart"})})).tween("brush:brush",(function(){var t=xr(s,a.x),n=xr(l,a.y);return e=r=null,function(e){s=a.x=t(e),l=a.y=n(e),i({type:"brush",mode:"resize"})}})).each("end.brush",(function(){e=a.i,r=a.j,i({type:"brush",mode:"resize"}),i({type:"brushend"})})):(i({type:"brushstart"}),i({type:"brush",mode:"resize"}),i({type:"brushend"}))}))},h.x=function(t){return arguments.length?(f=Ki[!(i=t)<<1|!a],h):i},h.y=function(t){return arguments.length?(f=Ki[!i<<1|!(a=t)],h):a},h.clamp=function(t){return arguments.length?(i&&a?(c=!!t[0],u=!!t[1]):i?c=!!t:a&&(u=!!t),h):i&&a?[c,u]:i?c:a?u:null},h.extent=function(t){var n,o,c,u,f;return arguments.length?(i&&(n=t[0],o=t[1],a&&(n=n[0],o=o[0]),e=[n,o],i.invert&&(n=i(n),o=i(o)),o<n&&(f=n,n=o,o=f),n==s[0]&&o==s[1]||(s=[n,o])),a&&(c=t[0],u=t[1],i&&(c=c[1],u=u[1]),r=[c,u],a.invert&&(c=a(c),u=a(u)),u<c&&(f=c,c=u,u=f),c==l[0]&&u==l[1]||(l=[c,u])),h):(i&&(e?(n=e[0],o=e[1]):(n=s[0],o=s[1],i.invert&&(n=i.invert(n),o=i.invert(o)),o<n&&(f=n,n=o,o=f))),a&&(r?(c=r[0],u=r[1]):(c=l[0],u=l[1],a.invert&&(c=a.invert(c),u=a.invert(u)),u<c&&(f=c,c=u,u=f))),i&&a?[[n,c],[o,u]]:i?[n,o]:a&&[c,u])},h.clear=function(){return h.empty()||(s=[0,0],l=[0,0],e=r=null),h},h.empty=function(){return!!i&&s[0]==s[1]||!!a&&l[0]==l[1]},t.rebind(h,n,"on")};var Ji={n:"ns-resize",e:"ew-resize",s:"ns-resize",w:"ew-resize",nw:"nwse-resize",ne:"nesw-resize",se:"nwse-resize",sw:"nesw-resize"},Ki=[["n","e","s","w","nw","ne","se","sw"],["e","w"],["n","s"],[]];function Qi(t){return JSON.parse(t.responseText)}function $i(t){var e=i.createRange();return e.selectNode(i.body),e.createContextualFragment(t.responseText)}t.text=ue((function(t){return t.responseText})),t.json=function(t,e){return fe(t,"application/json",Qi,e)},t.html=function(t,e){return fe(t,"text/html",$i,e)},t.xml=ue((function(t){return t.responseXML})),"object"==typeof e&&e.exports?e.exports=t:this.d3=t}).apply(self)},{}],59:[function(t,e,r){"use strict";e.exports=t("./quad")},{"./quad":60}],60:[function(t,e,r){"use strict";var n=t("binary-search-bounds"),i=t("clamp"),a=t("parse-rect"),o=t("array-bounds"),s=t("pick-by-alias"),l=t("defined"),c=t("flatten-vertex-data"),u=t("is-obj"),f=t("dtype"),h=t("math-log2");function p(t,e){for(var r=e[0],n=e[1],a=1/(e[2]-r),o=1/(e[3]-n),s=new Array(t.length),l=0,c=t.length/2;l<c;l++)s[2*l]=i((t[2*l]-r)*a,0,1),s[2*l+1]=i((t[2*l+1]-n)*o,0,1);return s}e.exports=function(t,e){e||(e={}),t=c(t,"float64"),e=s(e,{bounds:"range bounds dataBox databox",maxDepth:"depth maxDepth maxdepth level maxLevel maxlevel levels",dtype:"type dtype format out dst output destination"});var r=l(e.maxDepth,255),i=l(e.bounds,o(t,2));i[0]===i[2]&&i[2]++,i[1]===i[3]&&i[3]++;var d,m=p(t,i),g=t.length>>>1;e.dtype||(e.dtype="array"),"string"==typeof e.dtype?d=new(f(e.dtype))(g):e.dtype&&(d=e.dtype,Array.isArray(d)&&(d.length=g));for(var v=0;v<g;++v)d[v]=v;var y=[],x=[],b=[],_=[];!function t(e,n,i,a,o,s){if(!a.length)return null;var l=y[o]||(y[o]=[]),c=b[o]||(b[o]=[]),u=x[o]||(x[o]=[]),f=l.length;if(++o>r||s>1073741824){for(var h=0;h<a.length;h++)l.push(a[h]),c.push(s),u.push(null,null,null,null);return f}if(l.push(a[0]),c.push(s),a.length<=1)return u.push(null,null,null,null),f;for(var p=.5*i,d=e+p,g=n+p,v=[],_=[],w=[],T=[],k=1,A=a.length;k<A;k++){var M=a[k],S=m[2*M],E=m[2*M+1];S<d?E<g?v.push(M):_.push(M):E<g?w.push(M):T.push(M)}return s<<=2,u.push(t(e,n,p,v,o,s),t(e,g,p,_,o,s+1),t(d,n,p,w,o,s+2),t(d,g,p,T,o,s+3)),f}(0,0,1,d,0,1);for(var w=0,T=0;T<y.length;T++){var k=y[T];if(d.set)d.set(k,w);else for(var A=0,M=k.length;A<M;A++)d[A+w]=k[A];var S=w+y[T].length;_[T]=[w,S],w=S}return d.range=function(){var e,r=[],n=arguments.length;for(;n--;)r[n]=arguments[n];if(u(r[r.length-1])){var o=r.pop();r.length||null==o.x&&null==o.l&&null==o.left||(r=[o],e={}),e=s(o,{level:"level maxLevel",d:"d diam diameter r radius px pxSize pixel pixelSize maxD size minSize",lod:"lod details ranges offsets"})}else e={};r.length||(r=i);var c=a.apply(void 0,r),f=[Math.min(c.x,c.x+c.width),Math.min(c.y,c.y+c.height),Math.max(c.x,c.x+c.width),Math.max(c.y,c.y+c.height)],d=f[0],m=f[1],g=f[2],v=f[3],b=p([d,m,g,v],i),_=b[0],w=b[1],T=b[2],k=b[3],A=l(e.level,y.length);if(null!=e.d){var M;"number"==typeof e.d?M=[e.d,e.d]:e.d.length&&(M=e.d),A=Math.min(Math.max(Math.ceil(-h(Math.abs(M[0])/(i[2]-i[0]))),Math.ceil(-h(Math.abs(M[1])/(i[3]-i[1])))),A)}if(A=Math.min(A,y.length),e.lod)return E(_,w,T,k,A);var S=[];function L(e,r,n,i,a,o){if(null!==a&&null!==o&&!(_>e+n||w>r+n||T<e||k<r||i>=A||a===o)){var s=y[i];void 0===o&&(o=s.length);for(var l=a;l<o;l++){var c=s[l],u=t[2*c],f=t[2*c+1];u>=d&&u<=g&&f>=m&&f<=v&&S.push(c)}var h=x[i],p=h[4*a+0],b=h[4*a+1],M=h[4*a+2],E=h[4*a+3],P=C(h,a+1),I=.5*n,O=i+1;L(e,r,I,O,p,b||M||E||P),L(e,r+I,I,O,b,M||E||P),L(e+I,r,I,O,M,E||P),L(e+I,r+I,I,O,E,P)}}function C(t,e){for(var r=null,n=0;null===r;)if(r=t[4*e+n],++n>t.length)return null;return r}return L(0,0,1,0,0,1),S},d;function E(t,e,r,i,a){for(var o=[],s=0;s<a;s++){var l=b[s],c=_[s][0],u=L(t,e,s),f=L(r,i,s),h=n.ge(l,u),p=n.gt(l,f,h,l.length-1);o[s]=[h+c,p+c]}return o}function L(t,e,r){for(var n=1,i=.5,a=.5,o=.5,s=0;s<r;s++)n<<=2,n+=t<i?e<a?0:1:e<a?2:3,o*=.5,i+=t<i?-o:o,a+=e<a?-o:o;return n}}},{"array-bounds":72,"binary-search-bounds":100,clamp:121,defined:171,dtype:176,"flatten-vertex-data":244,"is-obj":436,"math-log2":446,"parse-rect":471,"pick-by-alias":475}],61:[function(t,e,r){"use strict";Object.defineProperty(r,"__esModule",{value:!0});var n=t("@turf/meta");function i(t){var e=0;if(t&&t.length>0){e+=Math.abs(a(t[0]));for(var r=1;r<t.length;r++)e-=Math.abs(a(t[r]))}return e}function a(t){var e,r,n,i,a,s,l=0,c=t.length;if(c>2){for(s=0;s<c;s++)s===c-2?(n=c-2,i=c-1,a=0):s===c-1?(n=c-1,i=0,a=1):(n=s,i=s+1,a=s+2),e=t[n],r=t[i],l+=(o(t[a][0])-o(e[0]))*Math.sin(o(r[1]));l=6378137*l*6378137/2}return l}function o(t){return t*Math.PI/180}r.default=function(t){return n.geomReduce(t,(function(t,e){return t+function(t){var e,r=0;switch(t.type){case"Polygon":return i(t.coordinates);case"MultiPolygon":for(e=0;e<t.coordinates.length;e++)r+=i(t.coordinates[e]);return r;case"Point":case"MultiPoint":case"LineString":case"MultiLineString":return 0}return 0}(e)}),0)}},{"@turf/meta":65}],62:[function(t,e,r){"use strict";Object.defineProperty(r,"__esModule",{value:!0});var n=t("@turf/meta");function i(t){var e=[1/0,1/0,-1/0,-1/0];return n.coordEach(t,(function(t){e[0]>t[0]&&(e[0]=t[0]),e[1]>t[1]&&(e[1]=t[1]),e[2]<t[0]&&(e[2]=t[0]),e[3]<t[1]&&(e[3]=t[1])})),e}i.default=i,r.default=i},{"@turf/meta":65}],63:[function(t,e,r){"use strict";Object.defineProperty(r,"__esModule",{value:!0});var n=t("@turf/meta"),i=t("@turf/helpers");r.default=function(t,e){void 0===e&&(e={});var r=0,a=0,o=0;return n.coordEach(t,(function(t){r+=t[0],a+=t[1],o++}),!0),i.point([r/o,a/o],e.properties)}},{"@turf/helpers":64,"@turf/meta":65}],64:[function(t,e,r){"use strict";function n(t,e,r){void 0===r&&(r={});var n={type:"Feature"};return(0===r.id||r.id)&&(n.id=r.id),r.bbox&&(n.bbox=r.bbox),n.properties=e||{},n.geometry=t,n}function i(t,e,r){if(void 0===r&&(r={}),!t)throw new Error("coordinates is required");if(!Array.isArray(t))throw new Error("coordinates must be an Array");if(t.length<2)throw new Error("coordinates must be at least 2 numbers long");if(!d(t[0])||!d(t[1]))throw new Error("coordinates must contain numbers");return n({type:"Point",coordinates:t},e,r)}function a(t,e,r){void 0===r&&(r={});for(var i=0,a=t;i<a.length;i++){var o=a[i];if(o.length<4)throw new Error("Each LinearRing of a Polygon must have 4 or more Positions.");for(var s=0;s<o[o.length-1].length;s++)if(o[o.length-1][s]!==o[0][s])throw new Error("First and last Position are not equivalent.")}return n({type:"Polygon",coordinates:t},e,r)}function o(t,e,r){if(void 0===r&&(r={}),t.length<2)throw new Error("coordinates must be an array of two or more positions");return n({type:"LineString",coordinates:t},e,r)}function s(t,e){void 0===e&&(e={});var r={type:"FeatureCollection"};return e.id&&(r.id=e.id),e.bbox&&(r.bbox=e.bbox),r.features=t,r}function l(t,e,r){return void 0===r&&(r={}),n({type:"MultiLineString",coordinates:t},e,r)}function c(t,e,r){return void 0===r&&(r={}),n({type:"MultiPoint",coordinates:t},e,r)}function u(t,e,r){return void 0===r&&(r={}),n({type:"MultiPolygon",coordinates:t},e,r)}function f(t,e){void 0===e&&(e="kilometers");var n=r.factors[e];if(!n)throw new Error(e+" units is invalid");return t*n}function h(t,e){void 0===e&&(e="kilometers");var n=r.factors[e];if(!n)throw new Error(e+" units is invalid");return t/n}function p(t){return 180*(t%(2*Math.PI))/Math.PI}function d(t){return!isNaN(t)&&null!==t&&!Array.isArray(t)}Object.defineProperty(r,"__esModule",{value:!0}),r.earthRadius=6371008.8,r.factors={centimeters:100*r.earthRadius,centimetres:100*r.earthRadius,degrees:r.earthRadius/111325,feet:3.28084*r.earthRadius,inches:39.37*r.earthRadius,kilometers:r.earthRadius/1e3,kilometres:r.earthRadius/1e3,meters:r.earthRadius,metres:r.earthRadius,miles:r.earthRadius/1609.344,millimeters:1e3*r.earthRadius,millimetres:1e3*r.earthRadius,nauticalmiles:r.earthRadius/1852,radians:1,yards:1.0936*r.earthRadius},r.unitsFactors={centimeters:100,centimetres:100,degrees:1/111325,feet:3.28084,inches:39.37,kilometers:.001,kilometres:.001,meters:1,metres:1,miles:1/1609.344,millimeters:1e3,millimetres:1e3,nauticalmiles:1/1852,radians:1/r.earthRadius,yards:1.0936133},r.areaFactors={acres:247105e-9,centimeters:1e4,centimetres:1e4,feet:10.763910417,hectares:1e-4,inches:1550.003100006,kilometers:1e-6,kilometres:1e-6,meters:1,metres:1,miles:386e-9,millimeters:1e6,millimetres:1e6,yards:1.195990046},r.feature=n,r.geometry=function(t,e,r){switch(void 0===r&&(r={}),t){case"Point":return i(e).geometry;case"LineString":return o(e).geometry;case"Polygon":return a(e).geometry;case"MultiPoint":return c(e).geometry;case"MultiLineString":return l(e).geometry;case"MultiPolygon":return u(e).geometry;default:throw new Error(t+" is invalid")}},r.point=i,r.points=function(t,e,r){return void 0===r&&(r={}),s(t.map((function(t){return i(t,e)})),r)},r.polygon=a,r.polygons=function(t,e,r){return void 0===r&&(r={}),s(t.map((function(t){return a(t,e)})),r)},r.lineString=o,r.lineStrings=function(t,e,r){return void 0===r&&(r={}),s(t.map((function(t){return o(t,e)})),r)},r.featureCollection=s,r.multiLineString=l,r.multiPoint=c,r.multiPolygon=u,r.geometryCollection=function(t,e,r){return void 0===r&&(r={}),n({type:"GeometryCollection",geometries:t},e,r)},r.round=function(t,e){if(void 0===e&&(e=0),e&&!(e>=0))throw new Error("precision must be a positive number");var r=Math.pow(10,e||0);return Math.round(t*r)/r},r.radiansToLength=f,r.lengthToRadians=h,r.lengthToDegrees=function(t,e){return p(h(t,e))},r.bearingToAzimuth=function(t){var e=t%360;return e<0&&(e+=360),e},r.radiansToDegrees=p,r.degreesToRadians=function(t){return t%360*Math.PI/180},r.convertLength=function(t,e,r){if(void 0===e&&(e="kilometers"),void 0===r&&(r="kilometers"),!(t>=0))throw new Error("length must be a positive number");return f(h(t,e),r)},r.convertArea=function(t,e,n){if(void 0===e&&(e="meters"),void 0===n&&(n="kilometers"),!(t>=0))throw new Error("area must be a positive number");var i=r.areaFactors[e];if(!i)throw new Error("invalid original units");var a=r.areaFactors[n];if(!a)throw new Error("invalid final units");return t/i*a},r.isNumber=d,r.isObject=function(t){return!!t&&t.constructor===Object},r.validateBBox=function(t){if(!t)throw new Error("bbox is required");if(!Array.isArray(t))throw new Error("bbox must be an Array");if(4!==t.length&&6!==t.length)throw new Error("bbox must be an Array of 4 or 6 numbers");t.forEach((function(t){if(!d(t))throw new Error("bbox must only contain numbers")}))},r.validateId=function(t){if(!t)throw new Error("id is required");if(-1===["string","number"].indexOf(typeof t))throw new Error("id must be a number or a string")}},{}],65:[function(t,e,r){"use strict";Object.defineProperty(r,"__esModule",{value:!0});var n=t("@turf/helpers");function i(t,e,r){if(null!==t)for(var n,a,o,s,l,c,u,f,h=0,p=0,d=t.type,m="FeatureCollection"===d,g="Feature"===d,v=m?t.features.length:1,y=0;y<v;y++){l=(f=!!(u=m?t.features[y].geometry:g?t.geometry:t)&&"GeometryCollection"===u.type)?u.geometries.length:1;for(var x=0;x<l;x++){var b=0,_=0;if(null!==(s=f?u.geometries[x]:u)){c=s.coordinates;var w=s.type;switch(h=!r||"Polygon"!==w&&"MultiPolygon"!==w?0:1,w){case null:break;case"Point":if(!1===e(c,p,y,b,_))return!1;p++,b++;break;case"LineString":case"MultiPoint":for(n=0;n<c.length;n++){if(!1===e(c[n],p,y,b,_))return!1;p++,"MultiPoint"===w&&b++}"LineString"===w&&b++;break;case"Polygon":case"MultiLineString":for(n=0;n<c.length;n++){for(a=0;a<c[n].length-h;a++){if(!1===e(c[n][a],p,y,b,_))return!1;p++}"MultiLineString"===w&&b++,"Polygon"===w&&_++}"Polygon"===w&&b++;break;case"MultiPolygon":for(n=0;n<c.length;n++){for(_=0,a=0;a<c[n].length;a++){for(o=0;o<c[n][a].length-h;o++){if(!1===e(c[n][a][o],p,y,b,_))return!1;p++}_++}b++}break;case"GeometryCollection":for(n=0;n<s.geometries.length;n++)if(!1===i(s.geometries[n],e,r))return!1;break;default:throw new Error("Unknown Geometry Type")}}}}}function a(t,e){var r;switch(t.type){case"FeatureCollection":for(r=0;r<t.features.length&&!1!==e(t.features[r].properties,r);r++);break;case"Feature":e(t.properties,0)}}function o(t,e){if("Feature"===t.type)e(t,0);else if("FeatureCollection"===t.type)for(var r=0;r<t.features.length&&!1!==e(t.features[r],r);r++);}function s(t,e){var r,n,i,a,o,s,l,c,u,f,h=0,p="FeatureCollection"===t.type,d="Feature"===t.type,m=p?t.features.length:1;for(r=0;r<m;r++){for(s=p?t.features[r].geometry:d?t.geometry:t,c=p?t.features[r].properties:d?t.properties:{},u=p?t.features[r].bbox:d?t.bbox:void 0,f=p?t.features[r].id:d?t.id:void 0,o=(l=!!s&&"GeometryCollection"===s.type)?s.geometries.length:1,i=0;i<o;i++)if(null!==(a=l?s.geometries[i]:s))switch(a.type){case"Point":case"LineString":case"MultiPoint":case"Polygon":case"MultiLineString":case"MultiPolygon":if(!1===e(a,h,c,u,f))return!1;break;case"GeometryCollection":for(n=0;n<a.geometries.length;n++)if(!1===e(a.geometries[n],h,c,u,f))return!1;break;default:throw new Error("Unknown Geometry Type")}else if(!1===e(null,h,c,u,f))return!1;h++}}function l(t,e){s(t,(function(t,r,i,a,o){var s,l=null===t?null:t.type;switch(l){case null:case"Point":case"LineString":case"Polygon":return!1!==e(n.feature(t,i,{bbox:a,id:o}),r,0)&&void 0}switch(l){case"MultiPoint":s="Point";break;case"MultiLineString":s="LineString";break;case"MultiPolygon":s="Polygon"}for(var c=0;c<t.coordinates.length;c++){var u={type:s,coordinates:t.coordinates[c]};if(!1===e(n.feature(u,i),r,c))return!1}}))}function c(t,e){l(t,(function(t,r,a){var o=0;if(t.geometry){var s=t.geometry.type;if("Point"!==s&&"MultiPoint"!==s){var l,c=0,u=0,f=0;return!1!==i(t,(function(i,s,h,p,d){if(void 0===l||r>c||p>u||d>f)return l=i,c=r,u=p,f=d,void(o=0);var m=n.lineString([l,i],t.properties);if(!1===e(m,r,a,d,o))return!1;o++,l=i}))&&void 0}}}))}function u(t,e){if(!t)throw new Error("geojson is required");l(t,(function(t,r,i){if(null!==t.geometry){var a=t.geometry.type,o=t.geometry.coordinates;switch(a){case"LineString":if(!1===e(t,r,i,0,0))return!1;break;case"Polygon":for(var s=0;s<o.length;s++)if(!1===e(n.lineString(o[s],t.properties),r,i,s))return!1}}}))}r.coordAll=function(t){var e=[];return i(t,(function(t){e.push(t)})),e},r.coordEach=i,r.coordReduce=function(t,e,r,n){var a=r;return i(t,(function(t,n,i,o,s){a=0===n&&void 0===r?t:e(a,t,n,i,o,s)}),n),a},r.featureEach=o,r.featureReduce=function(t,e,r){var n=r;return o(t,(function(t,i){n=0===i&&void 0===r?t:e(n,t,i)})),n},r.findPoint=function(t,e){if(e=e||{},!n.isObject(e))throw new Error("options is invalid");var r,i=e.featureIndex||0,a=e.multiFeatureIndex||0,o=e.geometryIndex||0,s=e.coordIndex||0,l=e.properties;switch(t.type){case"FeatureCollection":i<0&&(i=t.features.length+i),l=l||t.features[i].properties,r=t.features[i].geometry;break;case"Feature":l=l||t.properties,r=t.geometry;break;case"Point":case"MultiPoint":return null;case"LineString":case"Polygon":case"MultiLineString":case"MultiPolygon":r=t;break;default:throw new Error("geojson is invalid")}if(null===r)return null;var c=r.coordinates;switch(r.type){case"Point":return n.point(c,l,e);case"MultiPoint":return a<0&&(a=c.length+a),n.point(c[a],l,e);case"LineString":return s<0&&(s=c.length+s),n.point(c[s],l,e);case"Polygon":return o<0&&(o=c.length+o),s<0&&(s=c[o].length+s),n.point(c[o][s],l,e);case"MultiLineString":return a<0&&(a=c.length+a),s<0&&(s=c[a].length+s),n.point(c[a][s],l,e);case"MultiPolygon":return a<0&&(a=c.length+a),o<0&&(o=c[a].length+o),s<0&&(s=c[a][o].length-s),n.point(c[a][o][s],l,e)}throw new Error("geojson is invalid")},r.findSegment=function(t,e){if(e=e||{},!n.isObject(e))throw new Error("options is invalid");var r,i=e.featureIndex||0,a=e.multiFeatureIndex||0,o=e.geometryIndex||0,s=e.segmentIndex||0,l=e.properties;switch(t.type){case"FeatureCollection":i<0&&(i=t.features.length+i),l=l||t.features[i].properties,r=t.features[i].geometry;break;case"Feature":l=l||t.properties,r=t.geometry;break;case"Point":case"MultiPoint":return null;case"LineString":case"Polygon":case"MultiLineString":case"MultiPolygon":r=t;break;default:throw new Error("geojson is invalid")}if(null===r)return null;var c=r.coordinates;switch(r.type){case"Point":case"MultiPoint":return null;case"LineString":return s<0&&(s=c.length+s-1),n.lineString([c[s],c[s+1]],l,e);case"Polygon":return o<0&&(o=c.length+o),s<0&&(s=c[o].length+s-1),n.lineString([c[o][s],c[o][s+1]],l,e);case"MultiLineString":return a<0&&(a=c.length+a),s<0&&(s=c[a].length+s-1),n.lineString([c[a][s],c[a][s+1]],l,e);case"MultiPolygon":return a<0&&(a=c.length+a),o<0&&(o=c[a].length+o),s<0&&(s=c[a][o].length-s-1),n.lineString([c[a][o][s],c[a][o][s+1]],l,e)}throw new Error("geojson is invalid")},r.flattenEach=l,r.flattenReduce=function(t,e,r){var n=r;return l(t,(function(t,i,a){n=0===i&&0===a&&void 0===r?t:e(n,t,i,a)})),n},r.geomEach=s,r.geomReduce=function(t,e,r){var n=r;return s(t,(function(t,i,a,o,s){n=0===i&&void 0===r?t:e(n,t,i,a,o,s)})),n},r.lineEach=u,r.lineReduce=function(t,e,r){var n=r;return u(t,(function(t,i,a,o){n=0===i&&void 0===r?t:e(n,t,i,a,o)})),n},r.propEach=a,r.propReduce=function(t,e,r){var n=r;return a(t,(function(t,i){n=0===i&&void 0===r?t:e(n,t,i)})),n},r.segmentEach=c,r.segmentReduce=function(t,e,r){var n=r,i=!1;return c(t,(function(t,a,o,s,l){n=!1===i&&void 0===r?t:e(n,t,a,o,s,l),i=!0})),n}},{"@turf/helpers":64}],66:[function(t,e,r){"use strict";var n="undefined"==typeof WeakMap?t("weak-map"):WeakMap,i=t("gl-buffer"),a=t("gl-vao"),o=new n;e.exports=function(t){var e=o.get(t),r=e&&(e._triangleBuffer.handle||e._triangleBuffer.buffer);if(!r||!t.isBuffer(r)){var n=i(t,new Float32Array([-1,-1,-1,4,4,-1]));(e=a(t,[{buffer:n,type:t.FLOAT,size:2}]))._triangleBuffer=n,o.set(t,e)}e.bind(),t.drawArrays(t.TRIANGLES,0,3),e.unbind()}},{"gl-buffer":257,"gl-vao":343,"weak-map":598}],67:[function(t,e,r){e.exports=function(t){var e=0,r=0,n=0,i=0;return t.map((function(t){var a=(t=t.slice())[0],o=a.toUpperCase();if(a!=o)switch(t[0]=o,a){case"a":t[6]+=n,t[7]+=i;break;case"v":t[1]+=i;break;case"h":t[1]+=n;break;default:for(var s=1;s<t.length;)t[s++]+=n,t[s++]+=i}switch(o){case"Z":n=e,i=r;break;case"H":n=t[1];break;case"V":i=t[1];break;case"M":n=e=t[1],i=r=t[2];break;default:n=t[t.length-2],i=t[t.length-1]}return t}))}},{}],68:[function(t,e,r){var n=t("pad-left");e.exports=function(t,e,r){e="number"==typeof e?e:1,r=r||": ";var i=t.split(/\r?\n/),a=String(i.length+e-1).length;return i.map((function(t,i){var o=i+e,s=String(o).length;return n(o,a-s)+r+t})).join("\n")}},{"pad-left":469}],69:[function(t,e,r){"use strict";e.exports=function(t){var e=t.length;if(0===e)return[];if(1===e)return[0];for(var r=t[0].length,n=[t[0]],a=[0],o=1;o<e;++o)if(n.push(t[o]),i(n,r)){if(a.push(o),a.length===r+1)return a}else n.pop();return a};var n=t("robust-orientation");function i(t,e){for(var r=new Array(e+1),i=0;i<t.length;++i)r[i]=t[i];for(i=0;i<=t.length;++i){for(var a=t.length;a<=e;++a){for(var o=new Array(e),s=0;s<e;++s)o[s]=Math.pow(a+1-i,s);r[a]=o}if(n.apply(void 0,r))return!0}return!1}},{"robust-orientation":524}],70:[function(t,e,r){"use strict";e.exports=function(t,e){return n(e).filter((function(r){for(var n=new Array(r.length),a=0;a<r.length;++a)n[a]=e[r[a]];return i(n)*t<1}))};var n=t("delaunay-triangulate"),i=t("circumradius")},{circumradius:120,"delaunay-triangulate":172}],71:[function(t,e,r){e.exports=function(t,e){return i(n(t,e))};var n=t("alpha-complex"),i=t("simplicial-complex-boundary")},{"alpha-complex":70,"simplicial-complex-boundary":531}],72:[function(t,e,r){"use strict";e.exports=function(t,e){if(!t||null==t.length)throw Error("Argument should be an array");e=null==e?1:Math.floor(e);for(var r=Array(2*e),n=0;n<e;n++){for(var i=-1/0,a=1/0,o=n,s=t.length;o<s;o+=e)t[o]>i&&(i=t[o]),t[o]<a&&(a=t[o]);r[n]=a,r[e+n]=i}return r}},{}],73:[function(t,e,r){"use strict";e.exports=function(t,e,r){if("function"==typeof Array.prototype.findIndex)return t.findIndex(e,r);if("function"!=typeof e)throw new TypeError("predicate must be a function");var n=Object(t),i=n.length;if(0===i)return-1;for(var a=0;a<i;a++)if(e.call(r,n[a],a,n))return a;return-1}},{}],74:[function(t,e,r){"use strict";var n=t("array-bounds");e.exports=function(t,e,r){if(!t||null==t.length)throw Error("Argument should be an array");null==e&&(e=1);null==r&&(r=n(t,e));for(var i=0;i<e;i++){var a=r[e+i],o=r[i],s=i,l=t.length;if(a===1/0&&o===-1/0)for(s=i;s<l;s+=e)t[s]=t[s]===a?1:t[s]===o?0:.5;else if(a===1/0)for(s=i;s<l;s+=e)t[s]=t[s]===a?1:0;else if(o===-1/0)for(s=i;s<l;s+=e)t[s]=t[s]===o?0:1;else{var c=a-o;for(s=i;s<l;s+=e)isNaN(t[s])||(t[s]=0===c?.5:(t[s]-o)/c)}}return t}},{"array-bounds":72}],75:[function(t,e,r){e.exports=function(t,e){var r="number"==typeof t,n="number"==typeof e;r&&!n?(e=t,t=0):r||n||(t=0,e=0);var i=(e|=0)-(t|=0);if(i<0)throw new Error("array length must be positive");for(var a=new Array(i),o=0,s=t;o<i;o++,s++)a[o]=s;return a}},{}],76:[function(t,e,r){(function(r){(function(){"use strict";var n=t("object-assign");
/*!
 * The buffer module from node.js, for the browser.
 *
 * @author   Feross Aboukhadijeh <> <http://feross.org>
 * @license  MIT
 */function i(t,e){if(t===e)return 0;for(var r=t.length,n=e.length,i=0,a=Math.min(r,n);i<a;++i)if(t[i]!==e[i]){r=t[i],n=e[i];break}return r<n?-1:n<r?1:0}function a(t){return r.Buffer&&"function"==typeof r.Buffer.isBuffer?r.Buffer.isBuffer(t):!(null==t||!t._isBuffer)}var o=t("util/"),s=Object.prototype.hasOwnProperty,l=Array.prototype.slice,c="foo"===function(){}.name;function u(t){return Object.prototype.toString.call(t)}function f(t){return!a(t)&&("function"==typeof r.ArrayBuffer&&("function"==typeof ArrayBuffer.isView?ArrayBuffer.isView(t):!!t&&(t instanceof DataView||!!(t.buffer&&t.buffer instanceof ArrayBuffer))))}var h=e.exports=y,p=/\s*function\s+([^\(\s]*)\s*/;function d(t){if(o.isFunction(t)){if(c)return t.name;var e=t.toString().match(p);return e&&e[1]}}function m(t,e){return"string"==typeof t?t.length<e?t:t.slice(0,e):t}function g(t){if(c||!o.isFunction(t))return o.inspect(t);var e=d(t);return"[Function"+(e?": "+e:"")+"]"}function v(t,e,r,n,i){throw new h.AssertionError({message:r,actual:t,expected:e,operator:n,stackStartFunction:i})}function y(t,e){t||v(t,!0,e,"==",h.ok)}function x(t,e,r,n){if(t===e)return!0;if(a(t)&&a(e))return 0===i(t,e);if(o.isDate(t)&&o.isDate(e))return t.getTime()===e.getTime();if(o.isRegExp(t)&&o.isRegExp(e))return t.source===e.source&&t.global===e.global&&t.multiline===e.multiline&&t.lastIndex===e.lastIndex&&t.ignoreCase===e.ignoreCase;if(null!==t&&"object"==typeof t||null!==e&&"object"==typeof e){if(f(t)&&f(e)&&u(t)===u(e)&&!(t instanceof Float32Array||t instanceof Float64Array))return 0===i(new Uint8Array(t.buffer),new Uint8Array(e.buffer));if(a(t)!==a(e))return!1;var s=(n=n||{actual:[],expected:[]}).actual.indexOf(t);return-1!==s&&s===n.expected.indexOf(e)||(n.actual.push(t),n.expected.push(e),function(t,e,r,n){if(null==t||null==e)return!1;if(o.isPrimitive(t)||o.isPrimitive(e))return t===e;if(r&&Object.getPrototypeOf(t)!==Object.getPrototypeOf(e))return!1;var i=b(t),a=b(e);if(i&&!a||!i&&a)return!1;if(i)return t=l.call(t),e=l.call(e),x(t,e,r);var s,c,u=T(t),f=T(e);if(u.length!==f.length)return!1;for(u.sort(),f.sort(),c=u.length-1;c>=0;c--)if(u[c]!==f[c])return!1;for(c=u.length-1;c>=0;c--)if(s=u[c],!x(t[s],e[s],r,n))return!1;return!0}(t,e,r,n))}return r?t===e:t==e}function b(t){return"[object Arguments]"==Object.prototype.toString.call(t)}function _(t,e){if(!t||!e)return!1;if("[object RegExp]"==Object.prototype.toString.call(e))return e.test(t);try{if(t instanceof e)return!0}catch(t){}return!Error.isPrototypeOf(e)&&!0===e.call({},t)}function w(t,e,r,n){var i;if("function"!=typeof e)throw new TypeError('"block" argument must be a function');"string"==typeof r&&(n=r,r=null),i=function(t){var e;try{t()}catch(t){e=t}return e}(e),n=(r&&r.name?" ("+r.name+").":".")+(n?" "+n:"."),t&&!i&&v(i,r,"Missing expected exception"+n);var a="string"==typeof n,s=!t&&i&&!r;if((!t&&o.isError(i)&&a&&_(i,r)||s)&&v(i,r,"Got unwanted exception"+n),t&&i&&r&&!_(i,r)||!t&&i)throw i}h.AssertionError=function(t){this.name="AssertionError",this.actual=t.actual,this.expected=t.expected,this.operator=t.operator,t.message?(this.message=t.message,this.generatedMessage=!1):(this.message=function(t){return m(g(t.actual),128)+" "+t.operator+" "+m(g(t.expected),128)}(this),this.generatedMessage=!0);var e=t.stackStartFunction||v;if(Error.captureStackTrace)Error.captureStackTrace(this,e);else{var r=new Error;if(r.stack){var n=r.stack,i=d(e),a=n.indexOf("\n"+i);if(a>=0){var o=n.indexOf("\n",a+1);n=n.substring(o+1)}this.stack=n}}},o.inherits(h.AssertionError,Error),h.fail=v,h.ok=y,h.equal=function(t,e,r){t!=e&&v(t,e,r,"==",h.equal)},h.notEqual=function(t,e,r){t==e&&v(t,e,r,"!=",h.notEqual)},h.deepEqual=function(t,e,r){x(t,e,!1)||v(t,e,r,"deepEqual",h.deepEqual)},h.deepStrictEqual=function(t,e,r){x(t,e,!0)||v(t,e,r,"deepStrictEqual",h.deepStrictEqual)},h.notDeepEqual=function(t,e,r){x(t,e,!1)&&v(t,e,r,"notDeepEqual",h.notDeepEqual)},h.notDeepStrictEqual=function t(e,r,n){x(e,r,!0)&&v(e,r,n,"notDeepStrictEqual",t)},h.strictEqual=function(t,e,r){t!==e&&v(t,e,r,"===",h.strictEqual)},h.notStrictEqual=function(t,e,r){t===e&&v(t,e,r,"!==",h.notStrictEqual)},h.throws=function(t,e,r){w(!0,t,e,r)},h.doesNotThrow=function(t,e,r){w(!1,t,e,r)},h.ifError=function(t){if(t)throw t},h.strict=n((function t(e,r){e||v(e,!0,r,"==",t)}),h,{equal:h.strictEqual,deepEqual:h.deepStrictEqual,notEqual:h.notStrictEqual,notDeepEqual:h.notDeepStrictEqual}),h.strict.strict=h.strict;var T=Object.keys||function(t){var e=[];for(var r in t)s.call(t,r)&&e.push(r);return e}}).call(this)}).call(this,"undefined"!=typeof global?global:"undefined"!=typeof self?self:"undefined"!=typeof window?window:{})},{"object-assign":466,"util/":79}],77:[function(t,e,r){"function"==typeof Object.create?e.exports=function(t,e){t.super_=e,t.prototype=Object.create(e.prototype,{constructor:{value:t,enumerable:!1,writable:!0,configurable:!0}})}:e.exports=function(t,e){t.super_=e;var r=function(){};r.prototype=e.prototype,t.prototype=new r,t.prototype.constructor=t}},{}],78:[function(t,e,r){e.exports=function(t){return t&&"object"==typeof t&&"function"==typeof t.copy&&"function"==typeof t.fill&&"function"==typeof t.readUInt8}},{}],79:[function(t,e,r){(function(e,n){(function(){var i=/%[sdj%]/g;r.format=function(t){if(!v(t)){for(var e=[],r=0;r<arguments.length;r++)e.push(s(arguments[r]));return e.join(" ")}r=1;for(var n=arguments,a=n.length,o=String(t).replace(i,(function(t){if("%%"===t)return"%";if(r>=a)return t;switch(t){case"%s":return String(n[r++]);case"%d":return Number(n[r++]);case"%j":try{return JSON.stringify(n[r++])}catch(t){return"[Circular]"}default:return t}})),l=n[r];r<a;l=n[++r])m(l)||!b(l)?o+=" "+l:o+=" "+s(l);return o},r.deprecate=function(t,i){if(y(n.process))return function(){return r.deprecate(t,i).apply(this,arguments)};if(!0===e.noDeprecation)return t;var a=!1;return function(){if(!a){if(e.throwDeprecation)throw new Error(i);e.traceDeprecation?console.trace(i):console.error(i),a=!0}return t.apply(this,arguments)}};var a,o={};function s(t,e){var n={seen:[],stylize:c};return arguments.length>=3&&(n.depth=arguments[2]),arguments.length>=4&&(n.colors=arguments[3]),d(e)?n.showHidden=e:e&&r._extend(n,e),y(n.showHidden)&&(n.showHidden=!1),y(n.depth)&&(n.depth=2),y(n.colors)&&(n.colors=!1),y(n.customInspect)&&(n.customInspect=!0),n.colors&&(n.stylize=l),u(n,t,n.depth)}function l(t,e){var r=s.styles[e];return r?"\x1b["+s.colors[r][0]+"m"+t+"\x1b["+s.colors[r][1]+"m":t}function c(t,e){return t}function u(t,e,n){if(t.customInspect&&e&&T(e.inspect)&&e.inspect!==r.inspect&&(!e.constructor||e.constructor.prototype!==e)){var i=e.inspect(n,t);return v(i)||(i=u(t,i,n)),i}var a=function(t,e){if(y(e))return t.stylize("undefined","undefined");if(v(e)){var r="'"+JSON.stringify(e).replace(/^"|"$/g,"").replace(/'/g,"\\'").replace(/\\"/g,'"')+"'";return t.stylize(r,"string")}if(g(e))return t.stylize(""+e,"number");if(d(e))return t.stylize(""+e,"boolean");if(m(e))return t.stylize("null","null")}(t,e);if(a)return a;var o=Object.keys(e),s=function(t){var e={};return t.forEach((function(t,r){e[t]=!0})),e}(o);if(t.showHidden&&(o=Object.getOwnPropertyNames(e)),w(e)&&(o.indexOf("message")>=0||o.indexOf("description")>=0))return f(e);if(0===o.length){if(T(e)){var l=e.name?": "+e.name:"";return t.stylize("[Function"+l+"]","special")}if(x(e))return t.stylize(RegExp.prototype.toString.call(e),"regexp");if(_(e))return t.stylize(Date.prototype.toString.call(e),"date");if(w(e))return f(e)}var c,b="",k=!1,A=["{","}"];(p(e)&&(k=!0,A=["[","]"]),T(e))&&(b=" [Function"+(e.name?": "+e.name:"")+"]");return x(e)&&(b=" "+RegExp.prototype.toString.call(e)),_(e)&&(b=" "+Date.prototype.toUTCString.call(e)),w(e)&&(b=" "+f(e)),0!==o.length||k&&0!=e.length?n<0?x(e)?t.stylize(RegExp.prototype.toString.call(e),"regexp"):t.stylize("[Object]","special"):(t.seen.push(e),c=k?function(t,e,r,n,i){for(var a=[],o=0,s=e.length;o<s;++o)E(e,String(o))?a.push(h(t,e,r,n,String(o),!0)):a.push("");return i.forEach((function(i){i.match(/^\d+$/)||a.push(h(t,e,r,n,i,!0))})),a}(t,e,n,s,o):o.map((function(r){return h(t,e,n,s,r,k)})),t.seen.pop(),function(t,e,r){if(t.reduce((function(t,e){return e.indexOf("\n")>=0&&0,t+e.replace(/\u001b\[\d\d?m/g,"").length+1}),0)>60)return r[0]+(""===e?"":e+"\n ")+" "+t.join(",\n  ")+" "+r[1];return r[0]+e+" "+t.join(", ")+" "+r[1]}(c,b,A)):A[0]+b+A[1]}function f(t){return"["+Error.prototype.toString.call(t)+"]"}function h(t,e,r,n,i,a){var o,s,l;if((l=Object.getOwnPropertyDescriptor(e,i)||{value:e[i]}).get?s=l.set?t.stylize("[Getter/Setter]","special"):t.stylize("[Getter]","special"):l.set&&(s=t.stylize("[Setter]","special")),E(n,i)||(o="["+i+"]"),s||(t.seen.indexOf(l.value)<0?(s=m(r)?u(t,l.value,null):u(t,l.value,r-1)).indexOf("\n")>-1&&(s=a?s.split("\n").map((function(t){return"  "+t})).join("\n").substr(2):"\n"+s.split("\n").map((function(t){return"   "+t})).join("\n")):s=t.stylize("[Circular]","special")),y(o)){if(a&&i.match(/^\d+$/))return s;(o=JSON.stringify(""+i)).match(/^"([a-zA-Z_][a-zA-Z_0-9]*)"$/)?(o=o.substr(1,o.length-2),o=t.stylize(o,"name")):(o=o.replace(/'/g,"\\'").replace(/\\"/g,'"').replace(/(^"|"$)/g,"'"),o=t.stylize(o,"string"))}return o+": "+s}function p(t){return Array.isArray(t)}function d(t){return"boolean"==typeof t}function m(t){return null===t}function g(t){return"number"==typeof t}function v(t){return"string"==typeof t}function y(t){return void 0===t}function x(t){return b(t)&&"[object RegExp]"===k(t)}function b(t){return"object"==typeof t&&null!==t}function _(t){return b(t)&&"[object Date]"===k(t)}function w(t){return b(t)&&("[object Error]"===k(t)||t instanceof Error)}function T(t){return"function"==typeof t}function k(t){return Object.prototype.toString.call(t)}function A(t){return t<10?"0"+t.toString(10):t.toString(10)}r.debuglog=function(t){if(y(a)&&(a=e.env.NODE_DEBUG||""),t=t.toUpperCase(),!o[t])if(new RegExp("\\b"+t+"\\b","i").test(a)){var n=e.pid;o[t]=function(){var e=r.format.apply(r,arguments);console.error("%s %d: %s",t,n,e)}}else o[t]=function(){};return o[t]},r.inspect=s,s.colors={bold:[1,22],italic:[3,23],underline:[4,24],inverse:[7,27],white:[37,39],grey:[90,39],black:[30,39],blue:[34,39],cyan:[36,39],green:[32,39],magenta:[35,39],red:[31,39],yellow:[33,39]},s.styles={special:"cyan",number:"yellow",boolean:"yellow",undefined:"grey",null:"bold",string:"green",date:"magenta",regexp:"red"},r.isArray=p,r.isBoolean=d,r.isNull=m,r.isNullOrUndefined=function(t){return null==t},r.isNumber=g,r.isString=v,r.isSymbol=function(t){return"symbol"==typeof t},r.isUndefined=y,r.isRegExp=x,r.isObject=b,r.isDate=_,r.isError=w,r.isFunction=T,r.isPrimitive=function(t){return null===t||"boolean"==typeof t||"number"==typeof t||"string"==typeof t||"symbol"==typeof t||void 0===t},r.isBuffer=t("./support/isBuffer");var M=["Jan","Feb","Mar","Apr","May","Jun","Jul","Aug","Sep","Oct","Nov","Dec"];function S(){var t=new Date,e=[A(t.getHours()),A(t.getMinutes()),A(t.getSeconds())].join(":");return[t.getDate(),M[t.getMonth()],e].join(" ")}function E(t,e){return Object.prototype.hasOwnProperty.call(t,e)}r.log=function(){console.log("%s - %s",S(),r.format.apply(r,arguments))},r.inherits=t("inherits"),r._extend=function(t,e){if(!e||!b(e))return t;for(var r=Object.keys(e),n=r.length;n--;)t[r[n]]=e[r[n]];return t}}).call(this)}).call(this,t("_process"),"undefined"!=typeof global?global:"undefined"!=typeof self?self:"undefined"!=typeof window?window:{})},{"./support/isBuffer":78,_process:504,inherits:77}],80:[function(t,e,r){e.exports=function(t){return atob(t)}},{}],81:[function(t,e,r){"use strict";e.exports=function(t,e){for(var r=e.length,a=new Array(r+1),o=0;o<r;++o){for(var s=new Array(r+1),l=0;l<=r;++l)s[l]=t[l][o];a[o]=s}a[r]=new Array(r+1);for(o=0;o<=r;++o)a[r][o]=1;var c=new Array(r+1);for(o=0;o<r;++o)c[o]=e[o];c[r]=1;var u=n(a,c),f=i(u[r+1]);0===f&&(f=1);var h=new Array(r+1);for(o=0;o<=r;++o)h[o]=i(u[o])/f;return h};var n=t("robust-linear-solve");function i(t){for(var e=0,r=0;r<t.length;++r)e+=t[r];return e}},{"robust-linear-solve":523}],82:[function(t,e,r){"use strict";r.byteLength=function(t){var e=c(t),r=e[0],n=e[1];return 3*(r+n)/4-n},r.toByteArray=function(t){var e,r,n=c(t),o=n[0],s=n[1],l=new a(function(t,e,r){return 3*(e+r)/4-r}(0,o,s)),u=0,f=s>0?o-4:o;for(r=0;r<f;r+=4)e=i[t.charCodeAt(r)]<<18|i[t.charCodeAt(r+1)]<<12|i[t.charCodeAt(r+2)]<<6|i[t.charCodeAt(r+3)],l[u++]=e>>16&255,l[u++]=e>>8&255,l[u++]=255&e;2===s&&(e=i[t.charCodeAt(r)]<<2|i[t.charCodeAt(r+1)]>>4,l[u++]=255&e);1===s&&(e=i[t.charCodeAt(r)]<<10|i[t.charCodeAt(r+1)]<<4|i[t.charCodeAt(r+2)]>>2,l[u++]=e>>8&255,l[u++]=255&e);return l},r.fromByteArray=function(t){for(var e,r=t.length,i=r%3,a=[],o=0,s=r-i;o<s;o+=16383)a.push(u(t,o,o+16383>s?s:o+16383));1===i?(e=t[r-1],a.push(n[e>>2]+n[e<<4&63]+"==")):2===i&&(e=(t[r-2]<<8)+t[r-1],a.push(n[e>>10]+n[e>>4&63]+n[e<<2&63]+"="));return a.join("")};for(var n=[],i=[],a="undefined"!=typeof Uint8Array?Uint8Array:Array,o="ABCDEFGHIJKLMNOPQRSTUVWXYZabcdefghijklmnopqrstuvwxyz0123456789+/",s=0,l=o.length;s<l;++s)n[s]=o[s],i[o.charCodeAt(s)]=s;function c(t){var e=t.length;if(e%4>0)throw new Error("Invalid string. Length must be a multiple of 4");var r=t.indexOf("=");return-1===r&&(r=e),[r,r===e?0:4-r%4]}function u(t,e,r){for(var i,a,o=[],s=e;s<r;s+=3)i=(t[s]<<16&16711680)+(t[s+1]<<8&65280)+(255&t[s+2]),o.push(n[(a=i)>>18&63]+n[a>>12&63]+n[a>>6&63]+n[63&a]);return o.join("")}i["-".charCodeAt(0)]=62,i["_".charCodeAt(0)]=63},{}],83:[function(t,e,r){"use strict";var n=t("./lib/rationalize");e.exports=function(t,e){return n(t[0].mul(e[1]).add(e[0].mul(t[1])),t[1].mul(e[1]))}},{"./lib/rationalize":93}],84:[function(t,e,r){"use strict";e.exports=function(t,e){return t[0].mul(e[1]).cmp(e[0].mul(t[1]))}},{}],85:[function(t,e,r){"use strict";var n=t("./lib/rationalize");e.exports=function(t,e){return n(t[0].mul(e[1]),t[1].mul(e[0]))}},{"./lib/rationalize":93}],86:[function(t,e,r){"use strict";var n=t("./is-rat"),i=t("./lib/is-bn"),a=t("./lib/num-to-bn"),o=t("./lib/str-to-bn"),s=t("./lib/rationalize"),l=t("./div");e.exports=function t(e,r){if(n(e))return r?l(e,t(r)):[e[0].clone(),e[1].clone()];var c,u,f=0;if(i(e))c=e.clone();else if("string"==typeof e)c=o(e);else{if(0===e)return[a(0),a(1)];if(e===Math.floor(e))c=a(e);else{for(;e!==Math.floor(e);)e*=Math.pow(2,256),f-=256;c=a(e)}}if(n(r))c.mul(r[1]),u=r[0].clone();else if(i(r))u=r.clone();else if("string"==typeof r)u=o(r);else if(r)if(r===Math.floor(r))u=a(r);else{for(;r!==Math.floor(r);)r*=Math.pow(2,256),f+=256;u=a(r)}else u=a(1);f>0?c=c.ushln(f):f<0&&(u=u.ushln(-f));return s(c,u)}},{"./div":85,"./is-rat":87,"./lib/is-bn":91,"./lib/num-to-bn":92,"./lib/rationalize":93,"./lib/str-to-bn":94}],87:[function(t,e,r){"use strict";var n=t("./lib/is-bn");e.exports=function(t){return Array.isArray(t)&&2===t.length&&n(t[0])&&n(t[1])}},{"./lib/is-bn":91}],88:[function(t,e,r){"use strict";var n=t("bn.js");e.exports=function(t){return t.cmp(new n(0))}},{"bn.js":96}],89:[function(t,e,r){"use strict";var n=t("./bn-sign");e.exports=function(t){var e=t.length,r=t.words,i=0;if(1===e)i=r[0];else if(2===e)i=r[0]+67108864*r[1];else for(var a=0;a<e;a++){var o=r[a];i+=o*Math.pow(67108864,a)}return n(t)*i}},{"./bn-sign":88}],90:[function(t,e,r){"use strict";var n=t("double-bits"),i=t("bit-twiddle").countTrailingZeros;e.exports=function(t){var e=i(n.lo(t));if(e<32)return e;var r=i(n.hi(t));if(r>20)return 52;return r+32}},{"bit-twiddle":101,"double-bits":174}],91:[function(t,e,r){"use strict";t("bn.js");e.exports=function(t){return t&&"object"==typeof t&&Boolean(t.words)}},{"bn.js":96}],92:[function(t,e,r){"use strict";var n=t("bn.js"),i=t("double-bits");e.exports=function(t){var e=i.exponent(t);return e<52?new n(t):new n(t*Math.pow(2,52-e)).ushln(e-52)}},{"bn.js":96,"double-bits":174}],93:[function(t,e,r){"use strict";var n=t("./num-to-bn"),i=t("./bn-sign");e.exports=function(t,e){var r=i(t),a=i(e);if(0===r)return[n(0),n(1)];if(0===a)return[n(0),n(0)];a<0&&(t=t.neg(),e=e.neg());var o=t.gcd(e);if(o.cmpn(1))return[t.div(o),e.div(o)];return[t,e]}},{"./bn-sign":88,"./num-to-bn":92}],94:[function(t,e,r){"use strict";var n=t("bn.js");e.exports=function(t){return new n(t)}},{"bn.js":96}],95:[function(t,e,r){"use strict";var n=t("./lib/rationalize");e.exports=function(t,e){return n(t[0].mul(e[0]),t[1].mul(e[1]))}},{"./lib/rationalize":93}],96:[function(t,e,r){!function(e,r){"use strict";function n(t,e){if(!t)throw new Error(e||"Assertion failed")}function i(t,e){t.super_=e;var r=function(){};r.prototype=e.prototype,t.prototype=new r,t.prototype.constructor=t}function a(t,e,r){if(a.isBN(t))return t;this.negative=0,this.words=null,this.length=0,this.red=null,null!==t&&("le"!==e&&"be"!==e||(r=e,e=10),this._init(t||0,e||10,r||"be"))}var o;"object"==typeof e?e.exports=a:r.BN=a,a.BN=a,a.wordSize=26;try{o="undefined"!=typeof window&&void 0!==window.Buffer?window.Buffer:t("buffer").Buffer}catch(t){}function s(t,e){var r=t.charCodeAt(e);return r>=65&&r<=70?r-55:r>=97&&r<=102?r-87:r-48&15}function l(t,e,r){var n=s(t,r);return r-1>=e&&(n|=s(t,r-1)<<4),n}function c(t,e,r,n){for(var i=0,a=Math.min(t.length,r),o=e;o<a;o++){var s=t.charCodeAt(o)-48;i*=n,i+=s>=49?s-49+10:s>=17?s-17+10:s}return i}a.isBN=function(t){return t instanceof a||null!==t&&"object"==typeof t&&t.constructor.wordSize===a.wordSize&&Array.isArray(t.words)},a.max=function(t,e){return t.cmp(e)>0?t:e},a.min=function(t,e){return t.cmp(e)<0?t:e},a.prototype._init=function(t,e,r){if("number"==typeof t)return this._initNumber(t,e,r);if("object"==typeof t)return this._initArray(t,e,r);"hex"===e&&(e=16),n(e===(0|e)&&e>=2&&e<=36);var i=0;"-"===(t=t.toString().replace(/\s+/g,""))[0]&&(i++,this.negative=1),i<t.length&&(16===e?this._parseHex(t,i,r):(this._parseBase(t,e,i),"le"===r&&this._initArray(this.toArray(),e,r)))},a.prototype._initNumber=function(t,e,r){t<0&&(this.negative=1,t=-t),t<67108864?(this.words=[67108863&t],this.length=1):t<4503599627370496?(this.words=[67108863&t,t/67108864&67108863],this.length=2):(n(t<9007199254740992),this.words=[67108863&t,t/67108864&67108863,1],this.length=3),"le"===r&&this._initArray(this.toArray(),e,r)},a.prototype._initArray=function(t,e,r){if(n("number"==typeof t.length),t.length<=0)return this.words=[0],this.length=1,this;this.length=Math.ceil(t.length/3),this.words=new Array(this.length);for(var i=0;i<this.length;i++)this.words[i]=0;var a,o,s=0;if("be"===r)for(i=t.length-1,a=0;i>=0;i-=3)o=t[i]|t[i-1]<<8|t[i-2]<<16,this.words[a]|=o<<s&67108863,this.words[a+1]=o>>>26-s&67108863,(s+=24)>=26&&(s-=26,a++);else if("le"===r)for(i=0,a=0;i<t.length;i+=3)o=t[i]|t[i+1]<<8|t[i+2]<<16,this.words[a]|=o<<s&67108863,this.words[a+1]=o>>>26-s&67108863,(s+=24)>=26&&(s-=26,a++);return this.strip()},a.prototype._parseHex=function(t,e,r){this.length=Math.ceil((t.length-e)/6),this.words=new Array(this.length);for(var n=0;n<this.length;n++)this.words[n]=0;var i,a=0,o=0;if("be"===r)for(n=t.length-1;n>=e;n-=2)i=l(t,e,n)<<a,this.words[o]|=67108863&i,a>=18?(a-=18,o+=1,this.words[o]|=i>>>26):a+=8;else for(n=(t.length-e)%2==0?e+1:e;n<t.length;n+=2)i=l(t,e,n)<<a,this.words[o]|=67108863&i,a>=18?(a-=18,o+=1,this.words[o]|=i>>>26):a+=8;this.strip()},a.prototype._parseBase=function(t,e,r){this.words=[0],this.length=1;for(var n=0,i=1;i<=67108863;i*=e)n++;n--,i=i/e|0;for(var a=t.length-r,o=a%n,s=Math.min(a,a-o)+r,l=0,u=r;u<s;u+=n)l=c(t,u,u+n,e),this.imuln(i),this.words[0]+l<67108864?this.words[0]+=l:this._iaddn(l);if(0!==o){var f=1;for(l=c(t,u,t.length,e),u=0;u<o;u++)f*=e;this.imuln(f),this.words[0]+l<67108864?this.words[0]+=l:this._iaddn(l)}this.strip()},a.prototype.copy=function(t){t.words=new Array(this.length);for(var e=0;e<this.length;e++)t.words[e]=this.words[e];t.length=this.length,t.negative=this.negative,t.red=this.red},a.prototype.clone=function(){var t=new a(null);return this.copy(t),t},a.prototype._expand=function(t){for(;this.length<t;)this.words[this.length++]=0;return this},a.prototype.strip=function(){for(;this.length>1&&0===this.words[this.length-1];)this.length--;return this._normSign()},a.prototype._normSign=function(){return 1===this.length&&0===this.words[0]&&(this.negative=0),this},a.prototype.inspect=function(){return(this.red?"<BN-R: ":"<BN: ")+this.toString(16)+">"};var u=["","0","00","000","0000","00000","000000","0000000","00000000","000000000","0000000000","00000000000","000000000000","0000000000000","00000000000000","000000000000000","0000000000000000","00000000000000000","000000000000000000","0000000000000000000","00000000000000000000","000000000000000000000","0000000000000000000000","00000000000000000000000","000000000000000000000000","0000000000000000000000000"],f=[0,0,25,16,12,11,10,9,8,8,7,7,7,7,6,6,6,6,6,6,6,5,5,5,5,5,5,5,5,5,5,5,5,5,5,5,5],h=[0,0,33554432,43046721,16777216,48828125,60466176,40353607,16777216,43046721,1e7,19487171,35831808,62748517,7529536,11390625,16777216,24137569,34012224,47045881,64e6,4084101,5153632,6436343,7962624,9765625,11881376,14348907,17210368,20511149,243e5,28629151,33554432,39135393,45435424,52521875,60466176];function p(t,e,r){r.negative=e.negative^t.negative;var n=t.length+e.length|0;r.length=n,n=n-1|0;var i=0|t.words[0],a=0|e.words[0],o=i*a,s=67108863&o,l=o/67108864|0;r.words[0]=s;for(var c=1;c<n;c++){for(var u=l>>>26,f=67108863&l,h=Math.min(c,e.length-1),p=Math.max(0,c-t.length+1);p<=h;p++){var d=c-p|0;u+=(o=(i=0|t.words[d])*(a=0|e.words[p])+f)/67108864|0,f=67108863&o}r.words[c]=0|f,l=0|u}return 0!==l?r.words[c]=0|l:r.length--,r.strip()}a.prototype.toString=function(t,e){var r;if(e=0|e||1,16===(t=t||10)||"hex"===t){r="";for(var i=0,a=0,o=0;o<this.length;o++){var s=this.words[o],l=(16777215&(s<<i|a)).toString(16);r=0!==(a=s>>>24-i&16777215)||o!==this.length-1?u[6-l.length]+l+r:l+r,(i+=2)>=26&&(i-=26,o--)}for(0!==a&&(r=a.toString(16)+r);r.length%e!=0;)r="0"+r;return 0!==this.negative&&(r="-"+r),r}if(t===(0|t)&&t>=2&&t<=36){var c=f[t],p=h[t];r="";var d=this.clone();for(d.negative=0;!d.isZero();){var m=d.modn(p).toString(t);r=(d=d.idivn(p)).isZero()?m+r:u[c-m.length]+m+r}for(this.isZero()&&(r="0"+r);r.length%e!=0;)r="0"+r;return 0!==this.negative&&(r="-"+r),r}n(!1,"Base should be between 2 and 36")},a.prototype.toNumber=function(){var t=this.words[0];return 2===this.length?t+=67108864*this.words[1]:3===this.length&&1===this.words[2]?t+=4503599627370496+67108864*this.words[1]:this.length>2&&n(!1,"Number can only safely store up to 53 bits"),0!==this.negative?-t:t},a.prototype.toJSON=function(){return this.toString(16)},a.prototype.toBuffer=function(t,e){return n(void 0!==o),this.toArrayLike(o,t,e)},a.prototype.toArray=function(t,e){return this.toArrayLike(Array,t,e)},a.prototype.toArrayLike=function(t,e,r){var i=this.byteLength(),a=r||Math.max(1,i);n(i<=a,"byte array longer than desired length"),n(a>0,"Requested array length <= 0"),this.strip();var o,s,l="le"===e,c=new t(a),u=this.clone();if(l){for(s=0;!u.isZero();s++)o=u.andln(255),u.iushrn(8),c[s]=o;for(;s<a;s++)c[s]=0}else{for(s=0;s<a-i;s++)c[s]=0;for(s=0;!u.isZero();s++)o=u.andln(255),u.iushrn(8),c[a-s-1]=o}return c},Math.clz32?a.prototype._countBits=function(t){return 32-Math.clz32(t)}:a.prototype._countBits=function(t){var e=t,r=0;return e>=4096&&(r+=13,e>>>=13),e>=64&&(r+=7,e>>>=7),e>=8&&(r+=4,e>>>=4),e>=2&&(r+=2,e>>>=2),r+e},a.prototype._zeroBits=function(t){if(0===t)return 26;var e=t,r=0;return 0==(8191&e)&&(r+=13,e>>>=13),0==(127&e)&&(r+=7,e>>>=7),0==(15&e)&&(r+=4,e>>>=4),0==(3&e)&&(r+=2,e>>>=2),0==(1&e)&&r++,r},a.prototype.bitLength=function(){var t=this.words[this.length-1],e=this._countBits(t);return 26*(this.length-1)+e},a.prototype.zeroBits=function(){if(this.isZero())return 0;for(var t=0,e=0;e<this.length;e++){var r=this._zeroBits(this.words[e]);if(t+=r,26!==r)break}return t},a.prototype.byteLength=function(){return Math.ceil(this.bitLength()/8)},a.prototype.toTwos=function(t){return 0!==this.negative?this.abs().inotn(t).iaddn(1):this.clone()},a.prototype.fromTwos=function(t){return this.testn(t-1)?this.notn(t).iaddn(1).ineg():this.clone()},a.prototype.isNeg=function(){return 0!==this.negative},a.prototype.neg=function(){return this.clone().ineg()},a.prototype.ineg=function(){return this.isZero()||(this.negative^=1),this},a.prototype.iuor=function(t){for(;this.length<t.length;)this.words[this.length++]=0;for(var e=0;e<t.length;e++)this.words[e]=this.words[e]|t.words[e];return this.strip()},a.prototype.ior=function(t){return n(0==(this.negative|t.negative)),this.iuor(t)},a.prototype.or=function(t){return this.length>t.length?this.clone().ior(t):t.clone().ior(this)},a.prototype.uor=function(t){return this.length>t.length?this.clone().iuor(t):t.clone().iuor(this)},a.prototype.iuand=function(t){var e;e=this.length>t.length?t:this;for(var r=0;r<e.length;r++)this.words[r]=this.words[r]&t.words[r];return this.length=e.length,this.strip()},a.prototype.iand=function(t){return n(0==(this.negative|t.negative)),this.iuand(t)},a.prototype.and=function(t){return this.length>t.length?this.clone().iand(t):t.clone().iand(this)},a.prototype.uand=function(t){return this.length>t.length?this.clone().iuand(t):t.clone().iuand(this)},a.prototype.iuxor=function(t){var e,r;this.length>t.length?(e=this,r=t):(e=t,r=this);for(var n=0;n<r.length;n++)this.words[n]=e.words[n]^r.words[n];if(this!==e)for(;n<e.length;n++)this.words[n]=e.words[n];return this.length=e.length,this.strip()},a.prototype.ixor=function(t){return n(0==(this.negative|t.negative)),this.iuxor(t)},a.prototype.xor=function(t){return this.length>t.length?this.clone().ixor(t):t.clone().ixor(this)},a.prototype.uxor=function(t){return this.length>t.length?this.clone().iuxor(t):t.clone().iuxor(this)},a.prototype.inotn=function(t){n("number"==typeof t&&t>=0);var e=0|Math.ceil(t/26),r=t%26;this._expand(e),r>0&&e--;for(var i=0;i<e;i++)this.words[i]=67108863&~this.words[i];return r>0&&(this.words[i]=~this.words[i]&67108863>>26-r),this.strip()},a.prototype.notn=function(t){return this.clone().inotn(t)},a.prototype.setn=function(t,e){n("number"==typeof t&&t>=0);var r=t/26|0,i=t%26;return this._expand(r+1),this.words[r]=e?this.words[r]|1<<i:this.words[r]&~(1<<i),this.strip()},a.prototype.iadd=function(t){var e,r,n;if(0!==this.negative&&0===t.negative)return this.negative=0,e=this.isub(t),this.negative^=1,this._normSign();if(0===this.negative&&0!==t.negative)return t.negative=0,e=this.isub(t),t.negative=1,e._normSign();this.length>t.length?(r=this,n=t):(r=t,n=this);for(var i=0,a=0;a<n.length;a++)e=(0|r.words[a])+(0|n.words[a])+i,this.words[a]=67108863&e,i=e>>>26;for(;0!==i&&a<r.length;a++)e=(0|r.words[a])+i,this.words[a]=67108863&e,i=e>>>26;if(this.length=r.length,0!==i)this.words[this.length]=i,this.length++;else if(r!==this)for(;a<r.length;a++)this.words[a]=r.words[a];return this},a.prototype.add=function(t){var e;return 0!==t.negative&&0===this.negative?(t.negative=0,e=this.sub(t),t.negative^=1,e):0===t.negative&&0!==this.negative?(this.negative=0,e=t.sub(this),this.negative=1,e):this.length>t.length?this.clone().iadd(t):t.clone().iadd(this)},a.prototype.isub=function(t){if(0!==t.negative){t.negative=0;var e=this.iadd(t);return t.negative=1,e._normSign()}if(0!==this.negative)return this.negative=0,this.iadd(t),this.negative=1,this._normSign();var r,n,i=this.cmp(t);if(0===i)return this.negative=0,this.length=1,this.words[0]=0,this;i>0?(r=this,n=t):(r=t,n=this);for(var a=0,o=0;o<n.length;o++)a=(e=(0|r.words[o])-(0|n.words[o])+a)>>26,this.words[o]=67108863&e;for(;0!==a&&o<r.length;o++)a=(e=(0|r.words[o])+a)>>26,this.words[o]=67108863&e;if(0===a&&o<r.length&&r!==this)for(;o<r.length;o++)this.words[o]=r.words[o];return this.length=Math.max(this.length,o),r!==this&&(this.negative=1),this.strip()},a.prototype.sub=function(t){return this.clone().isub(t)};var d=function(t,e,r){var n,i,a,o=t.words,s=e.words,l=r.words,c=0,u=0|o[0],f=8191&u,h=u>>>13,p=0|o[1],d=8191&p,m=p>>>13,g=0|o[2],v=8191&g,y=g>>>13,x=0|o[3],b=8191&x,_=x>>>13,w=0|o[4],T=8191&w,k=w>>>13,A=0|o[5],M=8191&A,S=A>>>13,E=0|o[6],L=8191&E,C=E>>>13,P=0|o[7],I=8191&P,O=P>>>13,z=0|o[8],D=8191&z,R=z>>>13,F=0|o[9],B=8191&F,N=F>>>13,j=0|s[0],U=8191&j,V=j>>>13,H=0|s[1],q=8191&H,G=H>>>13,Y=0|s[2],W=8191&Y,X=Y>>>13,Z=0|s[3],J=8191&Z,K=Z>>>13,Q=0|s[4],$=8191&Q,tt=Q>>>13,et=0|s[5],rt=8191&et,nt=et>>>13,it=0|s[6],at=8191&it,ot=it>>>13,st=0|s[7],lt=8191&st,ct=st>>>13,ut=0|s[8],ft=8191&ut,ht=ut>>>13,pt=0|s[9],dt=8191&pt,mt=pt>>>13;r.negative=t.negative^e.negative,r.length=19;var gt=(c+(n=Math.imul(f,U))|0)+((8191&(i=(i=Math.imul(f,V))+Math.imul(h,U)|0))<<13)|0;c=((a=Math.imul(h,V))+(i>>>13)|0)+(gt>>>26)|0,gt&=67108863,n=Math.imul(d,U),i=(i=Math.imul(d,V))+Math.imul(m,U)|0,a=Math.imul(m,V);var vt=(c+(n=n+Math.imul(f,q)|0)|0)+((8191&(i=(i=i+Math.imul(f,G)|0)+Math.imul(h,q)|0))<<13)|0;c=((a=a+Math.imul(h,G)|0)+(i>>>13)|0)+(vt>>>26)|0,vt&=67108863,n=Math.imul(v,U),i=(i=Math.imul(v,V))+Math.imul(y,U)|0,a=Math.imul(y,V),n=n+Math.imul(d,q)|0,i=(i=i+Math.imul(d,G)|0)+Math.imul(m,q)|0,a=a+Math.imul(m,G)|0;var yt=(c+(n=n+Math.imul(f,W)|0)|0)+((8191&(i=(i=i+Math.imul(f,X)|0)+Math.imul(h,W)|0))<<13)|0;c=((a=a+Math.imul(h,X)|0)+(i>>>13)|0)+(yt>>>26)|0,yt&=67108863,n=Math.imul(b,U),i=(i=Math.imul(b,V))+Math.imul(_,U)|0,a=Math.imul(_,V),n=n+Math.imul(v,q)|0,i=(i=i+Math.imul(v,G)|0)+Math.imul(y,q)|0,a=a+Math.imul(y,G)|0,n=n+Math.imul(d,W)|0,i=(i=i+Math.imul(d,X)|0)+Math.imul(m,W)|0,a=a+Math.imul(m,X)|0;var xt=(c+(n=n+Math.imul(f,J)|0)|0)+((8191&(i=(i=i+Math.imul(f,K)|0)+Math.imul(h,J)|0))<<13)|0;c=((a=a+Math.imul(h,K)|0)+(i>>>13)|0)+(xt>>>26)|0,xt&=67108863,n=Math.imul(T,U),i=(i=Math.imul(T,V))+Math.imul(k,U)|0,a=Math.imul(k,V),n=n+Math.imul(b,q)|0,i=(i=i+Math.imul(b,G)|0)+Math.imul(_,q)|0,a=a+Math.imul(_,G)|0,n=n+Math.imul(v,W)|0,i=(i=i+Math.imul(v,X)|0)+Math.imul(y,W)|0,a=a+Math.imul(y,X)|0,n=n+Math.imul(d,J)|0,i=(i=i+Math.imul(d,K)|0)+Math.imul(m,J)|0,a=a+Math.imul(m,K)|0;var bt=(c+(n=n+Math.imul(f,$)|0)|0)+((8191&(i=(i=i+Math.imul(f,tt)|0)+Math.imul(h,$)|0))<<13)|0;c=((a=a+Math.imul(h,tt)|0)+(i>>>13)|0)+(bt>>>26)|0,bt&=67108863,n=Math.imul(M,U),i=(i=Math.imul(M,V))+Math.imul(S,U)|0,a=Math.imul(S,V),n=n+Math.imul(T,q)|0,i=(i=i+Math.imul(T,G)|0)+Math.imul(k,q)|0,a=a+Math.imul(k,G)|0,n=n+Math.imul(b,W)|0,i=(i=i+Math.imul(b,X)|0)+Math.imul(_,W)|0,a=a+Math.imul(_,X)|0,n=n+Math.imul(v,J)|0,i=(i=i+Math.imul(v,K)|0)+Math.imul(y,J)|0,a=a+Math.imul(y,K)|0,n=n+Math.imul(d,$)|0,i=(i=i+Math.imul(d,tt)|0)+Math.imul(m,$)|0,a=a+Math.imul(m,tt)|0;var _t=(c+(n=n+Math.imul(f,rt)|0)|0)+((8191&(i=(i=i+Math.imul(f,nt)|0)+Math.imul(h,rt)|0))<<13)|0;c=((a=a+Math.imul(h,nt)|0)+(i>>>13)|0)+(_t>>>26)|0,_t&=67108863,n=Math.imul(L,U),i=(i=Math.imul(L,V))+Math.imul(C,U)|0,a=Math.imul(C,V),n=n+Math.imul(M,q)|0,i=(i=i+Math.imul(M,G)|0)+Math.imul(S,q)|0,a=a+Math.imul(S,G)|0,n=n+Math.imul(T,W)|0,i=(i=i+Math.imul(T,X)|0)+Math.imul(k,W)|0,a=a+Math.imul(k,X)|0,n=n+Math.imul(b,J)|0,i=(i=i+Math.imul(b,K)|0)+Math.imul(_,J)|0,a=a+Math.imul(_,K)|0,n=n+Math.imul(v,$)|0,i=(i=i+Math.imul(v,tt)|0)+Math.imul(y,$)|0,a=a+Math.imul(y,tt)|0,n=n+Math.imul(d,rt)|0,i=(i=i+Math.imul(d,nt)|0)+Math.imul(m,rt)|0,a=a+Math.imul(m,nt)|0;var wt=(c+(n=n+Math.imul(f,at)|0)|0)+((8191&(i=(i=i+Math.imul(f,ot)|0)+Math.imul(h,at)|0))<<13)|0;c=((a=a+Math.imul(h,ot)|0)+(i>>>13)|0)+(wt>>>26)|0,wt&=67108863,n=Math.imul(I,U),i=(i=Math.imul(I,V))+Math.imul(O,U)|0,a=Math.imul(O,V),n=n+Math.imul(L,q)|0,i=(i=i+Math.imul(L,G)|0)+Math.imul(C,q)|0,a=a+Math.imul(C,G)|0,n=n+Math.imul(M,W)|0,i=(i=i+Math.imul(M,X)|0)+Math.imul(S,W)|0,a=a+Math.imul(S,X)|0,n=n+Math.imul(T,J)|0,i=(i=i+Math.imul(T,K)|0)+Math.imul(k,J)|0,a=a+Math.imul(k,K)|0,n=n+Math.imul(b,$)|0,i=(i=i+Math.imul(b,tt)|0)+Math.imul(_,$)|0,a=a+Math.imul(_,tt)|0,n=n+Math.imul(v,rt)|0,i=(i=i+Math.imul(v,nt)|0)+Math.imul(y,rt)|0,a=a+Math.imul(y,nt)|0,n=n+Math.imul(d,at)|0,i=(i=i+Math.imul(d,ot)|0)+Math.imul(m,at)|0,a=a+Math.imul(m,ot)|0;var Tt=(c+(n=n+Math.imul(f,lt)|0)|0)+((8191&(i=(i=i+Math.imul(f,ct)|0)+Math.imul(h,lt)|0))<<13)|0;c=((a=a+Math.imul(h,ct)|0)+(i>>>13)|0)+(Tt>>>26)|0,Tt&=67108863,n=Math.imul(D,U),i=(i=Math.imul(D,V))+Math.imul(R,U)|0,a=Math.imul(R,V),n=n+Math.imul(I,q)|0,i=(i=i+Math.imul(I,G)|0)+Math.imul(O,q)|0,a=a+Math.imul(O,G)|0,n=n+Math.imul(L,W)|0,i=(i=i+Math.imul(L,X)|0)+Math.imul(C,W)|0,a=a+Math.imul(C,X)|0,n=n+Math.imul(M,J)|0,i=(i=i+Math.imul(M,K)|0)+Math.imul(S,J)|0,a=a+Math.imul(S,K)|0,n=n+Math.imul(T,$)|0,i=(i=i+Math.imul(T,tt)|0)+Math.imul(k,$)|0,a=a+Math.imul(k,tt)|0,n=n+Math.imul(b,rt)|0,i=(i=i+Math.imul(b,nt)|0)+Math.imul(_,rt)|0,a=a+Math.imul(_,nt)|0,n=n+Math.imul(v,at)|0,i=(i=i+Math.imul(v,ot)|0)+Math.imul(y,at)|0,a=a+Math.imul(y,ot)|0,n=n+Math.imul(d,lt)|0,i=(i=i+Math.imul(d,ct)|0)+Math.imul(m,lt)|0,a=a+Math.imul(m,ct)|0;var kt=(c+(n=n+Math.imul(f,ft)|0)|0)+((8191&(i=(i=i+Math.imul(f,ht)|0)+Math.imul(h,ft)|0))<<13)|0;c=((a=a+Math.imul(h,ht)|0)+(i>>>13)|0)+(kt>>>26)|0,kt&=67108863,n=Math.imul(B,U),i=(i=Math.imul(B,V))+Math.imul(N,U)|0,a=Math.imul(N,V),n=n+Math.imul(D,q)|0,i=(i=i+Math.imul(D,G)|0)+Math.imul(R,q)|0,a=a+Math.imul(R,G)|0,n=n+Math.imul(I,W)|0,i=(i=i+Math.imul(I,X)|0)+Math.imul(O,W)|0,a=a+Math.imul(O,X)|0,n=n+Math.imul(L,J)|0,i=(i=i+Math.imul(L,K)|0)+Math.imul(C,J)|0,a=a+Math.imul(C,K)|0,n=n+Math.imul(M,$)|0,i=(i=i+Math.imul(M,tt)|0)+Math.imul(S,$)|0,a=a+Math.imul(S,tt)|0,n=n+Math.imul(T,rt)|0,i=(i=i+Math.imul(T,nt)|0)+Math.imul(k,rt)|0,a=a+Math.imul(k,nt)|0,n=n+Math.imul(b,at)|0,i=(i=i+Math.imul(b,ot)|0)+Math.imul(_,at)|0,a=a+Math.imul(_,ot)|0,n=n+Math.imul(v,lt)|0,i=(i=i+Math.imul(v,ct)|0)+Math.imul(y,lt)|0,a=a+Math.imul(y,ct)|0,n=n+Math.imul(d,ft)|0,i=(i=i+Math.imul(d,ht)|0)+Math.imul(m,ft)|0,a=a+Math.imul(m,ht)|0;var At=(c+(n=n+Math.imul(f,dt)|0)|0)+((8191&(i=(i=i+Math.imul(f,mt)|0)+Math.imul(h,dt)|0))<<13)|0;c=((a=a+Math.imul(h,mt)|0)+(i>>>13)|0)+(At>>>26)|0,At&=67108863,n=Math.imul(B,q),i=(i=Math.imul(B,G))+Math.imul(N,q)|0,a=Math.imul(N,G),n=n+Math.imul(D,W)|0,i=(i=i+Math.imul(D,X)|0)+Math.imul(R,W)|0,a=a+Math.imul(R,X)|0,n=n+Math.imul(I,J)|0,i=(i=i+Math.imul(I,K)|0)+Math.imul(O,J)|0,a=a+Math.imul(O,K)|0,n=n+Math.imul(L,$)|0,i=(i=i+Math.imul(L,tt)|0)+Math.imul(C,$)|0,a=a+Math.imul(C,tt)|0,n=n+Math.imul(M,rt)|0,i=(i=i+Math.imul(M,nt)|0)+Math.imul(S,rt)|0,a=a+Math.imul(S,nt)|0,n=n+Math.imul(T,at)|0,i=(i=i+Math.imul(T,ot)|0)+Math.imul(k,at)|0,a=a+Math.imul(k,ot)|0,n=n+Math.imul(b,lt)|0,i=(i=i+Math.imul(b,ct)|0)+Math.imul(_,lt)|0,a=a+Math.imul(_,ct)|0,n=n+Math.imul(v,ft)|0,i=(i=i+Math.imul(v,ht)|0)+Math.imul(y,ft)|0,a=a+Math.imul(y,ht)|0;var Mt=(c+(n=n+Math.imul(d,dt)|0)|0)+((8191&(i=(i=i+Math.imul(d,mt)|0)+Math.imul(m,dt)|0))<<13)|0;c=((a=a+Math.imul(m,mt)|0)+(i>>>13)|0)+(Mt>>>26)|0,Mt&=67108863,n=Math.imul(B,W),i=(i=Math.imul(B,X))+Math.imul(N,W)|0,a=Math.imul(N,X),n=n+Math.imul(D,J)|0,i=(i=i+Math.imul(D,K)|0)+Math.imul(R,J)|0,a=a+Math.imul(R,K)|0,n=n+Math.imul(I,$)|0,i=(i=i+Math.imul(I,tt)|0)+Math.imul(O,$)|0,a=a+Math.imul(O,tt)|0,n=n+Math.imul(L,rt)|0,i=(i=i+Math.imul(L,nt)|0)+Math.imul(C,rt)|0,a=a+Math.imul(C,nt)|0,n=n+Math.imul(M,at)|0,i=(i=i+Math.imul(M,ot)|0)+Math.imul(S,at)|0,a=a+Math.imul(S,ot)|0,n=n+Math.imul(T,lt)|0,i=(i=i+Math.imul(T,ct)|0)+Math.imul(k,lt)|0,a=a+Math.imul(k,ct)|0,n=n+Math.imul(b,ft)|0,i=(i=i+Math.imul(b,ht)|0)+Math.imul(_,ft)|0,a=a+Math.imul(_,ht)|0;var St=(c+(n=n+Math.imul(v,dt)|0)|0)+((8191&(i=(i=i+Math.imul(v,mt)|0)+Math.imul(y,dt)|0))<<13)|0;c=((a=a+Math.imul(y,mt)|0)+(i>>>13)|0)+(St>>>26)|0,St&=67108863,n=Math.imul(B,J),i=(i=Math.imul(B,K))+Math.imul(N,J)|0,a=Math.imul(N,K),n=n+Math.imul(D,$)|0,i=(i=i+Math.imul(D,tt)|0)+Math.imul(R,$)|0,a=a+Math.imul(R,tt)|0,n=n+Math.imul(I,rt)|0,i=(i=i+Math.imul(I,nt)|0)+Math.imul(O,rt)|0,a=a+Math.imul(O,nt)|0,n=n+Math.imul(L,at)|0,i=(i=i+Math.imul(L,ot)|0)+Math.imul(C,at)|0,a=a+Math.imul(C,ot)|0,n=n+Math.imul(M,lt)|0,i=(i=i+Math.imul(M,ct)|0)+Math.imul(S,lt)|0,a=a+Math.imul(S,ct)|0,n=n+Math.imul(T,ft)|0,i=(i=i+Math.imul(T,ht)|0)+Math.imul(k,ft)|0,a=a+Math.imul(k,ht)|0;var Et=(c+(n=n+Math.imul(b,dt)|0)|0)+((8191&(i=(i=i+Math.imul(b,mt)|0)+Math.imul(_,dt)|0))<<13)|0;c=((a=a+Math.imul(_,mt)|0)+(i>>>13)|0)+(Et>>>26)|0,Et&=67108863,n=Math.imul(B,$),i=(i=Math.imul(B,tt))+Math.imul(N,$)|0,a=Math.imul(N,tt),n=n+Math.imul(D,rt)|0,i=(i=i+Math.imul(D,nt)|0)+Math.imul(R,rt)|0,a=a+Math.imul(R,nt)|0,n=n+Math.imul(I,at)|0,i=(i=i+Math.imul(I,ot)|0)+Math.imul(O,at)|0,a=a+Math.imul(O,ot)|0,n=n+Math.imul(L,lt)|0,i=(i=i+Math.imul(L,ct)|0)+Math.imul(C,lt)|0,a=a+Math.imul(C,ct)|0,n=n+Math.imul(M,ft)|0,i=(i=i+Math.imul(M,ht)|0)+Math.imul(S,ft)|0,a=a+Math.imul(S,ht)|0;var Lt=(c+(n=n+Math.imul(T,dt)|0)|0)+((8191&(i=(i=i+Math.imul(T,mt)|0)+Math.imul(k,dt)|0))<<13)|0;c=((a=a+Math.imul(k,mt)|0)+(i>>>13)|0)+(Lt>>>26)|0,Lt&=67108863,n=Math.imul(B,rt),i=(i=Math.imul(B,nt))+Math.imul(N,rt)|0,a=Math.imul(N,nt),n=n+Math.imul(D,at)|0,i=(i=i+Math.imul(D,ot)|0)+Math.imul(R,at)|0,a=a+Math.imul(R,ot)|0,n=n+Math.imul(I,lt)|0,i=(i=i+Math.imul(I,ct)|0)+Math.imul(O,lt)|0,a=a+Math.imul(O,ct)|0,n=n+Math.imul(L,ft)|0,i=(i=i+Math.imul(L,ht)|0)+Math.imul(C,ft)|0,a=a+Math.imul(C,ht)|0;var Ct=(c+(n=n+Math.imul(M,dt)|0)|0)+((8191&(i=(i=i+Math.imul(M,mt)|0)+Math.imul(S,dt)|0))<<13)|0;c=((a=a+Math.imul(S,mt)|0)+(i>>>13)|0)+(Ct>>>26)|0,Ct&=67108863,n=Math.imul(B,at),i=(i=Math.imul(B,ot))+Math.imul(N,at)|0,a=Math.imul(N,ot),n=n+Math.imul(D,lt)|0,i=(i=i+Math.imul(D,ct)|0)+Math.imul(R,lt)|0,a=a+Math.imul(R,ct)|0,n=n+Math.imul(I,ft)|0,i=(i=i+Math.imul(I,ht)|0)+Math.imul(O,ft)|0,a=a+Math.imul(O,ht)|0;var Pt=(c+(n=n+Math.imul(L,dt)|0)|0)+((8191&(i=(i=i+Math.imul(L,mt)|0)+Math.imul(C,dt)|0))<<13)|0;c=((a=a+Math.imul(C,mt)|0)+(i>>>13)|0)+(Pt>>>26)|0,Pt&=67108863,n=Math.imul(B,lt),i=(i=Math.imul(B,ct))+Math.imul(N,lt)|0,a=Math.imul(N,ct),n=n+Math.imul(D,ft)|0,i=(i=i+Math.imul(D,ht)|0)+Math.imul(R,ft)|0,a=a+Math.imul(R,ht)|0;var It=(c+(n=n+Math.imul(I,dt)|0)|0)+((8191&(i=(i=i+Math.imul(I,mt)|0)+Math.imul(O,dt)|0))<<13)|0;c=((a=a+Math.imul(O,mt)|0)+(i>>>13)|0)+(It>>>26)|0,It&=67108863,n=Math.imul(B,ft),i=(i=Math.imul(B,ht))+Math.imul(N,ft)|0,a=Math.imul(N,ht);var Ot=(c+(n=n+Math.imul(D,dt)|0)|0)+((8191&(i=(i=i+Math.imul(D,mt)|0)+Math.imul(R,dt)|0))<<13)|0;c=((a=a+Math.imul(R,mt)|0)+(i>>>13)|0)+(Ot>>>26)|0,Ot&=67108863;var zt=(c+(n=Math.imul(B,dt))|0)+((8191&(i=(i=Math.imul(B,mt))+Math.imul(N,dt)|0))<<13)|0;return c=((a=Math.imul(N,mt))+(i>>>13)|0)+(zt>>>26)|0,zt&=67108863,l[0]=gt,l[1]=vt,l[2]=yt,l[3]=xt,l[4]=bt,l[5]=_t,l[6]=wt,l[7]=Tt,l[8]=kt,l[9]=At,l[10]=Mt,l[11]=St,l[12]=Et,l[13]=Lt,l[14]=Ct,l[15]=Pt,l[16]=It,l[17]=Ot,l[18]=zt,0!==c&&(l[19]=c,r.length++),r};function m(t,e,r){return(new g).mulp(t,e,r)}function g(t,e){this.x=t,this.y=e}Math.imul||(d=p),a.prototype.mulTo=function(t,e){var r=this.length+t.length;return 10===this.length&&10===t.length?d(this,t,e):r<63?p(this,t,e):r<1024?function(t,e,r){r.negative=e.negative^t.negative,r.length=t.length+e.length;for(var n=0,i=0,a=0;a<r.length-1;a++){var o=i;i=0;for(var s=67108863&n,l=Math.min(a,e.length-1),c=Math.max(0,a-t.length+1);c<=l;c++){var u=a-c,f=(0|t.words[u])*(0|e.words[c]),h=67108863&f;s=67108863&(h=h+s|0),i+=(o=(o=o+(f/67108864|0)|0)+(h>>>26)|0)>>>26,o&=67108863}r.words[a]=s,n=o,o=i}return 0!==n?r.words[a]=n:r.length--,r.strip()}(this,t,e):m(this,t,e)},g.prototype.makeRBT=function(t){for(var e=new Array(t),r=a.prototype._countBits(t)-1,n=0;n<t;n++)e[n]=this.revBin(n,r,t);return e},g.prototype.revBin=function(t,e,r){if(0===t||t===r-1)return t;for(var n=0,i=0;i<e;i++)n|=(1&t)<<e-i-1,t>>=1;return n},g.prototype.permute=function(t,e,r,n,i,a){for(var o=0;o<a;o++)n[o]=e[t[o]],i[o]=r[t[o]]},g.prototype.transform=function(t,e,r,n,i,a){this.permute(a,t,e,r,n,i);for(var o=1;o<i;o<<=1)for(var s=o<<1,l=Math.cos(2*Math.PI/s),c=Math.sin(2*Math.PI/s),u=0;u<i;u+=s)for(var f=l,h=c,p=0;p<o;p++){var d=r[u+p],m=n[u+p],g=r[u+p+o],v=n[u+p+o],y=f*g-h*v;v=f*v+h*g,g=y,r[u+p]=d+g,n[u+p]=m+v,r[u+p+o]=d-g,n[u+p+o]=m-v,p!==s&&(y=l*f-c*h,h=l*h+c*f,f=y)}},g.prototype.guessLen13b=function(t,e){var r=1|Math.max(e,t),n=1&r,i=0;for(r=r/2|0;r;r>>>=1)i++;return 1<<i+1+n},g.prototype.conjugate=function(t,e,r){if(!(r<=1))for(var n=0;n<r/2;n++){var i=t[n];t[n]=t[r-n-1],t[r-n-1]=i,i=e[n],e[n]=-e[r-n-1],e[r-n-1]=-i}},g.prototype.normalize13b=function(t,e){for(var r=0,n=0;n<e/2;n++){var i=8192*Math.round(t[2*n+1]/e)+Math.round(t[2*n]/e)+r;t[n]=67108863&i,r=i<67108864?0:i/67108864|0}return t},g.prototype.convert13b=function(t,e,r,i){for(var a=0,o=0;o<e;o++)a+=0|t[o],r[2*o]=8191&a,a>>>=13,r[2*o+1]=8191&a,a>>>=13;for(o=2*e;o<i;++o)r[o]=0;n(0===a),n(0==(-8192&a))},g.prototype.stub=function(t){for(var e=new Array(t),r=0;r<t;r++)e[r]=0;return e},g.prototype.mulp=function(t,e,r){var n=2*this.guessLen13b(t.length,e.length),i=this.makeRBT(n),a=this.stub(n),o=new Array(n),s=new Array(n),l=new Array(n),c=new Array(n),u=new Array(n),f=new Array(n),h=r.words;h.length=n,this.convert13b(t.words,t.length,o,n),this.convert13b(e.words,e.length,c,n),this.transform(o,a,s,l,n,i),this.transform(c,a,u,f,n,i);for(var p=0;p<n;p++){var d=s[p]*u[p]-l[p]*f[p];l[p]=s[p]*f[p]+l[p]*u[p],s[p]=d}return this.conjugate(s,l,n),this.transform(s,l,h,a,n,i),this.conjugate(h,a,n),this.normalize13b(h,n),r.negative=t.negative^e.negative,r.length=t.length+e.length,r.strip()},a.prototype.mul=function(t){var e=new a(null);return e.words=new Array(this.length+t.length),this.mulTo(t,e)},a.prototype.mulf=function(t){var e=new a(null);return e.words=new Array(this.length+t.length),m(this,t,e)},a.prototype.imul=function(t){return this.clone().mulTo(t,this)},a.prototype.imuln=function(t){n("number"==typeof t),n(t<67108864);for(var e=0,r=0;r<this.length;r++){var i=(0|this.words[r])*t,a=(67108863&i)+(67108863&e);e>>=26,e+=i/67108864|0,e+=a>>>26,this.words[r]=67108863&a}return 0!==e&&(this.words[r]=e,this.length++),this},a.prototype.muln=function(t){return this.clone().imuln(t)},a.prototype.sqr=function(){return this.mul(this)},a.prototype.isqr=function(){return this.imul(this.clone())},a.prototype.pow=function(t){var e=function(t){for(var e=new Array(t.bitLength()),r=0;r<e.length;r++){var n=r/26|0,i=r%26;e[r]=(t.words[n]&1<<i)>>>i}return e}(t);if(0===e.length)return new a(1);for(var r=this,n=0;n<e.length&&0===e[n];n++,r=r.sqr());if(++n<e.length)for(var i=r.sqr();n<e.length;n++,i=i.sqr())0!==e[n]&&(r=r.mul(i));return r},a.prototype.iushln=function(t){n("number"==typeof t&&t>=0);var e,r=t%26,i=(t-r)/26,a=67108863>>>26-r<<26-r;if(0!==r){var o=0;for(e=0;e<this.length;e++){var s=this.words[e]&a,l=(0|this.words[e])-s<<r;this.words[e]=l|o,o=s>>>26-r}o&&(this.words[e]=o,this.length++)}if(0!==i){for(e=this.length-1;e>=0;e--)this.words[e+i]=this.words[e];for(e=0;e<i;e++)this.words[e]=0;this.length+=i}return this.strip()},a.prototype.ishln=function(t){return n(0===this.negative),this.iushln(t)},a.prototype.iushrn=function(t,e,r){var i;n("number"==typeof t&&t>=0),i=e?(e-e%26)/26:0;var a=t%26,o=Math.min((t-a)/26,this.length),s=67108863^67108863>>>a<<a,l=r;if(i-=o,i=Math.max(0,i),l){for(var c=0;c<o;c++)l.words[c]=this.words[c];l.length=o}if(0===o);else if(this.length>o)for(this.length-=o,c=0;c<this.length;c++)this.words[c]=this.words[c+o];else this.words[0]=0,this.length=1;var u=0;for(c=this.length-1;c>=0&&(0!==u||c>=i);c--){var f=0|this.words[c];this.words[c]=u<<26-a|f>>>a,u=f&s}return l&&0!==u&&(l.words[l.length++]=u),0===this.length&&(this.words[0]=0,this.length=1),this.strip()},a.prototype.ishrn=function(t,e,r){return n(0===this.negative),this.iushrn(t,e,r)},a.prototype.shln=function(t){return this.clone().ishln(t)},a.prototype.ushln=function(t){return this.clone().iushln(t)},a.prototype.shrn=function(t){return this.clone().ishrn(t)},a.prototype.ushrn=function(t){return this.clone().iushrn(t)},a.prototype.testn=function(t){n("number"==typeof t&&t>=0);var e=t%26,r=(t-e)/26,i=1<<e;return!(this.length<=r)&&!!(this.words[r]&i)},a.prototype.imaskn=function(t){n("number"==typeof t&&t>=0);var e=t%26,r=(t-e)/26;if(n(0===this.negative,"imaskn works only with positive numbers"),this.length<=r)return this;if(0!==e&&r++,this.length=Math.min(r,this.length),0!==e){var i=67108863^67108863>>>e<<e;this.words[this.length-1]&=i}return this.strip()},a.prototype.maskn=function(t){return this.clone().imaskn(t)},a.prototype.iaddn=function(t){return n("number"==typeof t),n(t<67108864),t<0?this.isubn(-t):0!==this.negative?1===this.length&&(0|this.words[0])<t?(this.words[0]=t-(0|this.words[0]),this.negative=0,this):(this.negative=0,this.isubn(t),this.negative=1,this):this._iaddn(t)},a.prototype._iaddn=function(t){this.words[0]+=t;for(var e=0;e<this.length&&this.words[e]>=67108864;e++)this.words[e]-=67108864,e===this.length-1?this.words[e+1]=1:this.words[e+1]++;return this.length=Math.max(this.length,e+1),this},a.prototype.isubn=function(t){if(n("number"==typeof t),n(t<67108864),t<0)return this.iaddn(-t);if(0!==this.negative)return this.negative=0,this.iaddn(t),this.negative=1,this;if(this.words[0]-=t,1===this.length&&this.words[0]<0)this.words[0]=-this.words[0],this.negative=1;else for(var e=0;e<this.length&&this.words[e]<0;e++)this.words[e]+=67108864,this.words[e+1]-=1;return this.strip()},a.prototype.addn=function(t){return this.clone().iaddn(t)},a.prototype.subn=function(t){return this.clone().isubn(t)},a.prototype.iabs=function(){return this.negative=0,this},a.prototype.abs=function(){return this.clone().iabs()},a.prototype._ishlnsubmul=function(t,e,r){var i,a,o=t.length+r;this._expand(o);var s=0;for(i=0;i<t.length;i++){a=(0|this.words[i+r])+s;var l=(0|t.words[i])*e;s=((a-=67108863&l)>>26)-(l/67108864|0),this.words[i+r]=67108863&a}for(;i<this.length-r;i++)s=(a=(0|this.words[i+r])+s)>>26,this.words[i+r]=67108863&a;if(0===s)return this.strip();for(n(-1===s),s=0,i=0;i<this.length;i++)s=(a=-(0|this.words[i])+s)>>26,this.words[i]=67108863&a;return this.negative=1,this.strip()},a.prototype._wordDiv=function(t,e){var r=(this.length,t.length),n=this.clone(),i=t,o=0|i.words[i.length-1];0!==(r=26-this._countBits(o))&&(i=i.ushln(r),n.iushln(r),o=0|i.words[i.length-1]);var s,l=n.length-i.length;if("mod"!==e){(s=new a(null)).length=l+1,s.words=new Array(s.length);for(var c=0;c<s.length;c++)s.words[c]=0}var u=n.clone()._ishlnsubmul(i,1,l);0===u.negative&&(n=u,s&&(s.words[l]=1));for(var f=l-1;f>=0;f--){var h=67108864*(0|n.words[i.length+f])+(0|n.words[i.length+f-1]);for(h=Math.min(h/o|0,67108863),n._ishlnsubmul(i,h,f);0!==n.negative;)h--,n.negative=0,n._ishlnsubmul(i,1,f),n.isZero()||(n.negative^=1);s&&(s.words[f]=h)}return s&&s.strip(),n.strip(),"div"!==e&&0!==r&&n.iushrn(r),{div:s||null,mod:n}},a.prototype.divmod=function(t,e,r){return n(!t.isZero()),this.isZero()?{div:new a(0),mod:new a(0)}:0!==this.negative&&0===t.negative?(s=this.neg().divmod(t,e),"mod"!==e&&(i=s.div.neg()),"div"!==e&&(o=s.mod.neg(),r&&0!==o.negative&&o.iadd(t)),{div:i,mod:o}):0===this.negative&&0!==t.negative?(s=this.divmod(t.neg(),e),"mod"!==e&&(i=s.div.neg()),{div:i,mod:s.mod}):0!=(this.negative&t.negative)?(s=this.neg().divmod(t.neg(),e),"div"!==e&&(o=s.mod.neg(),r&&0!==o.negative&&o.isub(t)),{div:s.div,mod:o}):t.length>this.length||this.cmp(t)<0?{div:new a(0),mod:this}:1===t.length?"div"===e?{div:this.divn(t.words[0]),mod:null}:"mod"===e?{div:null,mod:new a(this.modn(t.words[0]))}:{div:this.divn(t.words[0]),mod:new a(this.modn(t.words[0]))}:this._wordDiv(t,e);var i,o,s},a.prototype.div=function(t){return this.divmod(t,"div",!1).div},a.prototype.mod=function(t){return this.divmod(t,"mod",!1).mod},a.prototype.umod=function(t){return this.divmod(t,"mod",!0).mod},a.prototype.divRound=function(t){var e=this.divmod(t);if(e.mod.isZero())return e.div;var r=0!==e.div.negative?e.mod.isub(t):e.mod,n=t.ushrn(1),i=t.andln(1),a=r.cmp(n);return a<0||1===i&&0===a?e.div:0!==e.div.negative?e.div.isubn(1):e.div.iaddn(1)},a.prototype.modn=function(t){n(t<=67108863);for(var e=(1<<26)%t,r=0,i=this.length-1;i>=0;i--)r=(e*r+(0|this.words[i]))%t;return r},a.prototype.idivn=function(t){n(t<=67108863);for(var e=0,r=this.length-1;r>=0;r--){var i=(0|this.words[r])+67108864*e;this.words[r]=i/t|0,e=i%t}return this.strip()},a.prototype.divn=function(t){return this.clone().idivn(t)},a.prototype.egcd=function(t){n(0===t.negative),n(!t.isZero());var e=this,r=t.clone();e=0!==e.negative?e.umod(t):e.clone();for(var i=new a(1),o=new a(0),s=new a(0),l=new a(1),c=0;e.isEven()&&r.isEven();)e.iushrn(1),r.iushrn(1),++c;for(var u=r.clone(),f=e.clone();!e.isZero();){for(var h=0,p=1;0==(e.words[0]&p)&&h<26;++h,p<<=1);if(h>0)for(e.iushrn(h);h-- >0;)(i.isOdd()||o.isOdd())&&(i.iadd(u),o.isub(f)),i.iushrn(1),o.iushrn(1);for(var d=0,m=1;0==(r.words[0]&m)&&d<26;++d,m<<=1);if(d>0)for(r.iushrn(d);d-- >0;)(s.isOdd()||l.isOdd())&&(s.iadd(u),l.isub(f)),s.iushrn(1),l.iushrn(1);e.cmp(r)>=0?(e.isub(r),i.isub(s),o.isub(l)):(r.isub(e),s.isub(i),l.isub(o))}return{a:s,b:l,gcd:r.iushln(c)}},a.prototype._invmp=function(t){n(0===t.negative),n(!t.isZero());var e=this,r=t.clone();e=0!==e.negative?e.umod(t):e.clone();for(var i,o=new a(1),s=new a(0),l=r.clone();e.cmpn(1)>0&&r.cmpn(1)>0;){for(var c=0,u=1;0==(e.words[0]&u)&&c<26;++c,u<<=1);if(c>0)for(e.iushrn(c);c-- >0;)o.isOdd()&&o.iadd(l),o.iushrn(1);for(var f=0,h=1;0==(r.words[0]&h)&&f<26;++f,h<<=1);if(f>0)for(r.iushrn(f);f-- >0;)s.isOdd()&&s.iadd(l),s.iushrn(1);e.cmp(r)>=0?(e.isub(r),o.isub(s)):(r.isub(e),s.isub(o))}return(i=0===e.cmpn(1)?o:s).cmpn(0)<0&&i.iadd(t),i},a.prototype.gcd=function(t){if(this.isZero())return t.abs();if(t.isZero())return this.abs();var e=this.clone(),r=t.clone();e.negative=0,r.negative=0;for(var n=0;e.isEven()&&r.isEven();n++)e.iushrn(1),r.iushrn(1);for(;;){for(;e.isEven();)e.iushrn(1);for(;r.isEven();)r.iushrn(1);var i=e.cmp(r);if(i<0){var a=e;e=r,r=a}else if(0===i||0===r.cmpn(1))break;e.isub(r)}return r.iushln(n)},a.prototype.invm=function(t){return this.egcd(t).a.umod(t)},a.prototype.isEven=function(){return 0==(1&this.words[0])},a.prototype.isOdd=function(){return 1==(1&this.words[0])},a.prototype.andln=function(t){return this.words[0]&t},a.prototype.bincn=function(t){n("number"==typeof t);var e=t%26,r=(t-e)/26,i=1<<e;if(this.length<=r)return this._expand(r+1),this.words[r]|=i,this;for(var a=i,o=r;0!==a&&o<this.length;o++){var s=0|this.words[o];a=(s+=a)>>>26,s&=67108863,this.words[o]=s}return 0!==a&&(this.words[o]=a,this.length++),this},a.prototype.isZero=function(){return 1===this.length&&0===this.words[0]},a.prototype.cmpn=function(t){var e,r=t<0;if(0!==this.negative&&!r)return-1;if(0===this.negative&&r)return 1;if(this.strip(),this.length>1)e=1;else{r&&(t=-t),n(t<=67108863,"Number is too big");var i=0|this.words[0];e=i===t?0:i<t?-1:1}return 0!==this.negative?0|-e:e},a.prototype.cmp=function(t){if(0!==this.negative&&0===t.negative)return-1;if(0===this.negative&&0!==t.negative)return 1;var e=this.ucmp(t);return 0!==this.negative?0|-e:e},a.prototype.ucmp=function(t){if(this.length>t.length)return 1;if(this.length<t.length)return-1;for(var e=0,r=this.length-1;r>=0;r--){var n=0|this.words[r],i=0|t.words[r];if(n!==i){n<i?e=-1:n>i&&(e=1);break}}return e},a.prototype.gtn=function(t){return 1===this.cmpn(t)},a.prototype.gt=function(t){return 1===this.cmp(t)},a.prototype.gten=function(t){return this.cmpn(t)>=0},a.prototype.gte=function(t){return this.cmp(t)>=0},a.prototype.ltn=function(t){return-1===this.cmpn(t)},a.prototype.lt=function(t){return-1===this.cmp(t)},a.prototype.lten=function(t){return this.cmpn(t)<=0},a.prototype.lte=function(t){return this.cmp(t)<=0},a.prototype.eqn=function(t){return 0===this.cmpn(t)},a.prototype.eq=function(t){return 0===this.cmp(t)},a.red=function(t){return new T(t)},a.prototype.toRed=function(t){return n(!this.red,"Already a number in reduction context"),n(0===this.negative,"red works only with positives"),t.convertTo(this)._forceRed(t)},a.prototype.fromRed=function(){return n(this.red,"fromRed works only with numbers in reduction context"),this.red.convertFrom(this)},a.prototype._forceRed=function(t){return this.red=t,this},a.prototype.forceRed=function(t){return n(!this.red,"Already a number in reduction context"),this._forceRed(t)},a.prototype.redAdd=function(t){return n(this.red,"redAdd works only with red numbers"),this.red.add(this,t)},a.prototype.redIAdd=function(t){return n(this.red,"redIAdd works only with red numbers"),this.red.iadd(this,t)},a.prototype.redSub=function(t){return n(this.red,"redSub works only with red numbers"),this.red.sub(this,t)},a.prototype.redISub=function(t){return n(this.red,"redISub works only with red numbers"),this.red.isub(this,t)},a.prototype.redShl=function(t){return n(this.red,"redShl works only with red numbers"),this.red.shl(this,t)},a.prototype.redMul=function(t){return n(this.red,"redMul works only with red numbers"),this.red._verify2(this,t),this.red.mul(this,t)},a.prototype.redIMul=function(t){return n(this.red,"redMul works only with red numbers"),this.red._verify2(this,t),this.red.imul(this,t)},a.prototype.redSqr=function(){return n(this.red,"redSqr works only with red numbers"),this.red._verify1(this),this.red.sqr(this)},a.prototype.redISqr=function(){return n(this.red,"redISqr works only with red numbers"),this.red._verify1(this),this.red.isqr(this)},a.prototype.redSqrt=function(){return n(this.red,"redSqrt works only with red numbers"),this.red._verify1(this),this.red.sqrt(this)},a.prototype.redInvm=function(){return n(this.red,"redInvm works only with red numbers"),this.red._verify1(this),this.red.invm(this)},a.prototype.redNeg=function(){return n(this.red,"redNeg works only with red numbers"),this.red._verify1(this),this.red.neg(this)},a.prototype.redPow=function(t){return n(this.red&&!t.red,"redPow(normalNum)"),this.red._verify1(this),this.red.pow(this,t)};var v={k256:null,p224:null,p192:null,p25519:null};function y(t,e){this.name=t,this.p=new a(e,16),this.n=this.p.bitLength(),this.k=new a(1).iushln(this.n).isub(this.p),this.tmp=this._tmp()}function x(){y.call(this,"k256","ffffffff ffffffff ffffffff ffffffff ffffffff ffffffff fffffffe fffffc2f")}function b(){y.call(this,"p224","ffffffff ffffffff ffffffff ffffffff 00000000 00000000 00000001")}function _(){y.call(this,"p192","ffffffff ffffffff ffffffff fffffffe ffffffff ffffffff")}function w(){y.call(this,"25519","7fffffffffffffff ffffffffffffffff ffffffffffffffff ffffffffffffffed")}function T(t){if("string"==typeof t){var e=a._prime(t);this.m=e.p,this.prime=e}else n(t.gtn(1),"modulus must be greater than 1"),this.m=t,this.prime=null}function k(t){T.call(this,t),this.shift=this.m.bitLength(),this.shift%26!=0&&(this.shift+=26-this.shift%26),this.r=new a(1).iushln(this.shift),this.r2=this.imod(this.r.sqr()),this.rinv=this.r._invmp(this.m),this.minv=this.rinv.mul(this.r).isubn(1).div(this.m),this.minv=this.minv.umod(this.r),this.minv=this.r.sub(this.minv)}y.prototype._tmp=function(){var t=new a(null);return t.words=new Array(Math.ceil(this.n/13)),t},y.prototype.ireduce=function(t){var e,r=t;do{this.split(r,this.tmp),e=(r=(r=this.imulK(r)).iadd(this.tmp)).bitLength()}while(e>this.n);var n=e<this.n?-1:r.ucmp(this.p);return 0===n?(r.words[0]=0,r.length=1):n>0?r.isub(this.p):void 0!==r.strip?r.strip():r._strip(),r},y.prototype.split=function(t,e){t.iushrn(this.n,0,e)},y.prototype.imulK=function(t){return t.imul(this.k)},i(x,y),x.prototype.split=function(t,e){for(var r=Math.min(t.length,9),n=0;n<r;n++)e.words[n]=t.words[n];if(e.length=r,t.length<=9)return t.words[0]=0,void(t.length=1);var i=t.words[9];for(e.words[e.length++]=4194303&i,n=10;n<t.length;n++){var a=0|t.words[n];t.words[n-10]=(4194303&a)<<4|i>>>22,i=a}i>>>=22,t.words[n-10]=i,0===i&&t.length>10?t.length-=10:t.length-=9},x.prototype.imulK=function(t){t.words[t.length]=0,t.words[t.length+1]=0,t.length+=2;for(var e=0,r=0;r<t.length;r++){var n=0|t.words[r];e+=977*n,t.words[r]=67108863&e,e=64*n+(e/67108864|0)}return 0===t.words[t.length-1]&&(t.length--,0===t.words[t.length-1]&&t.length--),t},i(b,y),i(_,y),i(w,y),w.prototype.imulK=function(t){for(var e=0,r=0;r<t.length;r++){var n=19*(0|t.words[r])+e,i=67108863&n;n>>>=26,t.words[r]=i,e=n}return 0!==e&&(t.words[t.length++]=e),t},a._prime=function(t){if(v[t])return v[t];var e;if("k256"===t)e=new x;else if("p224"===t)e=new b;else if("p192"===t)e=new _;else{if("p25519"!==t)throw new Error("Unknown prime "+t);e=new w}return v[t]=e,e},T.prototype._verify1=function(t){n(0===t.negative,"red works only with positives"),n(t.red,"red works only with red numbers")},T.prototype._verify2=function(t,e){n(0==(t.negative|e.negative),"red works only with positives"),n(t.red&&t.red===e.red,"red works only with red numbers")},T.prototype.imod=function(t){return this.prime?this.prime.ireduce(t)._forceRed(this):t.umod(this.m)._forceRed(this)},T.prototype.neg=function(t){return t.isZero()?t.clone():this.m.sub(t)._forceRed(this)},T.prototype.add=function(t,e){this._verify2(t,e);var r=t.add(e);return r.cmp(this.m)>=0&&r.isub(this.m),r._forceRed(this)},T.prototype.iadd=function(t,e){this._verify2(t,e);var r=t.iadd(e);return r.cmp(this.m)>=0&&r.isub(this.m),r},T.prototype.sub=function(t,e){this._verify2(t,e);var r=t.sub(e);return r.cmpn(0)<0&&r.iadd(this.m),r._forceRed(this)},T.prototype.isub=function(t,e){this._verify2(t,e);var r=t.isub(e);return r.cmpn(0)<0&&r.iadd(this.m),r},T.prototype.shl=function(t,e){return this._verify1(t),this.imod(t.ushln(e))},T.prototype.imul=function(t,e){return this._verify2(t,e),this.imod(t.imul(e))},T.prototype.mul=function(t,e){return this._verify2(t,e),this.imod(t.mul(e))},T.prototype.isqr=function(t){return this.imul(t,t.clone())},T.prototype.sqr=function(t){return this.mul(t,t)},T.prototype.sqrt=function(t){if(t.isZero())return t.clone();var e=this.m.andln(3);if(n(e%2==1),3===e){var r=this.m.add(new a(1)).iushrn(2);return this.pow(t,r)}for(var i=this.m.subn(1),o=0;!i.isZero()&&0===i.andln(1);)o++,i.iushrn(1);n(!i.isZero());var s=new a(1).toRed(this),l=s.redNeg(),c=this.m.subn(1).iushrn(1),u=this.m.bitLength();for(u=new a(2*u*u).toRed(this);0!==this.pow(u,c).cmp(l);)u.redIAdd(l);for(var f=this.pow(u,i),h=this.pow(t,i.addn(1).iushrn(1)),p=this.pow(t,i),d=o;0!==p.cmp(s);){for(var m=p,g=0;0!==m.cmp(s);g++)m=m.redSqr();n(g<d);var v=this.pow(f,new a(1).iushln(d-g-1));h=h.redMul(v),f=v.redSqr(),p=p.redMul(f),d=g}return h},T.prototype.invm=function(t){var e=t._invmp(this.m);return 0!==e.negative?(e.negative=0,this.imod(e).redNeg()):this.imod(e)},T.prototype.pow=function(t,e){if(e.isZero())return new a(1).toRed(this);if(0===e.cmpn(1))return t.clone();var r=new Array(16);r[0]=new a(1).toRed(this),r[1]=t;for(var n=2;n<r.length;n++)r[n]=this.mul(r[n-1],t);var i=r[0],o=0,s=0,l=e.bitLength()%26;for(0===l&&(l=26),n=e.length-1;n>=0;n--){for(var c=e.words[n],u=l-1;u>=0;u--){var f=c>>u&1;i!==r[0]&&(i=this.sqr(i)),0!==f||0!==o?(o<<=1,o|=f,(4===++s||0===n&&0===u)&&(i=this.mul(i,r[o]),s=0,o=0)):s=0}l=26}return i},T.prototype.convertTo=function(t){var e=t.umod(this.m);return e===t?e.clone():e},T.prototype.convertFrom=function(t){var e=t.clone();return e.red=null,e},a.mont=function(t){return new k(t)},i(k,T),k.prototype.convertTo=function(t){return this.imod(t.ushln(this.shift))},k.prototype.convertFrom=function(t){var e=this.imod(t.mul(this.rinv));return e.red=null,e},k.prototype.imul=function(t,e){if(t.isZero()||e.isZero())return t.words[0]=0,t.length=1,t;var r=t.imul(e),n=r.maskn(this.shift).mul(this.minv).imaskn(this.shift).mul(this.m),i=r.isub(n).iushrn(this.shift),a=i;return i.cmp(this.m)>=0?a=i.isub(this.m):i.cmpn(0)<0&&(a=i.iadd(this.m)),a._forceRed(this)},k.prototype.mul=function(t,e){if(t.isZero()||e.isZero())return new a(0)._forceRed(this);var r=t.mul(e),n=r.maskn(this.shift).mul(this.minv).imaskn(this.shift).mul(this.m),i=r.isub(n).iushrn(this.shift),o=i;return i.cmp(this.m)>=0?o=i.isub(this.m):i.cmpn(0)<0&&(o=i.iadd(this.m)),o._forceRed(this)},k.prototype.invm=function(t){return this.imod(t._invmp(this.m).mul(this.r2))._forceRed(this)}}(void 0===e||e,this)},{buffer:111}],97:[function(t,e,r){"use strict";var n=t("./lib/bn-sign");e.exports=function(t){return n(t[0])*n(t[1])}},{"./lib/bn-sign":88}],98:[function(t,e,r){"use strict";var n=t("./lib/rationalize");e.exports=function(t,e){return n(t[0].mul(e[1]).sub(t[1].mul(e[0])),t[1].mul(e[1]))}},{"./lib/rationalize":93}],99:[function(t,e,r){"use strict";var n=t("./lib/bn-to-num"),i=t("./lib/ctz");e.exports=function(t){var e=t[0],r=t[1];if(0===e.cmpn(0))return 0;var a=e.abs().divmod(r.abs()),o=a.div,s=n(o),l=a.mod,c=e.negative!==r.negative?-1:1;if(0===l.cmpn(0))return c*s;if(s){var u=i(s)+4,f=n(l.ushln(u).divRound(r));return c*(s+f*Math.pow(2,-u))}var h=r.bitLength()-l.bitLength()+53;f=n(l.ushln(h).divRound(r));return h<1023?c*f*Math.pow(2,-h):(f*=Math.pow(2,-1023),c*f*Math.pow(2,1023-h))}},{"./lib/bn-to-num":89,"./lib/ctz":90}],100:[function(t,e,r){"use strict";function n(t,e,r,n,i){for(var a=i+1;n<=i;){var o=n+i>>>1,s=t[o];(void 0!==r?r(s,e):s-e)>=0?(a=o,i=o-1):n=o+1}return a}function i(t,e,r,n,i){for(var a=i+1;n<=i;){var o=n+i>>>1,s=t[o];(void 0!==r?r(s,e):s-e)>0?(a=o,i=o-1):n=o+1}return a}function a(t,e,r,n,i){for(var a=n-1;n<=i;){var o=n+i>>>1,s=t[o];(void 0!==r?r(s,e):s-e)<0?(a=o,n=o+1):i=o-1}return a}function o(t,e,r,n,i){for(var a=n-1;n<=i;){var o=n+i>>>1,s=t[o];(void 0!==r?r(s,e):s-e)<=0?(a=o,n=o+1):i=o-1}return a}function s(t,e,r,n,i){for(;n<=i;){var a=n+i>>>1,o=t[a],s=void 0!==r?r(o,e):o-e;if(0===s)return a;s<=0?n=a+1:i=a-1}return-1}function l(t,e,r,n,i,a){return"function"==typeof r?a(t,e,r,void 0===n?0:0|n,void 0===i?t.length-1:0|i):a(t,e,void 0,void 0===r?0:0|r,void 0===n?t.length-1:0|n)}e.exports={ge:function(t,e,r,i,a){return l(t,e,r,i,a,n)},gt:function(t,e,r,n,a){return l(t,e,r,n,a,i)},lt:function(t,e,r,n,i){return l(t,e,r,n,i,a)},le:function(t,e,r,n,i){return l(t,e,r,n,i,o)},eq:function(t,e,r,n,i){return l(t,e,r,n,i,s)}}},{}],101:[function(t,e,r){"use strict";function n(t){var e=32;return(t&=-t)&&e--,65535&t&&(e-=16),16711935&t&&(e-=8),252645135&t&&(e-=4),858993459&t&&(e-=2),1431655765&t&&(e-=1),e}r.INT_BITS=32,r.INT_MAX=2147483647,r.INT_MIN=-1<<31,r.sign=function(t){return(t>0)-(t<0)},r.abs=function(t){var e=t>>31;return(t^e)-e},r.min=function(t,e){return e^(t^e)&-(t<e)},r.max=function(t,e){return t^(t^e)&-(t<e)},r.isPow2=function(t){return!(t&t-1||!t)},r.log2=function(t){var e,r;return e=(t>65535)<<4,e|=r=((t>>>=e)>255)<<3,e|=r=((t>>>=r)>15)<<2,(e|=r=((t>>>=r)>3)<<1)|(t>>>=r)>>1},r.log10=function(t){return t>=1e9?9:t>=1e8?8:t>=1e7?7:t>=1e6?6:t>=1e5?5:t>=1e4?4:t>=1e3?3:t>=100?2:t>=10?1:0},r.popCount=function(t){return 16843009*((t=(858993459&(t-=t>>>1&1431655765))+(t>>>2&858993459))+(t>>>4)&252645135)>>>24},r.countTrailingZeros=n,r.nextPow2=function(t){return t+=0===t,--t,t|=t>>>1,t|=t>>>2,t|=t>>>4,t|=t>>>8,(t|=t>>>16)+1},r.prevPow2=function(t){return t|=t>>>1,t|=t>>>2,t|=t>>>4,t|=t>>>8,(t|=t>>>16)-(t>>>1)},r.parity=function(t){return t^=t>>>16,t^=t>>>8,t^=t>>>4,27030>>>(t&=15)&1};var i=new Array(256);!function(t){for(var e=0;e<256;++e){var r=e,n=e,i=7;for(r>>>=1;r;r>>>=1)n<<=1,n|=1&r,--i;t[e]=n<<i&255}}(i),r.reverse=function(t){return i[255&t]<<24|i[t>>>8&255]<<16|i[t>>>16&255]<<8|i[t>>>24&255]},r.interleave2=function(t,e){return(t=1431655765&((t=858993459&((t=252645135&((t=16711935&((t&=65535)|t<<8))|t<<4))|t<<2))|t<<1))|(e=1431655765&((e=858993459&((e=252645135&((e=16711935&((e&=65535)|e<<8))|e<<4))|e<<2))|e<<1))<<1},r.deinterleave2=function(t,e){return(t=65535&((t=16711935&((t=252645135&((t=858993459&((t=t>>>e&1431655765)|t>>>1))|t>>>2))|t>>>4))|t>>>16))<<16>>16},r.interleave3=function(t,e,r){return t=1227133513&((t=3272356035&((t=251719695&((t=4278190335&((t&=1023)|t<<16))|t<<8))|t<<4))|t<<2),(t|=(e=1227133513&((e=3272356035&((e=251719695&((e=4278190335&((e&=1023)|e<<16))|e<<8))|e<<4))|e<<2))<<1)|(r=1227133513&((r=3272356035&((r=251719695&((r=4278190335&((r&=1023)|r<<16))|r<<8))|r<<4))|r<<2))<<2},r.deinterleave3=function(t,e){return(t=1023&((t=4278190335&((t=251719695&((t=3272356035&((t=t>>>e&1227133513)|t>>>2))|t>>>4))|t>>>8))|t>>>16))<<22>>22},r.nextCombination=function(t){var e=t|t-1;return e+1|(~e&-~e)-1>>>n(t)+1}},{}],102:[function(t,e,r){"use strict";var n=t("clamp");e.exports=function(t,e){e||(e={});var r,o,s,l,c,u,f,h,p,d,m,g=null==e.cutoff?.25:e.cutoff,v=null==e.radius?8:e.radius,y=e.channel||0;if(ArrayBuffer.isView(t)||Array.isArray(t)){if(!e.width||!e.height)throw Error("For raw data width and height should be provided by options");r=e.width,o=e.height,l=t,u=e.stride?e.stride:Math.floor(t.length/r/o)}else window.HTMLCanvasElement&&t instanceof window.HTMLCanvasElement?(f=(h=t).getContext("2d"),r=h.width,o=h.height,p=f.getImageData(0,0,r,o),l=p.data,u=4):window.CanvasRenderingContext2D&&t instanceof window.CanvasRenderingContext2D?(h=t.canvas,f=t,r=h.width,o=h.height,p=f.getImageData(0,0,r,o),l=p.data,u=4):window.ImageData&&t instanceof window.ImageData&&(p=t,r=t.width,o=t.height,l=p.data,u=4);if(s=Math.max(r,o),window.Uint8ClampedArray&&l instanceof window.Uint8ClampedArray||window.Uint8Array&&l instanceof window.Uint8Array)for(c=l,l=Array(r*o),d=0,m=c.length;d<m;d++)l[d]=c[d*u+y]/255;else if(1!==u)throw Error("Raw data can have only 1 value per pixel");var x=Array(r*o),b=Array(r*o),_=Array(s),w=Array(s),T=Array(s+1),k=Array(s);for(d=0,m=r*o;d<m;d++){var A=l[d];x[d]=1===A?0:0===A?i:Math.pow(Math.max(0,.5-A),2),b[d]=1===A?i:0===A?0:Math.pow(Math.max(0,A-.5),2)}a(x,r,o,_,w,k,T),a(b,r,o,_,w,k,T);var M=window.Float32Array?new Float32Array(r*o):new Array(r*o);for(d=0,m=r*o;d<m;d++)M[d]=n(1-((x[d]-b[d])/v+g),0,1);return M};var i=1e20;function a(t,e,r,n,i,a,s){for(var l=0;l<e;l++){for(var c=0;c<r;c++)n[c]=t[c*e+l];for(o(n,i,a,s,r),c=0;c<r;c++)t[c*e+l]=i[c]}for(c=0;c<r;c++){for(l=0;l<e;l++)n[l]=t[c*e+l];for(o(n,i,a,s,e),l=0;l<e;l++)t[c*e+l]=Math.sqrt(i[l])}}function o(t,e,r,n,a){r[0]=0,n[0]=-i,n[1]=+i;for(var o=1,s=0;o<a;o++){for(var l=(t[o]+o*o-(t[r[s]]+r[s]*r[s]))/(2*o-2*r[s]);l<=n[s];)s--,l=(t[o]+o*o-(t[r[s]]+r[s]*r[s]))/(2*o-2*r[s]);r[++s]=o,n[s]=l,n[s+1]=+i}for(o=0,s=0;o<a;o++){for(;n[s+1]<o;)s++;e[o]=(o-r[s])*(o-r[s])+t[r[s]]}}},{clamp:121}],103:[function(t,e,r){"use strict";e.exports=function(t){var e,r,n,i=t.length,a=0;for(e=0;e<i;++e)a+=t[e].length;var o=new Array(a),s=0;for(e=0;e<i;++e){var l=t[e],c=l.length;for(r=0;r<c;++r){var u=o[s++]=new Array(c-1),f=0;for(n=0;n<c;++n)n!==r&&(u[f++]=l[n]);if(1&r){var h=u[1];u[1]=u[0],u[0]=h}}}return o}},{}],104:[function(t,e,r){"use strict";e.exports=function(t,e,r){switch(arguments.length){case 1:return f(t);case 2:return"function"==typeof e?c(t,t,e,!0):h(t,e);case 3:return c(t,e,r,!1);default:throw new Error("box-intersect: Invalid arguments")}};var n,i=t("typedarray-pool"),a=t("./lib/sweep"),o=t("./lib/intersect");function s(t,e){for(var r=0;r<t;++r)if(!(e[r]<=e[r+t]))return!0;return!1}function l(t,e,r,n){for(var i=0,a=0,o=0,l=t.length;o<l;++o){var c=t[o];if(!s(e,c)){for(var u=0;u<2*e;++u)r[i++]=c[u];n[a++]=o}}return a}function c(t,e,r,n){var s=t.length,c=e.length;if(!(s<=0||c<=0)){var u=t[0].length>>>1;if(!(u<=0)){var f,h=i.mallocDouble(2*u*s),p=i.mallocInt32(s);if((s=l(t,u,h,p))>0){if(1===u&&n)a.init(s),f=a.sweepComplete(u,r,0,s,h,p,0,s,h,p);else{var d=i.mallocDouble(2*u*c),m=i.mallocInt32(c);(c=l(e,u,d,m))>0&&(a.init(s+c),f=1===u?a.sweepBipartite(u,r,0,s,h,p,0,c,d,m):o(u,r,n,s,h,p,c,d,m),i.free(d),i.free(m))}i.free(h),i.free(p)}return f}}}function u(t,e){n.push([t,e])}function f(t){return n=[],c(t,t,u,!0),n}function h(t,e){return n=[],c(t,e,u,!1),n}},{"./lib/intersect":106,"./lib/sweep":110,"typedarray-pool":590}],105:[function(t,e,r){"use strict";function n(t){return t?function(t,e,r,n,i,a,o,s,l,c,u){return i-n>l-s?function(t,e,r,n,i,a,o,s,l,c,u){for(var f=2*t,h=n,p=f*n;h<i;++h,p+=f){var d=a[e+p],m=a[e+p+t],g=o[h];t:for(var v=s,y=f*s;v<l;++v,y+=f){var x=c[e+y],b=c[e+y+t],_=u[v];if(!(b<d||m<x)){for(var w=e+1;w<t;++w){var T=a[w+p],k=a[w+t+p],A=c[w+y],M=c[w+t+y];if(k<A||M<T)continue t}var S=r(g,_);if(void 0!==S)return S}}}}(t,e,r,n,i,a,o,s,l,c,u):function(t,e,r,n,i,a,o,s,l,c,u){for(var f=2*t,h=s,p=f*s;h<l;++h,p+=f){var d=c[e+p],m=c[e+p+t],g=u[h];t:for(var v=n,y=f*n;v<i;++v,y+=f){var x=a[e+y],b=a[e+y+t],_=o[v];if(!(m<x||b<d)){for(var w=e+1;w<t;++w){var T=a[w+y],k=a[w+t+y],A=c[w+p],M=c[w+t+p];if(k<A||M<T)continue t}var S=r(_,g);if(void 0!==S)return S}}}}(t,e,r,n,i,a,o,s,l,c,u)}:function(t,e,r,n,i,a,o,s,l,c,u,f){return a-i>c-l?n?function(t,e,r,n,i,a,o,s,l,c,u){for(var f=2*t,h=n,p=f*n;h<i;++h,p+=f){var d=a[e+p],m=a[e+p+t],g=o[h];t:for(var v=s,y=f*s;v<l;++v,y+=f){var x=c[e+y],b=u[v];if(!(x<=d||m<x)){for(var _=e+1;_<t;++_){var w=a[_+p],T=a[_+t+p],k=c[_+y],A=c[_+t+y];if(T<k||A<w)continue t}var M=r(b,g);if(void 0!==M)return M}}}}(t,e,r,i,a,o,s,l,c,u,f):function(t,e,r,n,i,a,o,s,l,c,u){for(var f=2*t,h=n,p=f*n;h<i;++h,p+=f){var d=a[e+p],m=a[e+p+t],g=o[h];t:for(var v=s,y=f*s;v<l;++v,y+=f){var x=c[e+y],b=u[v];if(!(x<d||m<x)){for(var _=e+1;_<t;++_){var w=a[_+p],T=a[_+t+p],k=c[_+y],A=c[_+t+y];if(T<k||A<w)continue t}var M=r(g,b);if(void 0!==M)return M}}}}(t,e,r,i,a,o,s,l,c,u,f):n?function(t,e,r,n,i,a,o,s,l,c,u){for(var f=2*t,h=s,p=f*s;h<l;++h,p+=f){var d=c[e+p],m=u[h];t:for(var g=n,v=f*n;g<i;++g,v+=f){var y=a[e+v],x=a[e+v+t],b=o[g];if(!(d<=y||x<d)){for(var _=e+1;_<t;++_){var w=a[_+v],T=a[_+t+v],k=c[_+p],A=c[_+t+p];if(T<k||A<w)continue t}var M=r(m,b);if(void 0!==M)return M}}}}(t,e,r,i,a,o,s,l,c,u,f):function(t,e,r,n,i,a,o,s,l,c,u){for(var f=2*t,h=s,p=f*s;h<l;++h,p+=f){var d=c[e+p],m=u[h];t:for(var g=n,v=f*n;g<i;++g,v+=f){var y=a[e+v],x=a[e+v+t],b=o[g];if(!(d<y||x<d)){for(var _=e+1;_<t;++_){var w=a[_+v],T=a[_+t+v],k=c[_+p],A=c[_+t+p];if(T<k||A<w)continue t}var M=r(b,m);if(void 0!==M)return M}}}}(t,e,r,i,a,o,s,l,c,u,f)}}r.partial=n(!1),r.full=n(!0)},{}],106:[function(t,e,r){"use strict";e.exports=function(t,e,r,a,u,w,T,k,A){!function(t,e){var r=8*i.log2(e+1)*(t+1)|0,a=i.nextPow2(6*r);v.length<a&&(n.free(v),v=n.mallocInt32(a));var o=i.nextPow2(2*r);y.length<o&&(n.free(y),y=n.mallocDouble(o))}(t,a+T);var M,S=0,E=2*t;x(S++,0,0,a,0,T,r?16:0,-1/0,1/0),r||x(S++,0,0,T,0,a,1,-1/0,1/0);for(;S>0;){var L=6*(S-=1),C=v[L],P=v[L+1],I=v[L+2],O=v[L+3],z=v[L+4],D=v[L+5],R=2*S,F=y[R],B=y[R+1],N=1&D,j=!!(16&D),U=u,V=w,H=k,q=A;if(N&&(U=k,V=A,H=u,q=w),!(2&D&&(I=p(t,C,P,I,U,V,B),P>=I)||4&D&&(P=d(t,C,P,I,U,V,F))>=I)){var G=I-P,Y=z-O;if(j){if(t*G*(G+Y)<1<<22){if(void 0!==(M=l.scanComplete(t,C,e,P,I,U,V,O,z,H,q)))return M;continue}}else{if(t*Math.min(G,Y)<128){if(void 0!==(M=o(t,C,e,N,P,I,U,V,O,z,H,q)))return M;continue}if(t*G*Y<1<<22){if(void 0!==(M=l.scanBipartite(t,C,e,N,P,I,U,V,O,z,H,q)))return M;continue}}var W=f(t,C,P,I,U,V,F,B);if(P<W)if(t*(W-P)<128){if(void 0!==(M=s(t,C+1,e,P,W,U,V,O,z,H,q)))return M}else if(C===t-2){if(void 0!==(M=N?l.sweepBipartite(t,e,O,z,H,q,P,W,U,V):l.sweepBipartite(t,e,P,W,U,V,O,z,H,q)))return M}else x(S++,C+1,P,W,O,z,N,-1/0,1/0),x(S++,C+1,O,z,P,W,1^N,-1/0,1/0);if(W<I){var X=c(t,C,O,z,H,q),Z=H[E*X+C],J=h(t,C,X,z,H,q,Z);if(J<z&&x(S++,C,W,I,J,z,(4|N)+(j?16:0),Z,B),O<X&&x(S++,C,W,I,O,X,(2|N)+(j?16:0),F,Z),X+1===J){if(void 0!==(M=j?_(t,C,e,W,I,U,V,X,H,q[X]):b(t,C,e,N,W,I,U,V,X,H,q[X])))return M}else if(X<J){var K;if(j){if(K=m(t,C,W,I,U,V,Z),W<K){var Q=h(t,C,W,K,U,V,Z);if(C===t-2){if(W<Q&&void 0!==(M=l.sweepComplete(t,e,W,Q,U,V,X,J,H,q)))return M;if(Q<K&&void 0!==(M=l.sweepBipartite(t,e,Q,K,U,V,X,J,H,q)))return M}else W<Q&&x(S++,C+1,W,Q,X,J,16,-1/0,1/0),Q<K&&(x(S++,C+1,Q,K,X,J,0,-1/0,1/0),x(S++,C+1,X,J,Q,K,1,-1/0,1/0))}}else K=N?g(t,C,W,I,U,V,Z):m(t,C,W,I,U,V,Z),W<K&&(C===t-2?M=N?l.sweepBipartite(t,e,X,J,H,q,W,K,U,V):l.sweepBipartite(t,e,W,K,U,V,X,J,H,q):(x(S++,C+1,W,K,X,J,N,-1/0,1/0),x(S++,C+1,X,J,W,K,1^N,-1/0,1/0)))}}}}};var n=t("typedarray-pool"),i=t("bit-twiddle"),a=t("./brute"),o=a.partial,s=a.full,l=t("./sweep"),c=t("./median"),u=t("./partition"),f=u("!(lo>=p0)&&!(p1>=hi)"),h=u("lo===p0"),p=u("lo<p0"),d=u("hi<=p0"),m=u("lo<=p0&&p0<=hi"),g=u("lo<p0&&p0<=hi"),v=n.mallocInt32(1024),y=n.mallocDouble(1024);function x(t,e,r,n,i,a,o,s,l){var c=6*t;v[c]=e,v[c+1]=r,v[c+2]=n,v[c+3]=i,v[c+4]=a,v[c+5]=o;var u=2*t;y[u]=s,y[u+1]=l}function b(t,e,r,n,i,a,o,s,l,c,u){var f=2*t,h=l*f,p=c[h+e];t:for(var d=i,m=i*f;d<a;++d,m+=f){var g=o[m+e],v=o[m+e+t];if(!(p<g||v<p)&&(!n||p!==g)){for(var y,x=s[d],b=e+1;b<t;++b){g=o[m+b],v=o[m+b+t];var _=c[h+b],w=c[h+b+t];if(v<_||w<g)continue t}if(void 0!==(y=n?r(u,x):r(x,u)))return y}}}function _(t,e,r,n,i,a,o,s,l,c){var u=2*t,f=s*u,h=l[f+e];t:for(var p=n,d=n*u;p<i;++p,d+=u){var m=o[p];if(m!==c){var g=a[d+e],v=a[d+e+t];if(!(h<g||v<h)){for(var y=e+1;y<t;++y){g=a[d+y],v=a[d+y+t];var x=l[f+y],b=l[f+y+t];if(v<x||b<g)continue t}var _=r(m,c);if(void 0!==_)return _}}}}},{"./brute":105,"./median":107,"./partition":108,"./sweep":110,"bit-twiddle":101,"typedarray-pool":590}],107:[function(t,e,r){"use strict";e.exports=function(t,e,r,a,o,s){if(a<=r+1)return r;var l=r,c=a,u=a+r>>>1,f=2*t,h=u,p=o[f*u+e];for(;l<c;){if(c-l<8){i(t,e,l,c,o,s),p=o[f*u+e];break}var d=c-l,m=Math.random()*d+l|0,g=o[f*m+e],v=Math.random()*d+l|0,y=o[f*v+e],x=Math.random()*d+l|0,b=o[f*x+e];g<=y?b>=y?(h=v,p=y):g>=b?(h=m,p=g):(h=x,p=b):y>=b?(h=v,p=y):b>=g?(h=m,p=g):(h=x,p=b);for(var _=f*(c-1),w=f*h,T=0;T<f;++T,++_,++w){var k=o[_];o[_]=o[w],o[w]=k}var A=s[c-1];s[c-1]=s[h],s[h]=A,h=n(t,e,l,c-1,o,s,p);for(_=f*(c-1),w=f*h,T=0;T<f;++T,++_,++w){k=o[_];o[_]=o[w],o[w]=k}A=s[c-1];if(s[c-1]=s[h],s[h]=A,u<h){for(c=h-1;l<c&&o[f*(c-1)+e]===p;)c-=1;c+=1}else{if(!(h<u))break;for(l=h+1;l<c&&o[f*l+e]===p;)l+=1}}return n(t,e,r,u,o,s,o[f*u+e])};var n=t("./partition")("lo<p0");function i(t,e,r,n,i,a){for(var o=2*t,s=o*(r+1)+e,l=r+1;l<n;++l,s+=o)for(var c=i[s],u=l,f=o*(l-1);u>r&&i[f+e]>c;--u,f-=o){for(var h=f,p=f+o,d=0;d<o;++d,++h,++p){var m=i[h];i[h]=i[p],i[p]=m}var g=a[u];a[u]=a[u-1],a[u-1]=g}}},{"./partition":108}],108:[function(t,e,r){"use strict";e.exports=function(t){return n[t]};var n={"lo===p0":function(t,e,r,n,i,a,o){for(var s=2*t,l=s*r,c=l,u=r,f=e,h=r;n>h;++h,l+=s){if(i[l+f]===o)if(u===h)u+=1,c+=s;else{for(var p=0;s>p;++p){var d=i[l+p];i[l+p]=i[c],i[c++]=d}var m=a[h];a[h]=a[u],a[u++]=m}}return u},"lo<p0":function(t,e,r,n,i,a,o){for(var s=2*t,l=s*r,c=l,u=r,f=e,h=r;n>h;++h,l+=s){if(i[l+f]<o)if(u===h)u+=1,c+=s;else{for(var p=0;s>p;++p){var d=i[l+p];i[l+p]=i[c],i[c++]=d}var m=a[h];a[h]=a[u],a[u++]=m}}return u},"lo<=p0":function(t,e,r,n,i,a,o){for(var s=2*t,l=s*r,c=l,u=r,f=t+e,h=r;n>h;++h,l+=s){if(i[l+f]<=o)if(u===h)u+=1,c+=s;else{for(var p=0;s>p;++p){var d=i[l+p];i[l+p]=i[c],i[c++]=d}var m=a[h];a[h]=a[u],a[u++]=m}}return u},"hi<=p0":function(t,e,r,n,i,a,o){for(var s=2*t,l=s*r,c=l,u=r,f=t+e,h=r;n>h;++h,l+=s){if(i[l+f]<=o)if(u===h)u+=1,c+=s;else{for(var p=0;s>p;++p){var d=i[l+p];i[l+p]=i[c],i[c++]=d}var m=a[h];a[h]=a[u],a[u++]=m}}return u},"lo<p0&&p0<=hi":function(t,e,r,n,i,a,o){for(var s=2*t,l=s*r,c=l,u=r,f=e,h=t+e,p=r;n>p;++p,l+=s){var d=i[l+f],m=i[l+h];if(d<o&&o<=m)if(u===p)u+=1,c+=s;else{for(var g=0;s>g;++g){var v=i[l+g];i[l+g]=i[c],i[c++]=v}var y=a[p];a[p]=a[u],a[u++]=y}}return u},"lo<=p0&&p0<=hi":function(t,e,r,n,i,a,o){for(var s=2*t,l=s*r,c=l,u=r,f=e,h=t+e,p=r;n>p;++p,l+=s){var d=i[l+f],m=i[l+h];if(d<=o&&o<=m)if(u===p)u+=1,c+=s;else{for(var g=0;s>g;++g){var v=i[l+g];i[l+g]=i[c],i[c++]=v}var y=a[p];a[p]=a[u],a[u++]=y}}return u},"!(lo>=p0)&&!(p1>=hi)":function(t,e,r,n,i,a,o,s){for(var l=2*t,c=l*r,u=c,f=r,h=e,p=t+e,d=r;n>d;++d,c+=l){var m=i[c+h],g=i[c+p];if(!(m>=o||s>=g))if(f===d)f+=1,u+=l;else{for(var v=0;l>v;++v){var y=i[c+v];i[c+v]=i[u],i[u++]=y}var x=a[d];a[d]=a[f],a[f++]=x}}return f}}},{}],109:[function(t,e,r){"use strict";e.exports=function(t,e){e<=128?n(0,e-1,t):function t(e,r,u){var f=(r-e+1)/6|0,h=e+f,p=r-f,d=e+r>>1,m=d-f,g=d+f,v=h,y=m,x=d,b=g,_=p,w=e+1,T=r-1,k=0;l(v,y,u)&&(k=v,v=y,y=k);l(b,_,u)&&(k=b,b=_,_=k);l(v,x,u)&&(k=v,v=x,x=k);l(y,x,u)&&(k=y,y=x,x=k);l(v,b,u)&&(k=v,v=b,b=k);l(x,b,u)&&(k=x,x=b,b=k);l(y,_,u)&&(k=y,y=_,_=k);l(y,x,u)&&(k=y,y=x,x=k);l(b,_,u)&&(k=b,b=_,_=k);for(var A=u[2*y],M=u[2*y+1],S=u[2*b],E=u[2*b+1],L=2*v,C=2*x,P=2*_,I=2*h,O=2*d,z=2*p,D=0;D<2;++D){var R=u[L+D],F=u[C+D],B=u[P+D];u[I+D]=R,u[O+D]=F,u[z+D]=B}a(m,e,u),a(g,r,u);for(var N=w;N<=T;++N)if(c(N,A,M,u))N!==w&&i(N,w,u),++w;else if(!c(N,S,E,u))for(;;){if(c(T,S,E,u)){c(T,A,M,u)?(o(N,w,T,u),++w,--T):(i(N,T,u),--T);break}if(--T<N)break}s(e,w-1,A,M,u),s(r,T+1,S,E,u),w-2-e<=32?n(e,w-2,u):t(e,w-2,u);r-(T+2)<=32?n(T+2,r,u):t(T+2,r,u);T-w<=32?n(w,T,u):t(w,T,u)}(0,e-1,t)};function n(t,e,r){for(var n=2*(t+1),i=t+1;i<=e;++i){for(var a=r[n++],o=r[n++],s=i,l=n-2;s-- >t;){var c=r[l-2],u=r[l-1];if(c<a)break;if(c===a&&u<o)break;r[l]=c,r[l+1]=u,l-=2}r[l]=a,r[l+1]=o}}function i(t,e,r){e*=2;var n=r[t*=2],i=r[t+1];r[t]=r[e],r[t+1]=r[e+1],r[e]=n,r[e+1]=i}function a(t,e,r){e*=2,r[t*=2]=r[e],r[t+1]=r[e+1]}function o(t,e,r,n){e*=2,r*=2;var i=n[t*=2],a=n[t+1];n[t]=n[e],n[t+1]=n[e+1],n[e]=n[r],n[e+1]=n[r+1],n[r]=i,n[r+1]=a}function s(t,e,r,n,i){e*=2,i[t*=2]=i[e],i[e]=r,i[t+1]=i[e+1],i[e+1]=n}function l(t,e,r){e*=2;var n=r[t*=2],i=r[e];return!(n<i)&&(n!==i||r[t+1]>r[e+1])}function c(t,e,r,n){var i=n[t*=2];return i<e||i===e&&n[t+1]<r}},{}],110:[function(t,e,r){"use strict";e.exports={init:function(t){var e=i.nextPow2(t);o.length<e&&(n.free(o),o=n.mallocInt32(e));s.length<e&&(n.free(s),s=n.mallocInt32(e));l.length<e&&(n.free(l),l=n.mallocInt32(e));c.length<e&&(n.free(c),c=n.mallocInt32(e));u.length<e&&(n.free(u),u=n.mallocInt32(e));f.length<e&&(n.free(f),f=n.mallocInt32(e));var r=8*e;h.length<r&&(n.free(h),h=n.mallocDouble(r))},sweepBipartite:function(t,e,r,n,i,u,f,m,g,v){for(var y=0,x=2*t,b=t-1,_=x-1,w=r;w<n;++w){var T=u[w],k=x*w;h[y++]=i[k+b],h[y++]=-(T+1),h[y++]=i[k+_],h[y++]=T}for(w=f;w<m;++w){T=v[w]+(1<<28);var A=x*w;h[y++]=g[A+b],h[y++]=-T,h[y++]=g[A+_],h[y++]=T}var M=y>>>1;a(h,M);var S=0,E=0;for(w=0;w<M;++w){var L=0|h[2*w+1];if(L>=1<<28)p(l,c,E--,L=L-(1<<28)|0);else if(L>=0)p(o,s,S--,L);else if(L<=-(1<<28)){L=-L-(1<<28)|0;for(var C=0;C<S;++C){if(void 0!==(P=e(o[C],L)))return P}d(l,c,E++,L)}else{L=-L-1|0;for(C=0;C<E;++C){var P;if(void 0!==(P=e(L,l[C])))return P}d(o,s,S++,L)}}},sweepComplete:function(t,e,r,n,i,m,g,v,y,x){for(var b=0,_=2*t,w=t-1,T=_-1,k=r;k<n;++k){var A=m[k]+1<<1,M=_*k;h[b++]=i[M+w],h[b++]=-A,h[b++]=i[M+T],h[b++]=A}for(k=g;k<v;++k){A=x[k]+1<<1;var S=_*k;h[b++]=y[S+w],h[b++]=1|-A,h[b++]=y[S+T],h[b++]=1|A}var E=b>>>1;a(h,E);var L=0,C=0,P=0;for(k=0;k<E;++k){var I=0|h[2*k+1],O=1&I;if(k<E-1&&I>>1==h[2*k+3]>>1&&(O=2,k+=1),I<0){for(var z=-(I>>1)-1,D=0;D<P;++D){if(void 0!==(R=e(u[D],z)))return R}if(0!==O)for(D=0;D<L;++D){if(void 0!==(R=e(o[D],z)))return R}if(1!==O)for(D=0;D<C;++D){var R;if(void 0!==(R=e(l[D],z)))return R}0===O?d(o,s,L++,z):1===O?d(l,c,C++,z):2===O&&d(u,f,P++,z)}else{z=(I>>1)-1;0===O?p(o,s,L--,z):1===O?p(l,c,C--,z):2===O&&p(u,f,P--,z)}}},scanBipartite:function(t,e,r,n,i,l,c,u,f,m,g,v){var y=0,x=2*t,b=e,_=e+t,w=1,T=1;n?T=1<<28:w=1<<28;for(var k=i;k<l;++k){var A=k+w,M=x*k;h[y++]=c[M+b],h[y++]=-A,h[y++]=c[M+_],h[y++]=A}for(k=f;k<m;++k){A=k+T;var S=x*k;h[y++]=g[S+b],h[y++]=-A}var E=y>>>1;a(h,E);var L=0;for(k=0;k<E;++k){var C=0|h[2*k+1];if(C<0){var P=!1;if((A=-C)>=1<<28?(P=!n,A-=1<<28):(P=!!n,A-=1),P)d(o,s,L++,A);else{var I=v[A],O=x*A,z=g[O+e+1],D=g[O+e+1+t];t:for(var R=0;R<L;++R){var F=o[R],B=x*F;if(!(D<c[B+e+1]||c[B+e+1+t]<z)){for(var N=e+2;N<t;++N)if(g[O+N+t]<c[B+N]||c[B+N+t]<g[O+N])continue t;var j,U=u[F];if(void 0!==(j=n?r(I,U):r(U,I)))return j}}}}else p(o,s,L--,C-w)}},scanComplete:function(t,e,r,n,i,s,l,c,u,f,p){for(var d=0,m=2*t,g=e,v=e+t,y=n;y<i;++y){var x=y+(1<<28),b=m*y;h[d++]=s[b+g],h[d++]=-x,h[d++]=s[b+v],h[d++]=x}for(y=c;y<u;++y){x=y+1;var _=m*y;h[d++]=f[_+g],h[d++]=-x}var w=d>>>1;a(h,w);var T=0;for(y=0;y<w;++y){var k=0|h[2*y+1];if(k<0){if((x=-k)>=1<<28)o[T++]=x-(1<<28);else{var A=p[x-=1],M=m*x,S=f[M+e+1],E=f[M+e+1+t];t:for(var L=0;L<T;++L){var C=o[L],P=l[C];if(P===A)break;var I=m*C;if(!(E<s[I+e+1]||s[I+e+1+t]<S)){for(var O=e+2;O<t;++O)if(f[M+O+t]<s[I+O]||s[I+O+t]<f[M+O])continue t;var z=r(P,A);if(void 0!==z)return z}}}}else{for(x=k-(1<<28),L=T-1;L>=0;--L)if(o[L]===x){for(O=L+1;O<T;++O)o[O-1]=o[O];break}--T}}}};var n=t("typedarray-pool"),i=t("bit-twiddle"),a=t("./sort"),o=n.mallocInt32(1024),s=n.mallocInt32(1024),l=n.mallocInt32(1024),c=n.mallocInt32(1024),u=n.mallocInt32(1024),f=n.mallocInt32(1024),h=n.mallocDouble(8192);function p(t,e,r,n){var i=e[n],a=t[r-1];t[i]=a,e[a]=i}function d(t,e,r,n){t[r]=n,e[n]=r}},{"./sort":109,"bit-twiddle":101,"typedarray-pool":590}],111:[function(t,e,r){},{}],112:[function(t,e,r){(function(e){(function(){
/*!
 * The buffer module from node.js, for the browser.
 *
 * @author   Feross Aboukhadijeh <https://feross.org>
 * @license  MIT
 */
```
