## Supplementary material for "Spatial omics representation and functional tissue module inference using graph Fourier transform": Supp Note 1

**Section C1. A theoretical formulation for TM identification**

**C1.1 Rationale.** In spatially resolved transcriptomics (SRT) studies, a tissue module (TM) can be defined as a region or a set of discontinuous regions that are correlated with a group of SVGs sharing similar spatial distribution. The entity of a TM on SRT data is a subset of spots whose underlying features will support the spatial pattern and biological function of a TM. However, both the number of TMs and the entity of TMs are unknown. Therefore, in SpaGFT, we will identify a set of TMs that can cover the convoluted biological processes of a tissue of interest.

**C1.2 Theoretical foundation.** Let $S$ be the set of spots in an SRT dataset and equal the union of all TMs within the tissue. Herein, $S$is also defined as the universe for spots in all possible TMs. A collection $\mathcal{F}$ of $m$ spot sets whose union equals the universe $S$ and sets in $\mathcal{F}$ satisfy: (i) spots in each set have a spatial pattern supported by a group of SVGs sharing similar spatial distribution; (ii) each set is a potential TM which could be spatially discontinuous and have a specific biological function; and (iii) TMs may collaborate and form a higher-order functional tissue motif. The task that identifies TMs can be converted to find a sub-collection$\mathcal{F}^{'}$ of $\mathcal{F,}$ which satisfies: 1) the union of sets in $\mathcal{F}^{'}$ equals to the union of sets in $\mathcal{F}$; 2) $\mathcal{F}^{'}$ has the minimum cardinality, that is, $\mathcal{|F}^{'}|$ is the smallest. Such a task is a special case to a classical set covering problems in combinations, computer science, and operations research, which is one of Karp's 21 $\mathrm{NP}$-$complete$^1^.

**C1.3 Computational formulation.** Due to the set covering problems being $\mathrm{NP}$-$complete$, we propose SpaGFT as a heuristic algorithm to identify TMs by (*i*) identifying SVGs using the graph signal processing approach with a well theoretical foundation and (*ii*) predict TMs based on SVG clustering and optimizing the number of TMs through minimizing the overlapping spots among TMs. We will introduce SVG identification and evaluation in **Section C2** and SVG clustering and TMs optimization in **Section C3.**

**Section C2. SVG identification and evaluation using graph signal processing approach**

**C2.1 SVG identification**

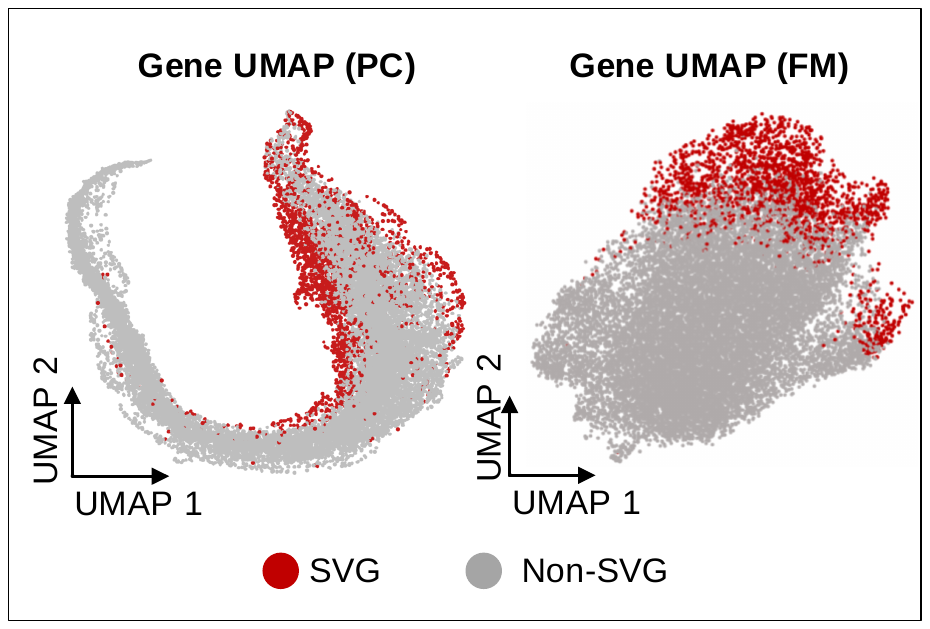

**Figure I**. Comparison of the UMAPs obtained using the top 207 principal components (PCs) (left) and the top 207 FMs (right) of the Mouse Visium data (HE-coronal, 2702 spots). The principal component analysis (PCA) dimensions were generated directly from the gene-spot expression matrix using PCA analysis in Scanpy. Red dots indicate the 2,118 SVGs identified by SpaGFT using the default settings and the other genes are represented by the grey color.

**C2.1.1 Rationale.** Graph Fourier transform can transform gene expression value and corresponding spatial patterns into simple but informative features in a new feature space. New features in such a space can make SVGs and non-SVGs more distinguishable compared to other feature spaces, such as principal components (PC) in the PCA feature space (**Fig. I**).

**C2.1.2 Theoretical foundation.** With the support of the theoretical foundation of graph Fourier transform^2-4^, SVG can be formulated as an approximate *k*-bandlimited graph signal, which is a smooth signal and can be represented as the linear combination of the first *k* Fourier mode (FMs). Such first k FMs usually refer to low-frequency FMs (**Fig. IIA**) rather than all or high-frequency FMs (**Fig. IIB**). We refer to the method for determining high-variant PCs from PCA analysis^5^. The determination of low-frequency and high-frequency FMs can be achieved by finding the inflection point based on the eigenvalue of the Laplacian matrix (**Fig. IIC**). Meanwhile, the contribution of FMs to SVG expression patterns can be measured by the corresponding Fourier coefficients (FCs). If a gene is an SVG, the low-frequency FCs usually have a larger value compared to high-frequency FC. Herein, SVG identification can be converted to the recognition of the *k*-bandlimited signal problem.

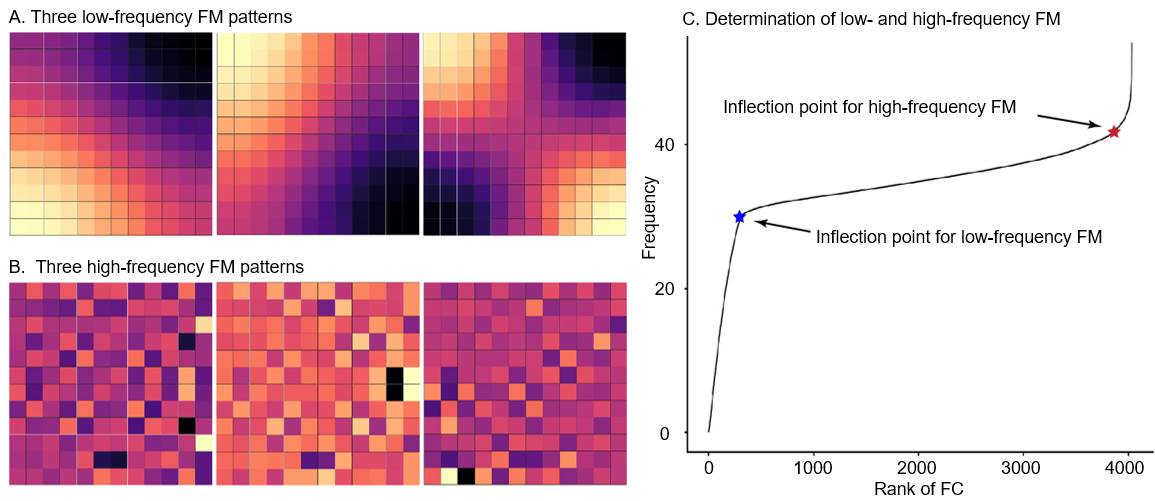

**Fig II**. **A and B demonstrate the low-frequency and high-frequency FM spatial patterns**. Low-frequency FMs refer to slow variation and can be used for recovering *k*-bandlimited graph signal. High-frequency FMs refer to rapid variation and represents noisy patterns. C. The screen plot shows the rank of eigenvalue (the x-axis) and corresponding values (the y-axis). The low-frequency FM can be determined by the first inflection point (blue star) and the high-frequency FM can be determined by the second inflection point (red star).

**C2.1.3 Computational formulation.** Given a graph $G=\left( V,E \right)$, where $V$ is the node set with $\left| V \right|=n$, $n$ is the number of spots, and $E$ is the edge set added based on the Euclidean distance, let $\boldsymbol{\mu}_{1}, \boldsymbol{\mu}_{2},\ldots,\boldsymbol{\mu}_{k}\in\mathbb{R}^{n}$be the first $k$ low-frequency FMs. For the expression values of a gene on $n$ spots $\boldsymbol{f}\in\mathbb{R}^{n}, \hat{f}_{i}=\left\langle\boldsymbol{\mu}_{i}, \right.\left. \boldsymbol{f} \right\rangle$is the graph FC to describe the contribution of $\boldsymbol{\mu}_{i}$ to recover $\boldsymbol{f}$, where $\left\langle\cdot\right.,\left. \cdot\right\rangle$ represents the dot product and $i=1, 2, \ldots, k.$ For a fixed error $\varepsilon>0$, a gene will be considered as an SVG if the corresponding $\boldsymbol{f}$ satisfies $\left\| \boldsymbol{f}-\sum_{i}^{k} \hat{f}_{i}\boldsymbol{\mu}_{i} \right\|<\varepsilon.$

**C2.2 SVG evaluation I: GFTscore**

**C2.2.1 Rationale of GFTscore.** Although we formulate the identification of SVG as the recognization of the *k*-bandlimited graph signal problem, characterizing the *k* FMs for *k*-bandlimited graph signals is challenging. Therefore, we plan to design the GFTscore to represent the contribution of first *k* FMs to the gene expression pattern. We expect a gene with a high GFTscore tends to be an SVG, and a gene with a low GFTscore tends to be a non-SVG.

**C2.2.2 Theoretical foundation of GFTscore.** The contribution of FMs can be measured by the FC^6^, which ranges from $\left[ -\infty,+\infty\right]$. To make FC comparable among different genes, FC will be normalized to [0,1]. Furthermore, as GFTscore aims to enhance the low-frequency FM contribution across all FMs, we refer to the low pass filter processing approach^4^ and assign different weights calculated from frequency to all FC. Specifically, low-frequency FC will be assigned a high weight, and high-frequency will be assigned a low weight. Therefore, SVG tends to have a high GFTscore, and non-SVG tends to show a low GFTscore in this definition.

**C2.2.3 Computational formulation of GFTscore.** We designed a GFTscore to quantitatively measure the randomness of gene expressions distributed, defined as

$$GFTscore\left( f_{g} \right)=\sum_{k=1}^{n} e^{-\lambda_{k}}\tilde{f}_{g}^{k},$$

where $\lambda_{k}$ is the pre-calculated eigenvalue of $\boldsymbol{L}$, and the normalized frequency signal $\tilde{f}_{g}^{k}$ is defined as:

$$\tilde{f}_{g}^{k}=\frac{{|\hat{f}}_{g}^{k}|}{\sum_{i=1}^{n} {|\hat{f}}_{g}^{i}|}.$$

The inflection point of GFTscore can be determined by the Kneedle algorithm^7^. According to a recent study^4^, the weight ($e^{-\lambda_{k}}$) of FC was one of common forms in grapg signal processing.

**C2.3 SVG evaluation II: hypothesis testing**

**C2.3.1 Rationale of hypothesis testing.** Although the above GFTscore is an indicator to rank and evaluate the potential SVGs, a rigorous statistical test is needed to calculate the *p*-value for SVGs and control type I error. We will use the Wilcoxon rank-sum test to test the differences between low-frequency FCs and high-frequency FCs to obtain statistical significance. If a gene has a high GFTscore and significantly adjusted *p*-value, the gene can be regarded as an SVG.

**C2.3.2 The null hypothesis of testing.** We use $x_{1},x_{2},\ldots,x_{n}$ to represent the expression of a random signal on $n$ spots. $x_{i}$ is followed by Gaussian distributions and can be regarded as independent and identically distributed (*i.i.d.*) random variables^8^. Implementing GFT on $\left( x_{1},x_{2},\ldots,x_{n} \right)$, we obtain $FC_{1}, FC_{2}, \cdots, FC_{p}$, where $p$ is the number of low-frequency FCs and reflects the contributions from low-frequency FMs. We also obtain the $FC_{p+1}, FC_{p+2}, \cdots, FC_{p+q}$ , where $q$ is the number of high-frequency FCs and reflects the contributions from noise. Hence, we form the null hypothesis that there exists no difference between low-frequency FCs and high-frequency FCs.

**C2.3.3 Proof of the null hypothesis.** Give a signal $\boldsymbol{x}={(x}_{1},x_{2},\ldots,x_{n}), where$ $x_{1},x_{2},\ldots,x_{n}$ are *i.i.d* random variables*.* Given any two FMs (corresponding to any non-zero eigenvalues), $\boldsymbol{u}_{k}=\left( u_{k}^{1},u_{k}^{2},\ldots,u_{k}^{n} \right)$ and $\boldsymbol{u}_{l}=\left( u_{l}^{1},u_{l}^{2},\ldots,u_{l}^{n} \right)$. According to the property of the Laplacian matrix defined in SpaGFT, (i) both$\boldsymbol{u}_{1}$ and $\boldsymbol{u}_{2}$ are orthogonal with vector $\left( 1,1,\ldots,1 \right)$, and one has that $\sum_{i}^{n} u_{k}^{i}=\sum_{i}^{n} u_{l}^{i}$*;* (ii) $\sum_{i}^{n} {{(u}_{k}^{i})}^{2}=\sum_{i}^{n} {{(u}_{l}^{i})}^{2}$ as they are both unit vectors. The $FC_{k}=\sum_{i}^{n} u_{k}^{i}x_{i}$ and $FC_{l}=\sum_{i}^{n} u_{l}^{i}x_{i}$ are two FCs corresponding to two FMs, respectively. In the following, we can observe two properties, including (i)$E\left( FC_{k} \right)= E\left( \sum_{i}^{n} u_{k}^{i}x_{i} \right)=\sum_{i}^{n} {[u}_{k}^{i}E\left( x_{i} \right)]=E\left( x_{1} \right)\sum_{i}^{n} u_{k}^{i}=E\left( x_{1} \right)\sum_{i}^{n} u_{l}^{i}=E\left( FC_{l} \right)$; (ii) $Var\left( FC_{k} \right)= Var\left( \sum_{i}^{n} u_{k}^{i}x_{i} \right)=\sum_{i}^{n} {{[(u}_{k}^{i})}^{2}Var\left( x_{i} \right)]=Var\left( x_{1} \right)\sum_{i}^{n} {{(u}_{k}^{i})}^{2}=Var\left( x_{1} \right)\sum_{i}^{n} {{(u}_{l}^{i})}^{2}=Var\left( FC_{l} \right)$. All FCs have the same mean and variance regardless of low-frequency and high-frequency. Therefore, there exists no difference between FCs corresponding to low-frequency FMs and FCs corresponding to high-frequency FMs under the null hypothesis.

**Section C3. SVG clustering and TMs optimization**

**C3.1 Rationale.** According to the TM definition, a group of SVGs sharing similar spatial distribution can support the pattern and function of a TM. However, the number and shape of TMs in tissue are unknown. Therefore, we propose a global optimization approach, which aims to find the best Louvain resolution to produce sufficient TMs and minimize overlapped spots among those TMs.

**C3.2 Theoretical foundation.** SVGs with similar patterns also have similar low-frequency FCs, which provides a fundamental basis of clustering as spectral clustering^9^. By performing the Louvain clustering algorithm, SVG with a similar low-frequency FC will be grouped and produce SVG clusters. The shared spatial pattern of an SVG cluster can be represented by (*i*) summing up the expression values of those SVGs using iGFT and (*ii*) binarizing summed expression values using k-means (set $k=2$). To determine the best number of TMs, we propose optimizing the resolution parameter of the Louvain algorithm using an objective function to minimize overlapped spots among identified TMs.

**C3.3 Computational formulation.** Denote $G^{'}$ as the set of SVGs identified by SpaGFT. For each resolution parameter $res>0$, $G^{'}$ can be partitioned to {$G_{1}^{'},G_{2}^{'},\ldots,G_{n_{res}}^{'}\}$ (i.e., $\bigcup_{k} G_{i}^{'}=G^{'}$ and $G_{k}^{'}\bigcap G_{l}^{'}=\emptyset, \forall k\neq l.$) by applying the Louvain algorithm on SVGs' signals in the frequency domain.Denote $X=(x_{s,g})\in\mathbb{R}^{\left| S \right|\times\left| G^{'} \right|}$ as the gene expression matrix, where $S$ is the set of all spots. In the following, for each SVG group $G_{k}^{'}$, $pseudo(s_{s,G_{k}^{'}})=\sum_{g\in G_{k}^{'}} log(x_{s,g})$ represents the pseudo expression value for spot $i$. Apply k-means algorithms with k=2 on $\{pseudo(s_{1,G_{k}^{'}}),pseudo(s_{2,G_{k}^{'}}), \ldots, pseudo(s_{\left| S \right|,G_{k}^{'}})\}$ to pick out one spot cluster whose spots highly express genes in SVG group $G_{k}^{'}$ and such spot cluster is identified as a TM, denoted as $S_{i}\in S$. Our objective function aims to find the best partition of $G^{'}$ such that the average overlap between any two $S_{i}, S_{j}$ is minimized:

$$\underset{res>0}{argmin} \frac{2\times\sum_{k\neq l} \left| S_{k}\cap S_{l} \right|}{n_{res}\times(n_{res}-1)}$$

1	Karp, R. M. in *Complexity of Computer Computations: Proceedings of a symposium on the Complexity of Computer Computations, held March 20–22, 1972, at the IBM Thomas J. Watson Research Center, Yorktown Heights, New York, and sponsored by the Office of Naval Research, Mathematics Program, IBM World Trade Corporation, and the IBM Research Mathematical Sciences Department*	(eds Raymond E. Miller, James W. Thatcher, & Jean D. Bohlinger) 85-103 (Springer US, 1972).

2	Segarra, S., Marques, A. G., Leus, G. & Ribeiro, A. Reconstruction of graph signals through percolation from seeding nodes. *IEEE Transactions on Signal Processing* **64**, 4363-4378 (2016).

3	Huang, L., Needell, D. & Tang, S. Robust recovery of bandlimited graph signals via randomized dynamical sampling. *arXiv preprint arXiv:2109.14079* (2021).

4	Ricaud, B., Borgnat, P., Tremblay, N., Gonçalves, P. & Vandergheynst, P. Fourier could be a data scientist: From graph Fourier transform to signal processing on graphs. *Comptes Rendus Physique* **20**, 474-488, doi:<https://doi.org/10.1016/j.crhy.2019.08.003> (2019).

5	Cattell, R. B. The scree test for the number of factors. *Multivariate behavioral research* **1**, 245-276 (1966).

6	Puy, G., Tremblay, N., Gribonval, R. & Vandergheynst, P. Random sampling of bandlimited signals on graphs. *Applied and Computational Harmonic Analysis* **44**, 446-475 (2018).

7	Satopaa, V., Albrecht, J., Irwin, D. & Raghavan, B. in *2011 31st international conference on distributed computing systems workshops.* 166-171 (IEEE).

8	Svensson, V., Teichmann, S. A. & Stegle, O. SpatialDE: identification of spatially variable genes. *Nature Methods* **15**, 343-346, doi:10.1038/nmeth.4636 (2018).

9	Ng, A., Jordan, M. & Weiss, Y. On spectral clustering: Analysis and an algorithm. *Advances in neural information processing systems* **14** (2001).
