## Supplementary material for "Spatial omics representation and functional tissue module inference using graph Fourier transform": Supp Note 2

**Gene function and tissue module (TM) annotation in benchmarking and case studies.**

**Annotation 1: SpaGFT unique identified gene**

*Nsmf* and *Tbr1* were identified by all six tools and recognized as cell-type-specific genes in the hippocampus, cortical region, and cerebral cortex^1-3^ (**Fig. 2f)**. In addition, SVGs uniquely identified by SpaGFT are also proven to be associated with specific brain region and functions, e.g., *Cartpt* (mainly expressed in the ventral tegmental area and involved in dopamine metabolism^4^), *Cbln2* (expressed in the cortical plate of the developing neocortex and displayed a high rostromedial to low caudolateral gradient^5^), *Ttr* (protective effect against the amyloidogenic peptide^6^), and *Pmch* (regulated distinct aspects of energy balance and vital behavior^7,8^) (**Supplementary Fig. 3**).

**Annotation 2: SpaGFT enhanced low-expression gene and removed noise background**

SpaGFT removed noisy background for the *Thra* gene, broadly expressed by the neural cell and detected across all brain structures^9^. On the contrary, for low expression genes (e.g., *Ano2* in the central lateral amygdala, the later septal nucleus, and olfactory sensory neurons^10,11^), SpaGFT can improve the signal magnitude (**Fig. 2i**).

**Annotation 3: TM 6 interpretation.**

*Ahi1* is most highly expressed in the brain, particularly in neurons from superior cerebellar peduncles^12^. The enrichment of ontologies (e.g., in SVG function enrichments section), KEGG pathways, and Reactome database (**Supplementary Fig. 5c**) was also performed for TM-associated SVGs to elucidate the underlying functions of the TM 6. TM 6 appears to be a module of regulatory neurons characteristic of the hypothalamus and amygdala, with gene set enrichments related to opioid receptors and GABA production and signaling. Moreover, the TM ID Card also showed the overlapped region with other TMs. For example, TM 8 also appears the most hypothalamic, highlighting anti-inflammatory activity, astrocytes, and regulatory neuronal circuits (**Supplementary Data 1**). TMs 6 and 8 overlapped into the hypothalamus, amygdala, and striatum, suggesting a high-order function region (e.g., emotion control for joy and fear^13^).

**Annotation 4: Tissue motif case interpretation.**

TM clusters 1 and 2 indicated the convolution of elements, reflecting a potential collaboration between the white matter and the hypothalamus region^14^. TM cluster 3 (enriched with excitatory neurons) also showed a strong association with TM cluster 2 (enriched with inhibitory neurons), indicating the neuronal circuit activity of inhibitory and excitatory neurons in the hypothalamus region^15,16^. In addition, TM cluster 3 displayed possible collaborations with either TM cluster 1 or 2, indicating potential connectivity among the partial cerebrum, hypothalamus region, and white matter^14,17^.

**Annotation 5: Three TM definitions in lymph node**

TM 3 (T cell zone) was highly correlated with eight T cell-related cell types (**Fig. 4b**). In addition, this TM pattern was supported by 177 SVGs, including several T cell zone markers relevant to T cell survival, such as *CD3E, IL7R, CCR7, and CCL19*^18^. The pathway analysis results showed the functional enrichment of 177 SVGs, which were positively enriched in the T cell activation and TCR signal pathway (**Supplementary Fig. 7b**). TM 5 was highly correlated with six cell2location cell types associated with the GC, including T follicular helper cells (T_CD4_TfH_GC), follicular dendritic cells (FDC), pre-GC B cells (B_GC_prePB), cycling B cells (B_cycling), dark zone B cells (B_GC-DZ), and light zone B cells (B_GC-LZ). SpaGFT identified 142 SVGs associated with TM 5, including *PCNA*, *CDK1*, and *CDC20*, which have previously been described as marker genes of cell proliferation and cell cycle pathway enrichment in the GC^19,20^ (**Supplementary Figure 7c**). TM 7 was highly correlated with five B cell types and associated with 95 SVGs. Contained in these SVGs were B cell markers (e.g., *CD19, CD79B, and CR2*) and pathway enrichments relevant to B cell activities (e.g., antigen processing and presentation) (**Supplementary Fig. 7d**)^21^. Altogether, we defined TMs 3, 5, and 7 as the T cell zone, GC, and B follicle, respectively.

**Annotation 6: Interpretation of other interaction regions in lymph node**

We observed that the antibody-secreting cell (ASC) pathway was enriched in the B follicle (**Fig. 4e**). Meanwhile, naïve, activated, and memory B cells were also enriched in the same region (**Fig. 4f** and **Supplementary Fig. 8**). The observation primarily demonstrated B follicles contained a large proportion of ASC for producing antibodies and memory B cells for quickly responding to antigens upon recall^22^. We also observed that GC formation-relevant pathways and cell types were enriched in the GC-B overlapping region. For example, several pathways, including GC formation, B cell proliferation, and T helper regulation in GC, showed a relatively high activity in the GC-B region^23-25^. Notably, the proportion of B cycling cells, light-zone specific B cells, follicular dendritic cells (FDC), and follicular T helper (TFH) also show increased abundance in the GC-B region. Furthermore, we investigated T cell functions in the T zone, which displayed enrichment of CD4 T cell markers, CD4 and CD5^26^, as well as a T cell activation gene module enrichment (**Supplementary Fig. 8**). Interestingly, the T cell migration pathway displayed high activity in B-T and tri-zone (i.e., B-T-GC), indicating that T cells will migrate to B follicle and GC to interact with B cells^27,28^.

**Annotation 7: Interpretation of other interaction regions in lymph node**

Based on the previous study, finer structures of the mature follicle in the human tonsil^29^, such as the light zone, were captured by high-resolution spatial-CITE-seq (~25$\mu$m per pixel size). Therefore, we mainly targeted functions within TMs relevant to follicles. As a results, we identified follicle-relevant TMs based on the protein signature and TM morphology. For example, CD9 and CD63 were found in the GC-crypt region (**Fig. 5g** and **Supplementary Table 18**), which were tetraspanins expressed in tonsillar B cells in both follicles and crypts (**Fig. 4g**), indicating a potential B cell migration zone^29,30^. CD23 and CD55 were found in the GC light zone^29^. Complement receptors CD21 and CD35 were found in whole follicles (**Fig. 4i**), indicating the existence of mature B cells and follicular dendritic cells in whole follicles^31,32^. Subsequently, we investigated the two overlapped regions, including (1) whole follicles and GC-light zone and (2) whole follicles and GC-crypt. It was well-known that the GC-light zone was a substructure of the whole follicle and executed the B cell selection function based on follicular helper T cells and follicular dendritic cells (FDCs) activity^33-36^. In another case of other overlapped regions between the whole follicle and GC-crypt, we found a potential B cell migration and spread between follicle and GC-crypt regions which might be modulated by tetraspanins such as CD9 and CD63 expressed on B cell membrane^30^.

1	Joglekar, A. *et al.* A spatially resolved brain region- and cell type-specific isoform atlas of the postnatal mouse brain. *Nature Communications* **12**, 463, doi:10.1038/s41467-020-20343-5 (2021).

2	Spilker, C. *et al.* A Jacob/Nsmf Gene Knockout Results in Hippocampal Dysplasia and Impaired BDNF Signaling in Dendritogenesis. *PLoS Genet* **12**, e1005907, doi:10.1371/journal.pgen.1005907 (2016).

3	Huang, T.-N. *et al.* Tbr1 haploinsufficiency impairs amygdalar axonal projections and results in cognitive abnormality. *Nature Neuroscience* **17**, 240-247, doi:10.1038/nn.3626 (2014).

4	Carpenter, M. D. *et al.* Nr4a1 suppresses cocaine-induced behavior via epigenetic regulation of homeostatic target genes. *Nature Communications* **11**, 504, doi:10.1038/s41467-020-14331-y (2020).

5	Miura, E., Iijima, T., Yuzaki, M. & Watanabe, M. Distinct expression of Cbln family mRNAs in developing and adult mouse brains. *European Journal of Neuroscience* **24**, 750-760, doi:<https://doi.org/10.1111/j.1460-9568.2006.04950.x> (2006).

6	Buxbaum, J. N. *et al.* Transthyretin protects Alzheimer's mice from the behavioral and biochemical effects of A&#x3b2; toxicity. *Proceedings of the National Academy of Sciences* **105**, 2681-2686, doi:doi:10.1073/pnas.0712197105 (2008).

7	Segal-Lieberman, G. *et al.* Melanin-concentrating hormone is a critical mediator of the leptin-deficient phenotype. *Proc Natl Acad Sci U S A* **100**, 10085-10090, doi:10.1073/pnas.1633636100 (2003).

8	González, J. A., Iordanidou, P., Strom, M., Adamantidis, A. & Burdakov, D. Awake dynamics and brain-wide direct inputs of hypothalamic MCH and orexin networks. *Nature Communications* **7**, 11395, doi:10.1038/ncomms11395 (2016).

9	Bernal, J. Thyroid hormone receptors in brain development and function. *Nat Clin Pract Endocrinol Metab* **3**, 249-259, doi:10.1038/ncpendmet0424 (2007).

10	Stephan, A. B. *et al.* ANO2 is the cilial calcium-activated chloride channel that may mediate olfactory amplification. *Proceedings of the National Academy of Sciences* **106**, 11776-11781 (2009).

11	Li, K. X. *et al.* TMEM16B regulates anxiety-related behavior and GABAergic neuronal signaling in the central lateral amygdala. *Elife* **8**, doi:10.7554/eLife.47106 (2019).

12	Ferland, R. J. *et al.* Abnormal cerebellar development and axonal decussation due to mutations in AHI1 in Joubert syndrome. *Nature Genetics* **36**, 1008-1013, doi:10.1038/ng1419 (2004).

13	Koelsch, S. & Skouras, S. Functional centrality of amygdala, striatum and hypothalamus in a "small-world" network underlying joy: an fMRI study with music. *Hum Brain Mapp* **35**, 3485-3498, doi:10.1002/hbm.22416 (2014).

14	Lemaire, J.-J. *et al.* White matter connectivity of human hypothalamus. *Brain Research* **1371**, 43-64, doi:<https://doi.org/10.1016/j.brainres.2010.11.072> (2011).

15	Belousov, A. B., O'Hara, B. F. & Denisova, J. V. Acetylcholine becomes the major excitatory neurotransmitter in the hypothalamus in vitro in the absence of glutamate excitation. *J Neurosci* **21**, 2015-2027, doi:10.1523/jneurosci.21-06-02015.2001 (2001).

16	Mickelsen, L. E. *et al.* Single-cell transcriptomic analysis of the lateral hypothalamic area reveals molecularly distinct populations of inhibitory and excitatory neurons. *Nature Neuroscience* **22**, 642-656, doi:10.1038/s41593-019-0349-8 (2019).

17	Bassett, D. S., Brown, J. A., Deshpande, V., Carlson, J. M. & Grafton, S. T. Conserved and variable architecture of human white matter connectivity. *NeuroImage* **54**, 1262-1279, doi:<https://doi.org/10.1016/j.neuroimage.2010.09.006> (2011).

18	Link, A. *et al.* Fibroblastic reticular cells in lymph nodes regulate the homeostasis of naive T cells. *Nature Immunology* **8**, 1255-1265, doi:10.1038/ni1513 (2007).

19	Klein, U. *et al.* Transcriptional analysis of the B cell germinal center reaction. *Proceedings of the National Academy of Sciences of the United States of America* **100**, 2639-2644, doi:10.1073/pnas.0437996100 (2003).

20	Holmes, A. B. *et al.* Single-cell analysis of germinal-center B cells informs on lymphoma cell of origin and outcome. *J Exp Med* **217**, doi:10.1084/jem.20200483 (2020).

21	Medaglia, C. *et al.* Spatial reconstruction of immune niches by combining photoactivatable reporters and scRNA-seq. *Science* **358**, 1622-1626, doi:10.1126/science.aao4277 (2017).

22	Palm, A. E. & Henry, C. Remembrance of Things Past: Long-Term B Cell Memory After Infection and Vaccination. *Front Immunol* **10**, 1787, doi:10.3389/fimmu.2019.01787 (2019).

23	Cumpelik, A. *et al.* Dynamic regulation of B cell complement signaling is integral to germinal center responses. *Nature Immunology* **22**, 757-768, doi:10.1038/s41590-021-00926-0 (2021).

24	Merkenschlager, J. *et al.* Dynamic regulation of TFH selection during the germinal centre reaction. *Nature* **591**, 458-463, doi:10.1038/s41586-021-03187-x (2021).

25	Park, C. S. & Choi, Y. S. How do follicular dendritic cells interact intimately with B cells in the germinal centre? *Immunology* **114**, 2-10, doi:10.1111/j.1365-2567.2004.02075.x (2005).

26	Henderson, Jacob G., Opejin, A., Jones, A., Gross, C. & Hawiger, D. CD5 Instructs Extrathymic Regulatory T Cell Development in Response to Self and Tolerizing Antigens. *Immunity* **42**, 471-483, doi:<https://doi.org/10.1016/j.immuni.2015.02.010> (2015).

27	Ramiscal, R. R. & Vinuesa, C. G. T-cell subsets in the germinal center. *Immunol Rev* **252**, 146-155, doi:10.1111/imr.12031 (2013).

28	Krummel, M. F., Bartumeus, F. & Gérard, A. T cell migration, search strategies and mechanisms. *Nature Reviews Immunology* **16**, 193-201, doi:10.1038/nri.2015.16 (2016).

29	Liu, Y. *et al.* High-plex protein and whole transcriptome co-mapping at cellular resolution with spatial CITE-seq. *Nature Biotechnology*, doi:10.1038/s41587-023-01676-0 (2023).

30	Zou, F. *et al.* Expression and Function of Tetraspanins and Their Interacting Partners in B Cells. *Frontiers in Immunology* **9**, doi:10.3389/fimmu.2018.01606 (2018).

31	Chen, Z., Koralov, S. B. & Kelsoe, G. Regulation of humoral immune responses by CD21/CD35. *Immunol Rev* **176**, 194-204, doi:10.1034/j.1600-065x.2000.00603.x (2000).

32	Roozendaal, R. & Carroll, M. C. Complement receptors CD21 and CD35 in humoral immunity. *Immunological Reviews* **219**, 157-166, doi:<https://doi.org/10.1111/j.1600-065X.2007.00556.x> (2007).

33	Johansson-Lindbom, B., Ingvarsson, S. & Borrebaeck, C. A. Germinal centers regulate human Th2 development. *J Immunol* **171**, 1657-1666, doi:10.4049/jimmunol.171.4.1657 (2003).

34	Allen, C. D., Okada, T. & Cyster, J. G. Germinal-center organization and cellular dynamics. *Immunity* **27**, 190-202, doi:10.1016/j.immuni.2007.07.009 (2007).

35	Natkunam, Y. The Biology of the Germinal Center. *Hematology* **2007**, 210-215, doi:10.1182/asheducation-2007.1.210 (2007).

36	Stebegg, M. *et al.* Regulation of the germinal center response. *Frontiers in immunology*, 2469 (2018).
