## Supplementary material for "Spatial omics representation and functional tissue module inference using graph Fourier transform": Terminology Box 1

**Terminology glossary used in SpaGFT**

**Length-scale** demonstrates the scaler of an observed tissue. Typically, a small length-scale structure can be captured using single-molecule resolution technology (e.g., MERFISH); and a large length-scale structure can be observed by cell/multiple-cells-level resolution technology (e.g., Visium and slide-seqV2)^1^.

**Tissue architecture** is defined as an organization of a specific tissue by describing the nature and the integrity of its cellular and extracellular compartments^2^.

A **spatial domain** is a segment of tissue architecture in spatially resolved transcriptomics (SRT) studies. A spatial domain is defined as a continuous region that is spatially coherent in both gene expression and histology^3^.

A **spatially variable gene (SVG)** represents a variable gene expression pattern, which varies along with the spatial distribution of tissue of interest^4^. Notably, SVG is not necessarily highly correlated to a spatial domain. Hence it may be distributed within a spatial domain or across multiple spatial domains.

A **tissue module (TM)** executes the specific biological function(s) within one spatial domain or across multiple spatial domains (e.g., germinal center in the lymph node). In SRT studies, a TM can be defined as a region or a set of discontinuous regions that are associated with a group of SVGs sharing similar spatial distribution.

A **tissue motif** is a conserved collection of TMs across multiple samples (e.g., the germinal center-B follicle-T cell zone of human lymph node samples). In a **tissue motif instance** of a specific sample, TMs are usually assembled by a specific tissue orchestration rule (e.g., spatial distance and cell-cell communication associated molecules) to form a complex tissue structure.

**Fourier mode (FM).** Given a KNN graph with *n* nodes (e.g., spots are nodes in SRT data and edges are added based on the Euclidean distance), a FM is a *n*-dimensional vector corresponding to an eigenvector of the Laplacian matrix of the graph. The **frequency** of a FM is defined as the corresponding eigenvalue. A FM is defined as a **low-frequency FM** if its corresponding eigenvalue is a small value and it represents a smooth and slow variation pattern (**Fig. IA**) on the graph. Similarly, a FM is defined as a **high-frequency FM** if its corresponding eigenvalue is a large value and it represents a rapid variation pattern on the graph (**Fig. IB**). The number of FMs is equal to the number of nodes in the graph, and FMs are ranked in increasing order of frequency. Notably, each FM is determined by the graph topology and does not depend on gene expression value.

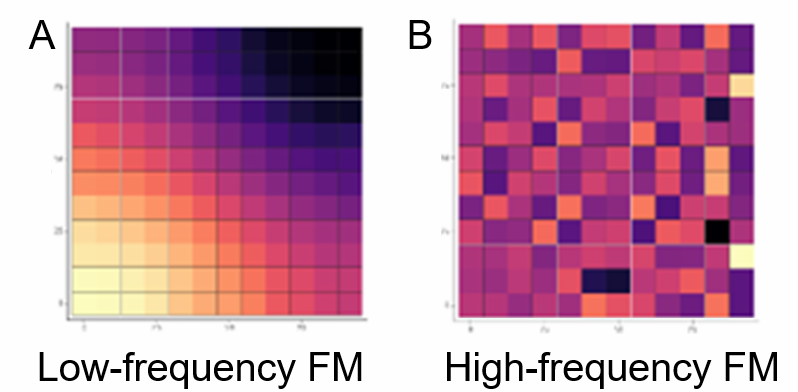

**Figure I**. A and B show Low-frequency FM and high-frequency FM.

**Graph Fourier transform (GFT).** In SRT studies, given the expression values of a gene on $n$ spots $f\in R^{n}$ on a KNN graph, the mathematical process of transforming $f$ to $\hat{f}$by equation $\hat{f}=U^{T}f$ is called **GFT**, where $U$ is the matrix consists of column vectors $u_{1},u_{2},\ldots,u_{n}$ ($u_{i}$ is the $i^{th}$FM). Each component of $\hat{f}$ is defined as a **Fourier coefficient (FC)**, and the $i^{th}$element of $\hat{f}$ is the FC of $i^{th}$FM. Hence, expression values of a specific gene on the KNN graph can be decomposed into a linear combination of FMs, where a FC can be used for measuring the contribution of the corresponding FM. **Inverse GFT (iGFT)** is the mathematical process of transforming $\hat{f}$ to $f$by the function $f=U^{T}\hat{f}$.

**GFTscore** of a specific gene is defined as $GFTscore\left( f \right)=\sum_{l=1}^{n} e^{-\lambda_{l}}\hat{f_{i}}$, where $\lambda_{l}$ is the $l^{th}$ frequency and $\hat{f_{i}}$ is the normalized $l^{th}$ FC.

1	Palla, G., Fischer, D. S., Regev, A. & Theis, F. J. Spatial components of molecular tissue biology. *Nature Biotechnology*, doi:10.1038/s41587-021-01182-1 (2022).

2	Hagios, C., Lochter, A. & Bissell, M. J. Tissue architecture: the ultimate regulator of epithelial function? *Philos Trans R Soc Lond B Biol Sci* **353**, 857-870, doi:10.1098/rstb.1998.0250 (1998).

3	Hu, J. *et al.* SpaGCN: Integrating gene expression, spatial location and histology to identify spatial domains and spatially variable genes by graph convolutional network. *Nature Methods*, doi:10.1038/s41592-021-01255-8 (2021).

4	Li, K. *et al.* Computational elucidation of spatial gene expression variation from spatially resolved transcriptomics data. *Molecular Therapy - Nucleic Acids* **27**, 404-411, doi:<https://doi.org/10.1016/j.omtn.2021.12.009> (2022).
