## Supplementary material for "Spatial omics representation and functional tissue module inference using graph Fourier transform": Terminology Box 2

**Terminology comparison between spatial domain and tissue module**

| **Terminology** | **Spatial domain** | **Tissue module (TM)** |
| --- | --- | --- |
| **Literature description** | A spatial domain is a segment of tissue architecture in spatially resolved transcriptomics (SRT) studies. A spatial domain is defined as a continuous region that is spatially coherent in both gene expression and histology^1^. | View of TM cell composition: A TM is a conserved region formed from cell types (e.g., immune and tumor) that is co-regulated by recruitment factors, such as cytokines^2^.  View of TM functions: A TM refers to a conserved functional niche across multiple samples and consists of recurrent cellular communities to execute the corresponding functions^3^. |
| **Examples** | Layer 1 is one of the layers in the human brain that serves as the locus of memory formation and storage in the neocortex^4^. | The germinal center in the lymph node is a typical TM for B cell maturation^3, 5^. |
| **Illustration** | 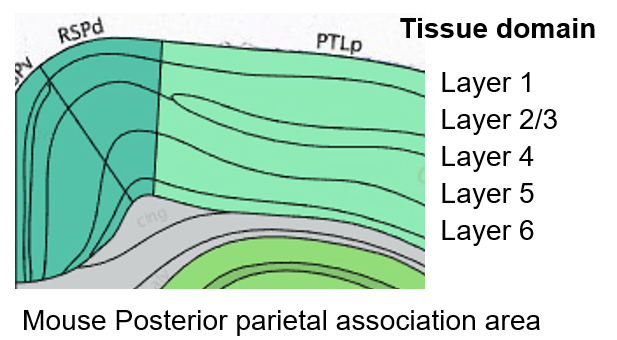  Generate from Allen Brain Atlas | 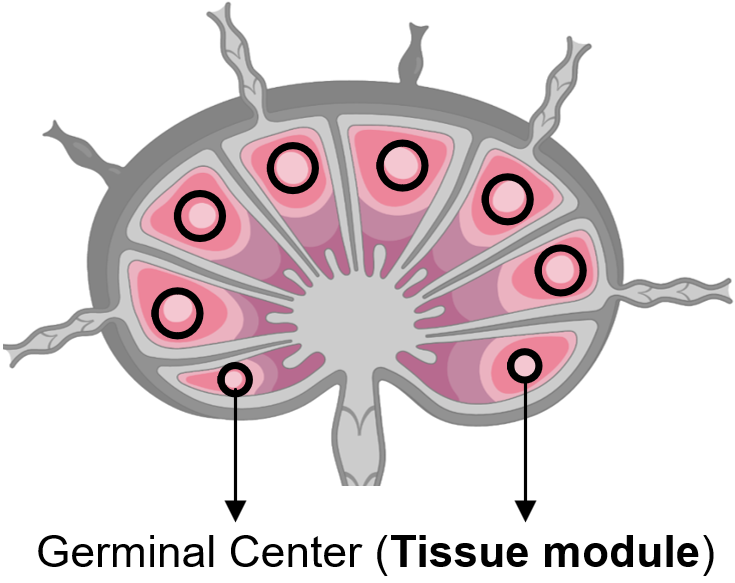  Generate from BioRender |
| **Difference** | (*i*) One spatial domain is a segment of tissue architecture and represents a substructural case of tissue architecture.  (*ii*) It requires spatial continuity.  (iii) SVG is not necessarily associated with a spatial domain. | (i) TMs vary in terms of their length-scale and cell composition. (ii) It may be within one spatial domain or across multiple spatial domains. (iii) A TM is not necessary to require spatial continuity. (iv) A group of SVGs is required to identify and elucidate TMs in SRT studies. (v) TMs may be adjacent or overlapped to form a high-order functional region, such as signal propagation, the cellular movement followed by secretion, or cell-cell signaling^6^. |
| **Applicable technologies** | Sequencing-based and  image-based spatial transcriptomics | Sequencing-based and  image-based spatial transcriptomics |
| **Computational formulation** | It can be formulated as spot clustering by measuring the similarity of the spot features, including but not limited to genes, spot distance, and histology. | (*i*) It can be formulated as tensor factorization based on spatial proteomics data across multiple samples^2^.  (ii) It can be formulated as the set covering problem by finding a set of TMs to cover the convoluted biological functions in an entire tissue (in our study). |
| **Computational framework** | **SpaGCN** is a graph convolutional network-based approach that considers both spatial location and histology information in clustering  **STAGATE^7^** used a graph attention auto-encoder framework STAGATE to accurately identify spatial domains by learning low-dimensional latent embeddings via integrating spatial information and gene expression profiles | **Tensor decomposition** uses tensor decomposition on a 3D tensor consisting of cell neighbor (cell adjacency graph), cell types, and patients to jointly identify cell neighbor-cell type compositions in each patient group separately^2^.  **SpaGFT** contains three steps. (*i*) It identifies SVGs using GFT; (*ii*) It uses the Louvain algorithm to cluster SVG using low-frequency FC, and each SVG cluster corresponds to a TM. (*iii*) An optimization method is used to optimize the resolution of the Louvain algorithm by minimizing the overlapped spots of identified TMs. |
| **high-ordered functional structure** | Continuous spatial domains will form tissue architecture, which is an organization of a specific tissue by describing the nature and the integrity of its cellular and extracellular compartments^8^. | A conserved collection of TMs across multiple samples will form a tissue motif (e.g., the germinal center-B follicle-T cell zone of human lymph node samples)^6^. |
| **Illustration** | 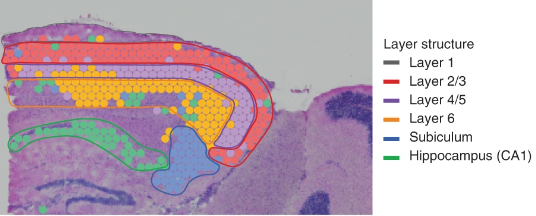  Figure from paper^1^.  SpaGCN results are used to demonstrate how the spatial domain forms the spatial architecture. The figure demonstrated identified six spatial domains from the mouse cortical region, including Layer 1, Layer 2/3, Layer 4/5, Layer 6, subiculum, and hippocampus. | 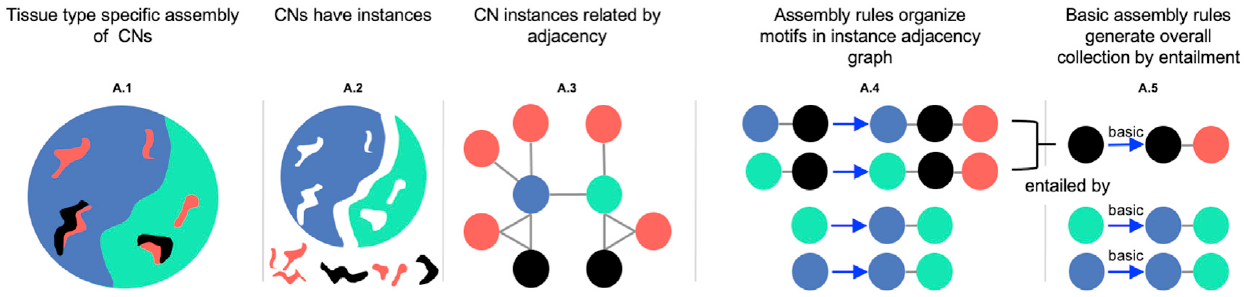  A.1 A.2 A.3 A.4  Figure from paper^6^.  (A.1) Tissues have cell neighbors (CN, defined as cell type composition in one region) assembled in different arrangements.  (A.2) CNs have connected components referred to as "instances."  (A.3) Instances in tissue are related by spatial adjacency forming a graph.  (A.4) Motifs in the graph are repeated colored subgraphs, and architectural rules correspond to a motif always found as part of a larger one. |

1.	Hu, J. et al. SpaGCN: Integrating gene expression, spatial location and histology to identify spatial domains and spatially variable genes by graph convolutional network. *Nature Methods* (2021).

2.	Schürch, C.M. et al. Coordinated Cellular Neighborhoods Orchestrate Antitumoral Immunity at the Colorectal Cancer Invasive Front. *Cell* **182**, 1341-1359.e1319 (2020).

3.	Palla, G., Fischer, D.S., Regev, A. & Theis, F.J. Spatial components of molecular tissue biology. *Nature Biotechnology* (2022).

4.	Shin, J.N., Doron, G. & Larkum, M.E. Memories off the top of your head. *Science* **374**, 538-539 (2021).

5.	Gars, E., Butzmann, A., Ohgami, R., Balakrishna, J.P. & O'Malley, D.P. The life and death of the germinal center. *Ann Diagn Pathol* **44**, 151421 (2020).

6.	Bhate, S.S., Barlow, G.L., Schürch, C.M. & Nolan, G.P. Tissue schematics map the specialization of immune tissue motifs and their appropriation by tumors. *Cell Systems* **13**, 109-130.e106 (2022).

7.	Dong, K. & Zhang, S. Deciphering spatial domains from spatially resolved transcriptomics with an adaptive graph attention auto-encoder. *Nature Communications* **13**, 1739 (2022).

8.	Hagios, C., Lochter, A. & Bissell, M.J. Tissue architecture: the ultimate regulator of epithelial function? *Philos Trans R Soc Lond B Biol Sci* **353**, 857-870 (1998).
